# Genome-Wide Discovery and Characterization of Terpene Synthases Contributing to Strawberry Aroma Metabolism

**DOI:** 10.1101/2025.06.03.654341

**Authors:** Mary A. Madera, Kristen L. Becker, Jenevieve D. Weissman, Randi A. Famula, Marta Bjornson, Glenn S. Cole, Mitchell J. Feldmann, Steven J. Knapp, Philipp Zerbe

**Author notes:** To whom correspondence may be addressed: Philipp Zerbe; Department of Plant Biology, University of California-Davis, One Shields Avenue, Davis, CA 95616, United States. **Author’s email addresses**.

## Abstract

Strawberry (*Fragaria* X *ananassa*) is a globally cultivated fruit crop valued for its unique flavor and aroma. Major groups of volatile organic compounds that contribute to strawberry aroma include terpenes, fatty acid esters, furanones, and benzenoids. Understanding the biosynthetic genes and pathways underlying strawberry terpene metabolism provides resources for developing precision breeding strategies to improve strawberry aroma traits. Here we describe the genome-wide identification and functional analysis of the terpene synthase (TPS) family that governs terpene chemical diversity in strawberries. Mining of the allo-octoploid genome of the cultivar ‘Royal Royce’ identified 75 TPS gene candidates. Biochemical characterization of 27 predicted mono-, sesqui– and di-TPS enzymes via *in vitro* activity assays and *in vivo* enzyme co-expression studies demonstrated a range of TPS activities that are relevant for the biosynthesis of two thirds of known strawberry terpenes as well as products not previously described in octoploid strawberry. Complementary metabolomic and transcriptomic studies across a diversity panel of strawberry accessions demonstrated substantial variation in the composition and abundance of more than 30 detected accession-specific and common terpene aromas, nine of which matching characterized TPS products. Analysis of a developmental time series of two accessions with contrasting aroma further revealed accession-specific alterations of terpene metabolism during fruit ripening. These results expand our understanding of strawberry aroma metabolism and characterized terpene aroma genes as a resource for the molecular breeding of desirable aroma traits.

**One-sentence summary:** Discovery and characterization of the terpene synthase family that controls the diversity of terpene aroma metabolism in accessions of strawberry.

**Significance Statement:** The diverse class of terpenoids contributes to complex crop traits, including plant growth, stress resilience, pollination, nutrition, and aroma. The discovery of terpene-biosynthetic pathways in octoploid strawberry (*Fragaria* X *ananassa*) provides foundational knowledge of the diversity of terpenoid metabolism in this important specialty crop toward applications in improving aroma traits.

## Introduction

Cultivated strawberry (*Fragaria ananassa* Duchesne ex Rozier*, Fa*) originated in Europe in the early 18^th^ century (Darrow, 1966; Hardigan et al., 2021b). Over the past 300 years, breeding efforts have improved many crop traits, making strawberry a popular fruit crop among consumers worldwide (Darrow, 1966; Ulrich and Olbricht, 2016; Yan et al., 2018). Global strawberry production reached over 14 M tons in 2023 with Asia, the Americas, and Europe being the primary producers (FAO, 2000). Cultivated strawberry is an allo-octoploid species that derived from the spontaneous hybridization of North American *F. virginiana* (*Fvr*) and South American *F. chiloensis* (*Fc*) (Fig. 1) (Darrow, 1966; Hardigan et al., 2021b). These octoploid progenitors evolved approximately a million years ago through several polyploidization events involving four diploids, *F. vesca (Fv), F. innumae,* and two other currently unidentified diploid sources (Njuguna et al., 2013; Tennessen et al., 2014; Sargent et al., 2016; Session and Rokhsar, 2023).

**Fig. 1.**
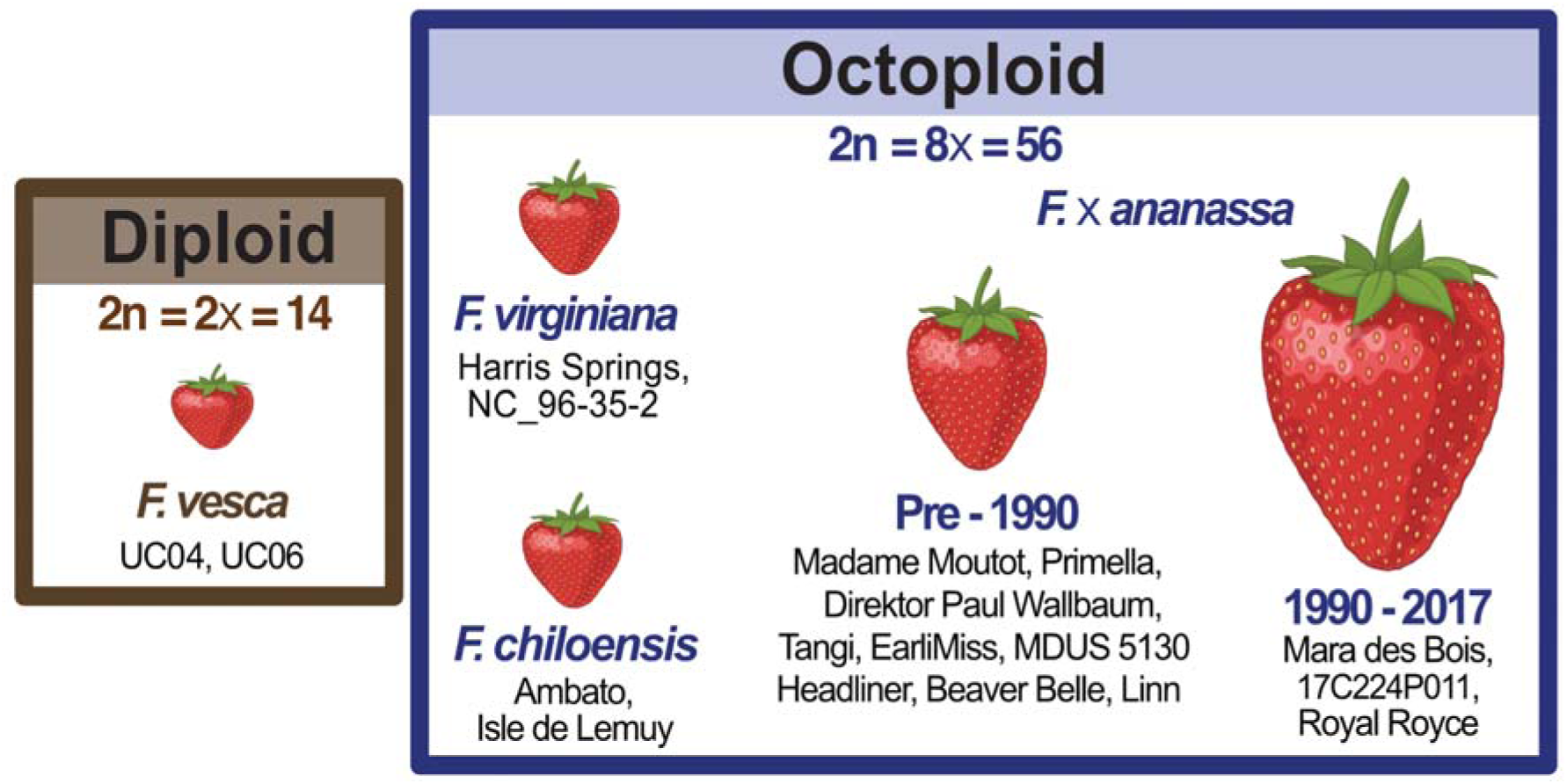
Schematic overview of strawberry (*Fragaria spp.*) accessions used in this study. Two diploid *F. vesca*, two octoploid *F. virginiana*, and two octoploid *F. chiloensis* accessions were used along with nine *F.* × *ananassa* cultivars bred before 1990 and 3 cultivars bred after 1990.

Strawberry breeding has greatly enhanced key horticultural traits, including fruit yield, berry size, firmness, and shelf life, and has largely expanded the range of available flavor and other quality traits (Feldmann et al., 2024). Closely associated with flavor that is largely driven by sugar and organic acid levels of the fruit (the fleshy receptacle), aroma is an important multifactorial trait that varies substantially across strawberry cultivars and is impacted by genotype, environment, and cultivation practices (Schwieterman et al., 2014; Ulrich and Olbricht, 2016; Fan et al., 2021b; Porter et al., 2023). Strawberry aroma is characterized by complex bouquets of over 300 identified volatile organic compounds (VOCs) that include fruity esters, caramellic furans, peachy lactones, green aldehydes, and floral or fruity terpenes (Pyysalo et al., 1979; Bood and Zabetakis, 2002; Schwieterman et al., 2014; Fan et al., 2021b). Among these VOCs, the large class of terpene metabolites has been an important target for aroma breeding (Schwieterman et al., 2014; Ulrich and Olbricht, 2016; Fan et al., 2021b). Over 25 volatile monoterpenes (C_10_) and sesquiterpenes (C_15_) have been reported in strawberry aroma profiles (Zabetakis and Holden, 1997; Negri et al., 2015; Urrutia et al., 2017; Duan et al., 2018; Yan et al., 2018). Among these linalool, terpineol, and nerolidol strongly contribute to citrus and floral aroma notes that are positively associated with sensory quality and flavor intensity (Schwieterman et al., 2014; Ulrich and Olbricht, 2016; Fan et al., 2021b). Commercial strawberry hybrids are typically dominated by linalool, nerolidol, and □-terpineol with fruity, floral, and jasmine-like aroma notes. By contrast, wild octoploid and diploid strawberry accessions often feature higher abundance of terpenes with pungent floral, woody, or minty aromas, including □– and li:-pinene, limonene, citronellol, myrtenol, myrtenyl acetate, and li:-phellandrene (Aharoni et al., 2004; Bianchi et al., 2014; Negri et al., 2015; Ulrich and Olbricht, 2016; Urrutia et al., 2017; Duan et al., 2018).

Long-standing research efforts have provided insights into the biosynthesis of aroma-related VOCs and advanced the breeding of strawberry aroma profiles (Aharoni et al., 2004; Beekwilder et al., 2004; Raab et al., 2006; Chambers et al., 2012; Cumplido-Laso et al., 2012; Zorrilla-Fontanesi et al., 2012; Cruz-Rus et al., 2017; Pillet et al., 2017; Barbey et al., 2021; Oh et al., 2021; Fan et al., 2021b; Fan et al., 2022; Porter et al., 2023). The chemical diversity of terpene VOCs is governed by the activity of terpene synthase (TPS) enzymes that convert a few conserved prenyl diphosphate precursors through complex, electrophilic cyclization and carbon-carbon rearrangement reactions (Chen et al., 2011; Karunanithi and Zerbe, 2019). These diverse TPS functions will have evolved through repeated events of gene and genome duplications and subsequent sub– and neo-functionalization (Zi et al., 2014; Karunanithi and Zerbe, 2019).

Therefore, it can be hypothesized that the polyploidization events that led to modern octoploid strawberry have resulted in an expansive TPS gene family (Njuguna et al., 2013; Tennessen et al., 2014; Sargent et al., 2016; Edger et al., 2019; Session and Rokhsar, 2023). Despite their importance for fruit aroma, our knowledge of strawberry terpene metabolism has remained incomplete. Known TPS functions include three functional nerolidol synthases (NESs), namely *F. vesca FvNES1* and *F.* × *ananassa FaNES1* and *FaNES2*, which feature a dual functionality, producing both the sesquiterpene nerolidol from farnesyl diphosphate (FPP) and the monoterpene linalool from geranyl diphosphate (GPP), respectively (Aharoni et al., 2004; Fan et al., 2022). However, gene expression of only *FaNES1* was shown to be high enough to quantitatively contribute to strawberry aroma. Although displaying lower transcript abundance, the near-identical *FaNES1t* gene has been correlated with a broader product range, including □-terpineol, li:-myrcene, and (*E*)-li:-farnesene (Fan et al., 2022). In addition, *F. vesca* pinene synthase, *FvPINS*, was identified as a monoterpene synthase, producing □-pinene, li:-phellandrene, and li:-myrcene found to be abundant in wild, ripe *F. vesca* berries (Aharoni et al., 2004), and a predicted germacrene D synthase, *FaTPS1,* was identified in pathogen-stressed strawberry fruits (Zhang et al., 2022).

To expand our understanding of terpene metabolism in octoploid strawberry, we here report on the identification of the strawberry *TPS* gene family. Biochemical characterization of 27 TPSs revealed common and accession-specific enzyme products beyond the known NES and PINS enzymes. Combined transcriptomics and metabolomics analyses across a diverse panel of wild and domesticated accessions highlight that differences in TPS functional specificity and gene family composition contribute to variations in terpene aroma profiles in strawberry.

## Results

### Discovery of the terpene synthase gene family in allo-octoploid strawberry

To identify the terpene-metabolic network in octoploid strawberry, we mined a haplotype-phased assembly of the *F.* × *ananassa* genome for the cultivar ‘Royal Royce’ (FaRR1; https://phytozome-next.jgi.doe.gov/info/FxananassaRoyalRoyce_v1_0; (Hardigan et al., 2021a)) for *TPS* gene candidates and genes of the upstream mevalonate (MVA) and methylerythritol phosphate (MEP) pathways that provide the conserved FPP and GPP precursors. BLAST searches against curated protein databases (Zerbe et al., 2013; Tiedge et al., 2020) identified a total of 62 candidates with best matches to MEP and MVA pathway genes, including one or more members of each gene family involved in these pathways (Supplemental Fig. 1, Supplemental Table 1). With the exception of *4-hydroxy-3-methylbut-2-enyl diphosphate synthase* (*HDS*), for which a homologous gene c opy was absent on FaRR1 chromosome 3C, all identified MEP and MVA pathway gene candidates were represented by four syntenic copies in the reference genome (Supplemental Figs. 2-3, Supplemental Table 2). Notably, eight copies of candidate genes were identified for *acetoacetyl-CoA thiolase* (*AACT*) (Supplemental Fig. 2), *3-hydroxy-3-methylglutaryl-CoA reductase* (*HMGR*) (Supplemental Fig. 2D-E), *FPP synthases* and *GPP* or geranylgeranyl diphosphate (*GGPP*) *synthases,* appearing in two groups of four syntenic genes each (Supplemental Figs. 2-4).

Next, BLAST searches of the FaRR1 genome against a TPS protein database (Zerbe et al., 2013) identified 104 predicted *TPS* genes, 75 of which represented full-length transcripts (Fig. 2, Supplemental Tables 3-4). Based on the presence of characteristic DDxxD, DDxD, and RRx_8_W sequence motifs, these 75 TPS candidates were classified as mono-or sesqui-TPSs (*FaTPS1-64*), class II diTPSs (copalyl diphosphate synthase [CPS], *FaCPS1-7*), or class I diTPSs (kaurene synthase-like [KSL], *FaKSL1-4*) (Fig. 2). This included the previously characterized NESs, *FaNES1* (*FaTPS17*), *FaNES1t* (*FaTPS18*), and *FaNES2* (*FaTPS11*) (Aharoni et al., 2004; Fan et al., 2022), and the predicted germacrene D synthase *FaTPS1* (*FaTPS26*) (Zhang et al., 2022). To improve consistency in TPS nomenclature, we here adopted the nomenclature suggested for maize and other species (Köllner et al., 2023) and designated the identified TPS genes as *FaTPS1-64*, *FaCPS1-*7, and *FaKSL1-4*. Accordingly, the previously characterized TPSs were renamed as *FaTPS17/NES1*, *FaTPS18/NES1t*, *FaTPS11/NES2*, *FvTPS50/PINS*, and *FaTPS26/TPS1*. Notably, *FaTPS17/NES1* and *FaTPS18/NES1t* are assigned to the same gene locus, *Fxa3Cg100266.1*, consistent with previous findings (Fan et al., 2022). Additionally, six other *TPS* genes were present in the FaRR1 genome as sequences substantially longer than known *TPS* genes and likely represent misassembled genes (Supplemental Fig. 5).

**Fig. 2.**
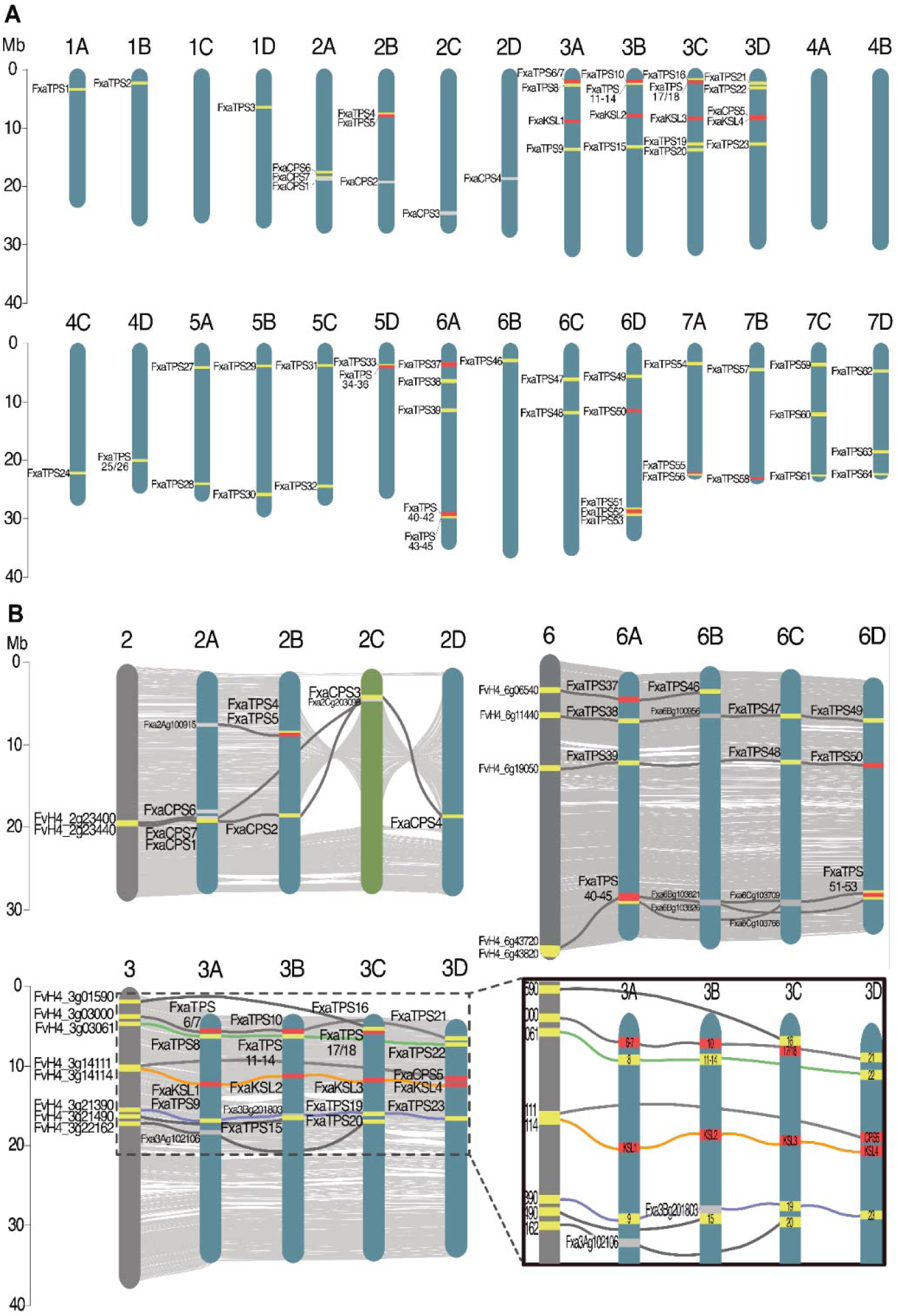
Genomic localization of identified terpene synthase (TPS) genes in the strawberry genome. (**A**) Relative location of functionally characterized (red) and predicted (yellow) full-length TPS genes in the *Fragaria* × *ananassa* cultivar ‘Royal Royce’ (FaRR1, (2021) genome. Abbreviations: CPS, copalyl diphosphate synthase; KSL, kaurene synthase-like. (**B**) Synteny plots of FaRR1 chromosomes 2 (top left), 6 (top right), and 3 (bottom) with TPS genes identified in the diploid *F. vesca* genome. Gray lines show syntenic relationships between TPS genes. Syntenic pseudogenes are shown (gray). Chromosome 2C shown in green to depict whole chromosome inversion. Gene-dense region of chromosome 3 shown in detail; colored lines depict synteny matches across all 5 chromosomes.

**Fig. 3.**
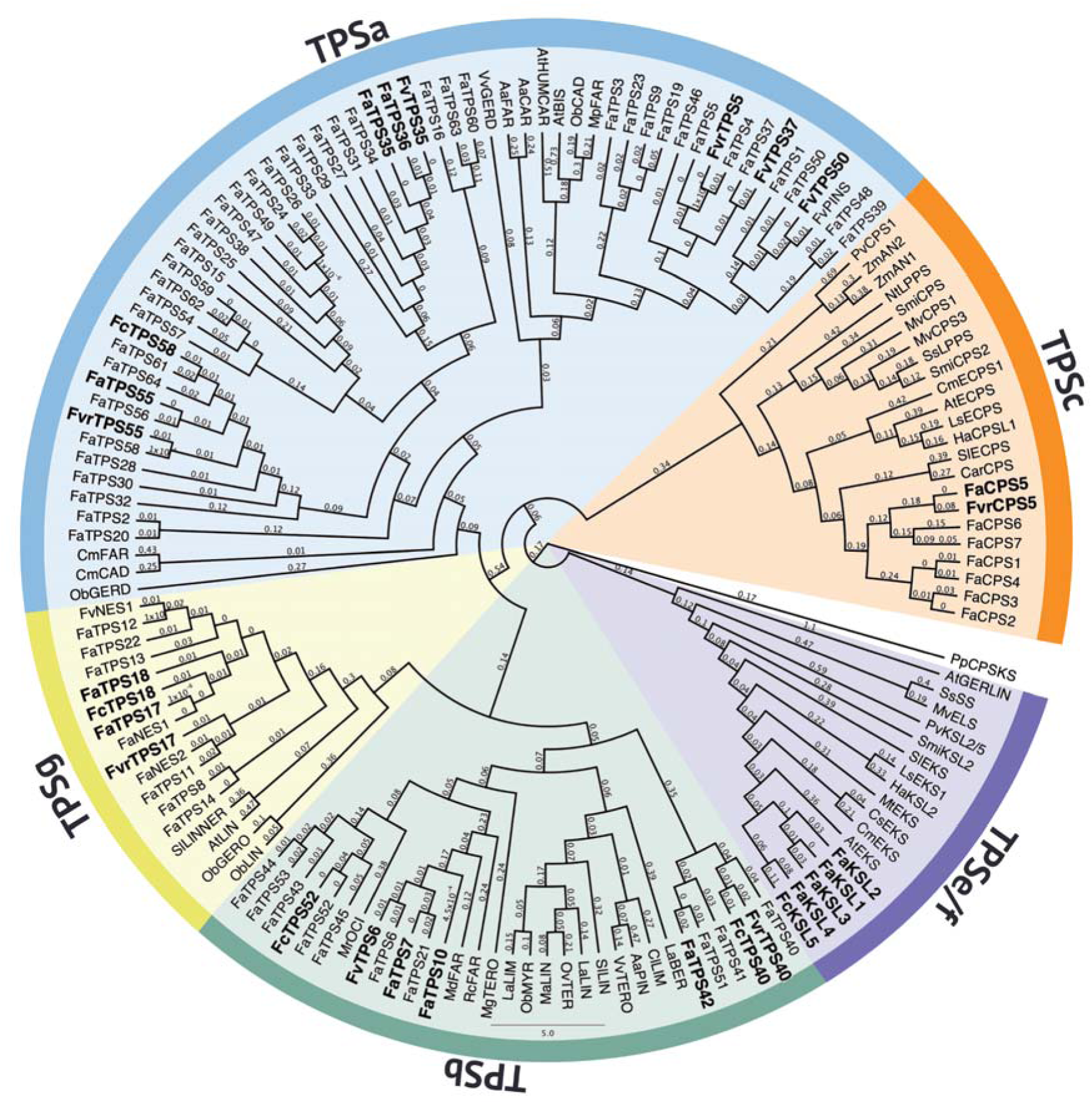
Maximum-likelihood phylogeny of strawberry terpene synthases (TPSs) and selected TPSs of the TPS-c, TPS-e/f, TPS-g, TPS-b and TPS-a subfamilies from related species. Tree rooted with the *ent*-CPP/*ent*-kaurene synthase from *Physcomitrella patens* (*PpCPSKS*). Bootstrap support values (500 repetitions) are depicted. Enzyme candidates biochemically characterized in this study are highlighted in bold. Accession numbers and protein sequences are given in Supplemental Table 1.

**Fig. 4.**
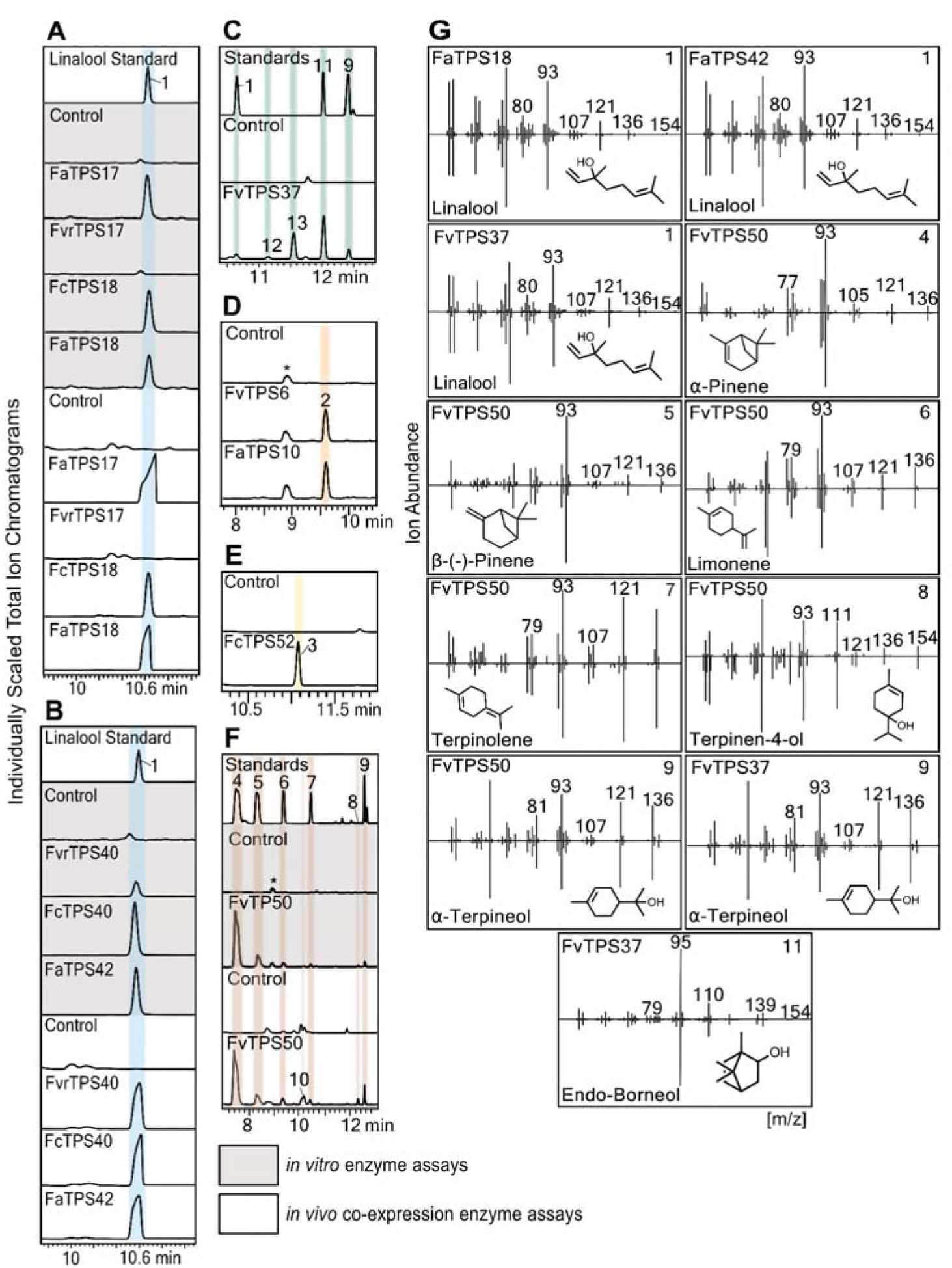
Functional characterization of monoterpene synthases (TPSs). (**A-F**) GC-MS traces of products resulting from either *in vitro* enzyme assays of individual recombinant TPSs with geranyl diphosphate (GPP) as a substrate (gray) or co-expression assays of individual TPSs and a GPP synthase in *E. coli* (white). (**G**) Mass spectra of enzyme products identified by comparison to authentic standards (white) or mass spectral databases (NIST, v17.1). FaTPS17, TPS18, and TPS42 taken from *F.* × *ananassa (Fa)* cultivar ‘Royal Royce’; FvrTPS17 *F. virginiana* (*Fvr*) accession ‘NC_96-35-2’; FcTPS18 *F. chiloensis* (*Fc*) ecotype ‘Ambato’; FvrTPS40 accession ‘Harris Springs’; FcTPS40 and FcTPS52 ecotype ‘Isle de Lemuy’; FvTPS6 and FvTPS50 diploid accession ‘UC06’; FaTPS10 cultivar ‘EarliMiss’.

The identified *TPS* genes were widely distributed across the FaRR1 genome with most genes located on chromosome 3 and 6 (Fig. 2). No predicted terpene-metabolic gene clusters were identified; however, several chromosomes featured closely co-localized TPS genes. Gene synteny analysis of the identified TPSs using the diploid *F. vesca* genome for accession ‘Hawaii-4’ (Edger et al., 2018) and the octoploid FaRR1 genome illustrated that 20 TPSs were present with four homoeologous copies across each FaRR1 subgenome and in the *F. vesca* genome, forming five syntenic groups with *F. vesca* genes *FvH4_3g23440* on chromosome 2, *FvH4_3g14116* on chromosome 3, *FvH4_5g06470* on chromosome 5, and *FvH4_7g03260* and *FvH4_7g33640* on chromosome 7 (Supplemental Fig. 6). The remaining *TPS* genes lacked homoeologous copies on at least one of the FaRR1 subgenomes and were present mostly in syntenic groups of two or three copies. This included *FvTPS50/PINS*, *FaTPS17/NES1*, *FaTPS18/NES1t* (Aharoni et al., 2004; Fan et al., 2022), and NES-like candidates found in the transcriptomes of *F. chiloensis* and *F. virginiana* (see below) and pinene synthase-like candidates from both *F. virginiana* and *F. vesca* (Supplemental Figs. 6-8). *FaTPS3-5, 11-15, 25, 33, 41-45, 53*, and *63* do not have syntenic equivalents in the diploid *F. vesca* genome, suggesting that they either evolved after polyploidization or originated from one of the other three diploid progenitor genomes (Supplemental Fig. 6; Supplemental Table 3).

To identify TPSs relevant for fruit aroma beyond one strawberry accession, we performed transcriptome analysis of greenhouse– and field-grown fruits of Royal Royce and 16 additional cultivars, ecotypes, or accessions selected to encompass a range of wild species and diverse breeding materials (Supplemental Table 5). RNAseq reads were assembled both *de novo* and aligned to the FaRR1 genome. Additionally, transcriptome sequencing was performed on Royal Royce and *F.* × *ananassa* cultivar ‘Mara des Bois’ fruits at green, white, underripe, ripe and overripe stages as described by Jiménez et al., 2024. Analysis of these transcriptome data identified 24 full-length transcripts with best matches to TPS genes. This included *FaTPS10, FaTPS55*, and *FaCPS5* with ≧95% sequence identity to the genes identified in the FaRR1 genome, as well as the individual transcripts for *FaTPS17/NES1* and *FaTPS18/NES1t* identified in the FaRR1 genome as a combined gene (*Fxa3Cg100266.1*). *FaTPS7* (*F.* × *ananassa* cultivar ‘Primella’; Prim) and *FaTPS36* (*F.* × *ananassa* cultivar ‘Direktor Paul Wallbaum’; DPW) showed lower sequence similarity (93-95%) to TPSs identified in the FaRR1 genome and may represent genes distinctive to these cultivars. Likewise, *FaTPS35* and *FaTPS42* identified in the Royal Royce transcriptome shared only >93% with the FaRR1 genomic TPS candidates and were tentatively considered distinct. The remaining 14 TPSs were identified in the transcriptomes of *F. chiloensis* (*Fc*)*, F. virginiana* (*Fvr*), and *F. vesca* (*Fv*). These TPSs were named according to the highest protein sequence similarity to TPSs identified in the FaRR1 genome: *F. chiloensis* ecotype ‘Ambato’ (Amb) *FcKSL5* and *FcTPS18*; *F. chiloensis* ecotype ‘Isle De Lemuy 02A White’ (ILE) *FcTPS40*, *FcTPS52*, and *FcTPS58*; *F. virginiana* accession ‘NC_96-35-2’ (NC) *FvrCPS5*, *FvrTPS17,* and *FvrTPS55*; *F. virginiana* accession ‘Harris Springs’ (HS) *FvrTPS5* and *FvrTPS40*; and *F. vesca* accession ‘UC06’ *FvTPS6*, *FvTPS35*, *FvTPS37*, and *FvTPS50* (Fig. 3, Supplemental Table 3 & 4). In summary, analysis of the FaRR1 genome and the obtained transcriptome data, we identified 76 predicted mono– and sesqui-TPSs and 13 class II and class I diTPSs with predicted functions in diterpenoid metabolism (Supplemental Table 4).I

Phylogenetic analysis of the identified TPS candidates was performed to substantiate functional annotations. Consistent with the presence of a βα-domain structure, a plastidial transit peptide, and a characteristic RRX_8_W motif (Chen et al., 2011; Karunanithi and Zerbe, 2019), 16 TPSs were placed together with known mono-TPSs of the TPS-b subfamily (Fig. 3). Sharing a βα-domain structure but lacking a transit peptide and RRX_8_W motif 50 TPSs clustered with members of the TPS-a1 subfamily of dicot sesqui-TPSs. Ten TPSs showed a close phylogenetic relationship with *FaNES1*, *FaNES2* and related enzymes of the TPS-g subfamily, suggesting similar catalytic functions.

In addition to mono– and sesqui-TPSs with predicted functions in strawberry aroma metabolism, 13 gene candidates were identified as predicted diTPSs. Of these *FaCPS1-7* and *FvrCPS5* (accession NC_96-35-2) were placed within the TPS-c subfamily and featured a DxDD motif characteristic for class II diTPSs (Zerbe and Bohlmann, 2015) (Fig. 3, Supplemental Table 3). With the exception of *FaCPS5* on chromosome 3D, all CPSs are localized on chromosome 2 homeologs and share >96% protein sequence similarity, suggesting a related function (Supplemental Fig. 7, Supplemental Table 6). Presence of a characteristic H-N dyad in the N-terminal catalytic domain of *FaCPS1-4* and *FaCPS6-7* suggested catalytic functions as diTPSs forming *ent*-CPP (Potter et al., 2014; Zi et al., 2014; Potter et al., 2016) (Supplemental Fig. 9). However, *FaCPS6* showed an N-terminal truncation, lacking part of the γ-domain likely rendering the protein inactive. *FaCPS5* and *FvrCPS5* shared only 70% sequence identity with the other CPSs and did not feature the H-N dyad or a His501 residue conserved in *syn*-CPP synthases (Potter et al., 2014; Zi et al., 2014; Potter et al., 2016), thus suggesting a distinct function (Supplemental Fig. 7 & 9). Lastly, five KSLs were identified that feature the conserved, catalytic DDxxD and NSE/DTE motifs and formed a separate group within the TPS-e/f subfamily of class I diTPSs (Fig. 3, Supplemental Fig. 9). *FaKSL1-3* co-localized on chromosome 3A-C, whereas *FaKSL4* was localized next to *FaCPS5* on chromosome 3D (Fig. 2, Supplemental Fig. 7). In addition, we identified *FcKSL5* in *F. chiloensis* ecotype Ambato. The *FcKSL5* gene sequence most closely matched *FaKSL4* in the FaRR1 genome. However, sequence analysis of *FaKSL4* indicated a possible misassembly rendering a gene candidate over three times the expected KSL length (Supplemental Fig. 5), preventing a clear analysis of the homology between these two diTPSs. *FcKSL5* adopts a βα-domain architecture known only in KSL enzymes involved in specialized diterpenoid metabolism (Zerbe and Bohlmann, 2015; Karunanithi and Zerbe, 2019) (Supplemental Fig. 9).

### Strawberry mono– and sesqui-terpene synthases produce known and previously unrecognized aroma metabolites

Based on predicted functions in terpene aroma metabolism, we selected 27 TPSs for biochemical characterization. Enzyme functional studies were conducted via *in vitro* enzyme assays using recombinant, purified proteins and commercial GPP and FPP substrates and, where further enzyme characterization was needed, *in vivo* co-expression assays using an *E. coli* platform engineered for terpene production (Cyr et al., 2007; Morrone et al., 2010; Murphy et al., 2019). Enzyme products were identified by comparison to authentic standards or, where so indicated, based on best matches to mass spectral databases.

GC-MS analysis of TPS enzyme products revealed catalytic activities for all tested TPSs with the exception of FvrTPS5 (accession Harris Springs) and FvrTPS17 (accession NC_96-35-2). *In vitro* and *in vivo* enzyme assays with GPP as a substrate verified linalool formation by FaTPS17/NES1 and FaTPS18/NES1t, and identified FaTPS42 (cultivar Royal Royce), FcTPS18 (ecotype Ambato), FvrTPS40 (accession Harris Springs), and FcTPS40 (ecotype Isle de Lemuy) as additional linalool synthases present in strawberry (Fig. 4). *In vitro* assays of FvTPS37 and FvTPS6 (accession UC06), FaTPS10 (cultivar ‘EarliMiss’, EM), and FcTPS52 (ecotype Isle de Lemuy) did not show detectable enzyme activity. However, co-expression studies of these TPSs with an *Abies grandis* GPP synthase identified FvTPS37 as a multi-product enzyme, forming □-terpineol, endo-borneol, and β-pinene hydroxide as major products with small amounts of linalool and predicted *trans*-2-pinanol as based on comparison to mass spectral databases (Fig. 4, Supplemental Fig. 10). In addition, FaTPS10 and FvTPS6 produced □-ocimene and FcTPS52 formed neo-allo-ocimene based on best database matches (Fig. 4, Supplemental Fig. 10). FvTPS50 (accession UC06) was identified *in vitro* and *in vivo* as a multi-product mono-TPS, forming □-pinene as a major product as well as small quantities of β-pinene, limonene, terpinolene, terpinen-4-ol, and □-terpineol.

*In vitro* and co-expression studies using FPP as a substrate showed that the identified linalool synthases, FaTPS17/NES1, FvrTPS17, FcTPS18, FaTPS18/NES1t, FvrTPS40, FcTPS40, and FaTPS42, also converted FPP to form *trans*-nerolidol (Fig. 5). This confirms the previously reported activities of FaTPS17/NES1 and FaTPS18/NES1t (Aharoni et al., 2004; Fan et al., 2022) and demonstrates that the strawberry genome contains additional dual function linalool/nerolidol synthases. *In vitro* enzyme assays identified FaTPS10, FaTPS7 (cultivar Primella) and FvTPS6 as □-farnesene synthases (Fig. 5). Additional *in vivo E. coli* co-expression assays of FaTPS10 and FvTPS6 with a maize FPP synthase further identified small quantities of 7-hydroxy-farnesene, □-*trans*-bergamotol, and □-sinensal as based on database references, which may represent additional TPS products or oxygenated by-products formed by endogenous *E. coli* enzymes (Fig. 5, Supplemental Fig. 11). Furthermore, FcTPS52 produced □-farnesene and Y-bisabolene as major products, as well as small amounts of □-*trans*-bergamotol and several other sesquiterpenes for which no significant mass spectral matches could be identified (Fig. 5, Supplemental Fig. 11). Protein yields generated for recombinant FvTPS35 (accession UC06), FaTPS35 (cultivar Royal Royce), and FaTPS36 (cultivar Direktor Paul Wallbaum) were consistently low. However, *in vitro* assays with FPP as a substrate resulted in the formation of β-elemene and humulene in addition to trace amounts of additional sesquiterpenes, likely including β-selinene and elemol (Fig. 5, Supplemental Fig. 12).

**Fig. 5.**
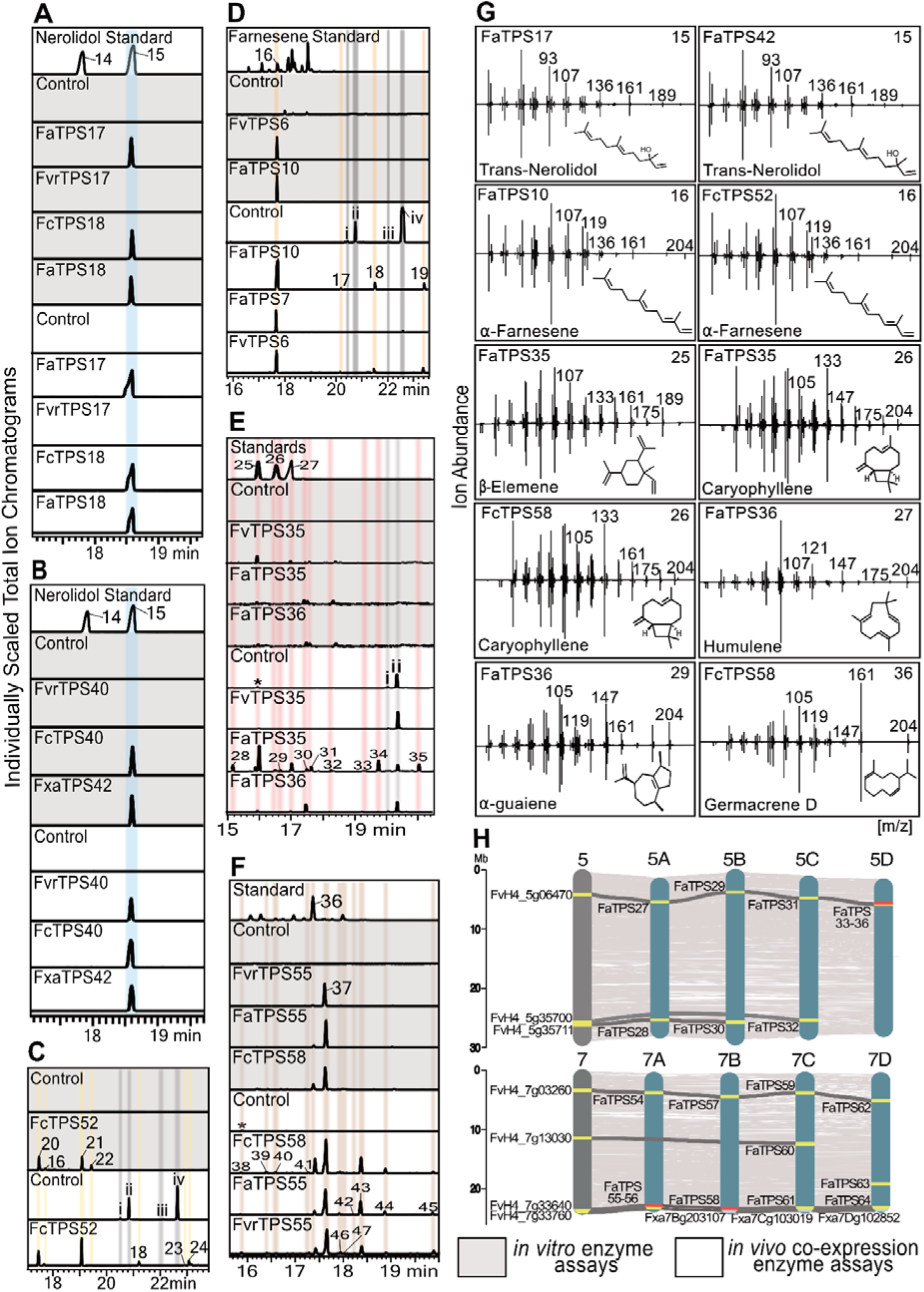
Functional characterization of sesquiterpene synthases (TPSs). (**A-F**) GC-MS traces of products resulting from either *in vitro* enzyme assays of individual recombinant TPSs with farnesyl diphosphate (FPP) as a substrate (gray) or co-expression assays of individual TPSs and a FPP synthase in *E. coli* (white). (**G**) Mass spectra of enzyme products identified by comparison to authentic standards (white) or mass spectral databases (NIST, v17.1) (purple). (**H)** Synteny plots of FaRR1 chromosomes 5 and 7 with *F. vesca.* The following compounds are likely degradation products of FPP produced in *E. coli* cultures: (i) *Cis-trans*-farnesene; (ii) Farnesol; (ii) 2,3-dihydro farnesyl acetate; (iv) *Trans*-farnesyl acetate. FaTPS17, TPS18, TPS35, and TPS42 taken from *F.* × *ananassa (Fa)* cultivar ‘Royal Royce’; FvrTPS17 and FvrTPS55 *F. virginiana* (*Fvr*) accession ‘NC_96-35-2’; FcTPS18 and FcTPS58 *F. chiloensis* (*Fc*) ecotype ‘Ambato’; FvrTPS40 accession ‘Harris Springs’; FcTPS40 and FcTPS52 ecotype ‘Isle de Lemuy’; FvTPS6, FvTPS35, and FvTPS50 *F. vesca* (*Fv*) accession ‘UC06’; FaTPS7 cultivar ‘Primella’; FaTPS10 cultivar ‘EarliMiss’; FaTPS55 cultivar ‘Madame Moutot’.

Additional co-expression assays confirmed the production of β-elemene and humulene by FaTPS35 alongside a range of additional sesquiterpenes, whereas co-expression of FaTPS36 showed β-selinene as the primary product. *In vitro* assays of FvrTPS55 (accession NC_96-35-2), FaTPS55 (cultivar ‘Madame Moutot’, MM), and FcTPS58 (ecotype Ambato) identified germacrene D and □-muurolene as major products as based on an authentic standard and mass spectral database matches, respectively (Fig. 5, Supplemental Fig. 12). Complementary co-expression assays confirmed these enzyme activities and showed additional lower abundant byproducts, including eremophilene, β-copaene, Y-muurolene, □-muurolol, epi-cubebol and shyobunol as based on mass spectral databases (Supplemental Fig. 12). Additional products that were present in trace amounts following TPS co-expression assays and control samples possibly represent isomerization or metabolism products of FPP, such as *cis*-*trans*-farnesene, farnesol, 2,3-dihydro farnesyl acetate, and *trans-*farnesyl acetate (Supplemental Figs. 11, 12).

### Pairwise activity of strawberry diterpene synthases produces specialized diterpenoids

Of the identified class II diTPSs, *FaCPS1-4* and *FaCPS7* featured a characteristic H-N dyad, supporting an *ent*-CPP synthase function with high probability and, therefore, were not biochemically analyzed in this study (Supplemental Fig. 9). Consistent with its functional prediction as a class II diTPS of specialized diterpenoid metabolism, *E. coli* co-expression of FaCPS5 or FvrCPS5 with a *Abies grandis* GGPP synthase resulted in the formation of *ent-neo-cis-cis*-clerodienyl diphosphate (CLPP) as identified by comparison to the previously reported products of the switchgrass (*Panicum virgatum*) *ent-neo-cis-trans*-CLPP synthase, PvCPS1, and a PvCPS1:F251V variant that forms *ent-neo-cis-cis*-CLPP (Fig. 6) (Pelot et al., 2018; Pelot et al., 2019).

**Fig. 6.**
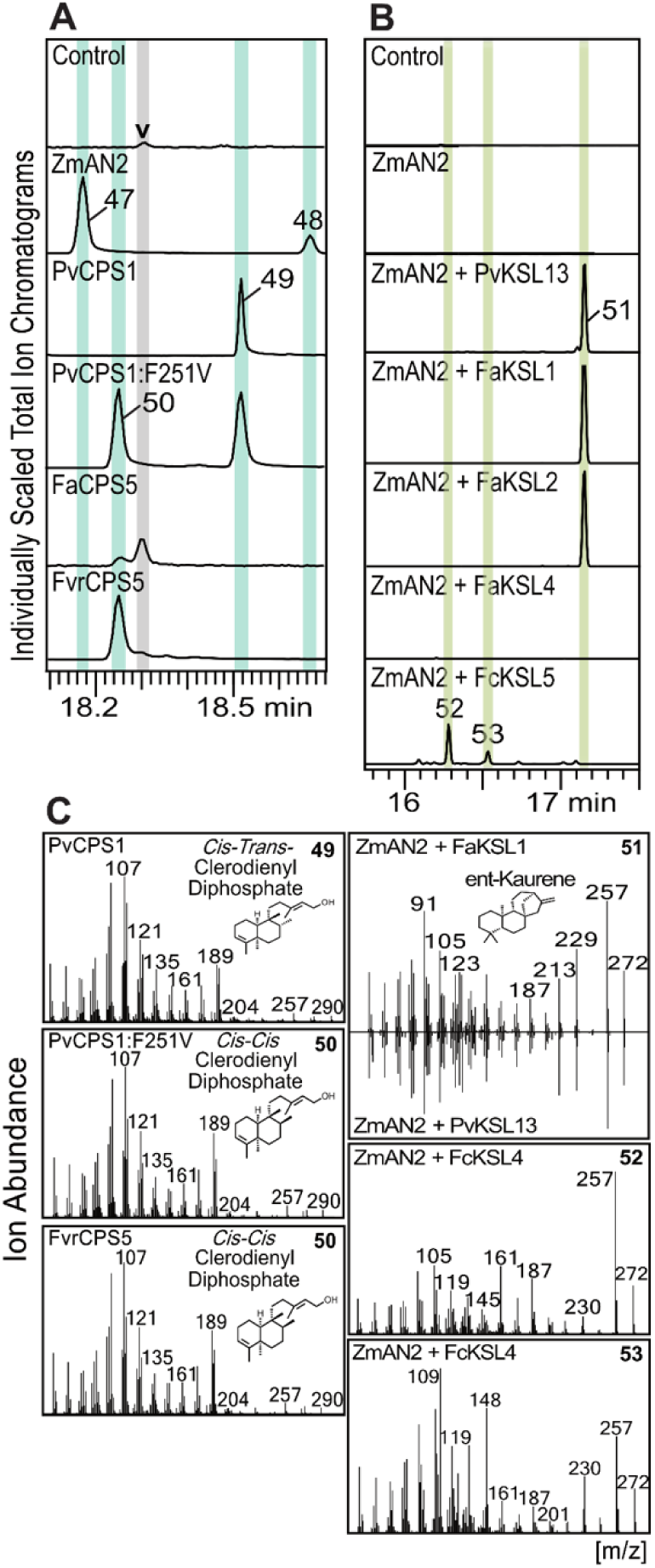
Functional characterization of diterpene synthases (diTPSs). (A) GC-MS traces of products resulting from co-expression assays of FaCPS5 or FvrCPS5 as compared to authentic standards produced by the *ent*-CPP synthase *Zea mays* AN2 (*ZmAN2*; Harris et al., 2005), the *ent-neo-cis-trans*-CLPP synthase *Panicum virgatum CPS1* (*PvCPS1*; Pelot et al., 2018) and the *PvCPS1* variant *PvCPS1:F251V* producing *ent-neo-cis-cis*-CLPP and *ent-neo-cis-trans*-CLPP (Pelot et al., 2016). (B) GC-MS traces of products resulting from co-expression assays of *FaKSL1-4* and *FcKSL5* with the *ent*-CPP synthase *Zea mays AN2* (*ZmAN2*; Harris et al., 2005). (C) Mass spectra of enzyme products identified by comparison to enzyme-produced standards. FaCPS5, and FaKSL1-4 taken from *F.* × *ananassa (Fa)* cultivar ‘Royal Royce’; FvrCPS5 *F. virginiana* (*Fvr*) accession ‘NC_96-35-2’; FcKSL5 *F. chiloensis* (*Fc*) ecotype ‘Ambato’.

Additional co-expression of the class I diTPSs FaKSL1, FaKSL4 or FcKSL5 did not show conversion of these class II diTPS products (Supplemental Fig. 13). Therefore, additional co-expression assays of FaKSL1-4 or FcKSL5 with an *ent*-CPP synthase from *Z. mays* (ZmTPS38/CPS2/AN2) and the *A. grandis* GGPP synthase were performed. No product formation was detected when co-expressing FaKSL3 or FaKSL4 (Fig. 6, Supplemental Fig. 13). By contrast, co-expression of ZmAN2 with FaKSL1 or FaKSL2 yielded *ent*-kaurene as verified by an authentic standard (Fig. 6). In addition, pairwise activity of ZmAN2 with FcKSL5 resulted in two pimaradiene-type diterpenoid products based on mass spectral comparison to known pimaradiene scaffolds. Low abundance of these products prevented further purification and structural verification by nuclear magnetic resonance (NMR) analysis.

### Terpene metabolite profiles underwent significant changes throughout strawberry domestication

To investigate the diversity of terpene compounds in strawberry aroma we selected a panel of 96 strawberry accessions comprising a variety of cultivars and wild ecotypes and accessions to represent a diverse range of aroma profiles across the germplasm curated and developed in the UC Davis Breeding Program (Supplemental Table 7). Ripe fruit harvested from field-grown plants was analyzed via SPME-GC-MS analysis and detected 171 VOCs, including short– and medium chain acetic, butyl, and hexyl acid esters such as methyl and hexyl butyrate, alcohols and aldehydes such as 2-hexanol and 2-hexenal, as well as mesifurane, y-decalactone, and 31 terpene metabolites (Supplemental Fig. 14, Supplemental Data 1, Supplemental Table 8).

Notably, a random forest analysis of the 171 identified metabolites demonstrated that linalool, nerolidol, and □-terpineol ranked among the ten most relevant metabolites defining the metabolic differences across the analyzed strawberry accessions, alongside other key strawberry aroma compounds such as Y-decalactone and mesifurane (Fig. 7). Principal components analysis (PCA) based upon the volatile random forest data showed that most samples cluster around 0 with *F. vesca* accession UC06 samples spreading out across PCA1, and cultivar EarliMiss stretching across PC2, which explain 4.3% and 3.4% of the variance, respectively (Fig 7).

**Fig. 7.**
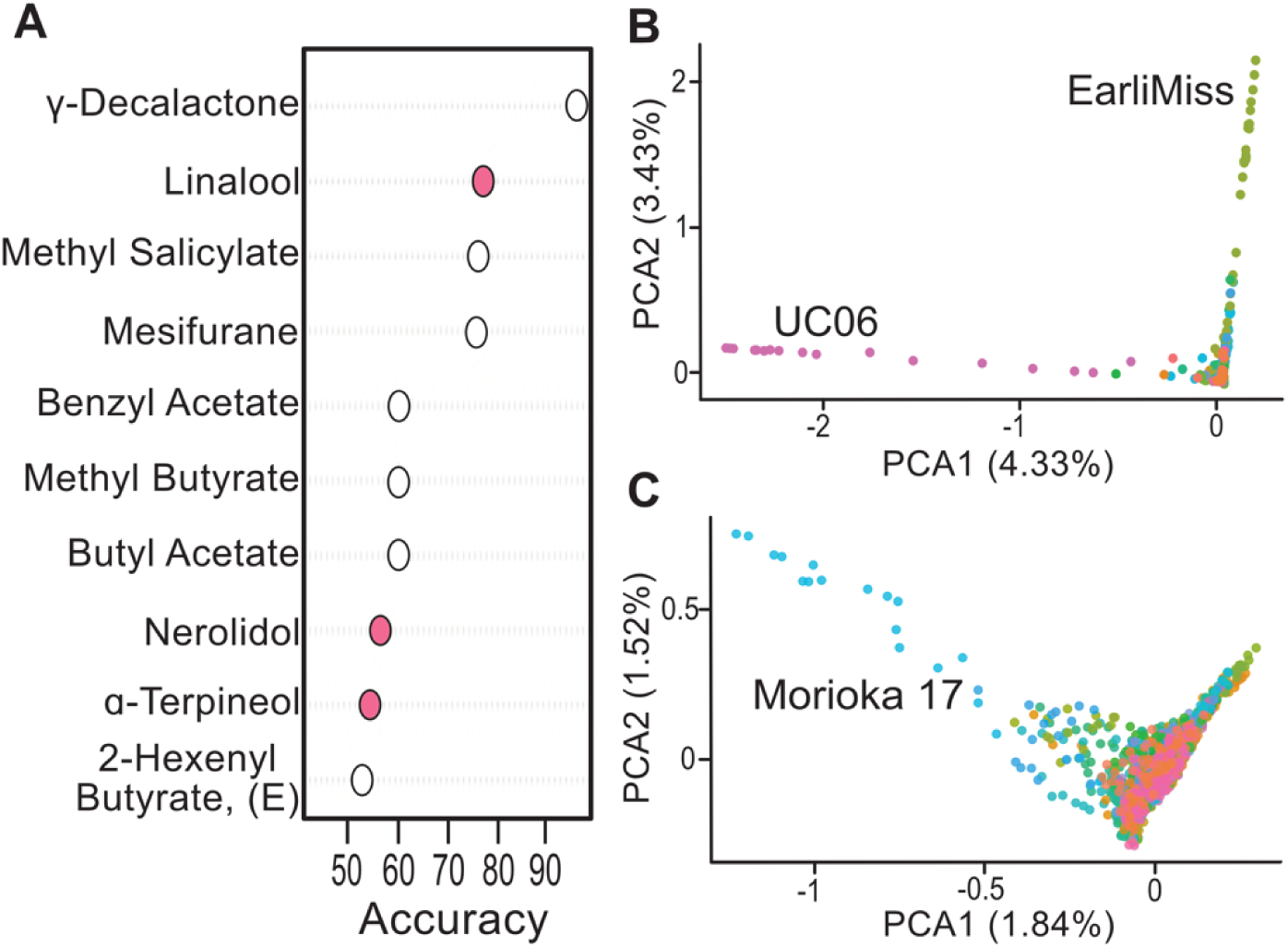
Volatile organic compound (VOC) analysis of ripe berries harvested from 96 different, field-grown accessions (**Supplemental Table 2.3**). (**A**) Scree plot of Mean Decrease Accuracy values from random forest analysis depicting the top distinguishing VOCs among the 96 tested accessions. Terpene VOCs are highlight in pink. (**B**) Principal Component Analysis (PCA) plot of VOC profiles from samples harvested in 2021 and 2022. (**C**) PCA plot of VOC profiles from samples harvested in 2021 and 2022, excluding the cultivar ‘EarliMiss’ and diploid accession ‘UC06’.

**Fig. 8.**
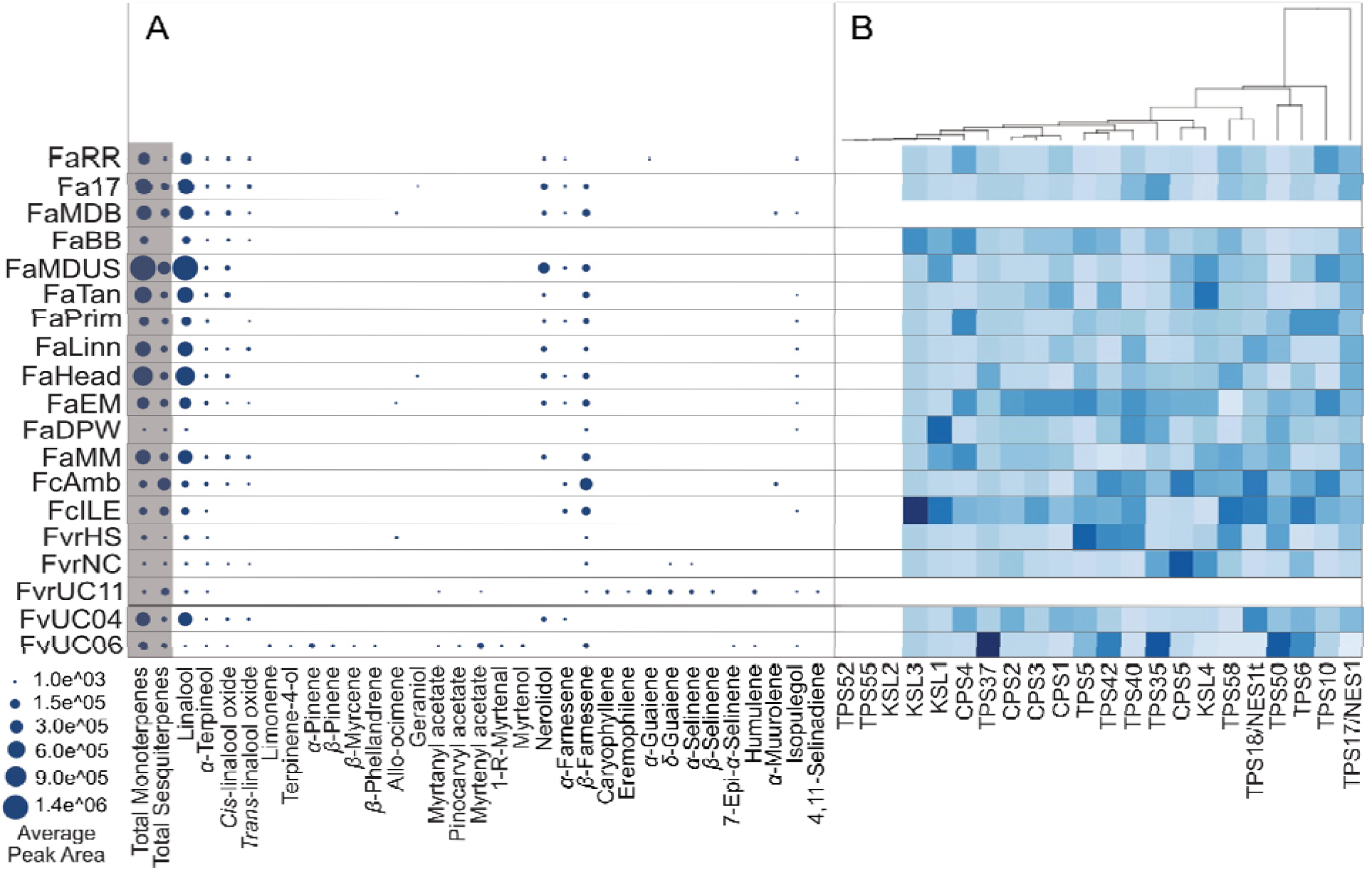
Abundance of terpene metabolites and relevant terpene synthase (TPS) genes in field-grown strawberry fruits. (**A**) Dot plot representing average normalized peak area of terpenes identified via SPME-GC-MS analysis of metabolites extracted from ripe fruit of field-grown strawberry plants. (**B**) Hierarchical cluster analysis performed on gene expression data of select TPS genes analyzed in this study. TPSs were identified based on mapping functionally characterized TPSs against the FaRR1 genome (Hardigan et al., 2021a). Gene expression data are based on three biological replicates. *F.* × *ananassa (Fa)* cultivars ‘Royal Royce’ (RR), ‘17C224P011’ (17), ‘Mara des Bois’ (MDB) for which no field-grown transcriptome data was available, ‘Beaver Belle’ (BB), ‘MDUS 5130’ (MDUS), ‘Tangi’ (Tan), ‘Primella’ (Prim), ‘Linn’, ‘Headliner’ (Head), ‘EarliMiss’ (EM), ‘Direktor Paul Wallbaum’ (DPW), and ‘Madame Moutot’ (MM). *F. chiloensis* (*Fc*) ecotypes ‘Ambato’ (Amb) and ‘Isle de Lemuy 02A White’ (ILE). *F. virginiana* (*Fvr*) accessions ‘Harris Springs’ (HS), ‘NC_96-35-2’ (NC), ‘UC11’. Diploid *F. vesca* (*Fv*) accessions ‘UC04’ and ‘UC06’.

When EarliMiss and UC06 were removed from the analysis, one additional cultivar, ‘Morioka 17’, was identified as distinctive (Fig. 7). UC06 featured the highest abundance of terpenes of all the samples with regards to terpene composition, consistent with its metabolic distinctiveness (Supplemental Data 1). For instance, UC06 was the only accession for which 1R(-)-myrtenal and both pinene metabolites were consistently detected and was one of less than five accessions for which β-myrcene, β-phellandrene, β-selinene, humulene, *trans*-pinocarvyl acetate, and (-)-myrtenol were detected. UC06 further contained ≥165 times higher amounts of myrtenyl acetate (as based on measured peak areas, Supplemental Data 1) as compared to all other tested accessions. In addition, cultivars EarliMiss and Morioka 17 differed from other accessions in the presence of *E*-β-farnesene and D-limonene and the lack of several fatty acid esters shared by most other accessions tested.

Linalool was the most abundant terpene in nearly all tested accessions and represented the most abundant terpene especially in domesticated *F.* × *ananassa* cultivars. Across the 2021 and 2022 sample collections the quantified average abundance of linalool and nerolidol showed high variation both among samples of individual accessions and across accessions (Fig. 8, Supplemental Data 1). Linalool abundance ranged from 1.3±0.8 to 5,500±4,500 ng/g dry weight (DW) and was not detectable in some accessions while accumulating to 1,960±3,160 ng/g DW in other samples. Additional abundant terpenes included □-terpineol, *cis*– and *trans*-linalool oxide, nerolidol, and □– and (*E*)-β-farnesene. In particular, modern cultivars ‘17C224P011’, ‘MDUS 5130’, and ‘Headliner’ showed the highest abundance of aroma-relevant linalool and nerolidol metabolites. By contrast, several wild or older domesticated accessions, including *F. chiloensis* Isle de Lemuy, *F.* X *ananassa* Direktor Paul Wallbaum, *F. virginiana* accessions Harris Springs and NC_96-35-2, and *F. vesca ‘*UC04’ showed no or low abundance of major terpenes such as linalool and nerolidol and featured overall low terpene diversity and abundance. Several terpene metabolites were only detected in select accessions, including allo-ocimene (cultivars EarliMiss and *‘*Mara des Bois’ and *F. virginiana* accession Harris Springs), geraniol (cultivars Headliner and 17C244P011), and □-muurolene (*F. chiloensis* ecotype Ambato and cultivar Mara des Bois). Two wild accessions showed a particularly high diversity of terpene metabolites. The diploid accession *F. vesca* UC06 contained a diverse monoterpene profile, featuring □-pinene, myrtenyl acetate, limonene, terpinen-4-ol, β-pinene, β-myrcene, β-phellandrene, *trans*-pinocarvyl acetate, 1-*R*-myrtenal, myrtanyl acetate, myrtenol, (*E*)-β-farnesene, 4,11-selinadiene, humulene, and isopulegol as based on authentic standards or best matches to reference mass spectra (Fig. 8). By contrast, the wild accession *F. virginiana ‘*UC11’ was low in monoterpenes but contained a range of sesquiterpenes not substantially detected in other accessions, including caryophyllene, eremophilene, □– and δ-guaiene, □– and β-selinene, 4,11-selinadiene, and humulene.

Additionally, slightly different VOC profiles were detected from fruits collected from the same plants when fruits were processed fresh or flash frozen (Fig. 8, Supplemental Fig. 15). For example, allo-ocimene was detected in higher amounts in flash frozen *F. virginiana* Harris Springs samples and two other accessions harvested for transcriptomic and VOC analysis, yet it was not detected in the corresponding fruits which were harvested and processed fresh for VOC analysis. Although freshly processed samples contained more terpenes (31), several terpenes, γ-dihydro-terpineol, carvomenthol, and γ-bisabolene, were detected only in flash frozen tissues.

To complement metabolite profiles, we used the transcriptome analysis described above, of field-grown, ripe fruits of two diploid *F. vesca*, two *F. chiloensis,* and two *F. virginiana* accessions and 12 *F.* × *ananassa* accessions developed between 1910 and 2017 (Fig. 8). Overall, the gene expression levels of identified TPSs did not show significant correlation with the terpene metabolite profiles. However, select TPSs displayed transcript abundance in alignment with accession-specific terpene profiles. For example, high expression of the pinene synthase *FaTPS50/PINS*, terpinolene synthase *FaTPS37*, ocimene synthase *FaTPS6*, and NES *FaTPS42* (Fig. 4) in diploid accession UC06 was aligned with the predominant presence of monoterpenes (Fig. 8).

Genotypes containing higher amounts of farnesene also showed higher abundance of the farnesene synthases *FaTPS6* and *FaTPS10*, including cultivars MDUS 5130, Primella, and EarliMiss, and *F. chiloensis* ecotypes Ambato and Isle de Lemuy. However, *FaTPS6* or *FaTPS10* were also abundant in cultivars producing no or low levels of farnesene such as Royal Royce and Madame Moutot. Similarly, with the exception of *F. vesca* accession UC06 that showed relatively high abundance of *FaTPS18/NESt*, genotypes abundant in linalool and nerolidol did not show high gene expression of NESs *FaTPS17/NES1*, *FaTPS18/NESt*, *FaTPS40,* and *FaTPS42*.

### Alteration of terpene metabolism during fruit ripening

To assess changes in terpene abundance during fruit ripening, two modern *F.* × *ananassa* cultivars with contrasting fruit properties, Royal Royce and Mara des Bois, were grown in the greenhouse and fruit samples were harvested at green, white, underripe, ripe and overripe stages (Fig. 9). SPME-GC-MS analysis identified 23 volatile terpenes, 17 of which matched compounds identified in field-grown, ripe fruits. Consistent with field-grown fruit, linalool was the most abundant terpene, followed by nerolidol, linalool oxide, and li:-pinene. Green fruits for both tested cultivars contained low quantities of linalool with 1-5 ng/g DW, whereas ripe fruits contained 484 ng/g DW linalool and 109 ng/g DW nerolidol in Mara des Bois and 2,100 ng/g DW linalool and 270 ng/g DW nerolidol in Royal Royce (Supplemental Data 2). While Mara des Bois fruit contained a lower diversity of terpenes as compared to Royal Royce, both cultivars showed low terpene abundance in green and white fruit. Royal Royce fruit showed an increase in linalool as well as linalool oxide and li:-pinene at the underripe stage. In ripe fruit, monoterpene abundance except for linalool was decreased, while an increase in nerolidol was observed. This trend continued in overripe fruit that showed a higher abundance in several sesquiterpenes. Although lower in terpene abundance, a similar pattern was observed in Mara des Bois berries with an increase in linalool and nerolidol in ripe and overripe fruit.

**Fig. 9.**
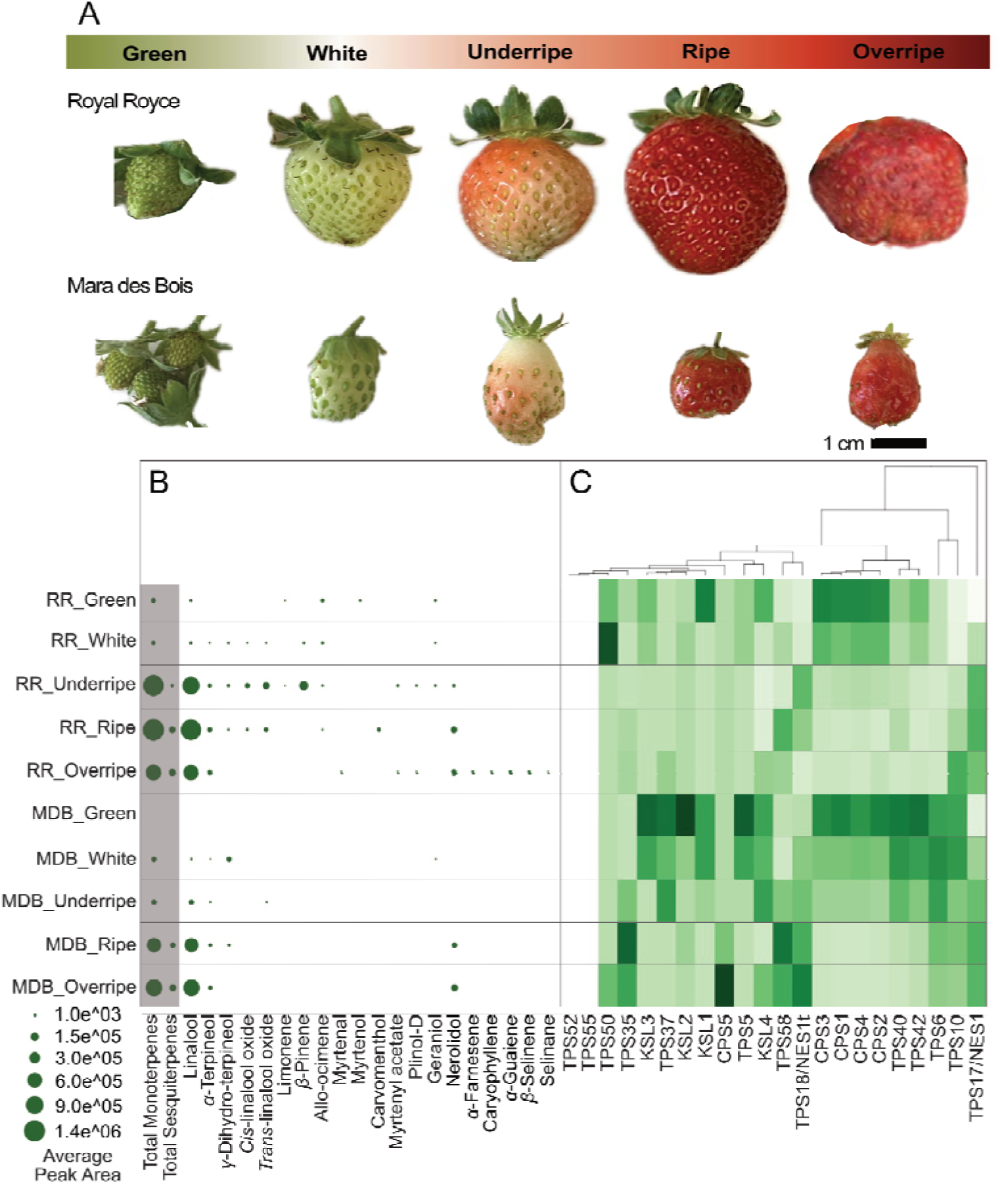
Alterations in terpene metabolism during fruit ripening. (**A**) Images of representative *Fragaria* × *ananassa* Royal Royce (RR) and Mara De Bois (MDB) berries at the green, white, underripe, ripe, and overripe developmental stages. (**B**) Dot plot showing average normalized peak area of terpenes identified via SPME-GC-MS analysis of metabolites extracted from berry tissue. Note that no MDB tissue was available for metabolite profiling of green fruit due to limited berry production in this cultivar. (**C**) Hierarchical cluster analysis performed on gene expression data of select terpene synthase (TPS) genes analyzed in this study using the same berry tissues. TPSs were identified based on mapping functionally characterized TPSs against the FaRR1 genome (Hardigan et al., 2021a). Gene expression data are based on three biological replicates.

Transcriptome analysis of the same fruit tissues revealed that core rate-limiting genes of the MEP pathway, *Deoxyxylulose 5-phosphate synthase* (*DXS*) and *Deoxyxylulose 5-phosphate reductoisomerase* (*DXR*), showed highest transcript abundance in green fruit in either cultivar with a continued decrease in gene expression during ripening (Supplemental Fig. 16).

MEP genes *4-hydroxy-3-methylbut-2-enyl diphosphate reductase 1 (HDR1)* and *HDR3* showed slightly higher expression in Mara des Bois fruits through ripening whereas *HDR2* and *HDR4* expression gradually increased in Royal Royce fruits during ripening with the highest normalized counts seen in overripe fruits. By contrast, *HMG-CoA reductase* (*HMGR*), as a key rate-limiting gene of the MVA pathway, increased in transcript abundance during ripening with highest expression levels in overripe fruit. Gene expression patterns of *TPSs* did not show a clear correlation with terpene abundance (Fig. 9). However, a moderate increase in the transcript abundance of the NESs, *FaTPS17/NES1* and *FaTPS18/NES1t*, was observed during fruit ripening in both cultivars that is aligned with an accumulation of linalool and nerolidol. In addition, expression of *FaTPS50* was high in white Royal Royce fruit possibly associated with the increase in li:-pinene in underripe fruit. Notably, gene expression of *FaCPS1-4* and *FaKSL1-4* was highest in green fruit of Royal Royce and/or Mara des Bois. In contrast, transcript abundance of *FaCPS5* with a proposed function in specialized diterpenoid metabolism was highest in overripe Mara des Bois, but not Royal Royce, fruit.

## Discussion

Species-specific blends of volatile terpene metabolites serve as chemical cues in various plant-environment interactions, including roles in long-distance chemical defenses and pollinator interactions (Tholl, 2015). Beyond their physiological importance, terpene VOCs are core components of aroma traits in many fruit crops including strawberry (Nieuwenhuizen et al., 2013; Ilc et al., 2016; Ulrich and Olbricht, 2016; Fan et al., 2021a). Leveraging advanced omics and synthetic biology approaches now enables a detailed investigation of the diversity of crop aroma and the underlying genes, enzymes, and pathways. The discovery of the strawberry TPS gene family provides a genomic atlas for the biosynthesis of aroma-relevant terpenes. The presence of 75 full-length TPS gene candidates in the octoploid *F.* × *ananassa* ‘Royal Royce’ genome (FaRR1) places the strawberry TPS family at the higher end of the TPS families identified in other species of the rosid clade, including eucalyptus (*Eucalyptus grandis*) with 113 TPSs (Kulheim et al., 2015), grape (*Vitis vinifera*) with 39 TPSs (Martin et al., 2010), apple (*Malus domestica*) with 55 TPSs (Nieuwenhuizen et al., 2013), and almond (*Prunus dulcis*) with seven characterized and ∼23 predicted TPSs (Nawade et al., 2019). Notably, the genome of the ancestral diploid *F. vesca* contains a smaller TPS family of 43 predicted members (Zhang et al., 2022), 25 of which have orthologs in the octoploid FaRR1 genome (Supplemental Fig. 6).

Additionally, the presence of TPSs (*FaTPS3-5, 11-15, 25, 33, 41-45, 53*, and *63*) in the FaRR1 genome that lack syntenic equivalents in *F. vesca* support the expansion of the TPS gene family in octoploid strawberry through a combination of polyploidization and tandem gene duplication events (Fig. 2, Supplemental Fig. 6). For instance, closely co-localized TPS genes are present on the distal ends of chromosome 3 homeologs, including the □-farnesene synthases *TPS6, TPS7,* and *TPS10*; the NESs *FaTPS17/NES1* and *FaTPS18/NES1t*, as well as *FaTPS11/NES2* and *FaTPS12-14* co-located on chromosome 3B, suggesting their evolutionary origin from repeated gene duplications.

In addition to the previously reported NESs, *FaTPS17/NES1*, *FaTPS18/NES1t*, and *FaTPS11/NES2*, biochemical characterization showed that TPS40 and FaTPS42 also form nerolidol and linalool. TPS40 and FaTPS4 are phylogenetically distant from known NES enzymes within the TPS-b family, highlighting that the strawberry genome contains a larger group of NESs, thus providing genetic redundancy for the biosynthesis for these major terpenes in strawberry and likely contributing to the high abundance of these aroma compounds in most strawberry cultivars and accessions. Indeed, higher gene expression of especially *FaTPS40* and *FaTPS42* in early cultivars and wild relatives as compared to modern cultivars indicates an important role in nerolidol and linalool accumulation (Fig. 8).

Beyond the identification of additional NES enzymes, biochemical analysis of 20 mono– and sesqui-TPSs selected for their predicted functions and expression in strawberry fruits revealed the biosynthesis of 47 terpene metabolites, including known strawberry terpenes and metabolites that, to the best of our knowledge, have not previously been reported in strawberry such as □– and δ-guaiene, □-muurolene, β-selinene, eremophilene, isopulegol, 7-epi-–selinane, pinocarvyl acetate, and 4,11-selinadiene (Fig. 10). The identified TPS products reveal the biosynthetic origin of over half of the terpene metabolites detected in the strawberry accessions analyzed in this study (Fig. 8), notably including linalool, □-terpineol, limonene, □[r]-pinene, terpinen-4-ol, allo-ocimene, nerolidol, □-farnesene, caryophyllene, humulene, □/δ-guaiene, [r]-selinene, eremophilene, and □-muurolene. The functional identification of FvTPS6, FaTPS7, and FaTPS10 as □-farnesene synthases that form a syntenic grouping with *FaTPS21* in the FaRR1 genome provides biochemical evidence for previously reported functional predictions on the basis of eQTL analyses (Barbey et al., 2021). In addition to FaTPS6, 7 and 10, FcTPS52, identified in the *F. chiloensis* ecotype Isle de Lemuy, also produced □-farnesene but represented a multi-product TPS also forming other sesquiterpenes such as (*Z*)-–*trans*-bergamotol, (*Z*,*E*)-– farnesene, *E*-–bisabolene, thus possibly contributing to the diversity of strawberry aroma profiles (Supplemental Fig. 11).

**Fig. 10.**
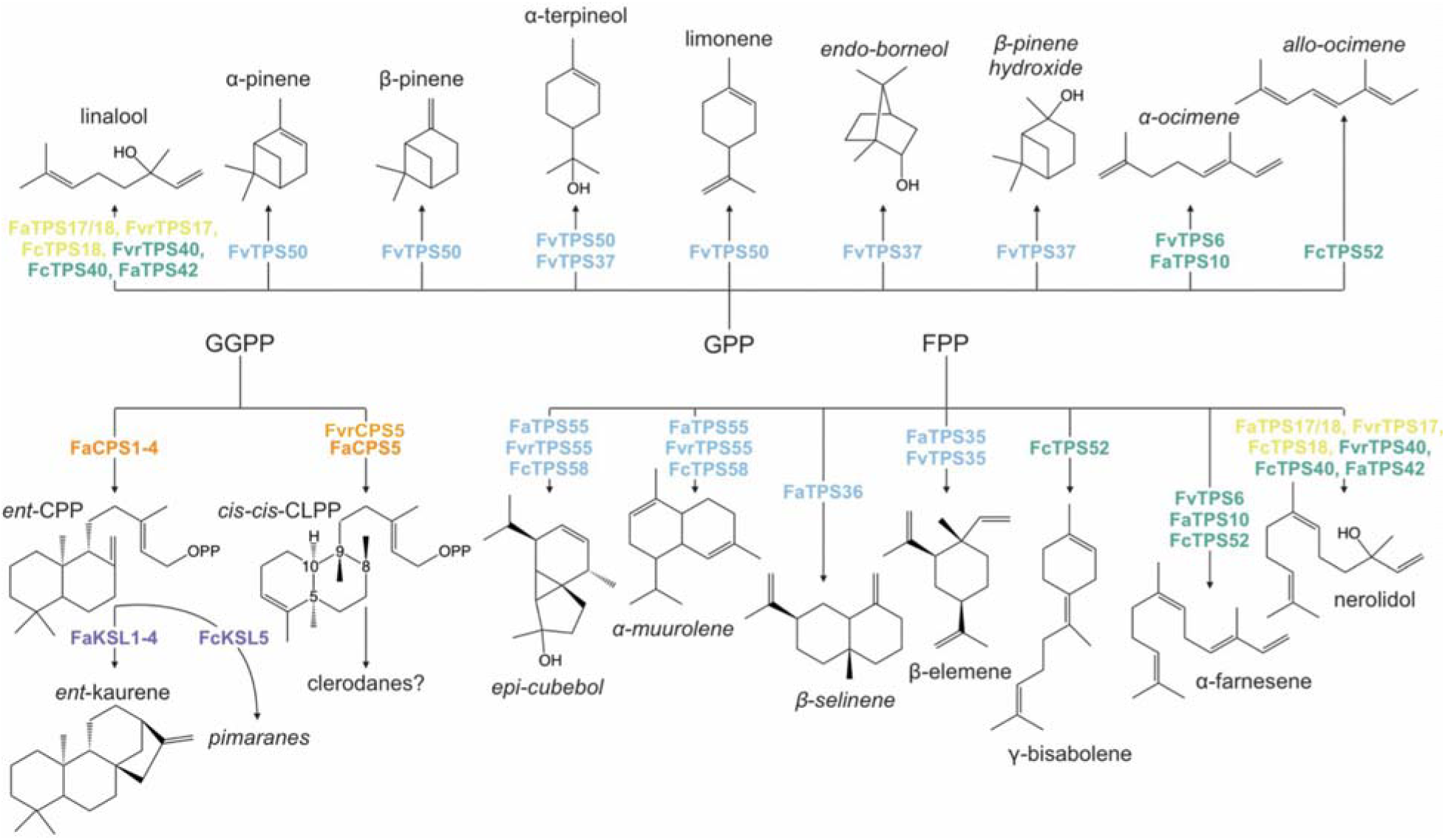
Strawberry terpene metabolism. Shown are major terpene products formed by the 27 terpene synthase (TPS) enzymes functionally analyzed in this study. TPS enzymes are colored corresponding to TPS clades assigned in Fig. 2.3. Abbreviations: TPS, terpene synthase; GGPP, geranylgeranyl diphosphate; GPP, geranyl diphosphate; FPP, farnesyl diphosphate; CPP, copalyl diphosphate; CPS, copalyl diphosphate synthase; KSL, kaurene synthase-like.

Production of □-pinene and several low abundant byproducts by FvTPS50/PINS confirm earlier studies that demonstrated the function of FvTPS50/PINS, and a downstream-acting cytochrome P450 monooxygenase and alcohol acyltransferase, in the biosynthesis of myrtenol and myrtenyl acetate (Aharoni et al., 2004). Transcript and metabolite profiling of diploid accession UC06 showed the highest abundance of *FaTPS50* transcript and the □/li:-pinene, myrtenol, and myrtenyl acetate products, suggesting that this pathway is highly productive in this accession. In addition to FvTPS50/PINS, FvTPS37 produced linalool, □-terpineol, and several other monoterpenes, thus representing another enzymatic source of consumer-preferred aroma metabolites in strawberry. Differential gene expression of *FvTPS37* across the breeding material may contribute to the function of *FvTPS37* in strawberry aroma metabolism (Fig. 7 & 8). In addition, production of as many as 23 different sesquiterpene products by *FaTPS35*, *FaTPS36*, *TPS55*, and *FcTPS58* reveals the genetic basis for the biosynthesis of more complex terpene blends in some of the tested accessions such as *F. virginiana* UC11 and NC_96-35-2 and diploid UC06 (Supplemental Fig. 12). Collectively, the identified TPS functions demonstrate that catalytic redundancy and functional diversity within the strawberry TPS family provide the biochemical basis for the complex aroma profiles in different cultivars and accessions, especially in wild and diploid strawberries that often feature a higher diversity of terpenes (Aharoni et al., 2004; Bianchi et al., 2014; Negri et al., 2015; Ulrich and Olbricht, 2016; Urrutia et al., 2017; Duan et al., 2018).

In addition to a large family of mono– and sesqui-TPS enzymes, genome mining identified ten diTPS candidate genes in the FaRR1 genome. Sequence analysis suggested that FaCPS1-4 encode for *ent*-CPP synthases with possible functions in gibberellin (GA) phytohormone as well as specialized diterpenoid metabolism based on related enzymes identified in other plant species (Peters, 2010; Karunanithi and Zerbe, 2019). By contrast, FvrCPS5 (NC_96-35-2) and FaCPS5 (Royal Royce) were predicted as specialized diTPSs and indeed produced *ent-neo-cis-cis*-clerodienyl diphosphate (CC-CLPP), a member of the group of clerodane diterpenoids that are abundant in species of mint and a few grain crops where they have probable functions in stress response mechanisms (Peters, 2010; Pelot et al., 2017; Johnson et al., 2019a; Johnson et al., 2019b; Muchlinski et al., 2021; Tiedge et al., 2022). Co-localization of *FaCPS5* with a predicted class I diTPS, *FaKSL4*, on chromosome 3D suggested a possible pairwise activity of the encoded enzymes in clerodane biosynthesis. However, co-expression studies in *N. benthamiana* or an engineered *E. coli* platform did not reveal any coupled reaction products, thus requiring more expansive future studies to elucidate the diterpenoid-metabolic network in strawberry.

Congruent with past research, terpene-derived aromas in ripe fruits are dominated by linalool, nerolidol, and □-terpineol in *Fragaria* × *ananassa* fruits, whereas wild diploid accessions are dominated by olefinic monoterpenes such as □/li:-pinene (Aharoni et al., 2004; Chambers et al., 2012) (Fig. 8). Much diversity in both quantity and complexity of terpene profiles exists across the diversity panel analyzed in this study as exemplified by linalool amounts varying by a factor of more than 500 between the highest and lowest linalool-producers. This reflects recent breeding efforts having selected for high linalool cultivars based upon known positive correlations with this compound and consumer preferences (Schwieterman et al., 2014; Ulrich and Olbricht, 2016; Fan et al., 2021b). Similarly, significant differences in nerolidol abundance were detected between most non-*F.* × *ananassa* accessions and more modern breeding lines such as ‘EarliGlow’ and 17C224P001 (Supplemental Data 3). The functional characterization of previously reported and unidentified TPSs provides a better comprehension of the functional range of the strawberry TPS family and its contribution to terpene variation across different cultivars and accessions, thus providing a resource to link biochemical functional annotations with rapidly expanding genomic and genetic insights into key regulators of terpene aroma profiles. A broader knowledge of the metabolic diversity across the existing germplasm can further aid the identification of strawberry lines for breeding of advanced, consumer-preferred strawberry aromas.

In addition to the terpene diversity across different strawberries, analysis of a fruit ripening time course provided insight into the terpene-metabolic alterations during fruit ripening. Consistent with the high abundance in ripe fruits, linalool and nerolidol abundance increased throughout fruit ripening in both modern cultivars tested here with the largest increase starting in unripe fruits (Fig. 9). High gene expression of the NESs *FaTPS17/NES1* and *FaTPS18/NES1t* in underripe and ripe fruits indicates a likely involvement of these genes in driving metabolite production during fruit ripening. Likewise, increase in linalool oxides, limonene, myrtenol and li:-pinene in green or white Royal Royce fruits correlated with increasing transcript abundance of *FaTPS50*, supporting a biosynthetic relationship. Interestingly, several diTPS transcripts were also higher in green fruits in both tested cultivars. Consistent with their predicted functions as *ent*-CPSs, it can be speculated that these genes are involved in GA phytohormone biosynthesis at the early stages of fruit development (Csukasi et al., 2011; Liao et al., 2018). However, given the specialized function of FaCPS5 and the formation of many specialized diterpenoids from *ent*-CPP in other plant species (Peters, 2010; Pelot et al., 2017; Johnson et al., 2019a; Johnson et al., 2019b; Muchlinski et al., 2021; Tiedge et al.), possible roles in stress response mechanisms in developing fruits cannot be excluded.

## Experimental Procedures

### Plant material

A panel of 96 strawberry accessions comprising a selection of both ecotypes and modern cultivars derived from the UC Davis Strawberry Breeding Program collection were grown under established commercial conditions at Wolfskill Experimental Orchards in Winters, California in 2021 and UC Davis Vegetable Crop Fields in Davis, CA in 2022 (Supplemental Table 7) (Petrasch et al., 2022). Ripe fruits from each accession were harvested between 8-10 AM throughout the growing season every two to three weeks between April and June and stored at 4 □ with 70% relative humidity until processing. Twice in early May during the middle of the 2021 season, additional ripe fruits from 17 accessions which span the range of strawberry domestication and were expected to harbor a diversity of volatile profiles were harvested for transcriptome analysis (Supplemental Table 5). For this purpose, berries were immediately sliced, frozen in liquid N_2_, transported on dry ice and stored at –80 □ until use for metabolite and transcript analysis. In addition to field harvests, two cultivars, Royal Royce and Mara de Bois, were grown at UC Davis greenhouses. Two plants of each cultivar were planted into pots in Fall 2020 and kept throughout the winter so that berries were harvested as needed throughout the first week of March 2021 through the first week of June 2021. Fruits were harvested fully green, white, turning or underripe, fully ripe, and overripe and processed as described above for further transcriptomic and aroma analysis (Jiménez et al., 2024).

### Metabolite analysis

For metabolite profiling of field-grown plants, 3-9 fully ripe berries (depending on different fruit production across the tested accessions) were harvested and pooled from four individual plants of each accession. Berries were then divided into three technical replicates and halved for subsequent metabolite or transcriptome analysis. For metabolite analysis, the samples were weighed and supplemented with an equal amount (w/v) of 20% NaCl and 50 pp Y-undecalactone (final concentration, in methanol; Sigma-Aldrich, St. Louis, MO, USA) as an internal standard (Jetti et al., 2007). Samples were homogenized in a blender (SharkNinja, Needham, Massachusetts, USA) and 5 mL of the solution transferred to pre-weighed 20 mL glass crimp-cap SPME vials (Agilent Technologies, Santa Clara, CA, USA). Samples were capped, labeled, and stored at –20.

Samples were analyzed using solid-phase microextraction-gas chromatography-mass spectrometry (SPME-GC-MS) on an Agilent Intuvo 9000 GC coupled with a 5977b MS with a 350 XTR Electron Ionization (EI) source (Agilent Technologies). Samples were incubated at 30□ for 5 min during SPME fiber adsorption with a 50/30 µm DVB/Carboxen-WR/PDMS SPME fiber (Supelco, Bellefonte, PA, USA), followed by injection and 6 min desorption into the inlet at 240 □using helium as a carrier gas at a flow rate of 1.2 mL per min. Compound separation was achieved on a DB-WAX UltraInert column (30 m x 0.25 mm x 0.25 µm column; Agilent Technologies). Oven settings were held at 40 □for 2 min, 4 /min to 70□, 6 /min to 180, 15 /min to 220, and held at 220 for 2 min. Mass ranges were recorded from 40-400 *m*/*z* with an EI energy of 70 eV and detector temperature set to 230. Compounds were identified based on available authentic standards (Supplemental Table 9) or matches to reference mass spectral databases of the National Institute of Standards and Technology (NIST, version 17.1). Compound abundance was calculated based on relative peak areas and integration using MassHunter Unknowns Analysis software with default deconvolution settings (Agilent Technologies). Metabolite abundance values were normalized against the internal standard and sample dry weight after drying samples for 48 hrs at 100, and subtraction of saline buffer content and vial weight. The same total ion chromatograms were secondarily analyzed against eight terpene standard curves and normalized against the same dried berry weights used above to report compound concentrations in ng/g dry berry weight. Relative quantification of eight quantified metabolites were performed using MassHunter Quantification and Unknown Analysis software (Agilent Technologies) and are shown in Supplemental Table 9. Cleaned, normalized GC-MS data were analyzed using the randomForest package (Liaw and Wiener, 2002) in the R Statistical Software (v.4.4.0; (R Core Team, 2021) with default parameters of ntree 5000. The square root of total variables entered were tested at each split. Thus, for the analysis of all 171 detected VOCs 13 randomly chosen aroma metabolites were tested at each split. Mean Decreased Accuracy (MDA), or the level of accuracy lost at each tree node when a particular metabolite is omitted, were extracted from each random forest analysis to estimate metabolite impact within sample aroma profiles. Principal Component Analyses and Multidimensional Scaling plots were generated within the randomForest package and R Statistical Software.

### RNA isolation and transcriptome analysis

The second half of harvested flash frozen berries (see above) were ground into a fine powder in liquid N_2_ using mortars and pestles. Approximately 2 g of ground tissue was mixed with 6 mL warm extraction buffer (2% v/v CTAB, 2% PVP, 100 mM Tris, 40% NaCl, 25 mM EDTA, 0.2 g spermidine, 2% mercaptoethanol) and incubated at 65 for 5 min (Blanco-Ulate et al., 2013). Equal volumes of 24:1 chloroform:isoamyl alcohol was added, mixed, and centrifuged at 4,000 rpm for 30 min at 4. The supernatant was collected, supplemented with 24:1 chloroform:isoamyl alcohol and centrifuged again. The supernatant was then supplemented with 10% 1 M Potassium Acetate, centrifuged at 4,000 rpm for 20 min at 4, and the resulting supernatant added to a ¼ volume of 10 M Lithium Chloride. The sample was mixed gently and precipitated overnight at –20 before centrifugation at 4,000 rpm for 45 min at 4. The resulting pellet was resuspended in 60 µL RNAse-free water and cleaned using the Zymo Research Quick-RNA Mini-Prep Kit and DNase1 treatment (Zymo Research Corporation, Irvine, CA) as per manufacturer’s instructions.

Library preparation and transcriptome sequencing was performed by the NovoGene Corporation Inc. using NovaSeq 6000 paired-end short read sequencing using the NEBNext and AMPure XP suite of reagents and protocols (NEB, USA). Transcripts were cleaned using the HTStream suite to remove adapters, polyA tails, and reads under 50 bp. FastQC and MultiQC were used to assess RNA read quality before proceeding (Andrews, 2010). The remaining high-quality reads (20-29.6M 150 bp reads with an average of ∼24M reads x2 per sample) were aligned to the *FaRR1* genome and read counts were obtained using the STAR package with default parameters (Dobin et al., 2013; Petersen et al., 2015). Additionally, to identify transcripts that did not align to the *FaRR1* genome, a Trinity *de novo* assembly (Grabherr et al., 2011; Haas et al., 2013) was performed with default parameters and assembled using bowtie2 for all reads of each genotype (Langmead and Salzberg, 2012). Cleaned reads were normalized using the TMM function with default parameters in the edgeR software package (Robinson et al., 2010) and averaged across replicates for each genotype, after which any contigs or genes with fewer than 10 counts for at least one sample were discarded. The remaining data were then both mapped to the *FaRR1* genome and assembled *de novo* to identify transcripts that had less than 95% sequence identity with any of the 75 full-length *FaRR1* genomic TPS candidates (Supplemental Table 3 & 4). Gene counts were normalized and analyzed in R using TMM in edgeR (Robinson et al., 2010), topGO (Alexa and Rahnenfuhrer, 2022), volcano, and heatmap (Perry, 2024) packages.

### Gene identification, synthesis, and construct design

To identify TPS gene candidates and core genes of the MEP and MVA pathways, the *FaRR1* genome (2021) and the transcriptome data generated in this study were BLAST searched against a curated database of protein sequences that represent the functional space of TPSs and MEP/MVA enzymes across higher plants (Zerbe et al., 2013). The obtained transcripts were manually curated for presence of characteristic catalytic domains, sequence completeness, and phylogenetic relationships as described previously (Zerbe et al., 2013). The resulting 104 possible TPS were narrowed further based on sequence length: mono– and sesqui-TPS candidates with a 450-750 amino acids (AA) length and diTPS with a length of 550-850 AA were selected for further analysis, resulting in 27 full-length TPS genes (Supplemental Table 4). Transcripts featuring expression counts lower than 10 or lacking *de novo* transcripts were also not considered for further study.

Full-length or N-terminally truncated (removal of the predicted plastidial transit peptides; Supplemental Table 4) genes were obtained via DNA synthesis and codon optimized for expression in *E. coli* (Twist Biosciences, South San Francisco, CA, USA). For protein purification and *in vitro* enzyme assays, TPS genes were inserted into the pET28a(+) expression vector (EMD Millipore, Burlington, MA, USA) in frame with the N-terminal poly-His tag. For *in vivo* co-expression analysis in *E. coli*, synthesized TPS genes were inserted into the pACYC vector (EMD Millipore, Burlington, MA, USA) containing previously confirmed functional copies of either *Abies grandis* (Grand Fir) GPP synthase, *Zea mays* FPP synthase (*ZmFPP4*), or the *Abies grandis* GGPP synthase in the second multiple cloning site.

### In vitro enzyme assays

Synthesized TPS constructs in the pET28a(+) vector were expressed in *E. coli* BL21DE3-C41 cells (Sigma-Aldrich) and Ni^2+^-NTA affinity purified as follows: 100 mL cultures were induced with 1 mM isopropyl-β-D-1-thiogalactopyranoside (IPTG; Sigma-Aldrich) and grown for 24 hours at 16°C. Cultures were pelleted at 4 and resuspended in 40 mL of wash buffer (20 mM Tris Base, 50 mM KCl, pH 7) then resuspended in 4 mL of Phosphate-buffered saline (PBS) lysis buffer (50 mM H_2_NaPO_4_, 200 mM NaCl, 10% glycerol, 15 mM Imidazole, pH 7) with freshly added 0.5 mM phenylmethylsulfonyl fluoride (PMSF; Sigma-Aldrich), and 1 mM Dithiothreitol (DTT; Sigma-Aldrich) and sonicated in an ice bath at 20% amplitude for a total of 2 min. Crude lysates were pelleted and stored at –80. Protein was purified at 4 using equilibrated 750 µL Ni-NTA agarose. Lysates were added to Ni-NTA beads with 4 mL of lysis buffer and batch bound for 1 hour. Lysates were allowed to settle for 30 min before they were washed with 2 mL of wash buffer 1 (50 mM H_2_NaPO_4_, 300 mM NaCl, 5% glycerol, 25 mM Imidazole, pH 7) and followed with 4 mL of wash buffer 2 (50 mM H_2_NaPO_4_, 300 mM NaCl, 5% glycerol, 35 mM Imidazole, pH 7). Protein was eluted using Elution Buffer (50 mM H_2_NaPO_4_, 300 mM NaCl, 350 mM Imidazole, pH 7). Presence of all purified proteins were confirmed via Western Blot using the BioRad TransBlot and Alkaline-Phosphatase system from BioRad Technologies (Hercules, CA) and Alkaline Phosphatase Anti-6x His tag^®^ antibody (AD1.1.10; Abcam Limited, Cambridge, UK).

Single-vial mono– and sesqui-TPS enzyme assays were performed as previously described (Martin et al., 2004). In brief, 100 μg of recombinant proteins were combined with 15 μM GPP or FPP (Sigma-Aldrich) as substrates in a total volume of 1 mL of assay buffer (25 mM HEPES, pH 7.2, 100 mM KCl, 10 mM MnCl2, 10% glycerol, 5 mM DTT) and overlaid with 600 µL of hexane (Thermo Fisher Scientific) to trap terpene products during the assays. Each sample was supplanted with 0.03 ppm isobutylbenzene (Sigma-Aldrich) as internal standard. After 2 h incubation at room temperature, enzyme products were extracted by vortexing and analyzed via GC-MS as described below.

### In vivo enzyme co-expression analysis

For co-expression assays of selected TPSs, *E. coli* BL21DE3-C41 cells were co-transformed with individual TPS constructs in the pACYC vector, as well as plasmids carrying either *Abies grandis* GPP synthase, *Zea mays* FPP synthase (*ZmFPP4*), or the *Abies grandis* GGPP synthase (pGG plasmid) and key enzymes of the MEP pathway (pIRS plasmid) as described previously (Cyr et al., 2007; Morrone et al., 2010). For combinatorial expression of diTPSs, class I diTPSs were cloned into the pET28a(+) plasmid and class II diTPSs were inserted into the pACYC vector to maintain replication complementarity. For all combinations, co-expression assays were performed as described previously (Pelot et al., 2017). In brief, cultures were grown at 37 °C and 180 rpm in 50 mL of Terrific Broth medium to an OD_600_ of ∼0.6 before cooling the cultures to 16°C and induction with 1 mM IPTG for 72 h with the addition of 1.25 mL 1M Sodium pyruvate and 50 μL MgCl_2_. Products were extracted with 50 mL of hexane containing 17.5 ng/μL 1-eicosene (Sigma-Aldrich) as internal standard, air dried, and resuspended in 1 mL of n-hexane for GC-MS analysis.

### GC-MS analysis of terpene synthase products

Analysis of extracted enzyme products was conducted via liquid-injection GC-MS on an Agilent 7890B GC coupled with a 5977A Extractor EI MS (Agilent Technologies). Samples were injected into the inlet at 250 using Helium as a carrier gas at a flow rate of 1.2 mL/min. Compound separation was achieved on a HP-5MS column (30 m x 0.25 mm x 0.25 µm column; Agilent Technologies). Oven settings for analysis of the mono– and sesqui-terpenes were held at 50 for 2 min, 8 /min to 200, 25 /min to 250, and held at 250 for 1 min. Oven settings for analysis of diterpene products were held at 50 for 3 min, 15 /min to 300 and held at 300 for 4 min. Mass ranges were recorded after a 6-min solvent delay for mono– and sesqui-terpenes and 13 min for diterpenes from 40-400 *m*/*z* with an EI energy of 70 eV and detector temperature set to 230. Total ion chromatogram peaks were integrated using MassHunter Unknowns Analysis software (Agilent Technologies) and compounds were identified based on comparison to available authentic standards (Supplemental Table 9) or significant matches to reference mass spectral databases of the National Institute of Standards and Technology (NIST, version 17.1).

### Heat map and hierarchical cluster analysis

Gene candidates of the MEP and MVA pathways were plotted across the FaRR1 genome using Chromplot in R (Oróstica and Verdugo, 2016) (Supplemental Table 1). Heatmaps and hierarchical cluster analysis on selected transcriptome data was analyzed using the heatmap package in R (Perry, 2024).

### Phylogenetic analysis

For phylogenetic studies, protein sequences of the identified TPSs and known TPSs from other Rosaceae and plant species were aligned using the MUSCLE package with neighbor joining parameters (Edgar, 2004) and cleaned with BMGE default parameters (Criscuolo and Gribaldo, 2010). A maximum likelihood phylogenetic tree was generated with the NGphylogeny.fr user interface PHYML with Smart Model Selection (SMS), Subtree Pruning and Regrafting (SPR), minimum theoretical information criterion (AIC) and 500 bootstraps (Guindon et al., 2010; Lefort et al., 2017; Lemoine et al., 2018; Lemoine et al., 2019). The tree was visualized using Newick displays and formatted using Geneious (Junier and Zdobnov, 2010; 2012).

## Accession Numbers

The RNA-sequencing data were submitted to the National Center for Biotechnology Information Sequence Read Archive (NCBI SRA; ncbi.nlm.nih.gov/sra), BioProject: PRJNA1172127. Nucleotide sequences for functionally characterized transcripts reported in this study were submitted to the NCBI GenBank (SUB14747557) with the following accession numbers: PQ433313 (*FcKSL5*); PQ433314 (*FcTPS18*); PQ433315 (*FcTPS40*); PQ433316 (*FcTPS52*); PQ433317 (*FcTPS58*); PQ433318 (*FvTPS6*); PQ433319 (*FvTPS35*); PQ433320 (*FvTPS37*); PQ433321 (*FvTPS50*); PQ433322 (*FvrCPS5*); PQ433323 (*FvrTPS5*); PQ433324 (*FvrTPS17*); PQ433325 (*FvrTPS40*); PQ433326 (*FvrTPS55*); PQ433327 (*FaCPS5*); PQ433328 (*FaKSL1*); PQ433329 (*FaKSL2*); PQ433330 (*FaKSL3*); PQ433331 (*FaKSL4*); PQ433332 (*FaTPS7*); PQ433333 (*FaTPS10*); PQ433334 (*FaTPS17*); PQ433335 (*FaTPS18*); PQ433336 (*FaTPS35*); PQ433337 (*FaTPS36*); PQ433338 (*FaTPS42*); PQ433339 (*FaTPS55*).

## Acknowledgements

We are grateful for the outstanding support of the field superintendents and staff at the UC Davis Wolfskill Experiment Orchard (Winters, CA) and UC Davis Department of Plant Sciences (Davis, CA) and the production managers and farm managers and field staff at Mar Vista Farms (Santa Maria, CA). We further thank Dr. Reuben Peters (Iowa State University) for providing the pIRS and pACYC plasmids. We also thank Anna Cowie and Gabby Wyatt for assistance analyzing the diterpene data and Mishi Vachev for help with synteny analysis. This research was supported by a California Strawberry Commission grant (#A19-3961-001) to P.Z. and S.J.K., grants to S.J.K from the United States Department of Agriculture (USDA; http://dx.doi.org/10.13039/100000199) National Institute of Food and Agriculture (NIFA) Specialty Crops Research Initiative (SCRI) (#2017-51181B6833), S.J.K, M.J.F., and M.B. from the USDA NIFA SCRI (#2022-51181-38328-0), and S.J.K., G.S.C., M.J.F., and M.B. from the California Strawberry Commission (http://dx.doi.org/10.13039/100006760), as well as a UC Davis Jastro-Shields Graduate Research Award to M.A.M.

## Author Contributions

P.Z. and S.J.K conceived the original research and oversaw data analysis. M.A.M. and K.L.B performed the tissue harvest and metabolite analyses. G.S.C and R.A.F maintained field and greenhouse plant material. M.A.M, R.A.F, M.B, and M.J.F. performed transcriptome studies. M.A.M. performed enzyme biochemical assays. M.A.M and P.Z. wrote the manuscript with input from all authors. All authors have read and approved the manuscript.

## Supporting Information

Additional Supporting Information may be found in the online version of this article.

**Supplemental Table 1:** Upstream Terpene biosynthetic genes of the mevalonate (MVA) and methylerythritol phosphate (MEP) pathways.

**Supplemental Table 2:** Amino acid sequence similarity matrices of identified/tested TPS and syntenic pseudogenes from synteny analysis.

**Supplemental Table 3:** Terpene synthase sequences identified in this study.

**Supplemental Table 4:** Sequences of tested TPS genes.

**Supplemental Table 5:** Strawberry accessions used for transcriptomics.

**Supplemental Table 6:** Amino acid sequence similarity matrices of upstream terpene biosynthetic genes.

**Supplemental Table 7:** Harvest and accession information for this study.

**Supplemental Table 8:** Volatile compound names, CAS numbers, SPME-GC-MS retention times, and compound classifiers detected in this study.

**Supplemental Table 9:** Reagents and authentic standards used in this study.

**Supplemental Data 1:** SPME-GC-MS Data from 96 field-grown accessions.

**Supplemental Data 2:** SPME-GC-MS Data from developmental time-course cultivars.

**Supplemental Data 3:** SPME-GC-MS Data from 17 accessions harvested and flash frozen in the field for transcriptomic analysis.

**Supplemental Figure 1:** Genome localization of functionally tested and syntenic TPS genes and upstream genes of the MVA and MEP terpene metabolism pathway.

**Supplemental Figure 2:** Microsynteny plots of diploid *F. vesca* and FaRR1 Royal Royce of Mevalonate pathway genes.

**Supplemental Figure 3:** Microsynteny plots of diploid *F. vesca* and FaRR1 Royal Royce of MEP pathway genes.

**Supplemental Figure 4:** Microsynteny plots of diploid *F. vesca* and FaRR1 Royal Royce of prenyltransferases.

**Supplemental Figure 5:** Amino acid sequence alignments of the seven putatively misassembled TPS sequences.

**Supplemental Figure 6:** Synteny plots of all 7 FaRR1 chromosomes with TPS genes identified in the diploid *F. vesca* genome.

**Supplemental Figure 7:** Microsynteny plots of diploid *F. vesca* and FaRR1 Royal Royce of diterpene synthases.

**Supplemental Figure 8:** Microsynteny plots of diploid *F. vesca* and FaRR1 Royal Royce of all functionally analyzed mono-, sesqui-terpene synthases.

**Supplemental Figure 9:** Sequence alignment of diterpene candidates.

**Supplemental Figure 10:** Functional characterization of terpene synthase (TPS) products resulting from either *in vitro* enzyme assays of individual recombinant TPSs with GPP as a substrate or co-expression assays of individual TPSs and a GPP synthase in *E. coli*.

**Supplemental Figure 11:** Functional characterization of TPS-g and TPS-b products resulting from either *in vitro* enzyme assays of individual recombinant TPSs with FPP as a substrate or co-expression assays of individual TPSs and a FPP synthase in *E. coli*.

**Supplemental Figure 12:** Functional characterization of TPS-a products resulting from either *in vitro* enzyme assays of individual recombinant TPSs with FPP as a substrate or co-expression assays of individual TPSs and a FPP synthase in *E. coli*.

**Supplemental Figure 13:** Functional characterization of diterpene synthase products resulting from either *in vitro* enzyme assays of individual recombinant TPSs with GGPP as a substrate or co-expression assays of individual TPSs and a GGPP synthase in *E. coli*.

**Supplemental Figure 14:** Total SPME-GC-MS chromatograms of a sample from each of the 4 *Fragaria* species represented in this study and identified terpenes and associated spectra.

**Supplemental Figure 15:** Terpene metabolite and underlying gene abundance in field-grown strawberry fruits which were flash frozen immediately after harvest for transcriptomic analysis.

**Supplemental Figure 16:** Hierarchical cluster analysis of select upstream terpene biosynthetic and terpene synthase (TPS) genes analyzed from various developmental stages.

## Literature Cited

1. Aharoni A, Giri AP, Verstappen FWA, Bertea CM, Sevenier R, Sun Z, Jongsma MA, Schwab W, Bouwmeester HJ (2004) Gain and loss of fruit flavor compounds produced by wild and cultivated strawberry species. Plant Cell 16: 3110–3131

2. Alexa A, Rahnenfuhrer J (2022) Gene set enrichment analysis with topGO.

3. Andrews S (2010) FastQC A quality control tool for high throughput sequence data.

4. Barbey CR, Hogshead MH, Harrison B, Schwartz AE, Verma S, Oh Y, Lee S, Folta KM, Whitaker VM (2021) Genetic Analysis of Methyl Anthranilate, Mesifurane, Linalool, and Other Flavor Compounds in Cultivated Strawberry (Fragaria × ananassa). Front Plant Sci 12: 615749

5. Beekwilder J, Alvarez-Huerta M, Neef E, Verstappen FWA, Bouwmeester HJ, Aharoni A (2004) Functional characterization of enzymes forming volatile esters from strawberry and banana. Plant Physiol 135: 1865–1878

6. Bianchi G, Lovazzano A, Lanubile A, Marocco A (2014) Aroma Quality of Fruits of Wild and Cultivated Strawberry (FRAGARIA SPP.) in Relation to the Flavour-Related Gene Expression. Journal of Horticultural Research 22: 77–84

7. Blanco-Ulate B, Vincenti E, Powell ALT, Cantu D (2013) Tomato transcriptome and mutant analyses suggest a role for plant stress hormones in the interaction between fruit and Botrytis cinerea. Front Plant Sci 4: 142

8. Bood KG, Zabetakis I (2002) The biosynthesis of strawberry flavor (II): Biosynthetic and molecular biology studies. J Food Sci 67: 2–8

9. Chambers A, Whitaker VM, Gibbs B, Plotto A, Folta KM (2012) Detection of the linalool producing *NES1* variant across diverse strawberry (*Fragaria* spp.) accessions. Plant Breed 131: 437–443

10. Chen F, Tholl D, Bohlmann J, Pichersky E (2011) The family of terpene synthases in plants: a mid-size family of genes for specialized metabolism that is highly diversified throughout the kingdom. Plant J 66: 212–229

11. Criscuolo A, Gribaldo S (2010) BMGE (Block Mapping and Gathering with Entropy): a new software for selection of phylogenetic informative regions from multiple sequence alignments. BMC Evol Biol 10: 210

12. Cruz-Rus E, Sesmero R, Ángel-Pérez JA, Sánchez-Sevilla JF, Ulrich D, Amaya I (2017) Validation of a PCR test to predict the presence of flavor volatiles mesifurane and γ-decalactone in fruits of cultivated strawberry (Fragaria × ananassa). Mol Breed 37: 131

13. Csukasi F, Osorio S, Gutierrez JR, Kitamura J, Giavalisco P, Nakajima M, Fernie AR, Rathjen JP, Botella MA, Valpuesta V, et al (2011) Gibberellin biosynthesis and signalling during development of the strawberry receptacle. New Phytol 191: 376–390

14. Cumplido-Laso G, Medina-Puche L, Moyano E, Hoffmann T, Sinz Q, Ring L, Studart-Wittkowski C, Caballero JL, Schwab W, Muñoz-Blanco J, et al (2012) The fruit ripening-related gene FaAAT2 encodes an acyl transferase involved in strawberry aroma biogenesis. J Exp Bot 63: 4275–4290

15. Cyr A, Wilderman PR, Determan M, Peters RJ (2007) A modular approach for facile biosynthesis of labdane-related diterpenes. J Am Chem Soc 129: 6684–6685

16. Darrow GM (1966) The Strawberry: History, Breeding, and Physiology.

17. Dobin A, Davis CA, Schlesinger F, Drenkow J, Zaleski C, Jha S, Batut P, Chaisson M, Gingeras TR (2013) STAR: ultrafast universal RNA-seq aligner. Bioinformatics 29: 15–21

18. Duan W, Sun P, Chen L, Gao S, Shao W, Li J (2018) Comparative analysis of fruit volatiles and related gene expression between the wild strawberry Fragaria pentaphylla and cultivated Fragaria × ananassa. Eur Food Res Technol 244: 57–72

19. Edgar RC (2004) MUSCLE: multiple sequence alignment with high accuracy and high throughput. Nucleic Acids Res 32: 1792–1797

20. Edger PP, Poorten TJ, VanBuren R, Hardigan MA, Colle M, McKain MR, Smith RD, Teresi SJ, Nelson ADL, Wai CM, et al (2019) Origin and evolution of the octoploid strawberry genome. Nat Genet 51: 541–547

21. Edger PP, VanBuren R, Colle M, Poorten TJ, Wai CM, Niederhuth CE, Alger EI, Ou S, Acharya CB, Wang J, et al (2018) Single-molecule sequencing and optical mapping yields an improved genome of woodland strawberry (Fragaria vesca) with chromosome-scale contiguity. Gigascience 7: 1–7

22. Fan Z, Hasing T, Johnson TS, Garner DM, Schwieterman ML, Barbey CR, Colquhoun TA, Sims CA, Resende MFR, Whitaker VM (2021a) Strawberry sweetness and consumer preference are enhanced by specific volatile compounds. Hortic Res 8: 66

23. Fan Z, Plotto A, Bai J, Whitaker VM (2021b) Volatiles Influencing Sensory Attributes and Bayesian Modeling of the Soluble Solids-Sweetness Relationship in Strawberry. Front Plant Sci 12: 640704

24. Fan Z, Tieman DM, Knapp SJ, Zerbe P, Famula R, Barbey CR, Folta KM, Amadeu RR, Lee M, Oh Y, et al (2022) A multi-omics framework reveals strawberry flavor genes and their regulatory elements. New Phytol 236: 1089–1107

25. Fao Food And Agriculture Organization Staff (2000) Faostat: FAO Statistical Databases.

26. Feldmann MJ, Pincot DDA, Cole GS, Knapp SJ (2024) Genetic gains underpinning a little-known strawberry Green Revolution. Nat Commun 15: 2468

27. Grabherr MG, Haas BJ, Yassour M, Levin JZ, Thompson DA, Amit I, Adiconis X, Fan L, Raychowdhury R, Zeng Q, et al (2011) Full-length transcriptome assembly from RNA-Seq data without a reference genome. Nat Biotechnol 29: 644–652

28. Guindon S, Dufayard J-F, Lefort V, Anisimova M, Hordijk W, Gascuel O (2010) New algorithms and methods to estimate maximum-likelihood phylogenies: assessing the performance of PhyML 3.0. Syst Biol 59: 307–321

29. Haas BJ, Papanicolaou A, Yassour M, Grabherr M, Blood PD, Bowden J, Couger MB, Eccles D, Li B, Lieber M, et al (2013) De novo transcript sequence reconstruction from RNA-seq using the Trinity platform for reference generation and analysis. Nat Protoc 8: 1494–1512

30. Hardigan MA, Feldmann MJ, Pincot DDA, Famula RA, Vachev MV, Madera MA, Zerbe P, Mars K, Peluso P, Rank D, et al (2021a) Blueprint for Phasing and Assembling the Genomes of Heterozygous Polyploids: Application to the Octoploid Genome of Strawberry. Genomics

31. Hardigan MA, Lorant A, Pincot DDA, Feldmann MJ, Famula RA, Acharya CB, Lee S, Verma S, Whitaker VM, Bassil N, et al (2021b) Unraveling the Complex Hybrid Ancestry and Domestication History of Cultivated Strawberry. Mol Biol Evol 38: 2285– 2305

32. Ilc T, Werck-Reichhart D, Navrot N (2016) Meta-Analysis of the Core Aroma Components of Grape and Wine Aroma. Front Plant Sci 7: 1472

33. Jetti RR, Yang E, Kurnianta A, Finn C, Qian MC (2007) Quantification of selected aroma-active compounds in strawberries by headspace solid-phase microextraction gas chromatography and correlation with sensory descriptive analysis. J Food Sci 72: S487–96

34. Jiménez NP, Bjornson M, Famula RA, Pincot DDA, Hardigan MA, Madera MA, Lopez Ramirez CM, Cole GS, Feldmann MJ, Knapp SJ (2024) Loss-of-function mutations in the fruit softening gene *POLYGALACTURONASE1* doubled fruit firmness in strawberry. Hortic Res. doi: 10.1093/hr/uhae315

35. Johnson SR, Bhat WW, Bibik J, Turmo A, Hamberger B, Evolutionary Mint Genomics C, Hamberger B (2019a) A database-driven approach identifies additional diterpene synthase activities in the mint family (Lamiaceae). J Biol Chem 294: 1349–1362

36. Johnson SR, Bhat WW, Sadre R, Miller GP, Garcia AS, Hamberger B (2019b) Promiscuous terpene synthases from Prunella vulgaris highlight the importance of substrate and compartment switching in terpene synthase evolution. New Phytol. doi: 10.1111/nph.15778

37. Junier T, Zdobnov EM (2010) The Newick utilities: high-throughput phylogenetic tree processing in the UNIX shell. Bioinformatics 26: 1669–1670

38. Karunanithi PS, Zerbe P (2019) Terpene Synthases as Metabolic Gatekeepers in the Evolution of Plant Terpenoid Chemical Diversity. Front Plant Sci 10: 1166

39. Köllner TG, Gershenzon J, Peters RJ, Zerbe P, Schmelz EA (2023) The terpene synthase gene family in maize – a clarification of existing community nomenclature. BMC Genomics 24: 744

40. Kulheim C, Padovan A, Hefer C, Krause ST, Kollner TG, Myburg AA, Degenhardt J, Foley WJ (2015) The Eucalyptus terpene synthase gene family. BMC Genomics 16: 450

41. Langmead B, Salzberg SL (2012) Fast gapped-read alignment with Bowtie 2. Nat Methods 9: 357–359

42. Lefort V, Longueville J-E, Gascuel O (2017) SMS: Smart Model Selection in PhyML. Mol Biol Evol 34: 2422–2424

43. Lemoine F, Correia D, Lefort V, Doppelt-Azeroual O, Mareuil F, Cohen-Boulakia S, Gascuel O (2019) NGPhylogeny.fr: new generation phylogenetic services for non-specialists. Nucleic Acids Res 47: W260–W265

44. Lemoine F, Domelevo Entfellner J-B, Wilkinson E, Correia D, Dávila Felipe M, De Oliveira T, Gascuel O (2018) Renewing Felsenstein’s phylogenetic bootstrap in the era of big data. Nature 556: 452–456

45. Liao X, Li M, Liu B, Yan M, Yu X, Zi H, Liu R, Yamamuro C (2018) Interlinked regulatory loops of ABA catabolism and biosynthesis coordinate fruit growth and ripening in woodland strawberry. Proc Natl Acad Sci U S A 115: E11542–E11550

46. Liaw A, Wiener M (2002) Classification and Regression by randomForest.

47. Martin DM, Aubourg S, Schouwey MB, Daviet L, Schalk M, Toub O, Lund ST, Bohlmann J (2010) Functional annotation, genome organization and phylogeny of the grapevine (Vitis vinifera) terpene synthase gene family based on genome assembly, FLcDNA cloning, and enzyme assays. BMC Plant Biol 10: 226

48. Martin DM, Faldt J, Bohlmann J (2004) Functional characterization of nine Norway Spruce TPS genes and evolution of gymnosperm terpene synthases of the TPS-d subfamily. Plant Physiol 135: 1908–1927

49. Morrone D, Lowry L, Determan MK, Hershey DM, Xu M, Peters RJ (2010) Increasing diterpene yield with a modular metabolic engineering system in E. coli: comparison of MEV and MEP isoprenoid precursor pathway engineering. Appl Microbiol Biotechnol 85: 1893–1906

50. Muchlinski A, Jia M, Tiedge K, Fell JS, Pelot KA, Chew L, Davisson D, Chen Y, Siegel J, Lovell JT, et al (2021) Cytochrome P450-catalyzed biosynthesis of furanoditerpenoids in the bioenergy crop switchgrass (Panicum virgatum L.). Plant J 108: 1053–1068

51. Murphy KM, Chung S, Fogla S, Minsky HB, Zhu KY, Zerbe P (2019) A Customizable Approach for the Enzymatic Production and Purification of Diterpenoid Natural Products. J Vis Exp. doi: 10.3791/59992

52. Nawade B, Yahyaa M, Reuveny H, Shaltiel-Harpaz L, Eisenbach O, Faigenboim A, Bar-Yaakov I, Holland D, Ibdah M (2019) Profiling of volatile terpenes from almond (Prunus dulcis) young fruits and characterization of seven terpene synthase genes. Plant Sci 287: 110187

53. Negri AS, Allegra D, Simoni L, Rusconi F, Tonelli C, Espen L, Galbiati M (2015) Comparative analysis of fruit aroma patterns in the domesticated wild strawberries â€œProfumata di Tortonaâ€ (F. moschata) and â€œRegina delle Valliâ€ (F. vesca). Front Plant Sci 6: 56

54. Nieuwenhuizen NJ, Green SA, Chen X, Bailleul EJD, Matich AJ, Wang MY, Atkinson RG (2013) Functional genomics reveals that a compact terpene synthase gene family can account for terpene volatile production in apple. Plant Physiol 161: 787–804

55. Njuguna W, Liston A, Cronn R, Ashman T-L, Bassil N (2013) Insights into phylogeny, sex function and age of Fragaria based on whole chloroplast genome sequencing. Mol Phylogenet Evol 66: 17–29

56. Oh Y, Barbey CR, Chandra S, Bai J, Fan Z, Plotto A, Pillet J, Folta KM, Whitaker VM, Lee S (2021) Genomic Characterization of the Fruity Aroma Gene, FaFAD1, Reveals a Gene Dosage Effect on γ-Decalactone Production in Strawberry (Fragaria × ananassa). Front Plant Sci 12: 639345

57. Oróstica KY, Verdugo RA (2016) chromPlot: visualization of genomic data in chromosomal context. Bioinformatics 32: 2366–2368

58. Pelot KA, Chen R, Hagelthorn DM, Young CA, Addison JB, Muchlinski A, Tholl D, Zerbe P (2018) Functional diversity of diterpene synthases in the biofuel crop switchgrass. Plant Physiol. doi: 10.1104/pp.18.00590

59. Pelot KA, Hagelthorn DM, Hong YJ, Tantillo DJ, Zerbe P (2019) Diterpene Synthase-Catalyzed Biosynthesis of Distinct Clerodane Stereoisomers. Chembiochem 20: 111–117

60. Pelot KA, Mitchell R, Kwon M, Hagelthorn LM, Wardman JF, Chiang A, Bohlmann J, Ro D-K, Zerbe P (2017) Biosynthesis of the psychotropic plant diterpene salvinorin A: Discovery and characterization of the Salvia divinorum clerodienyl diphosphate synthase. Plant J 89: 885–897

61. Perry M (2024) heatmaps: Flexible Heatmaps for Functional Genomics and Sequence Features. doi: 10.18129/B9.BIOC.HEATMAPS

62. Petersen KR, Streett DA, Gerritsen AT, Hunter SS, Settles ML (2015) Super deduper, fast PCR duplicate detection in fastq files. Proceedings of the 6th ACM Conference on Bioinformatics, Computational Biology and Health Informatics. Association for Computing Machinery, New York, NY, USA, pp 491–492

63. Peters RJ (2010) Two rings in them all: the labdane-related diterpenoids. Nat Prod Rep 27: 1521–1530

64. Petrasch S, Mesquida-Pesci SD, Pincot DDA, Feldmann MJ, López CM, Famula R, Hardigan MA, Cole GS, Knapp SJ, Blanco-Ulate B (2022) Genomic prediction of strawberry resistance to postharvest fruit decay caused by the fungal pathogen Botrytis cinerea. G3. doi: 10.1093/g3journal/jkab378

65. Pillet J, Chambers AH, Barbey C, Bao Z, Plotto A, Bai J, Schwieterman M, Johnson T, Harrison B, Whitaker VM, et al (2017) Identification of a methyltransferase catalyzing the final step of methyl anthranilate synthesis in cultivated strawberry. BMC Plant Biol 17: 147

66. Porter M, Fan Z, Lee S, Whitaker VM (2023) Strawberry breeding for improved flavor. Crop Sci 63: 1949–1963

67. Potter KC, Jia M, Hong YJ, Tantillo D, Peters RJ (2016) Product Rearrangement from Altering a Single Residue in the Rice syn-Copalyl Diphosphate Synthase. Org Lett 18: 1060–1063

68. Potter K, Criswell J, Zi J, Stubbs A, Peters RJ (2014) Novel product chemistry from mechanistic analysis of ent-copalyl diphosphate synthases from plant hormone biosynthesis. Angew Chem Int Ed Engl 53: 7198–7202

69. Pyysalo T, Honkanen E, Hirvi T (1979) Volatiles of wild strawberries, Fragaria vesca L., compared to those of cultivated berries, Fragaria.times. ananassa cv Senga Sengana. J Agric Food Chem 27: 19–22

70. Raab T, López-Ráez JA, Klein D, Caballero JL, Moyano E, Schwab W, Muñoz-Blanco J (2006) FaQR, required for the biosynthesis of the strawberry flavor compound 4-hydroxy-2,5-dimethyl-3(2H)-furanone, encodes an enone oxidoreductase. Plant Cell 18: 1023–1037

71. R Core Team (2021) R: A language and environment for statistical computing. https://www.R-project.org/.

72. Robinson MD, McCarthy DJ, Smyth GK (2010) edgeR: a Bioconductor package for differential expression analysis of digital gene expression data. Bioinformatics 26: 139–140

73. Sargent DJ, Yang Y, Šurbanovski N, Bianco L, Buti M, Velasco R, Giongo L, Davis TM (2016) HaploSNP affinities and linkage map positions illuminate subgenome composition in the octoploid, cultivated strawberry (Fragaria×ananassa). Plant Sci 242: 140–150

74. Schwieterman ML, Colquhoun TA, Jaworski EA, Bartoshuk LM, Gilbert JL, Tieman DM, Odabasi AZ, Moskowitz HR, Folta KM, Klee HJ, et al (2014) Strawberry flavor: diverse chemical compositions, a seasonal influence, and effects on sensory perception. PLoS One 9: e88446

75. Session AM, Rokhsar DS (2023) Transposon signatures of allopolyploid genome evolution. Nat Commun 14: 3180

76. Tennessen JA, Govindarajulu R, Ashman T-L, Liston A (2014) Evolutionary origins and dynamics of octoploid strawberry subgenomes revealed by dense targeted capture linkage maps. Genome Biol Evol 6: 3295–3313

77. Tholl D (2015) Biosynthesis and biological functions of terpenoids in plants. Adv Biochem Eng Biotechnol 148: 63–106

78. Tiedge K, Li X, Merrill AT, Davisson D, Chen Y, Yu P, Tantillo DJ, Last RL, Zerbe P (2022) Comparative transcriptomics and metabolomics reveal specialized metabolite drought stress responses in switchgrass (Panicum virgatum). New Phytol 236: 1393–1408

79. Tiedge K, Li X, Merrill AT, Davisson D, Chen Y, Yu P, Tantillo DJ, Last RL, Zerbe P Comparative transcriptomics and metabolomics reveal specialized metabolite drought stress responses in switchgrass (Panicum virgatum L.). doi: 10.1101/2022.04.20.488672

80. Tiedge K, Muchlinski A, Zerbe P (2020) Genomics-enabled analysis of specialized metabolism in bioenergy crops: current progress and challenges. Synth Biol 5: ysaa005

81. Ulrich D, Olbricht K (2016) A search for the ideal flavor of strawberry – Comparison of consumer acceptance and metabolite patterns in Fragaria × ananassa Duch. J Appl Bot Food Qual 89: 223–234

82. Urrutia M, Rambla JL, Alexiou KG, Granell A, Monfort A (2017) Genetic analysis of the wild strawberry (Fragaria vesca) volatile composition. Plant Physiol Biochem 121: 99–117

83. Yan J-W, Ban Z-J, Lu H-Y, Li D, Poverenov E, Luo Z-S, Li L (2018) The aroma volatile repertoire in strawberry fruit: a review. J Sci Food Agric 98: 4395–4402

84. Zabetakis I, Holden M (1997) Strawberry flavour: Analysis and biosynthesis. J Sci Food Agric 74: 421–434

85. Zerbe P, Bohlmann J (2015) Plant diterpene synthases: exploring modularity and metabolic diversity for bioengineering. Trends Biotechnol 33: 419–428

86. Zerbe P, Hamberger B, Yuen MMS, Chiang A, Sandhu HK, Madilao LL, Nguyen A, Hamberger B, Bach SS, Bohlmann J (2013) Gene discovery of modular diterpene metabolism in nonmodel systems. Plant Physiol 162: 1073–1091

87. Zhang A, Xiong Y, Fang J, Jiang X, Wang T, Liu K, Peng H, Zhang X (2022) Diversity and Functional Evolution of Terpene Synthases in Rosaceae. Plants. doi: 10.3390/plants11060736

88. Zi J, Mafu S, Peters RJ (2014) To gibberellins and beyond! Surveying the evolution of (di)terpenoid metabolism. Annu Rev Plant Biol 65: 259–286

89. Zorrilla-Fontanesi Y, Rambla J-L, Cabeza A, Medina JJ, Sánchez-Sevilla JF, Valpuesta V, Botella MA, Granell A, Amaya I (2012) Genetic analysis of strawberry fruit aroma and identification of O-methyltransferase FaOMT as the locus controlling natural variation in mesifurane content. Plant Physiol 159: 851–870

