## Supplemental Tables & Figures for "Genome-Wide Discovery and Characterization of Terpene Synthases Contributing to Strawberry Aroma Metabolism"

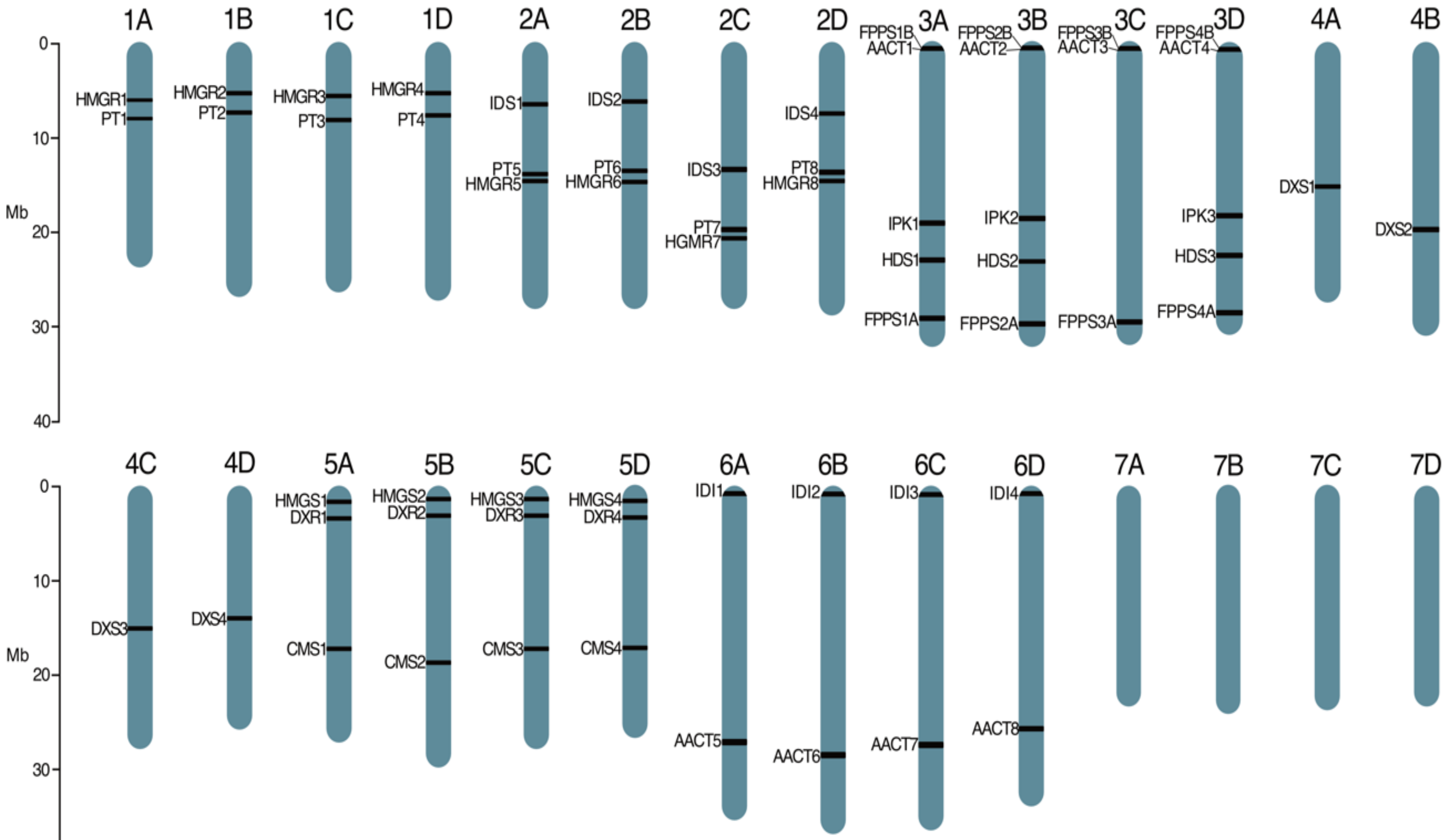
1A
1B
1C
1D
2A
2B
2C
2D
3A
3B
3C
3D
4A
4B
Mb
0
10
20
30
40
HMGR1
PT1
HMGR2
PT2
HMGR3
PT3
HMGR4
PT4
IDS1
PT5
HMGR5
IDS2
PT6
HMGR6
IDS3
PT7
HGMR7
IDS4
PT8
HMGR8
FPPS1B
AACT1
IPK1
HDS1
FPPS1A
FPPS2B
AACT2
IPK2
HDS2
FPPS2A
FPPS3B
AACT3
FPPS3A
FPPS4B
AACT4
IPK3
HDS3
FPPS4A
DXS1
DXS2
4C
4D
5A
5B
5C
5D
6A
6B
6C
6D
7A
7B
7C
7D
DXS3
DXS4
HMGS1
DXR1
CMS1
HMGS2
DXR2
CMS2
HMGS3
DXR3
CMS3
HMGS4
DXR4
CMS4
IDI1
AACT5
IDI2
AACT6
IDI3
AACT7
IDI4
AACT8

**Supplemental Fig. 1.** Genome localization of functionally tested TPS (red), genes predicted to be syntenic to functionally tested TPS genes (yellow), and upstream genes of the MVA and MEP terpene metabolism pathway (black). *AACT, Acetoacetyl-CoA Thiolase; CMS, 4-diphosphocytidyl-2-C-methyl-D-erythritol synthase; DXR, 1-deoxy-D-xylulose 5-phosphate reductoisomerase; DXS, 1-deoxy-D-xylulose 5-phosphate synthase; HDS, 4-hydroxy-3-methylbut-2-enyl diphosphate synthase; FPPS, farnesyl diphosphate synthase; HMGR, 3-hydroxy-3-methylglutaryl-CoA reductase; HMGS, 3-hydroxy-3-methylglutaryl-CoA synthase; IDI, isopentenyl diphosphate isomerase; IDS, 4-hydroxy-3-methylbut-2-enyl diphosphate reductase; IPK, isopentenyl monophosphate kinase; PT, Prenyltransferase (Geranyl Diphosphate Synthase/Geranylgeranyl diphosphate synthase).*

**A** AACT - Acetoacetyl CoA Thiolase

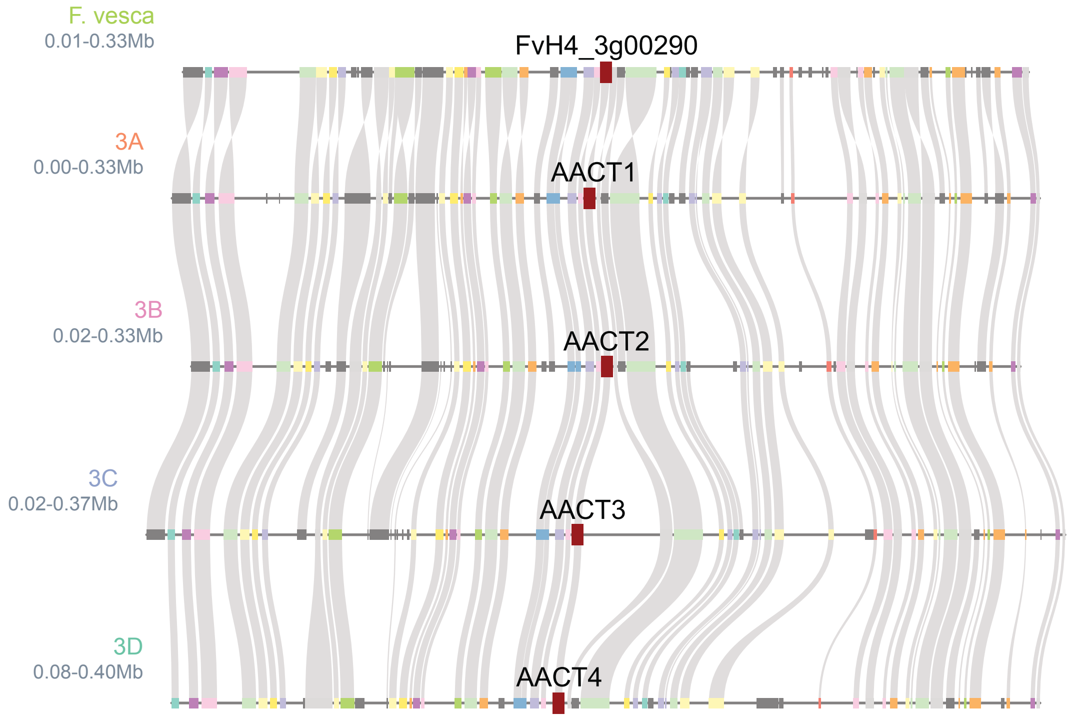

**B** AACT - Acetoacetyl CoA Thiolase

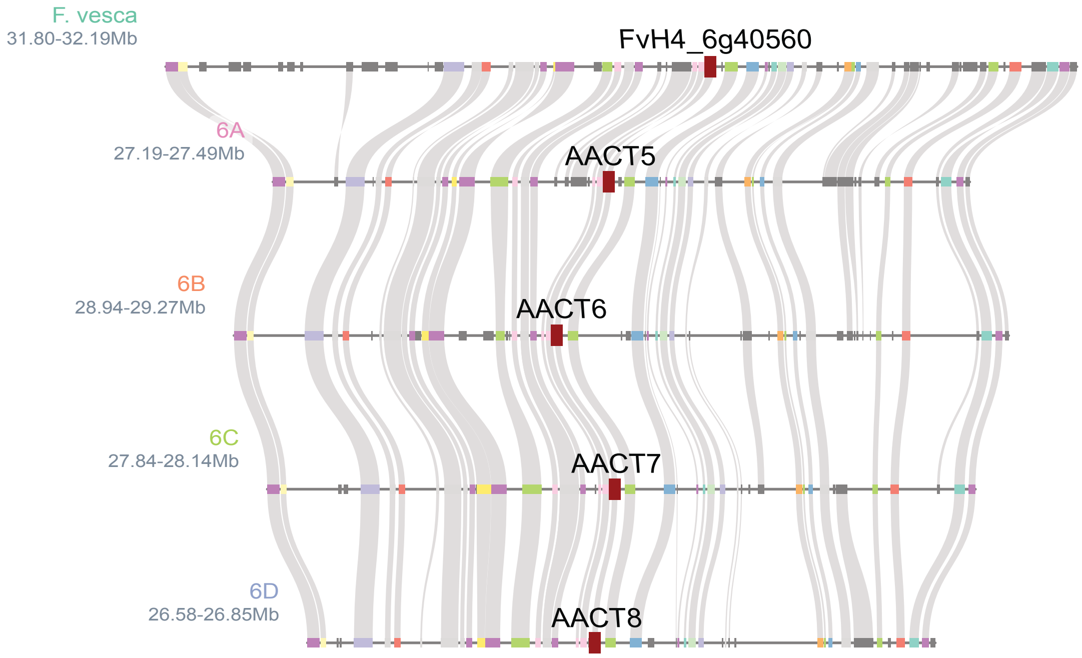

**C** HMGS - 3-hydroxy-3-methylglutaryl-CoA Synthase

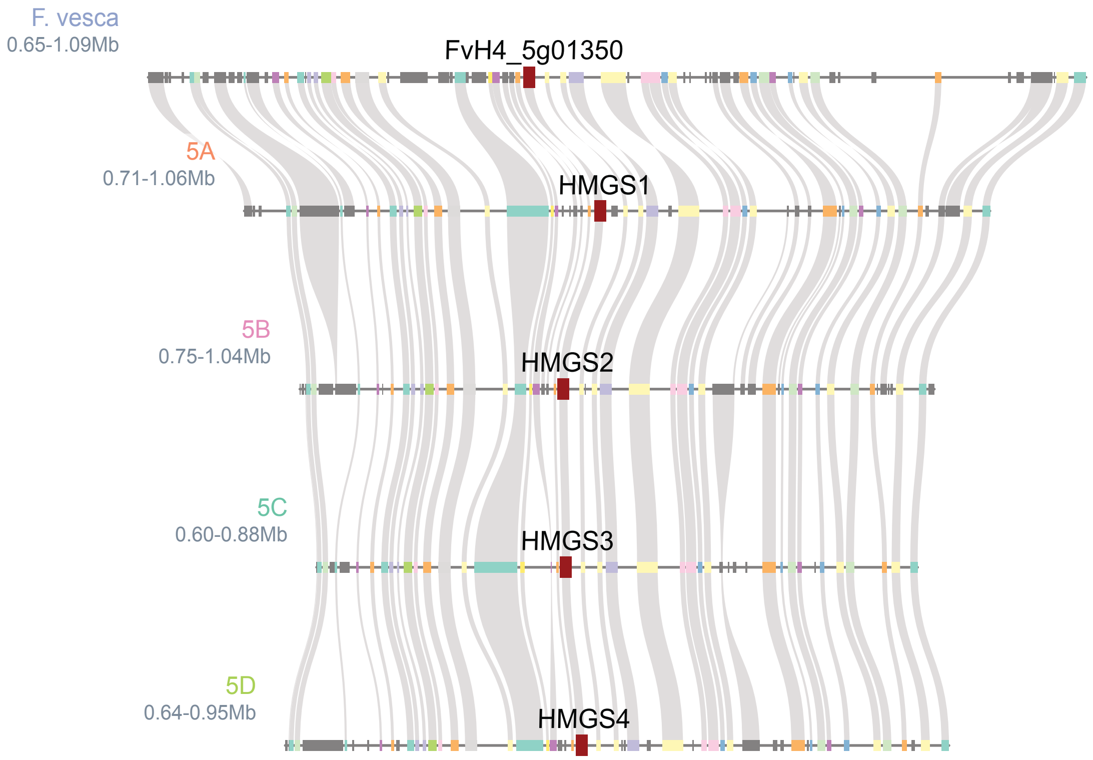

**D** HMGR - 3-hydroxy-3-methylglutaryl-CoA reductase

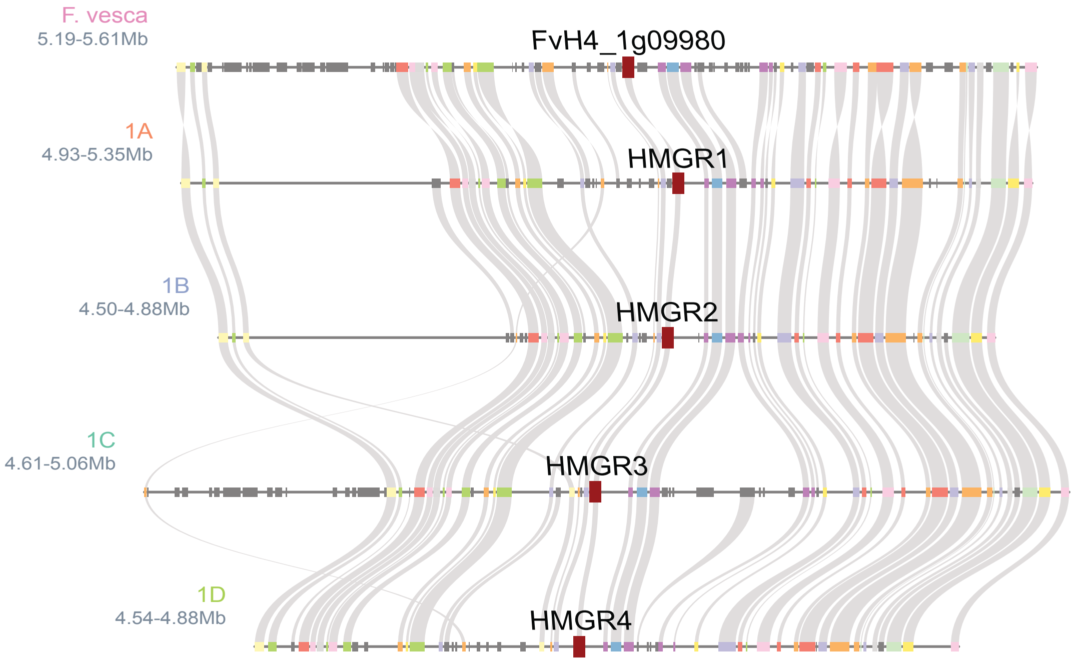

**E** HMGR - 3-hydroxy-3-methylglutaryl-CoA reductase

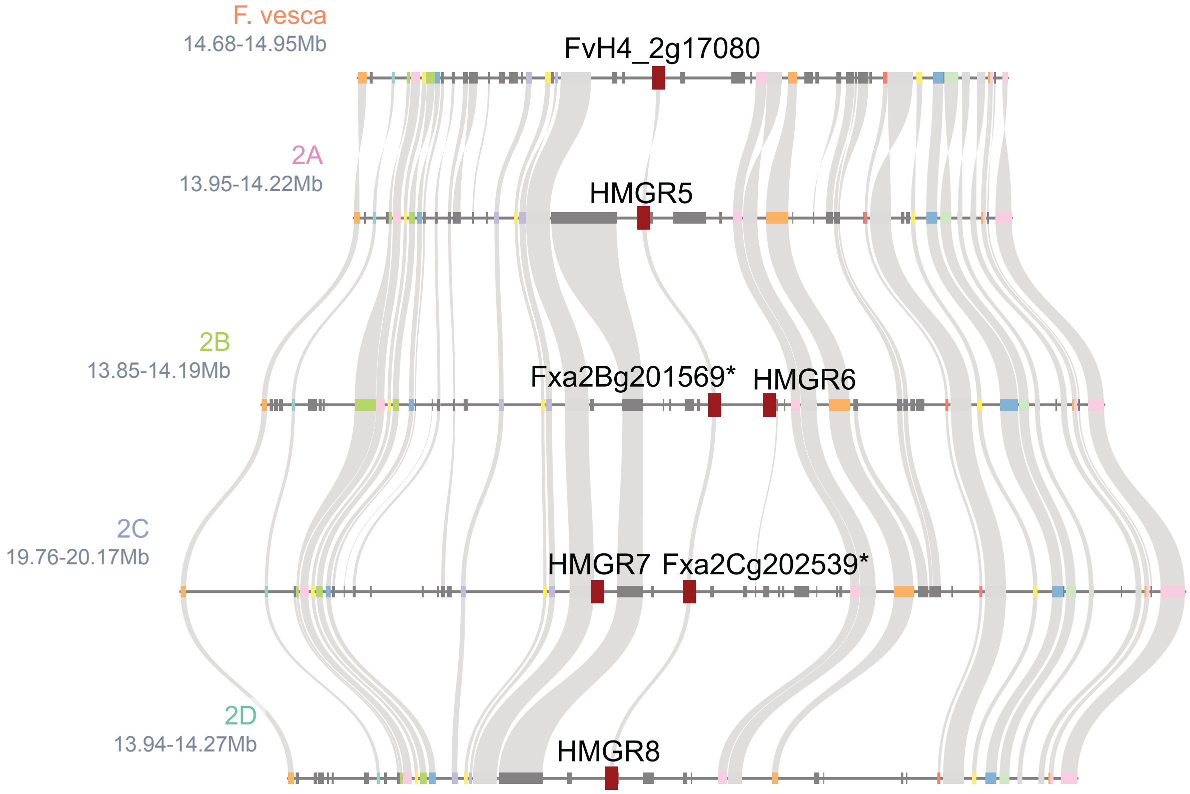

**Supplemental Fig. 2.** Microsynteny plots of diploid *F. vesca* and FaRR1 Royal Royce of Mevalonate pathway genes. Genes of interest are highlighted in red.

**A** DXS - 1-deoxy-D-xylulose 5-phosphate synthase

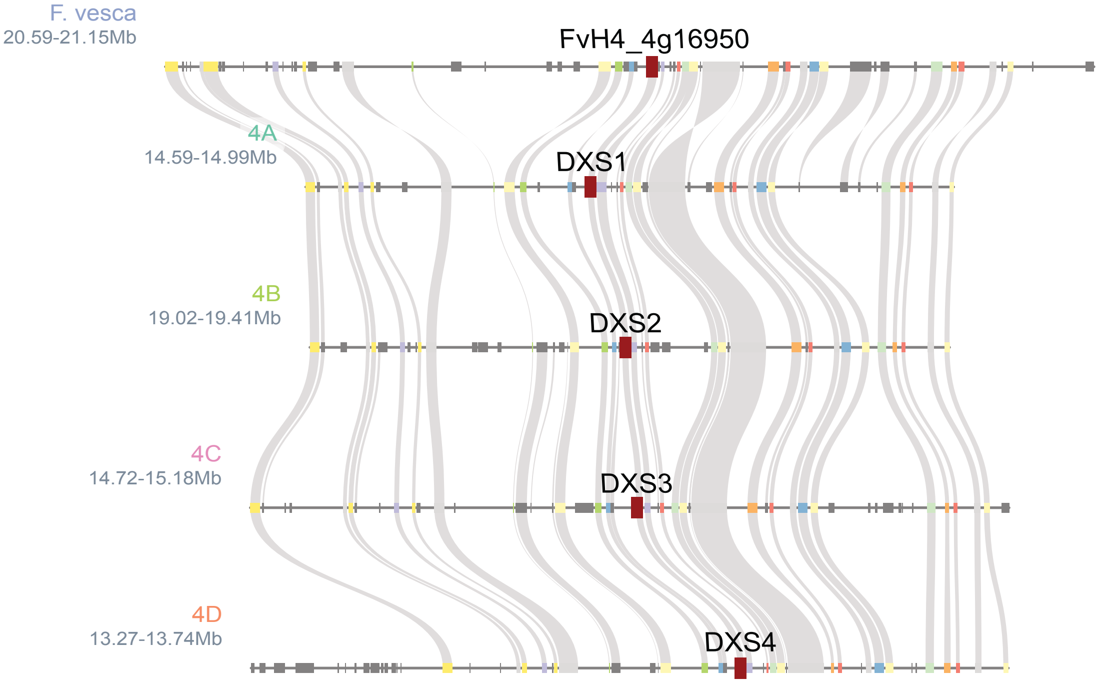

**B** DXR - 1-deoxy-D-xylulose 5-phosphate reductoisomerase

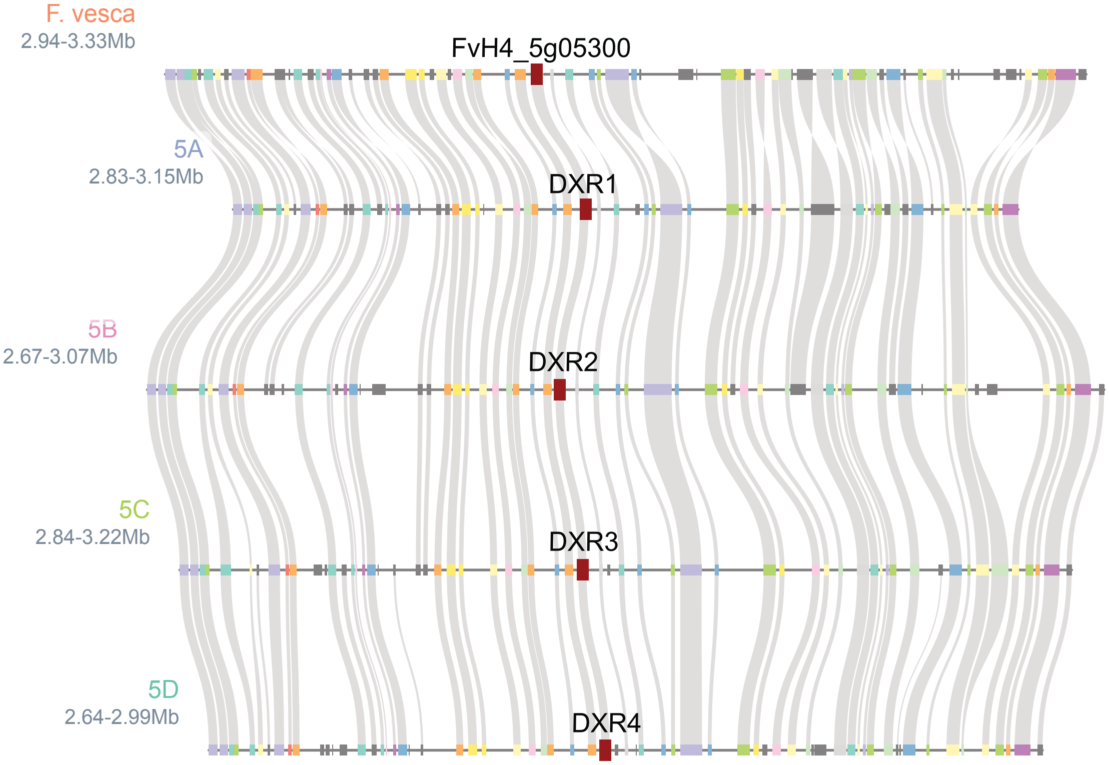

**C** CMS - 4-diphosphocytidyl-2-C-methyl-D-erythitol synthase

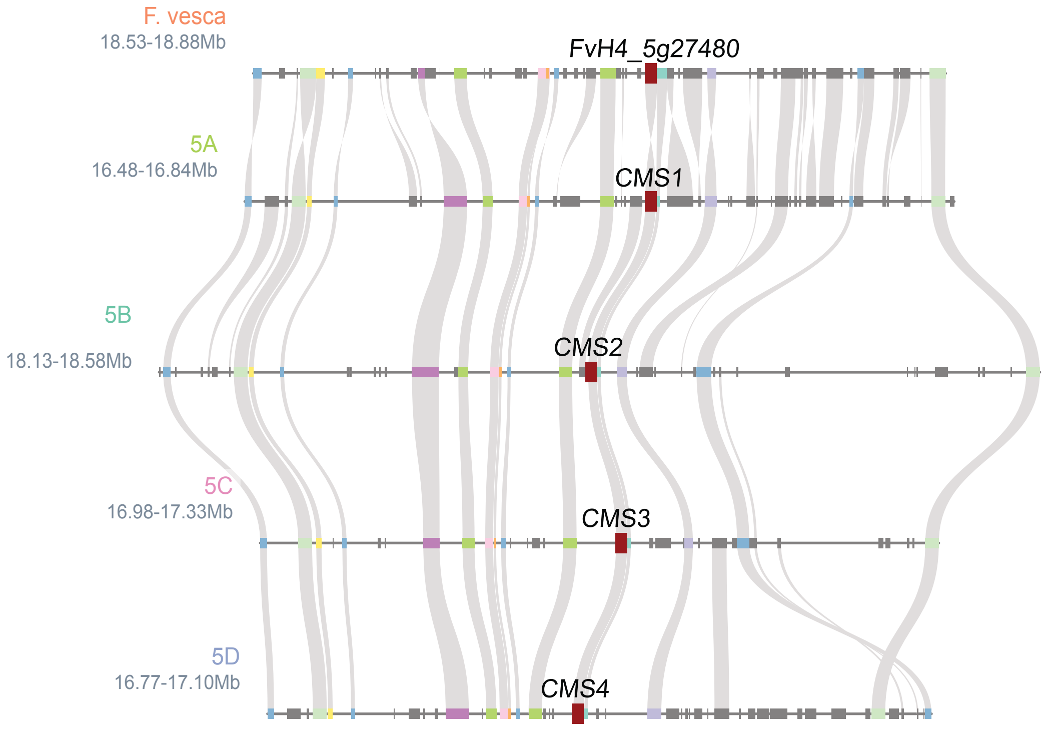

**D** HDS - 4-hydroxy-3-methylbut-2-enyl diphosphate synthase

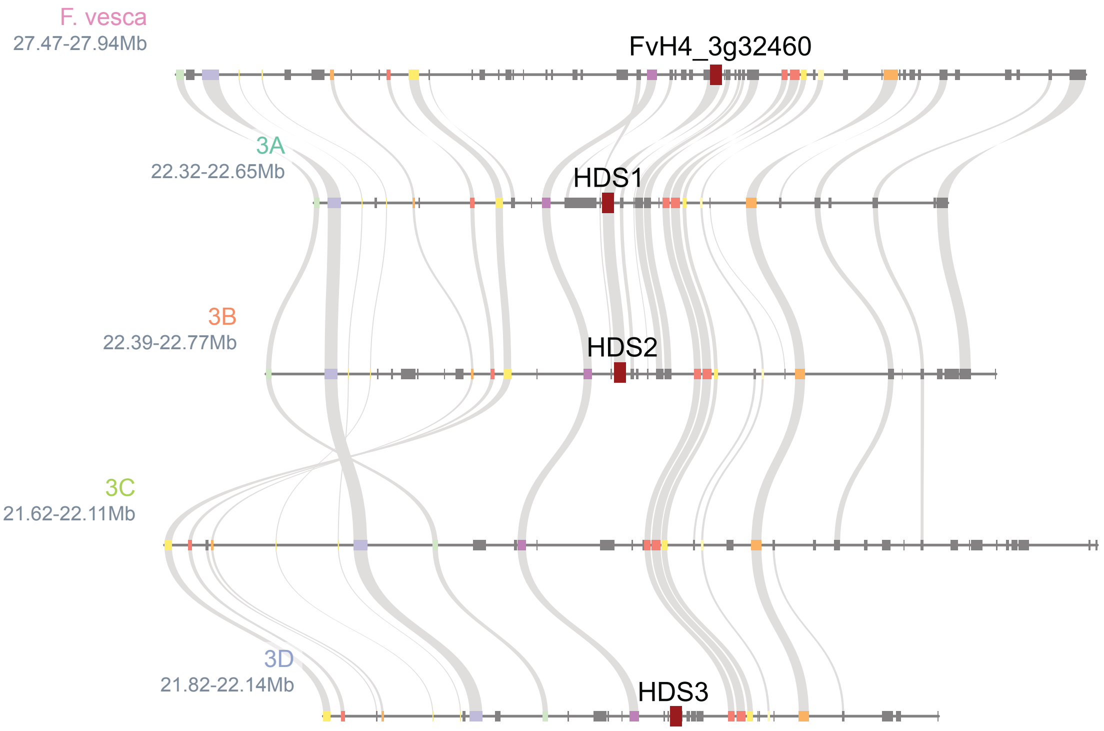

**E** HDR - 4-hydroxy-3-methylbut-2-enyl diphosphate reductase

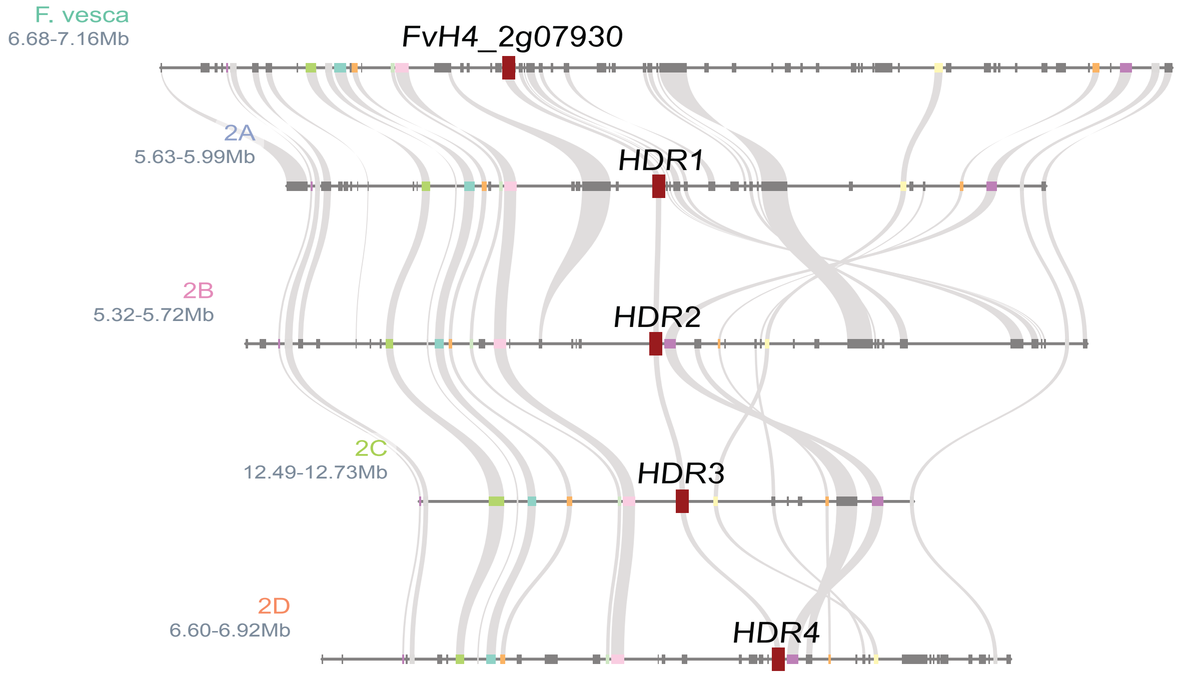

**Supplemental Fig. 3.** Microsynteny plots of diploid *F. vesca* and FaRR1 Royal Royce of MEP pathway genes. Genes of interest are highlighted in red.

**A** Geranyl/Geranylgeranyl Diphosphate Synthase

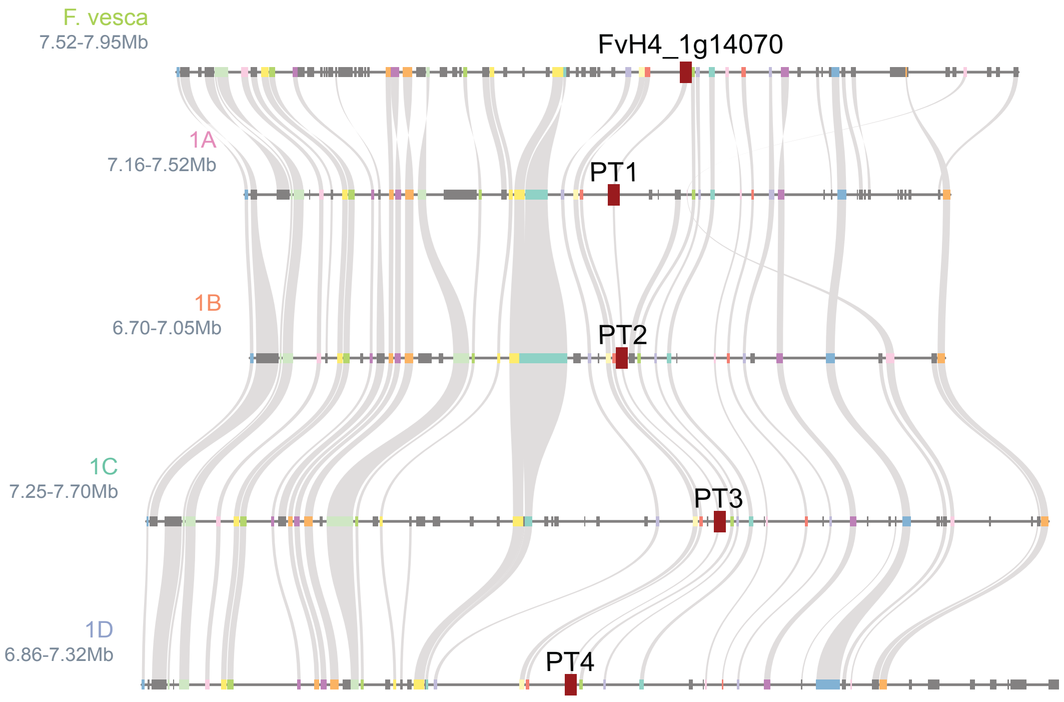

**B** Geranyl/Geranylgeranyl Diphosphate Synthase

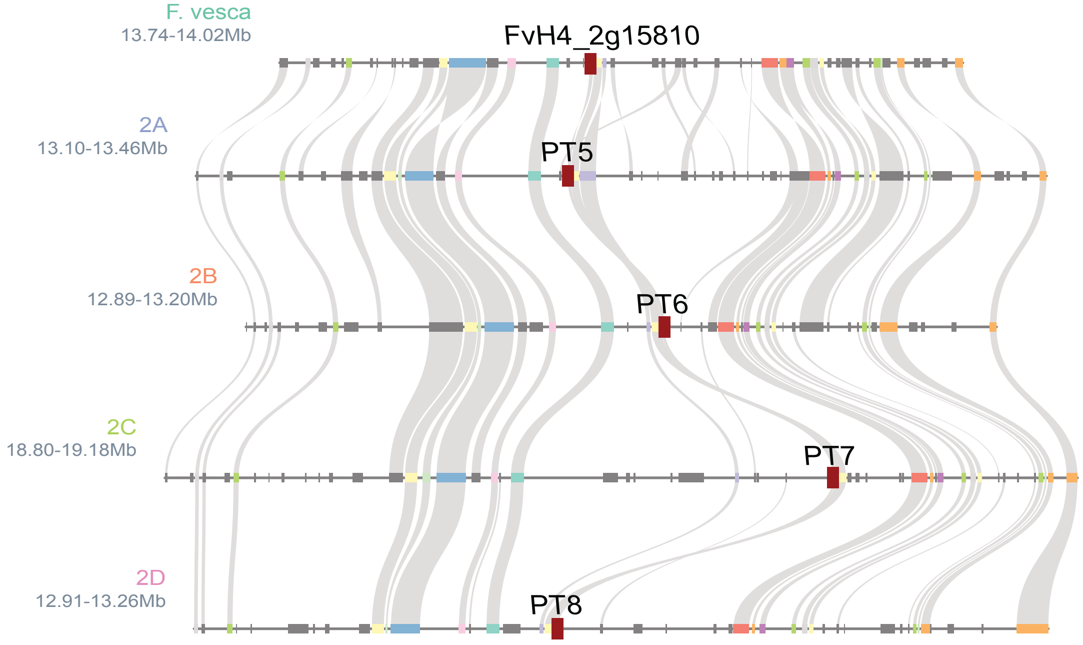

**C** FPPS - Farnesyl Diphosphate Synthase

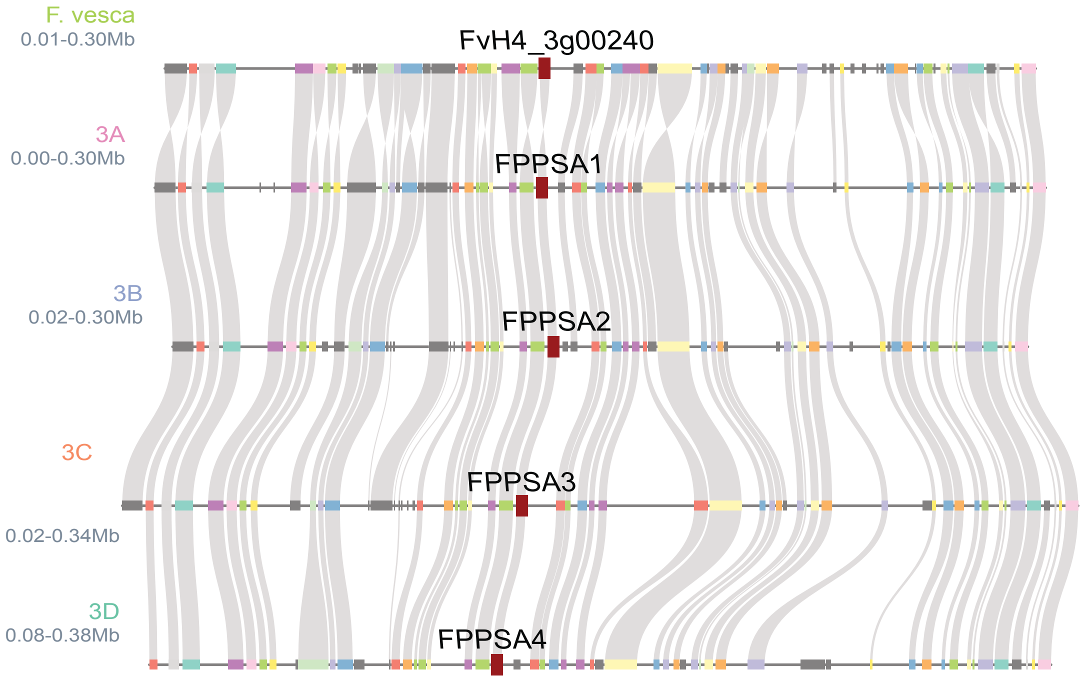

**D** FPPS - Farnesyl Diphosphate Synthase

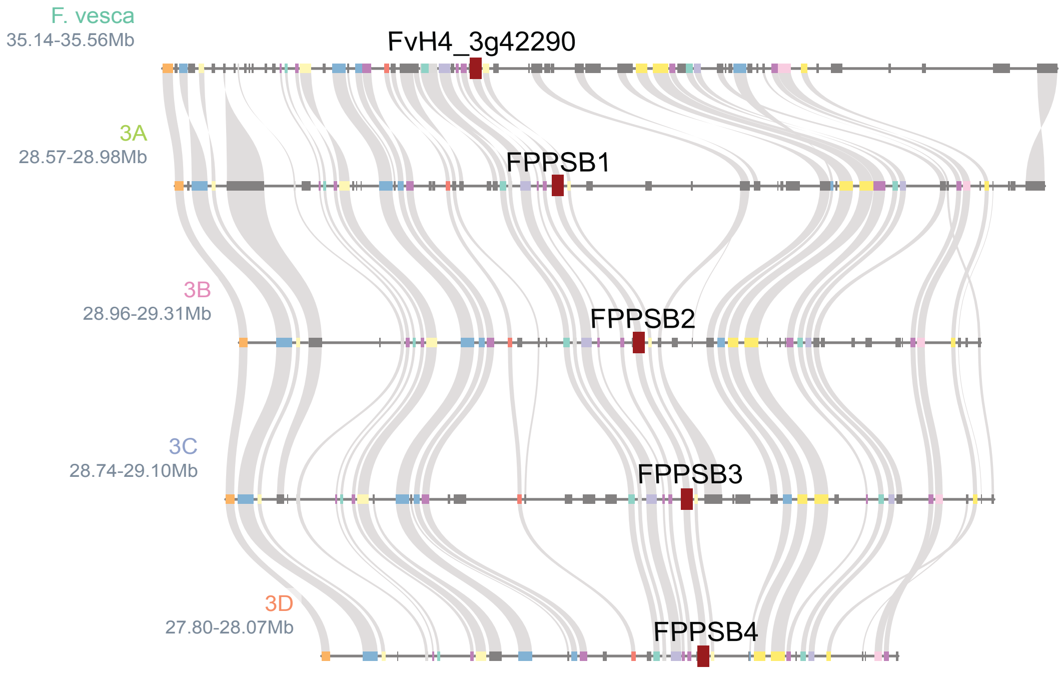

**E** IDI - Isopentenyl Diphosphate Isomerase

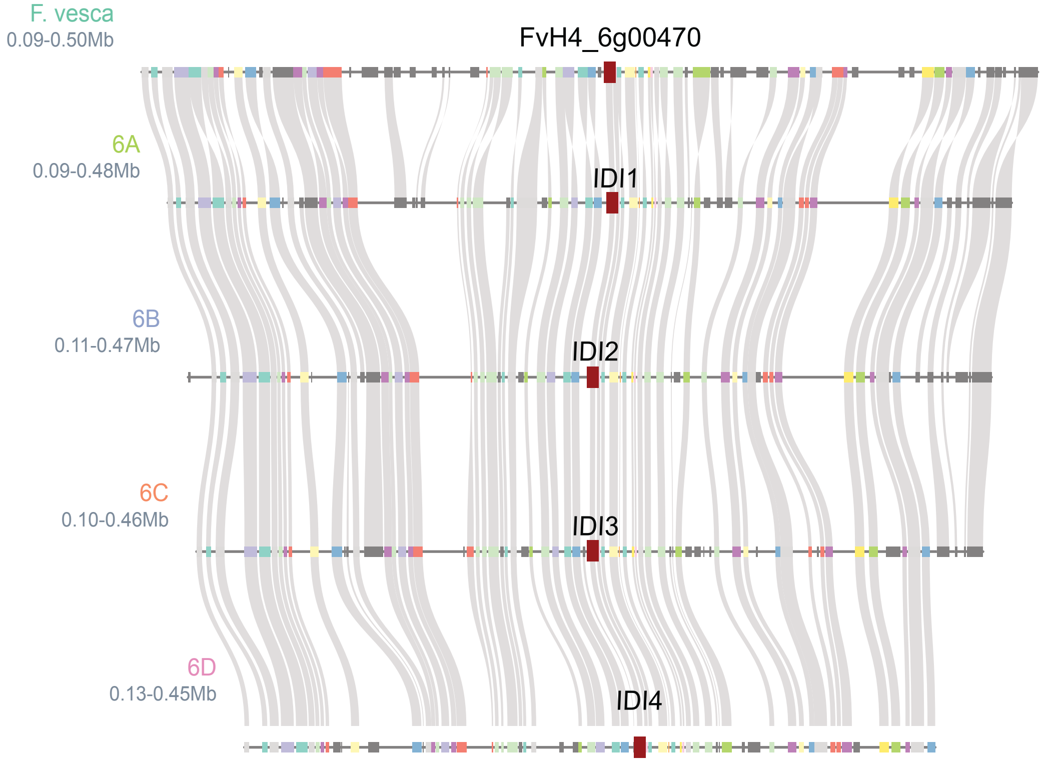

**F** IPK - Isopentenyl Monophosphate Kinase

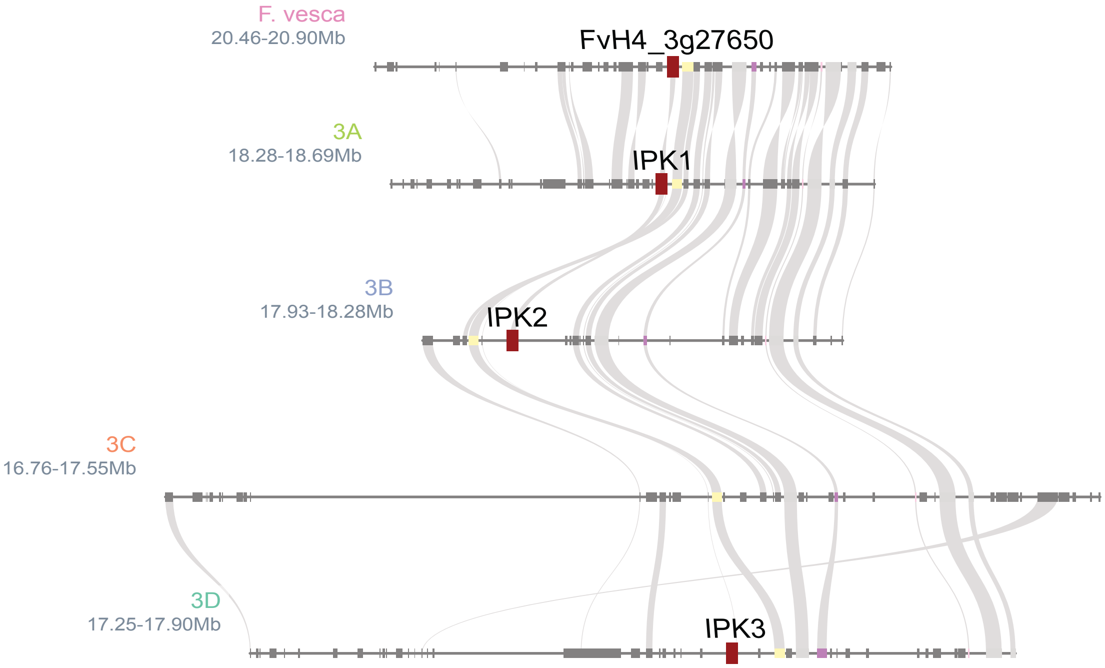

**Supplemental Fig. 4.** Microsynteny plots of diploid *F. vesca* and FaRR1 Royal Royce of (**A-D**) prenyltransferases, (**E**) Isopentenyl diphosphate isomerase and (**F**) Isopentenyl monophosphate kinase. Genes of interest are highlighted in red.

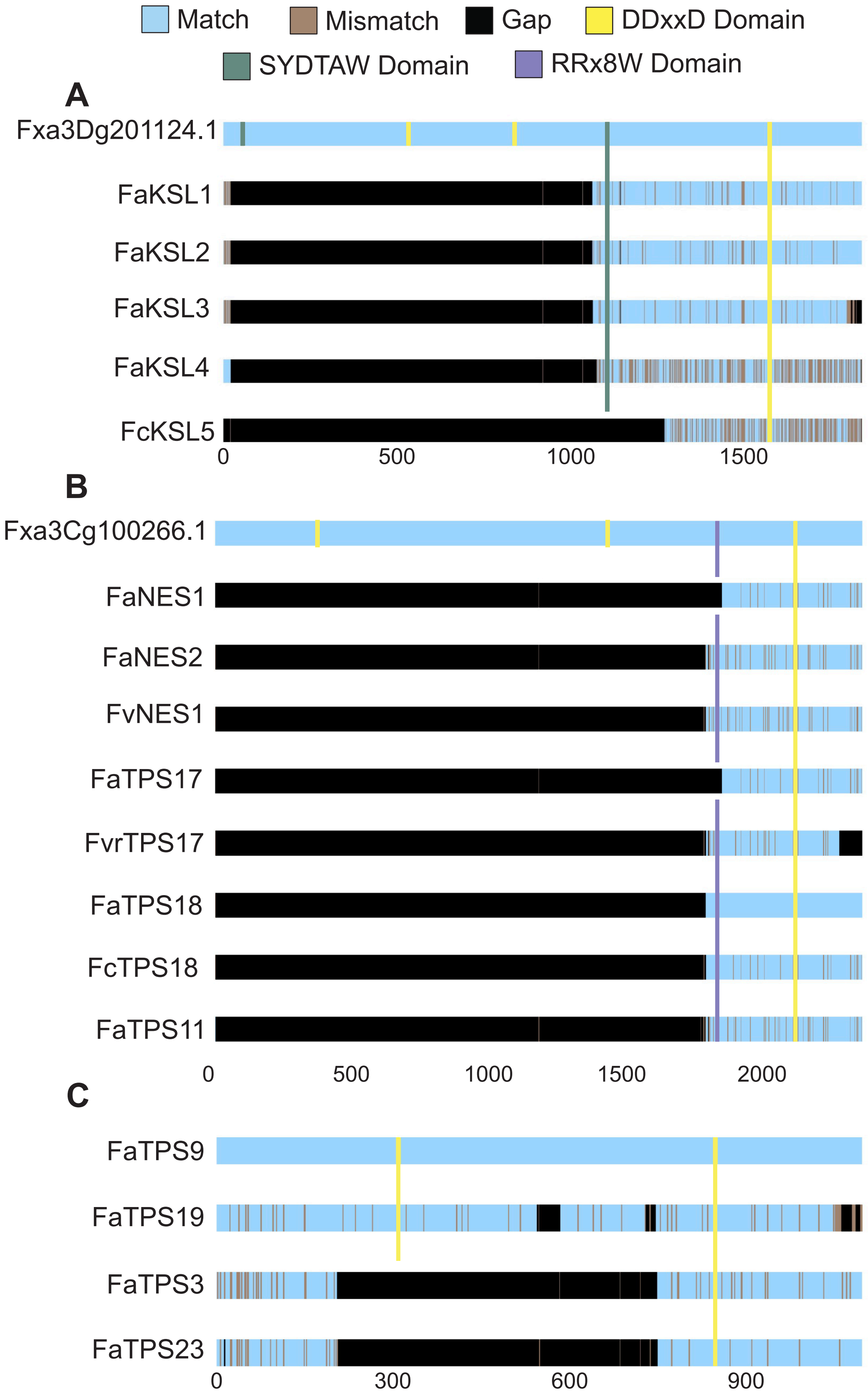

Match
Mismatch
Gap
DDxxD Domain
SYDTAW Domain
RRx8W Domain
A
Fxa3Dg201124.1
FaKSL1
FaKSL2
FaKSL3
FaKSL4
FcKSL5
0
500
1000
1500
B
Fxa3Cg100266.1
FaNES1
FaNES2
FvNES1
FaTPS17
FvrTPS17
FaTPS18
FcTPS18
FaTPS11
0
500
1000
1500
2000
C
FaTPS9
FaTPS19
FaTPS3
FaTPS23
0
300
600
900

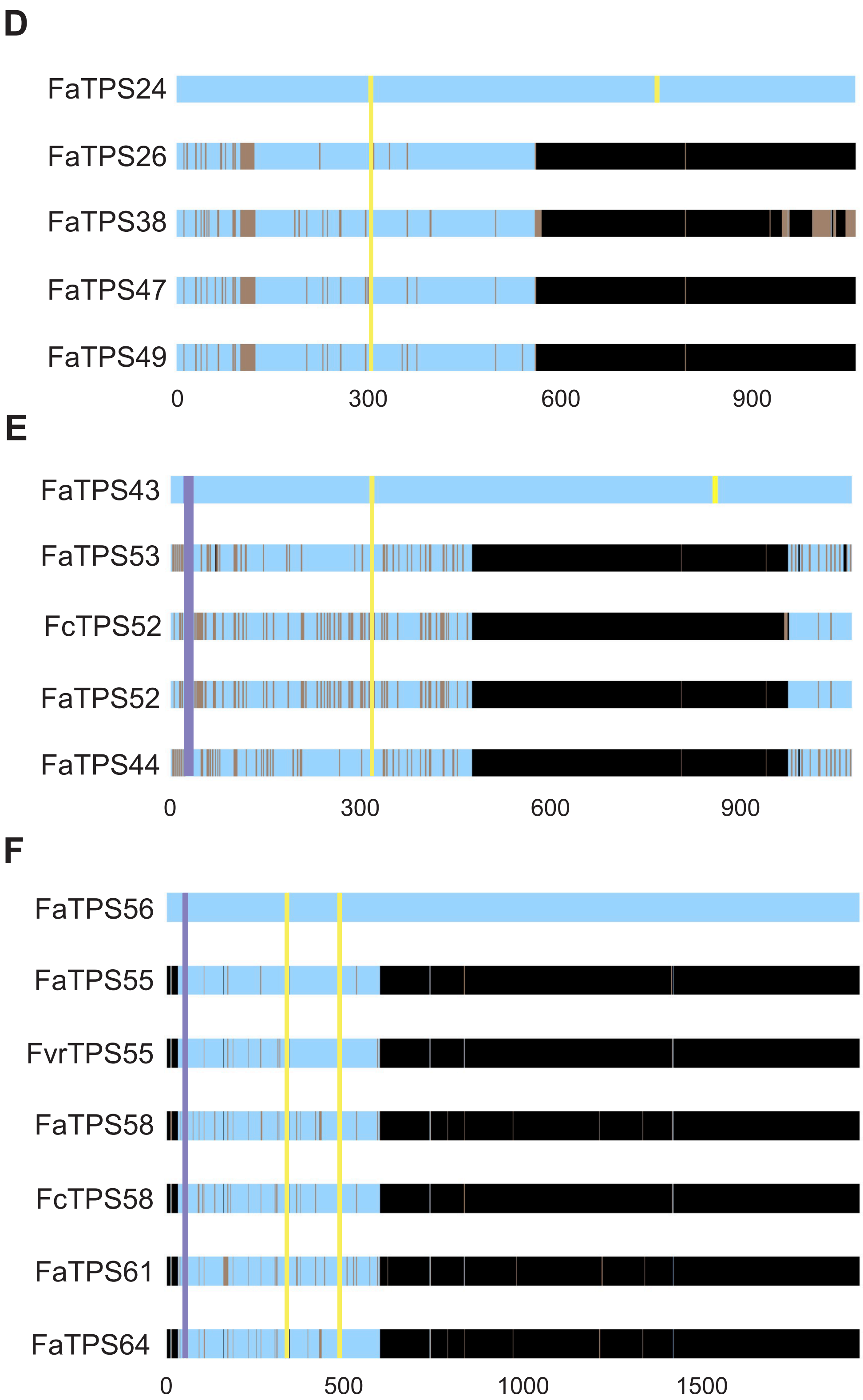
D
FaTPS24
FaTPS26
FaTPS38
FaTPS47
FaTPS49
0
300
600
900
E
FaTPS43
FaTPS53
FcTPS52
FaTPS52
FaTPS44
0
300
600
900
F
FaTPS56
FaTPS55
FvrTPS55
FaTPS58
FcTPS58
FaTPS61
FaTPS64
0
500
1000
1500

**Supplemental Fig. 5.** Amino acid sequence alignments of the seven putatively misassembled TPS sequences with the closest matches to either previously identified genes or those phylogenetically most closely related. Color coding: Black = missing regions in the alignment; brown = amino acid differences; DDxxD motif = yellow; DTAW motif = green; RRx8W motif = purple. Misassembled KSL4 (Fxa3Dg201124.1) (**A**) and NES (Fxa3Cg100266.1) (**B**) before sequence correction for gene synthesis are shown with other strawberry KSL and NES candidates, respectively. (**C, D, F**) Four TPS-a clade candidates *TPS9, TPS19, TPS24, TPS56*. (**E**) TPS-b clade *TPS43*.

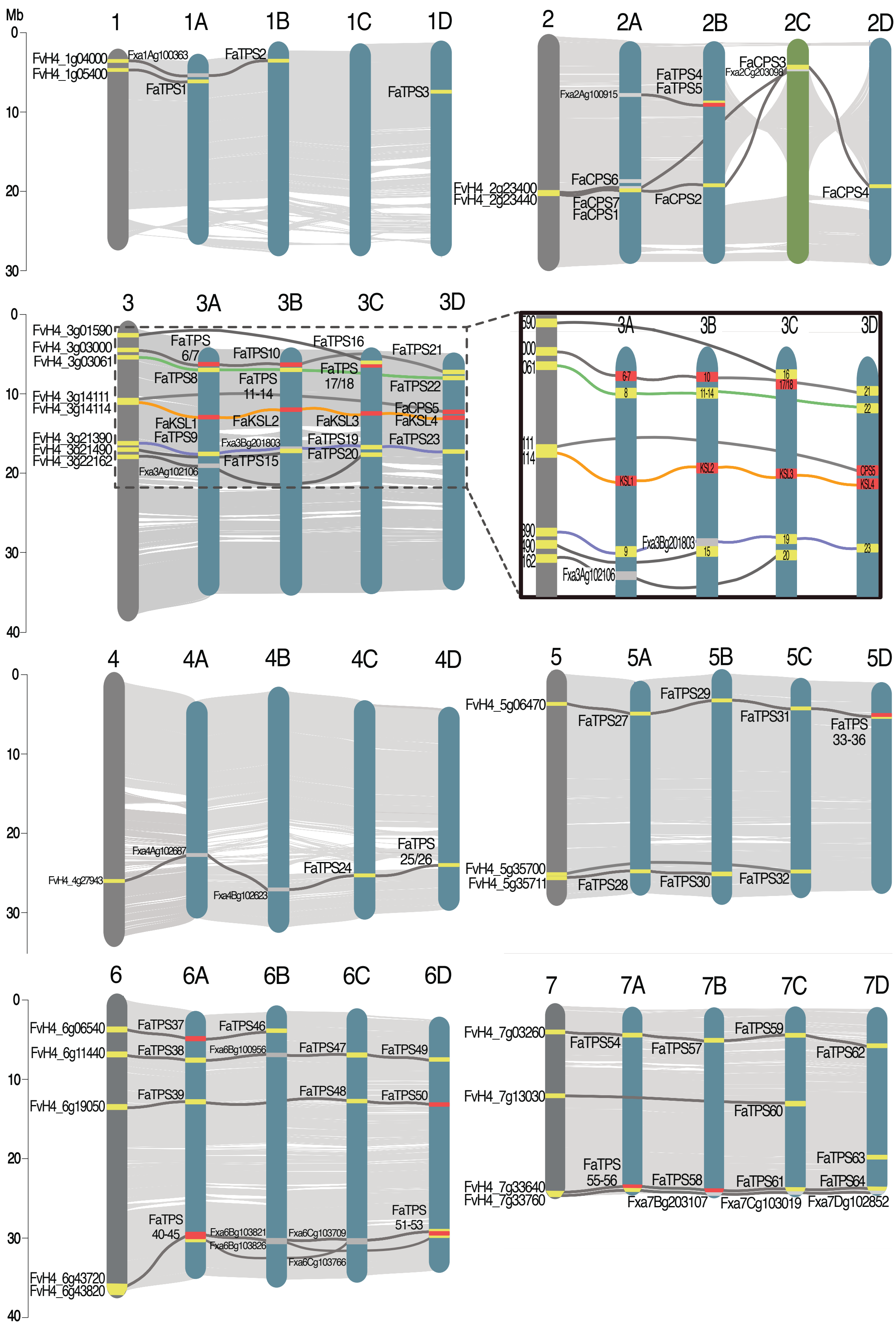
Mb
1
1A
1B
1C
1D
FvH4_1g04000
FvH4_1g05400
Fxa1Ag100363
FaTPS1
FaTPS2
FaTPS3
2
2A
2B
2C
2D
Fxa2Ag100915
FaTPS4
FaTPS5
FaCPS3
Fxa2Cg203098
FaCPS6
FvH4_2g23400
FvH4_2g23440
FaCPS7
FaCPS1
FaCPS2
FaCPS4
3
3A
3B
3C
3D
FvH4_3g01590
FvH4_3g03000
FvH4_3g03061
FvH4_3g14111
FvH4_3g14114
FvH4_3g21390
FvH4_3g21490
FvH4_3g22162
FaTPS 6/7
FaTPS8
FaTPS10
FaTPS 11-14
FaTPS16
FaTPS 17/18
FaTPS21
FaTPS22
FaCPS5
FaKSL1
FaKSL2
FaKSL3
FaKSL4
FaTPS9
Fxa3Bg201803
FaTPS15
FaTPS19
FaTPS20
FaTPS23
Fxa3Ag102106
4
4A
4B
4C
4D
Fxa4Ag102687
FvH4_4g27943
Fxa4Bg102623
FaTPS24
FaTPS 25/26
5
5A
5B
5C
5D
FvH4_5g06470
FaTPS27
FaTPS29
FaTPS31
FaTPS 33-36
FvH4_5g35700
FvH4_5g35711
FaTPS28
FaTPS30
FaTPS32
6
6A
6B
6C
6D
FvH4_6g06540
FvH4_6g11440
FvH4_6g19050
FvH4_6g43720
FvH4_6g43820
FaTPS37
FaTPS38
FaTPS39
FaTPS46
Fxa6Bg100956
FaTPS47
FaTPS48
FaTPS49
FaTPS50
FaTPS 40-45
Fxa6Bg103821
Fxa6Bg103826
Fxa6Cg103709
Fxa6Cg103766
FaTPS 51-53
7
7A
7B
7C
7D
FvH4_7g03260
FvH4_7g13030
FvH4_7g33640
FvH4_7g33760
FaTPS54
FaTPS57
FaTPS59
FaTPS62
FaTPS60
FaTPS 55-56
FaTPS58
FaTPS61
FaTPS63
FaTPS64
Fxa7Bg203107
Fxa7Cg103019
Fxa7Dg102852

**Supplemental Fig. 6.** Synteny plots of all 7 FaRR1 chromosomes with TPS genes identified in the diploid *F. vesca* genome. Gray lines show syntenic relationships between TPS genes. Syntenic pseudogenes are shown (gray). Chromosome 2C shown in green to depict whole chromosome inversion. Gene-dense region of chromosome 3 shown in detail; colored lines depict synteny matches across all 5 chromosomes.

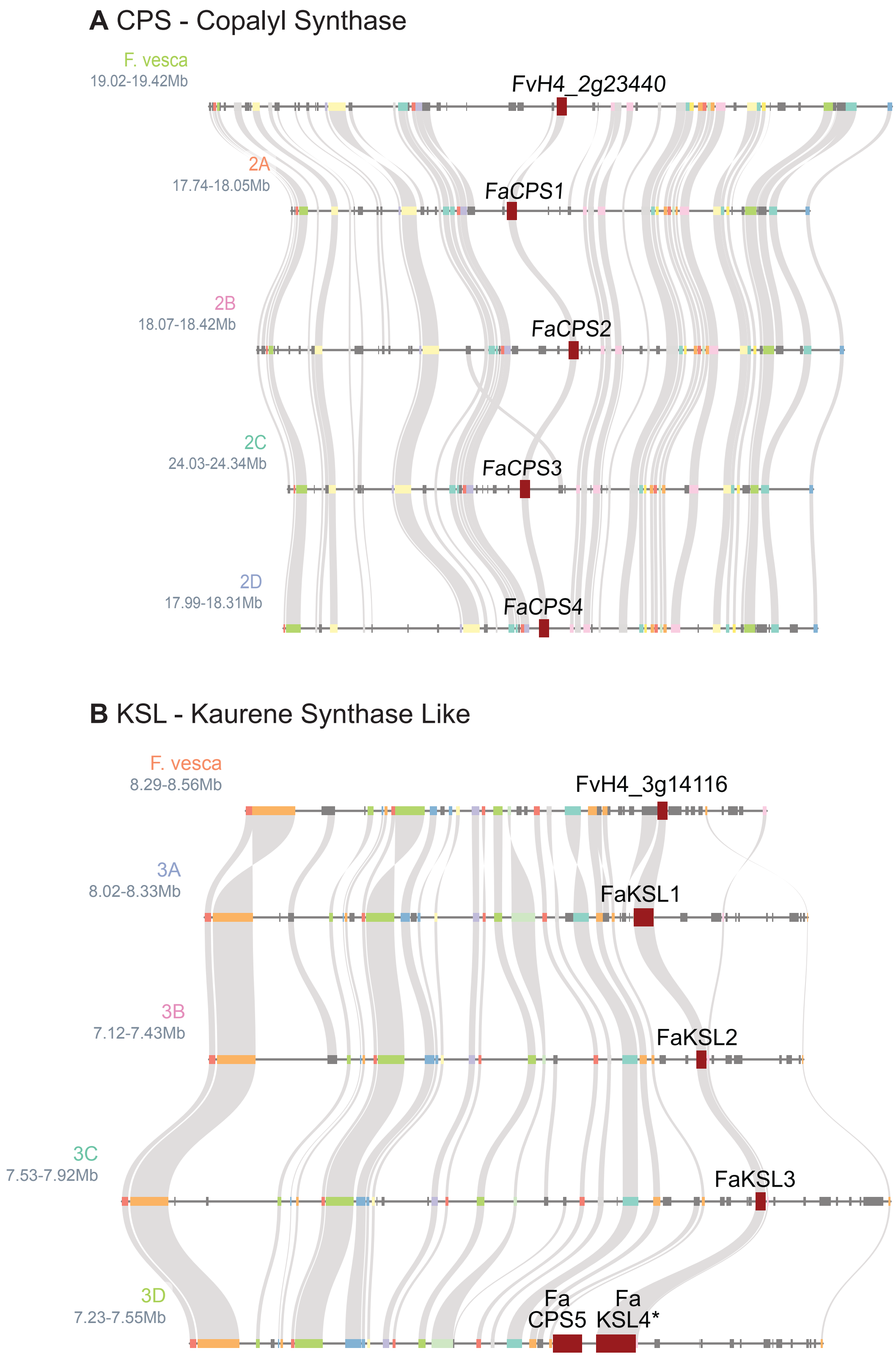

A CPS - Copalyl Synthase
F. vesca
19.02-19.42Mb
FvH4_2g23440
2A
17.74-18.05Mb
FaCPS1
2B
18.07-18.42Mb
FaCPS2
2C
24.03-24.34Mb
FaCPS3
2D
17.99-18.31Mb
FaCPS4
B KSL - Kaurene Synthase Like
F. vesca
8.29-8.56Mb
FvH4_3g14116
3A
8.02-8.33Mb
FaKSL1
3B
7.12-7.43Mb
FaKSL2
3C
7.53-7.92Mb
FaKSL3
3D
7.23-7.55Mb
Fa
CPS5
Fa
KSL4*

**Supplemental Fig. 7.** Microsynteny plots of diploid *F. vesca* and FaRR1 Royal Royce of diterpene synthases (**A**) Copalyl synthases (CPS) and (**B**) functionally tested Kaurene Synthases (KSL) tested in this study. Genes of interest are highlighted in red.

**A** NES - Nerolidol Synthase Like

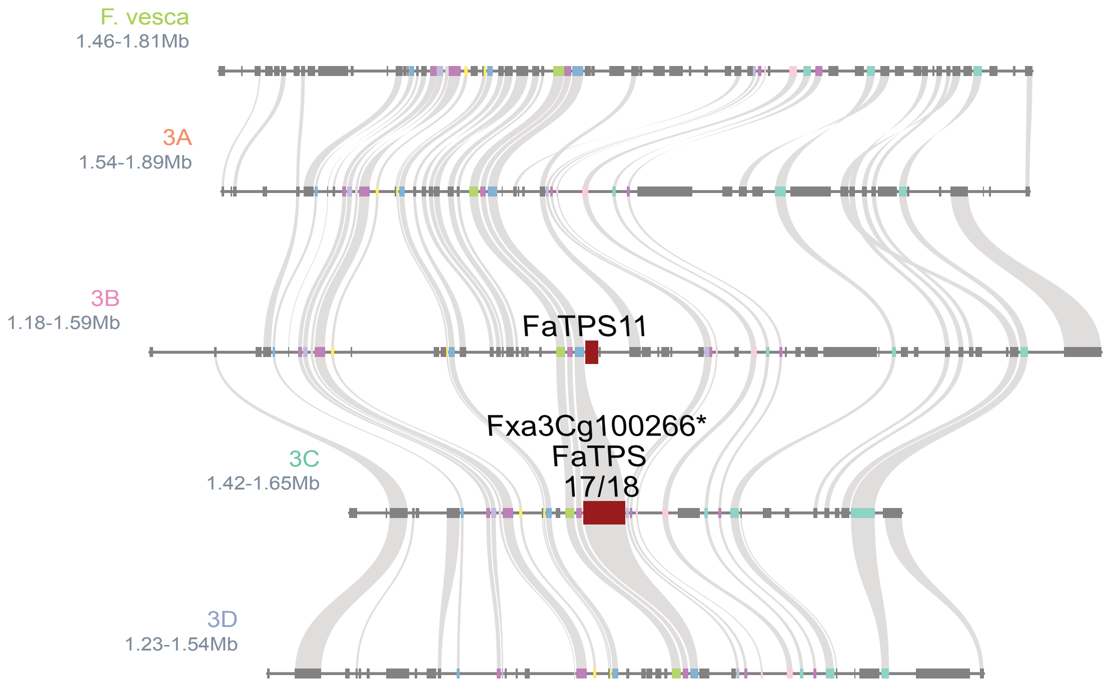

**B** FAR - Farnesene Synthase Like

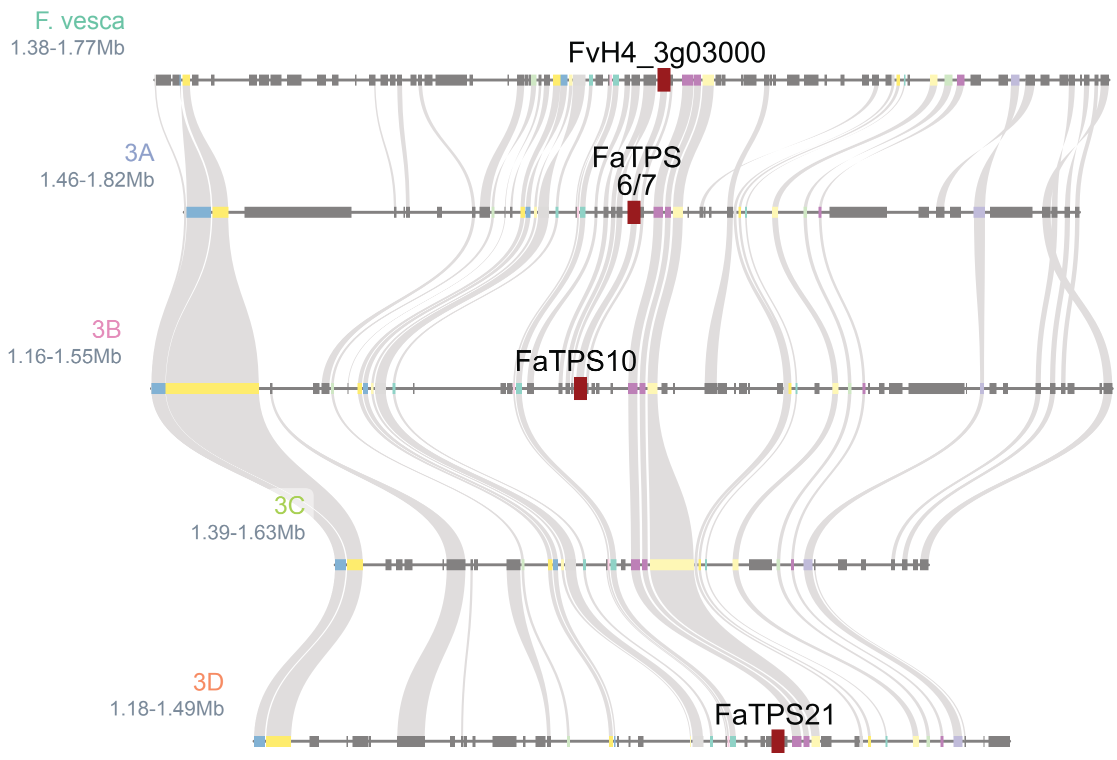

**C** PINS - Pinene Synthase Like

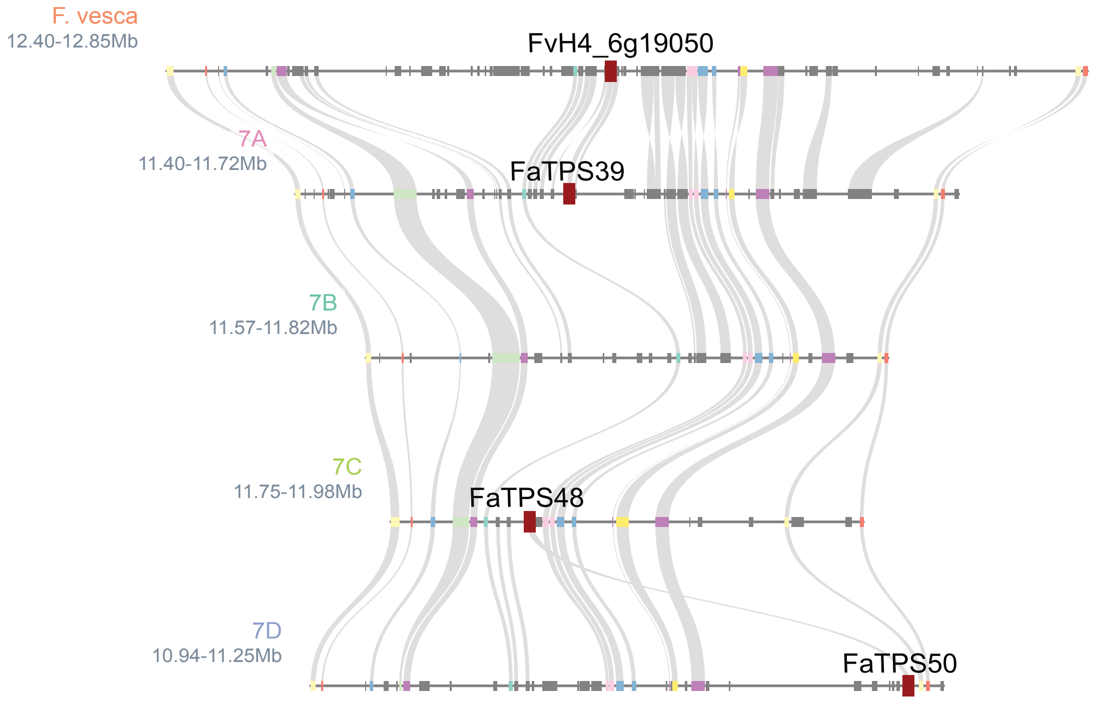

**D** TPSa Candidate

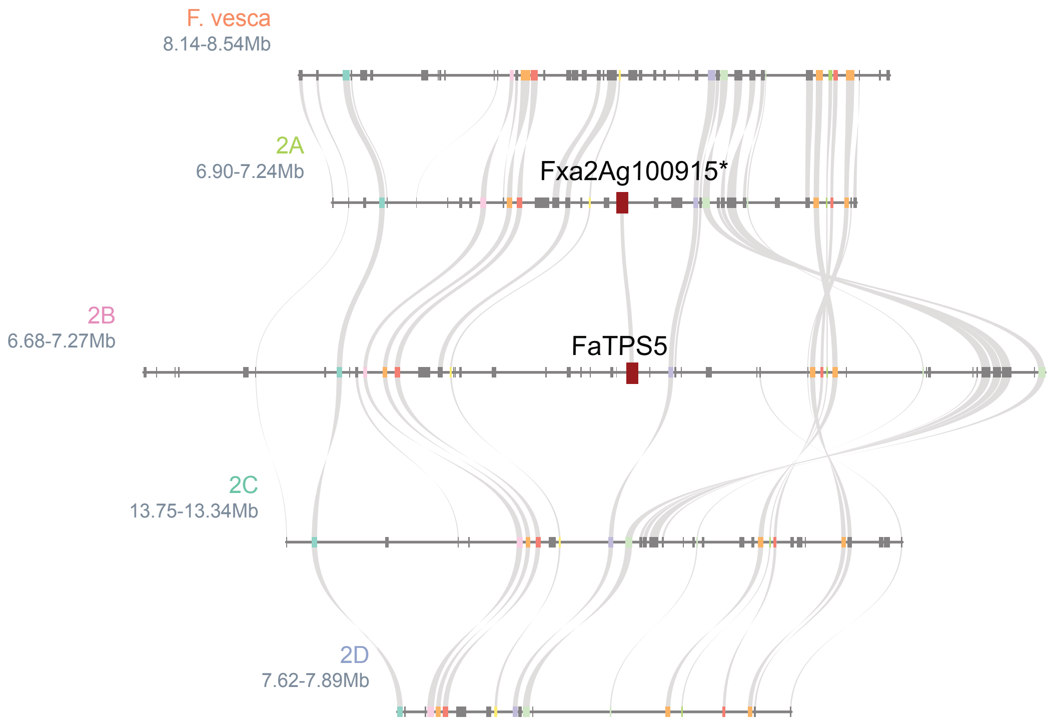

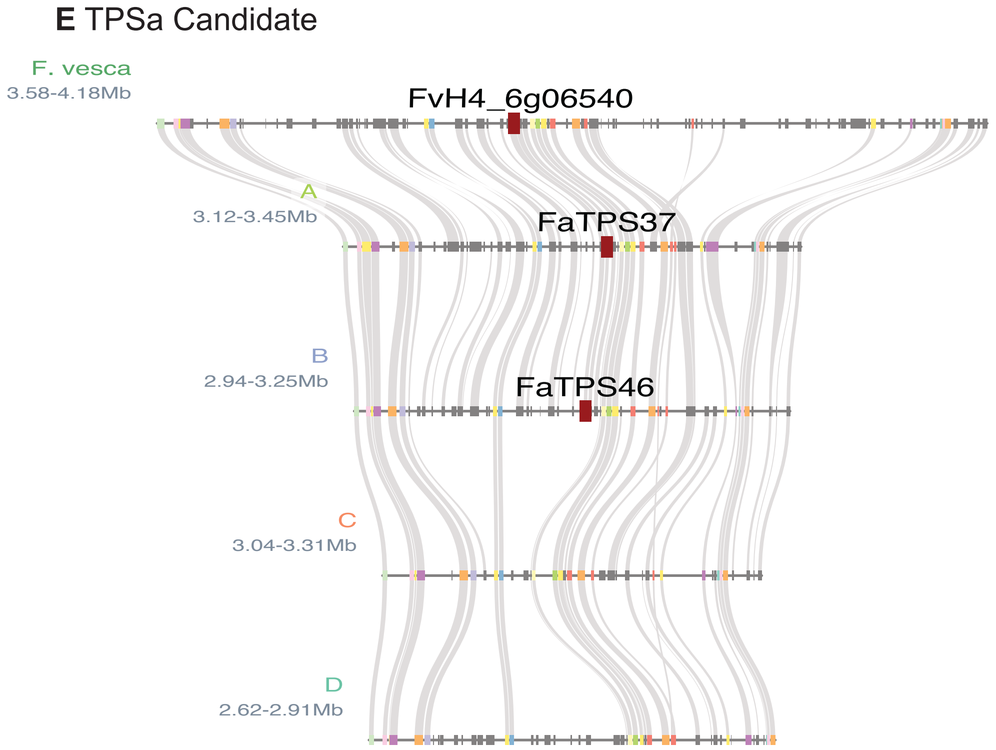
E TPSa Candidate
F. vesca
3.58-4.18Mb
FvH4_6g06540
A
3.12-3.45Mb
FaTPS37
B
2.94-3.25Mb
FaTPS46
C
3.04-3.31Mb
D
2.62-2.91Mb

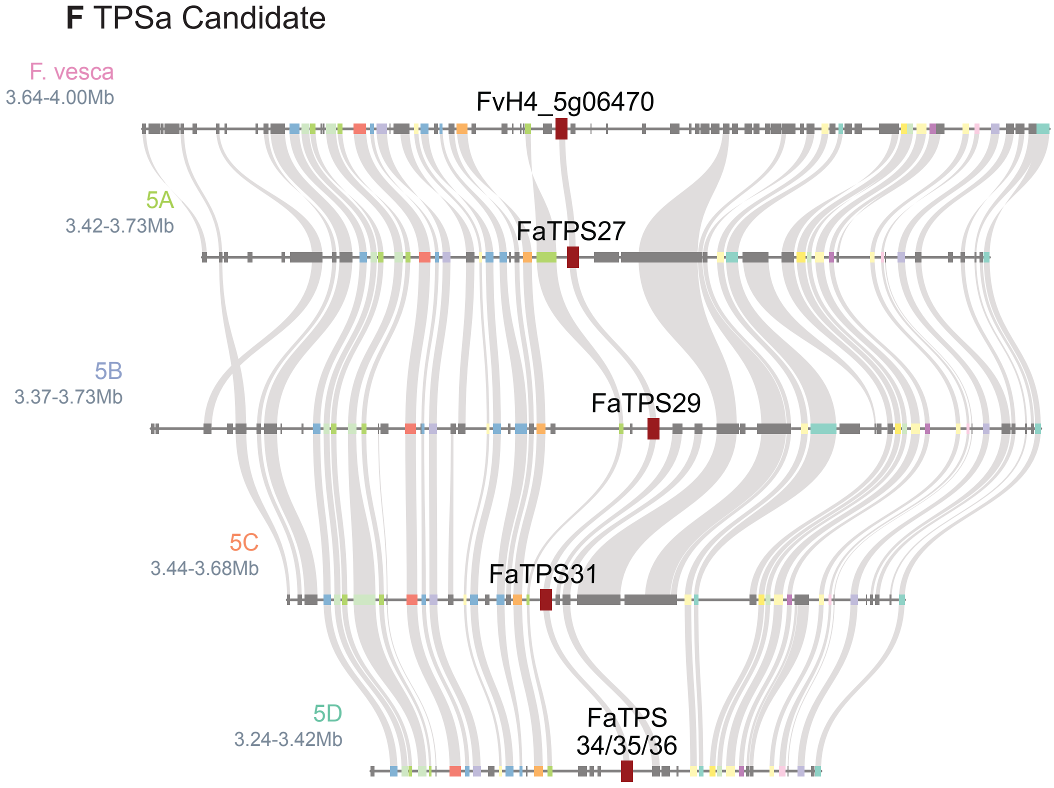
F TPSa Candidate
F. vesca
3.64-4.00Mb
FvH4_5g06470
5A
3.42-3.73Mb
FaTPS27
5B
3.37-3.73Mb
FaTPS29
5C
3.44-3.68Mb
FaTPS31
5D
3.24-3.42Mb
FaTPS
34/35/36

**G** TPSa Candidate

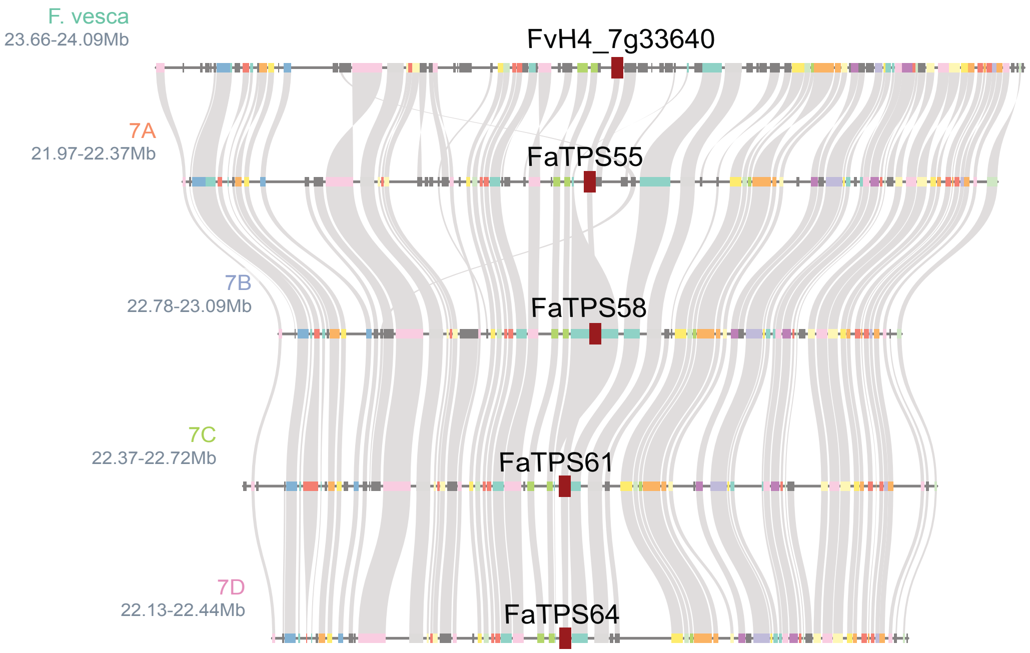

**H** TPSb Candidate

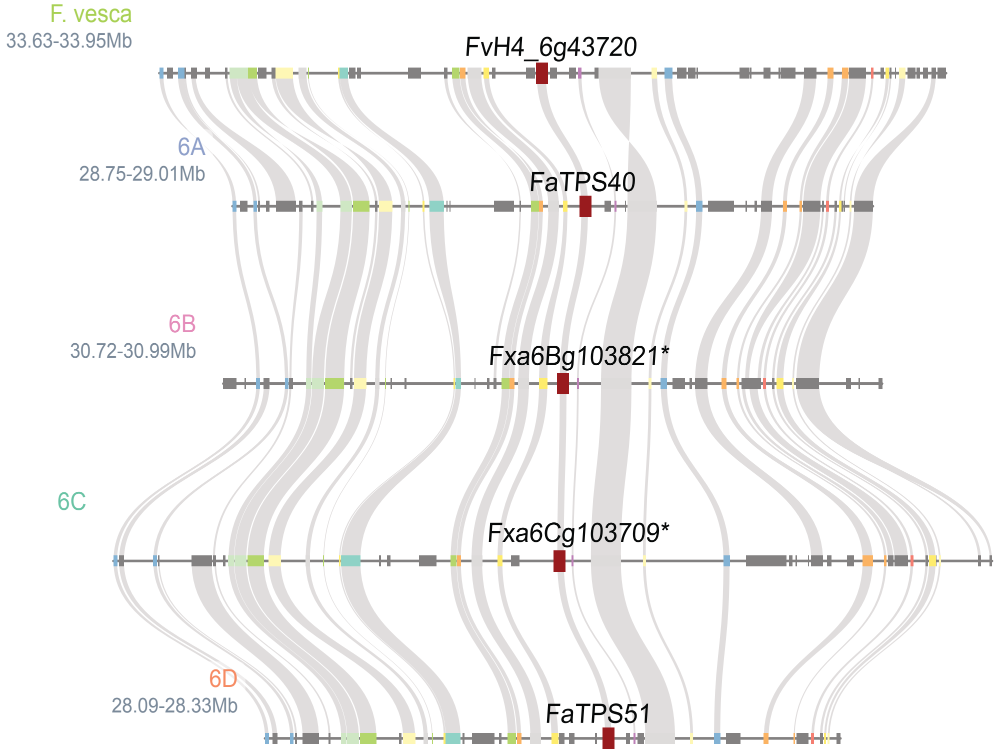

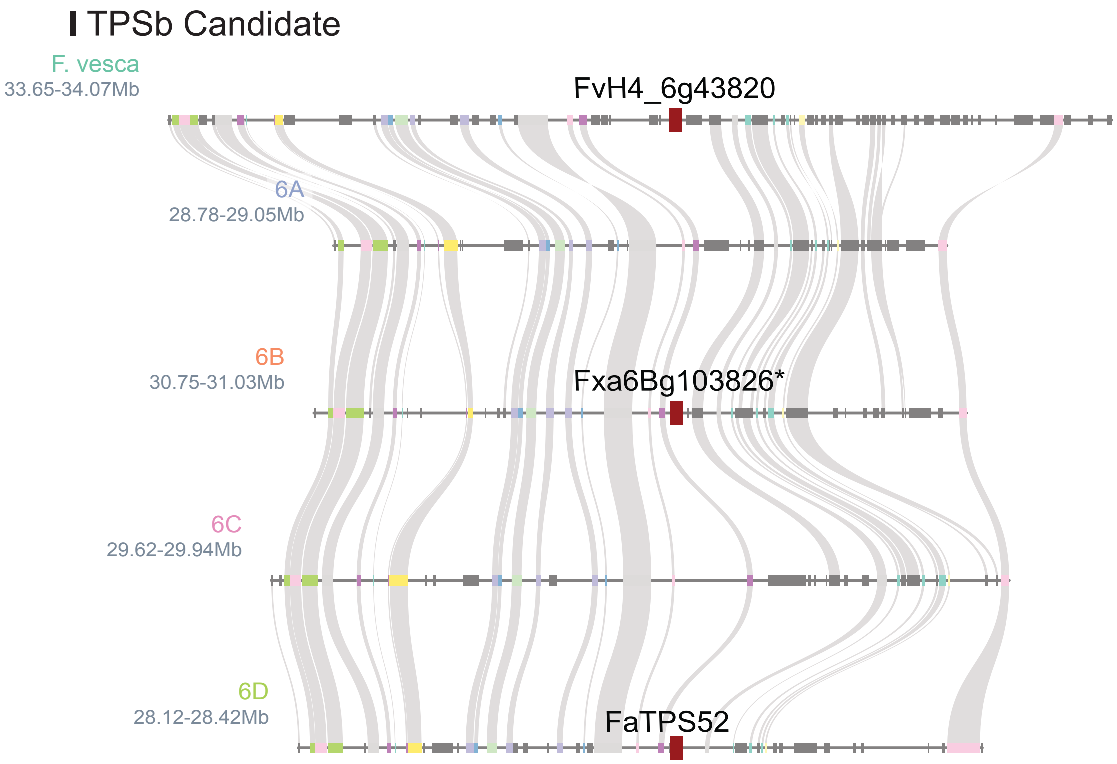

**Supplemental Fig. 8.** Microsynteny plots of diploid *F. vesca* and FaRR1 Royal Royce of all functionally analyzed mono-, sesqui-terpene synthases.

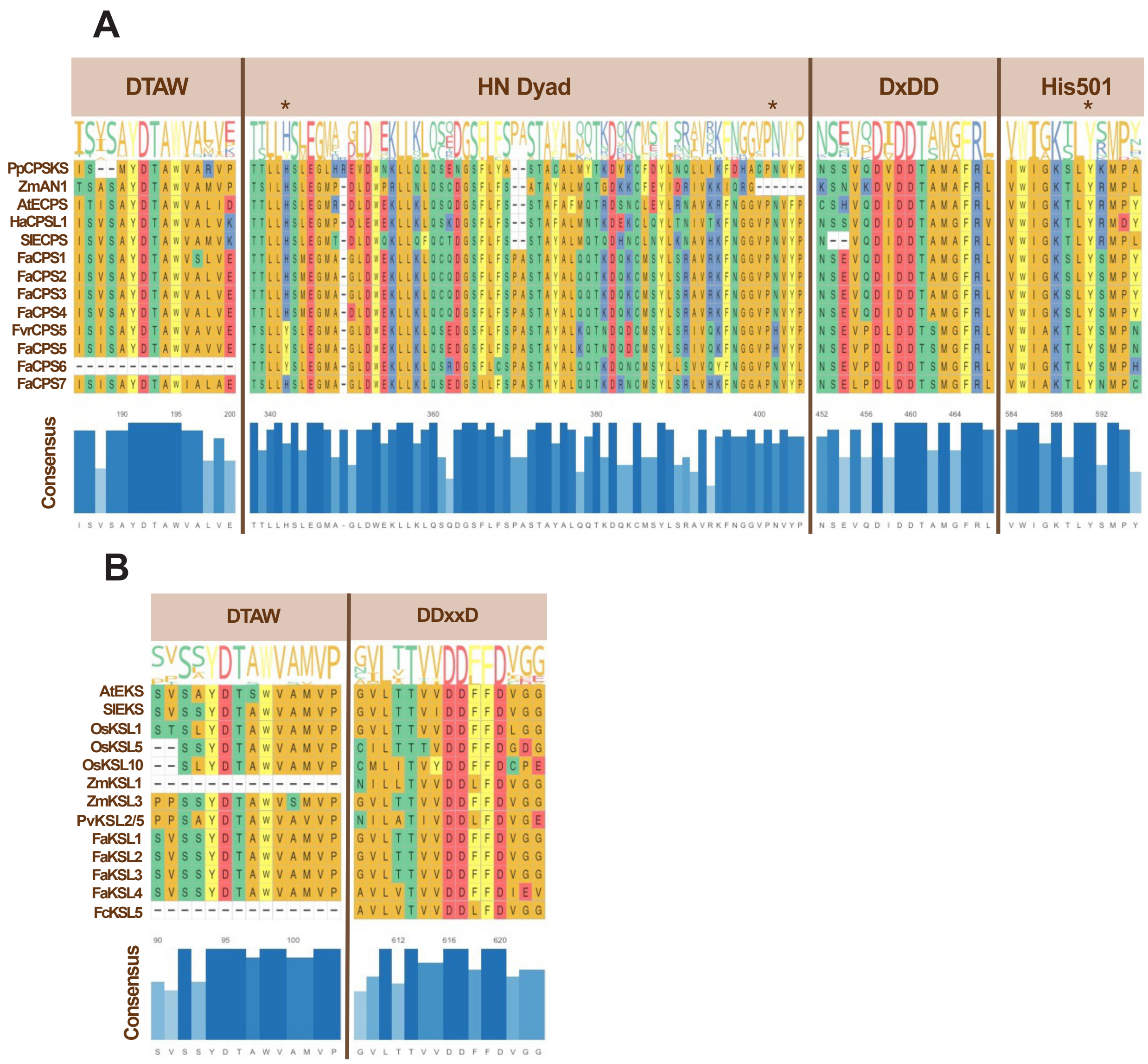

**Supplemental Fig. 9**. Protein sequence alignment of identified diterpene synthase candidates. (**A**) FaCPS candidates and reference sequences from *Physcomitrella patens*, *Zea mays*, *Arabidopsis thaliana*, *Helianthus annuus*, *Oryza sativa* and *Solanum lycopersicum* showing the DTAW domain, H-N dyad and DDxD motif. Additional OsCPS4 His501 residue representing *syn*-CPS activity is shown. (**B**) FaKSL candidates and reference sequences shown with DTAW domain and catalytic DxxDD motif.

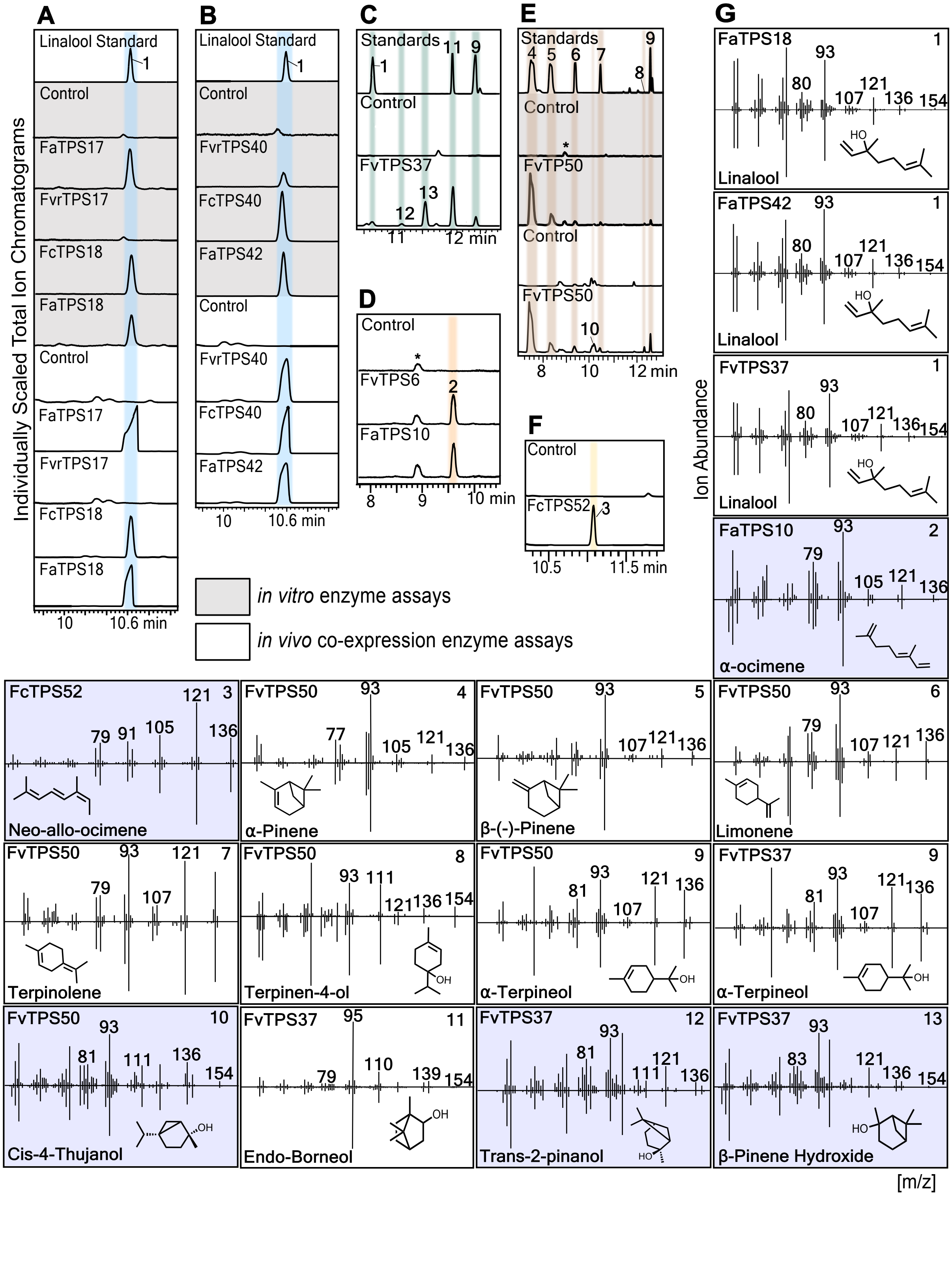
A
B
C
D
E
F
G
Linalool Standard
Control
FaTPS17
FvrTPS17
FcTPS18
FaTPS18
Individually Scaled Total Ion Chromatograms
10
10.6 min
FvrTPS40
FcTPS40
FaTPS42
Standards
FvTPS37
11
12 min
FvTPS6
FaTPS10
8
9
10 min
FvTP50
FvTPS50
12 min
FcTPS52
10.5
11.5 min
in vitro enzyme assays
in vivo co-expression enzyme assays
Ion Abundance
Linalool
α-ocimene
Neo-allo-ocimene
α-Pinene
β-(-)-Pinene
Limonene
Terpinolene
Terpinen-4-ol
α-Terpineol
Cis-4-Thujanol
Endo-Borneol
Trans-2-pinanol
β-Pinene Hydroxide
[m/z]

**Supplemental Fig. 10**. Functional characterization of terpene synthases (TPSs). (**A-F**) GC-MS traces of products resulting from either *in vitro* enzyme assays of individual recombinant TPSs with geranyl diphosphate (GPP) as a substrate (gray) or co-expression assays of individual TPSs and a GPP synthase in *E. coli* (white). (**G**) Mass spectra of enzyme products identified by comparison to authentic standards (white) or mass spectral databases (NIST, v17.1). FaTPS17, TPS18, and TPS42 taken from *F. × ananassa (Fa)* cultivar 'Royal Royce'; FvrTPS17 *F. virginiana* (*Fvr*) accession 'NC_96-35-2'; FcTPS18 *F. chiloensis* (*Fc*) ecotype 'Ambato'; FvrTPS40 accession 'Harris Springs'; FcTPS40 and FcTPS52 ecotype 'Isle de Lemuy'; FvTPS6 and FvTPS50 diploid accession 'UC06'; FaTPS10 cultivar 'EarliMiss'.

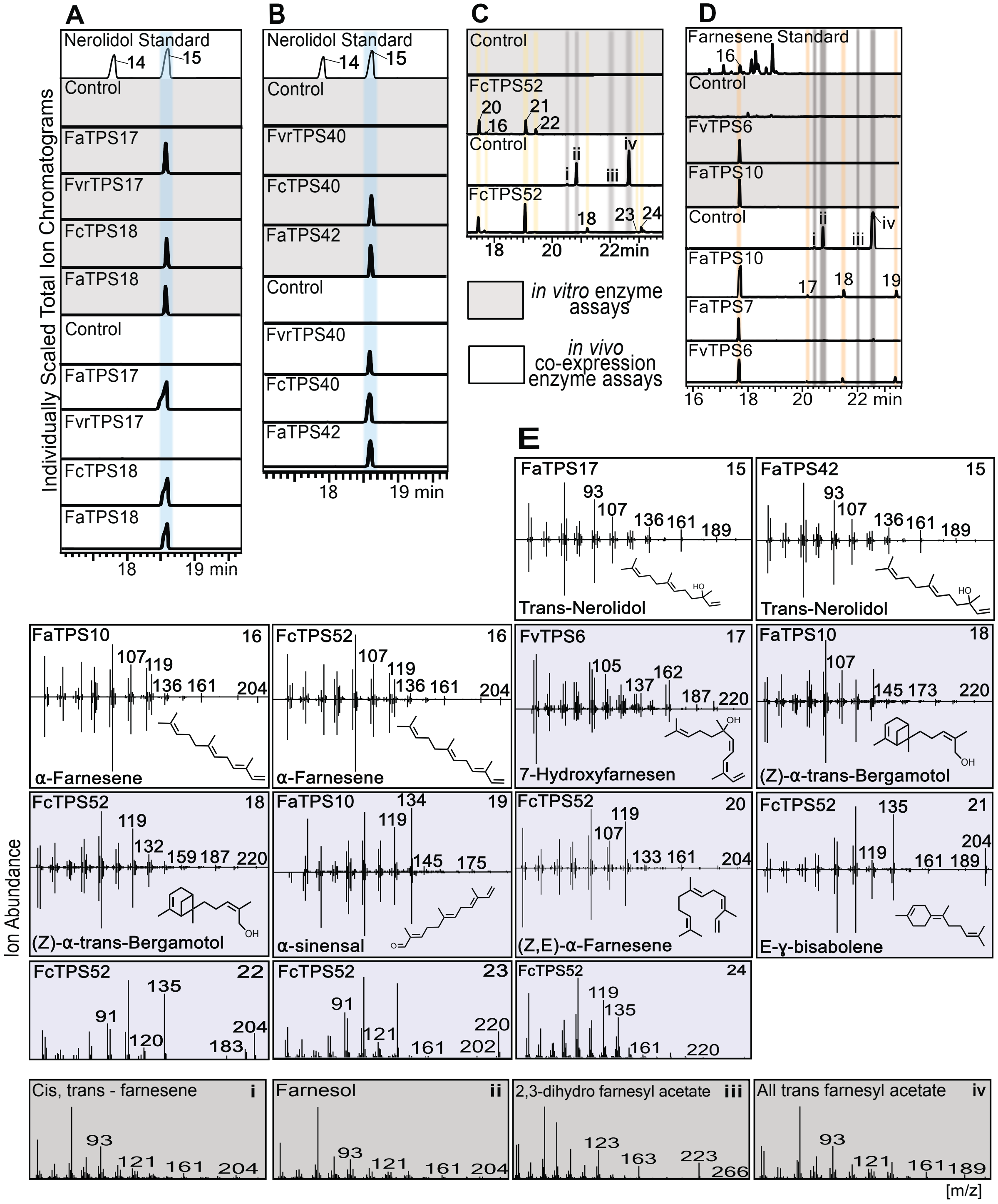
A
B
C
D
E
Nerolidol Standard
Farnesene Standard
Control
FaTPS17
FvrTPS17
FcTPS18
FaTPS18
FvrTPS40
FcTPS40
FaTPS42
FcTPS52
FvTPS6
FaTPS10
FaTPS7
Individually Scaled Total Ion Chromatograms
18
19 min
20
22min
16
22 min
in vitro enzyme assays
in vivo co-expression enzyme assays
Trans-Nerolidol
α-Farnesene
7-Hydroxyfarnesen
(Z)-α-trans-Bergamotol
α-sinensal
(Z,E)-α-Farnesene
E-ɣ-bisabolene
Ion Abundance
Cis, trans - farnesene
Farnesol
2,3-dihydro farnesyl acetate
All trans farnesyl acetate
[m/z]

**Supplemental Fig. 11**. Functional characterization of TPS-g and TPS-b clade terpene synthases (TPSs). (**A-F**) GC-MS traces of products resulting from either *in vitro* enzyme assays of individual recombinant TPSs with farnesyl diphosphate (FPP) as a substrate (gray) or co-expression assays of individual TPSs and a FPP synthase in *E. coli* (white). (**G**) Mass spectra of enzyme products identified by comparison to authentic standards (white) or mass spectral databases (NIST, v17.1) (purple). The following compounds are likely degradation products of FPP produced in *E. coli* cultures: (i) *Cis-trans*-farnesene; (ii) Farnesol; (iii) 2,3-dihydro farnesyl acetate; (iv) *Trans*-farnesyl acetate. FaTPS17, TPS18, and TPS42 taken from *F. × ananassa (Fa)* cultivar 'Royal Royce'; FvrTPS17 *F. virginiana* (*Fvr*) accession 'NC_96-35-2'; FcTPS18 *F. chiloensis* (*Fc*) ecotype 'Ambato'; FvrTPS40 accession 'Harris Springs'; FcTPS40 and FcTPS52 ecotype 'Isle de Lemuy'; FvTPS6 and FvTPS50 *F. vesca* (*Fv*) accession 'UC06'; FaTPS7 cultivar 'Primella'; FaTPS10 cultivar 'EarliMiss'.

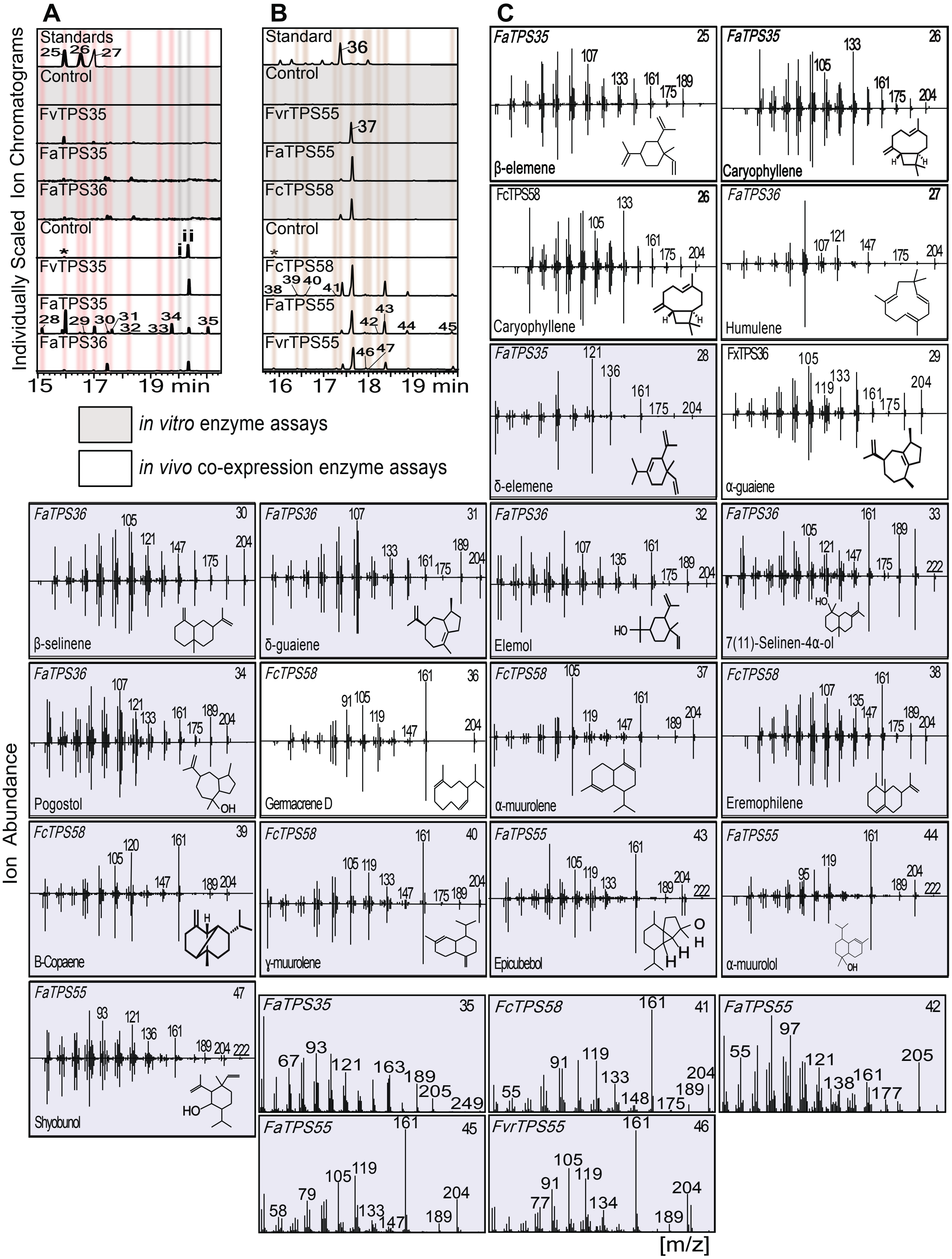
A
B
C
Individually Scaled Ion Chromatograms
Standards
Control
FvTPS35
FaTPS35
FaTPS36
Standard
FvrTPS55
FaTPS55
FcTPS58
15 17 19 min
16 17 18 19 min
in vitro enzyme assays
in vivo co-expression enzyme assays
Ion Abundance
β-elemene
Caryophyllene
Humulene
δ-elemene
α-guaiene
β-selinene
δ-guaiene
Elemol
7(11)-Selinen-4α-ol
Pogostol
Germacrene D
α-muurolene
Eremophilene
B-Copaene
γ-muurolene
Epicubebol
α-muurolol
Shyobunol
[m/z]

**Supplemental Fig. 12** Functional characterization of TPS-a terpene synthases (TPSs). (**A-B**) GC-MS traces of products resulting from either *in vitro* enzyme assays of individual recombinant TPSs with farnesyl diphosphate (FPP) as a substrate (gray) or co-expression assays of individual TPSs and a FPP synthase in *E. coli* (white). (**C**) Mass spectra of enzyme products identified by comparison to authentic standards (white) or mass spectral databases (NIST, v17.1) (purple). The following compounds are likely degradation products of FPP produced in *E. coli* cultures: (i) *Cis-trans*-farnesene; (ii) Farnesol. FvTPS35 from *F. vesca* (*Fv*) accession 'UC06'; FaTPS35 from *F.* × *ananassa (Fa)* cultivar 'Royal Royce'; FaTPS36 from *F.* × *ananassa (Fa)* 'Direktor Paul Wallbaum*';* FvrTPS55 *F. virginiana* (*Fvr* ) accession 'NC_96-35-2'; FaTPS55 cultivar 'Madame Moutot', FcTPS58 *F. chiloensis* (*Fc*) ecotype 'Ambato'.

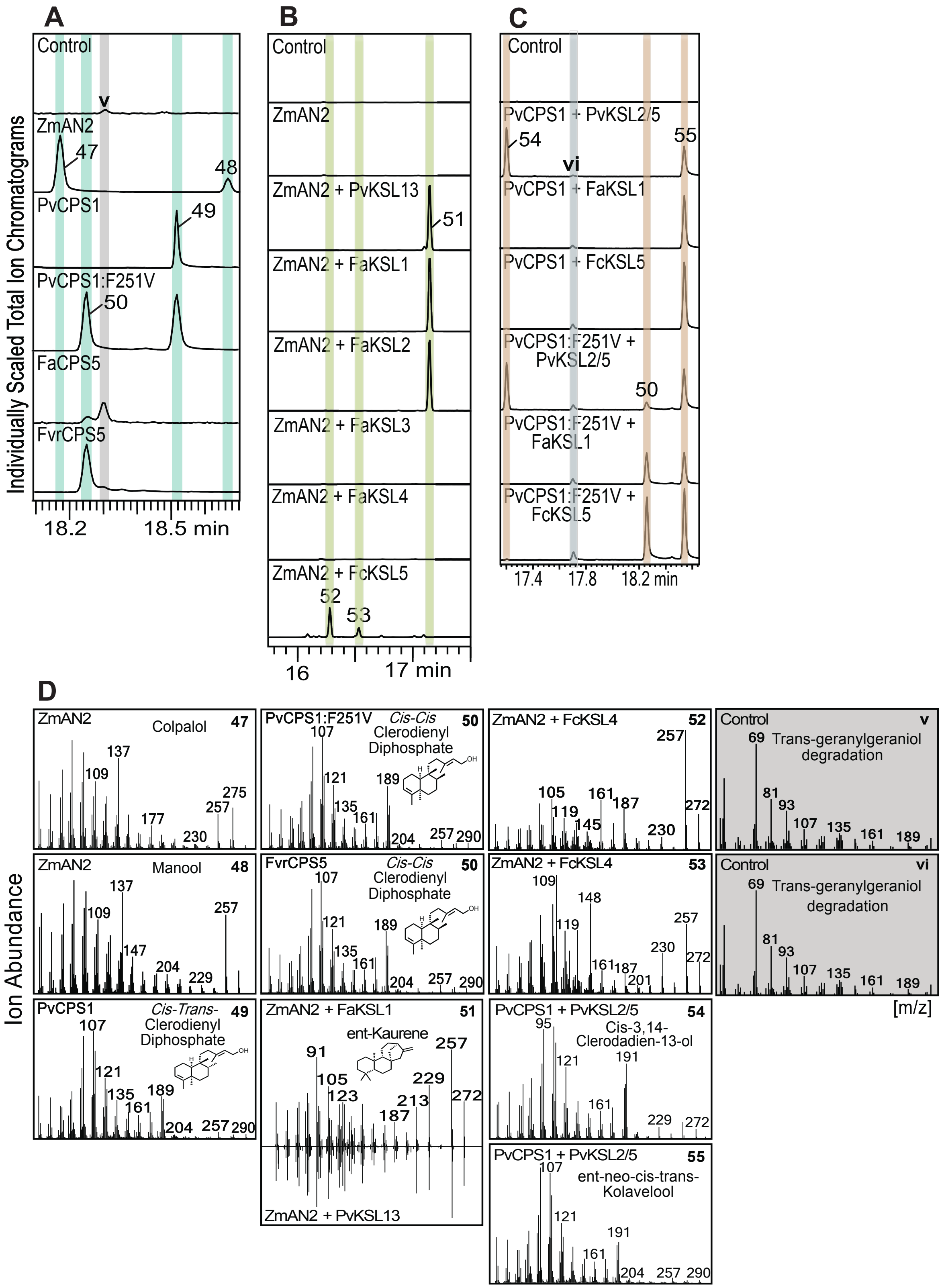
A
B
C
D
Individually Scaled Total Ion Chromatograms
Control
ZmAN2
PvCPS1
PvCPS1:F251V
FaCPS5
FvrCPS5
18.2
18.5 min
ZmAN2 + PvKSL13
ZmAN2 + FaKSL1
ZmAN2 + FaKSL2
ZmAN2 + FaKSL3
ZmAN2 + FaKSL4
ZmAN2 + FcKSL5
16
17 min
PvCPS1 + PvKSL2/5
PvCPS1 + FaKSL1
PvCPS1 + FcKSL5
PvCPS1:F251V + PvKSL2/5
PvCPS1:F251V + FaKSL1
PvCPS1:F251V + FcKSL5
17.4
17.8
18.2 min
Ion Abundance
Colpalol
Manool
Cis-Trans- Clerodienyl Diphosphate
Cis-Cis Clerodienyl Diphosphate
ent-Kaurene
ZmAN2 + FcKSL4
Cis-3,14-Clerodadien-13-ol
ent-neo-cis-trans-Kolavelool
Trans-geranylgeraniol degradation
[m/z]

**Supplemental Fig. 13** Functional characterization of diterpene synthases (diTPSs). (**A**) GC-MS traces of products resulting from co-expression assays of FaCPS5 or FvrCPS5 as compared to authentic standards produced by the *ent*-CPP synthase *Zea mays* AN2 (*ZmAN2*; Harris et al., 2005), the *ent-neo-cis-trans*-CLPP synthase *Panicum virgatum CPS1* (*PvCPS1*; Pelot et al., 2018) and the *PvCPS1* variant *PvCPS1:F251V* producing *ent-neo-cis-cis*-CLPP and *ent-neo-cis-trans*-CLPP (Pelot et al., 2016). (**B**) GC-MS traces of products resulting from co-expression assays of *FaKSL1-4* and *FcKSL5* with the *ent*-CPP synthase *Zea mays AN2* (*ZmAN2*; Harris et al., 2005). (**C**) GC-MS traces of products resulting from co-expression assays of specialized *P. virgatum* diTPSs tested in combinations with predicted FaKSLs to examine possible specialized FaCPS5 function. (**D**) Mass spectra of enzyme products identified by comparison to enzyme-produced standards. FaCPS5, and *FaKSL1-4* taken from *F.* × *ananassa (Fa)* cultivar 'Royal Royce'; FvrCPS5 *F. virginiana* (*Fvr* ) accession 'NC_96-35-2'; FcKSL5 *F. chiloensis* (*Fc*) ecotype 'Ambato'.

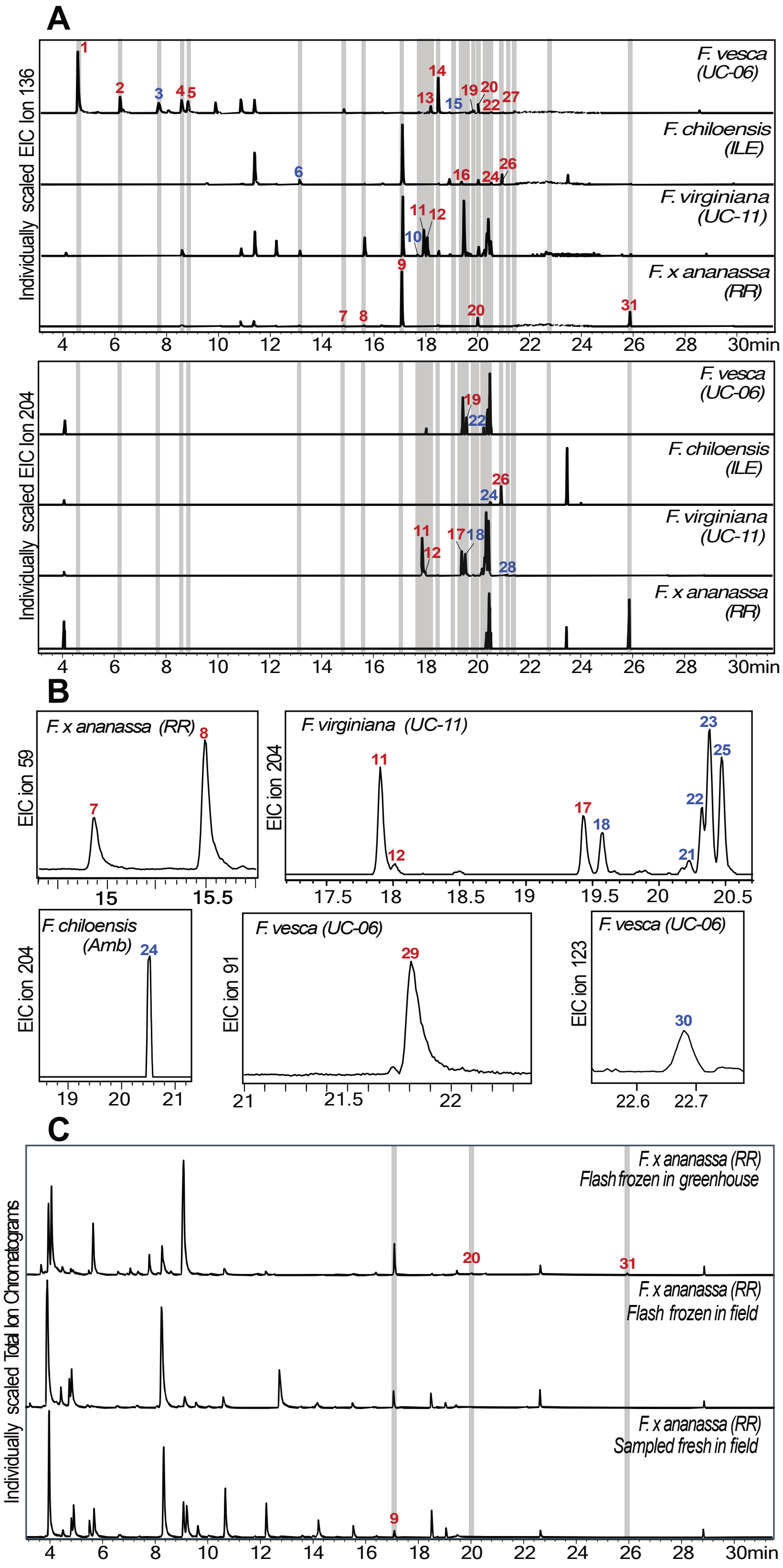

**1. α-Pinene (4.4)**
**2. β-Pinene (6.2)**
3. β-Myrcene (7.5)
**4. D-Limonene (8.4)**
**5. β-Phellandrene(8.7)**
6. Allo-Ocimene (13.1)
**7. Cis-Linalool Oxide (14.7)**
**8. Trans-Linalool Oxide (15.4)**
**9. Linalool (17.0)**
10. Isopulegol (17.7)
**11. α-Guaiene (17.9)**
**12. Caryophyllene (18.0)**
**13. Terpinen-4-ol (18.1)**
**14. 1-R-Myrtenal (18.6)**
15. Trans-Pinocarvyl Acetate (19.0)
**16. E-β-Farnesene (19.3)**
**17. Humulene (19.4)**
18. 4,11Selinadiene (19.6)
**19. Myrtenyl Acetate (19.7)**
**20. α-Terpineol (20.0)**
21. Eremophilene (20.2)
22. β-Selinene (20.3)
23. δ-Guaiene (20.4)
24. α-Muurolene (20.5)
25. Selinane (20.5)
**26. α-Farnesene (20.9)**
27. Trans-Myrtanyl Acetate (20.9)
28. 7-epi-α-Selinene (21.3)
**29. (-)-Myrtenol (21.7)**
30. Geraniol (22.6)
**31. Nerolidol (25.9)**

**Non-Terpene Peaks**
(4.0) Silica Fiber Contamination
(5.0) Silica Fiber Contamination
(8.1) Silica Fiber Contamination
(9.8) Ethyl Hexanoate
(10.8) Hexyl Acetate
(11.5) Tridecane
(12.3) Trans-2-Hexenyl Acetate
(16.3) Silica Fiber Contamination
(18.1) Methyl Caproate
(18.5) Benzyl Acetate
(18.9) Ethyl Caproate
(19.5) Trans-2-Hexenyl- Caproate
(20.5) Benzyl Acetate
(21.3) Methyl Salicylate
(22-23) Silica Fiber Contamination
(28.7) Internal Standard
(30) Silica Fiber Contamination

D

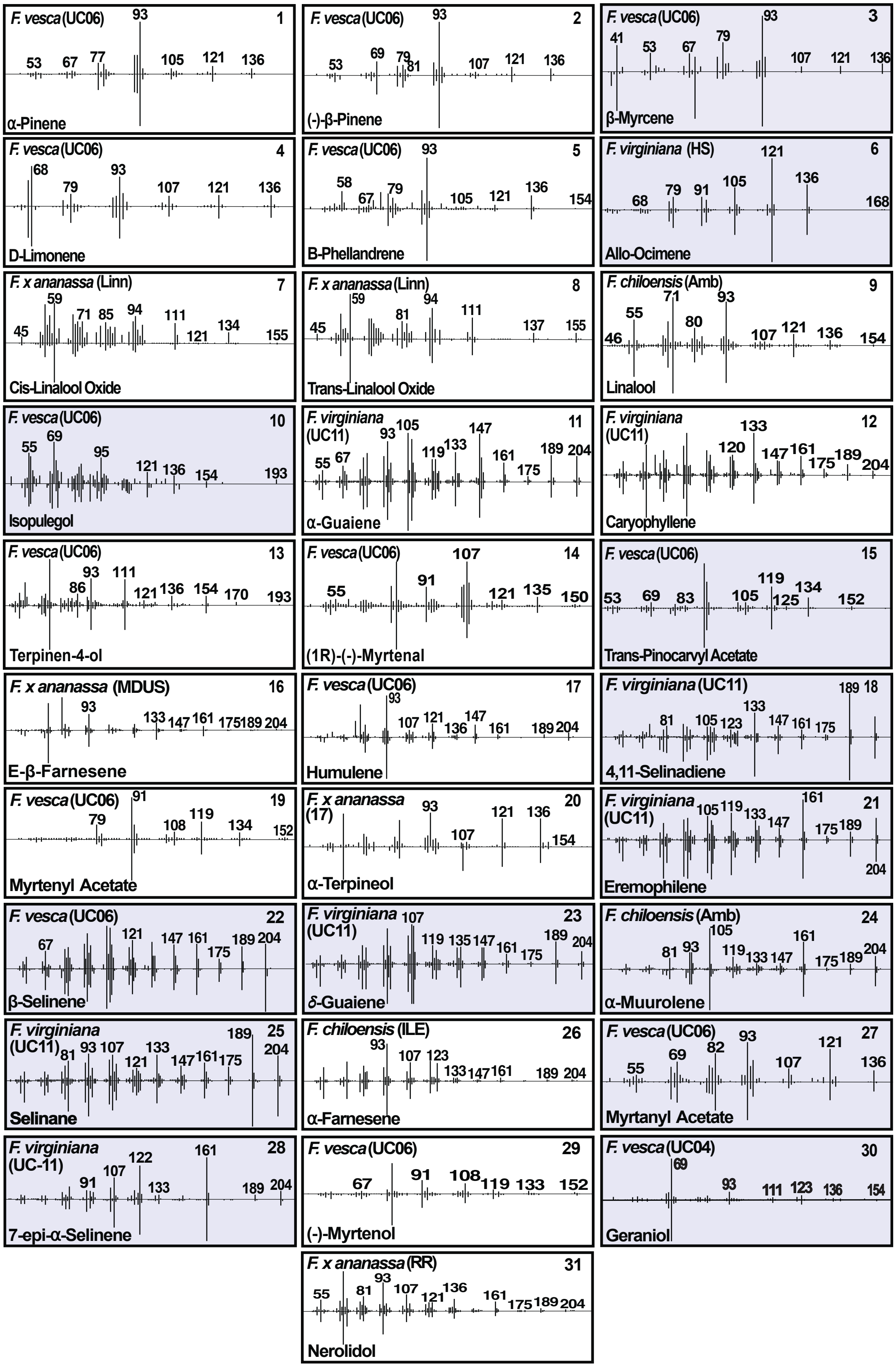

1 F. vesca (UC06) α-Pinene 53 67 77 93 105 121 136
2 F. vesca (UC06) (-)-β-Pinene 53 69 79 81 93 107 121 136
3 F. vesca (UC06) β-Myrcene 41 53 67 79 93 107 121 136
4 F. vesca (UC06) D-Limonene 68 79 93 107 121 136
5 F. vesca (UC06) B-Phellandrene 58 67 79 93 105 121 136 154
6 F. virginiana (HS) Allo-Ocimene 68 79 91 105 121 136 168
7 F. x ananassa (Linn) Cis-Linalool Oxide 45 59 71 85 94 111 121 134 155
8 F. x ananassa (Linn) Trans-Linalool Oxide 45 59 81 94 111 137 155
9 F. chiloensis (Amb) Linalool 46 55 71 80 93 107 121 136 154
10 F. vesca (UC06) Isopulegol 55 69 95 121 136 154 193
11 F. virginiana (UC11) α-Guaiene 55 67 93 105 119 133 147 161 175 189 204
12 F. virginiana (UC11) Caryophyllene 120 133 147 161 175 189 204
13 F. vesca (UC06) Terpinen-4-ol 86 93 111 121 136 154 170 193
14 F. vesca (UC06) (1R)-(-)-Myrtenal 55 91 107 121 135 150
15 F. vesca (UC06) Trans-Pinocarvyl Acetate 53 69 83 105 119 125 134 152
16 F. x ananassa (MDUS) E-β-Farnesene 93 133 147 161 175 189 204
17 F. vesca (UC06) Humulene 93 107 121 136 147 161 189 204
18 F. virginiana (UC11) 4,11-Selinadiene 81 105 123 133 147 161 175 189
19 F. vesca (UC06) Myrtenyl Acetate 79 91 108 119 134 152
20 F. x ananassa (17) α-Terpineol 93 107 121 136 154
21 F. virginiana (UC11) Eremophilene 105 119 133 147 161 175 189 204
22 F. vesca (UC06) β-Selinene 67 121 147 161 175 189 204
23 F. virginiana (UC11) δ-Guaiene 107 119 135 147 161 175 189 204
24 F. chiloensis (Amb) α-Muurolene 81 93 105 119 133 147 161 175 189 204
25 F. virginiana (UC11) Selinane 81 93 107 121 133 147 161 175 189 204
26 F. chiloensis (ILE) α-Farnesene 93 107 123 133 147 161 189 204
27 F. vesca (UC06) Myrtanyl Acetate 55 69 82 93 107 121 136
28 F. virginiana (UC-11) 7-epi-α-Selinene 91 107 122 133 161 189 204
29 F. vesca (UC06) (-)-Myrtenol 67 91 108 119 133 152
30 F. vesca (UC04) Geraniol 69 93 111 123 136 154
31 F. x ananassa (RR) Nerolidol 55 81 93 107 121 136 161 175 189 204

**Supplemental Fig. 14.** Total SPME-GC-MS chromatograms of a sample from each of the 4 *Fragaria* species represented in this study. 31 terpenes were identified and are shown and listed with retention times. Standard confirmed terpenes are labeled in red and bold, best spectral matches in blue. Non-terpene peaks shown are also listed. (**A**) Individually scaled extracted ion chromatograms for all terpenes found via SPME-GC-MS for monoterpene ion 136 and sesquiterpene ion 204. (**B**) Small or missing terpene peaks from TIC are shown. (**C**) Total ion chromatograms shown for ripe, *Fa* Royal Royce samples processed from freshly harvested field-grown fruits, flash-frozen field-grown fruits, and flash-frozen greenhouse-grown fruits. (**D**) NIST spectral matches with standard confirmed terpenes in white (NIST, v17.1).

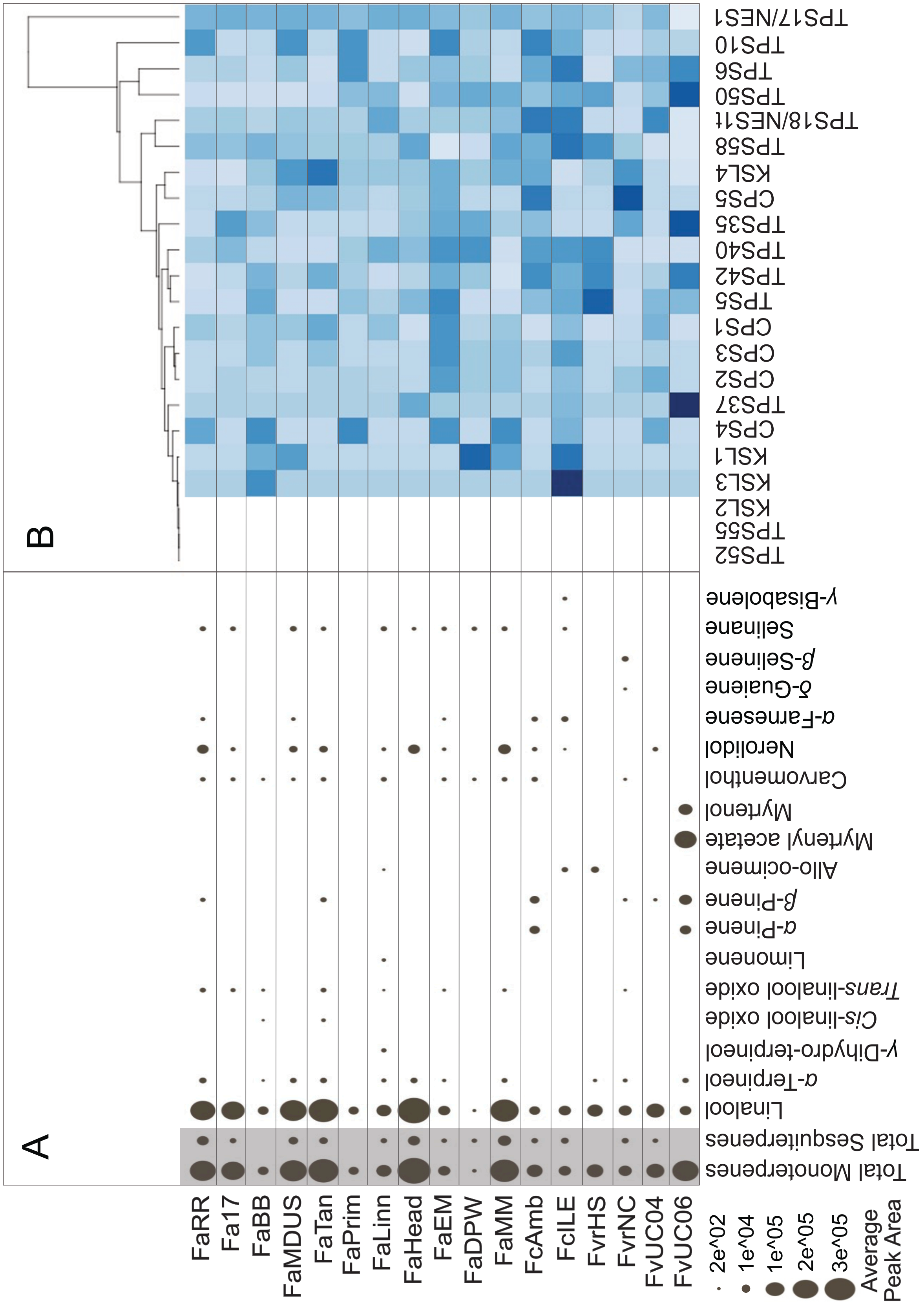
A
B
FaRR
Fa17
FaBB
FaMDUS
FaTan
FaPrim
FaLinn
FaHead
FaEM
FaDPW
FaMM
FcAmb
FclLE
FvrHS
FvrNC
FvUC04
FvUC06
2e^02
1e^04
1e^05
2e^05
3e^05
Average Peak Area
Total Monoterpenes
Total Sesquiterpenes
Linalool
α-Terpineol
γ-Dihydro-terpineol
Cis-linalool oxide
Trans-linalool oxide
Limonene
α-Pinene
β-Pinene
Allo-ocimene
Myrtenyl acetate
Myrtenol
Carvomenthol
Nerolidol
α-Farnesene
δ-Guaiene
β-Selinene
Selinane
γ-Bisabolene
TPS52
TPS55
KSL2
KSL3
KSL1
CPS4
TPS37
CPS2
CPS3
CPS1
TPS5
TPS42
TPS40
TPS35
CPS5
KSL4
TPS58
TPS18/NES1t
TPS50
TPS6
TPS10
TPS17/NES1

**Supplemental Fig. 15.** Abundance of terpene metabolites and relevant terpene synthase (TPS) genes in field-grown strawberry fruits which were flash frozen immediately after harvest for transcriptomic analysis. (**A**) Dot plot representing average normalized peak area of terpenes identified via SPME-GC-MS analysis of metabolites extracted from harvested from flash frozen tissue. (**B**) Hierarchical cluster analysis performed on gene expression data of select TPS genes analyzed in this study. TPSs were identified based on mapping functionally characterized TPSs against the FaRR1 genome (Hardigan et al. 2021). Gene expression data are based on three biological replicates. *F.* × *ananassa (Fa)* cultivars 'Royal Royce' (RR), '17C224P011' (17), 'Mara des Bois' (MDB) for which no field-grown transcriptome data was available, 'Beaver Belle' (BB), 'MDUS 5130' (MDUS), 'Tangi' (Tan), 'Primella' (Prim), 'Linn', 'Headliner' (Head), 'EarliMiss' (EM), 'Direktor Paul Wallbaum' (DPW), and 'Madame Moutot' (MM). *F. chiloensis* (*Fc*) ecotypes 'Ambato' (Amb) and 'Isle de Lemuy 02A White' (ILE). *F. virginiana* (*Fvr* ) accessions 'Harris Springs' (HS), 'NC_96-35-2' (NC), 'UC11'. Diploid *F. vesca* (*Fv*) accessions 'UC04' and 'UC06'

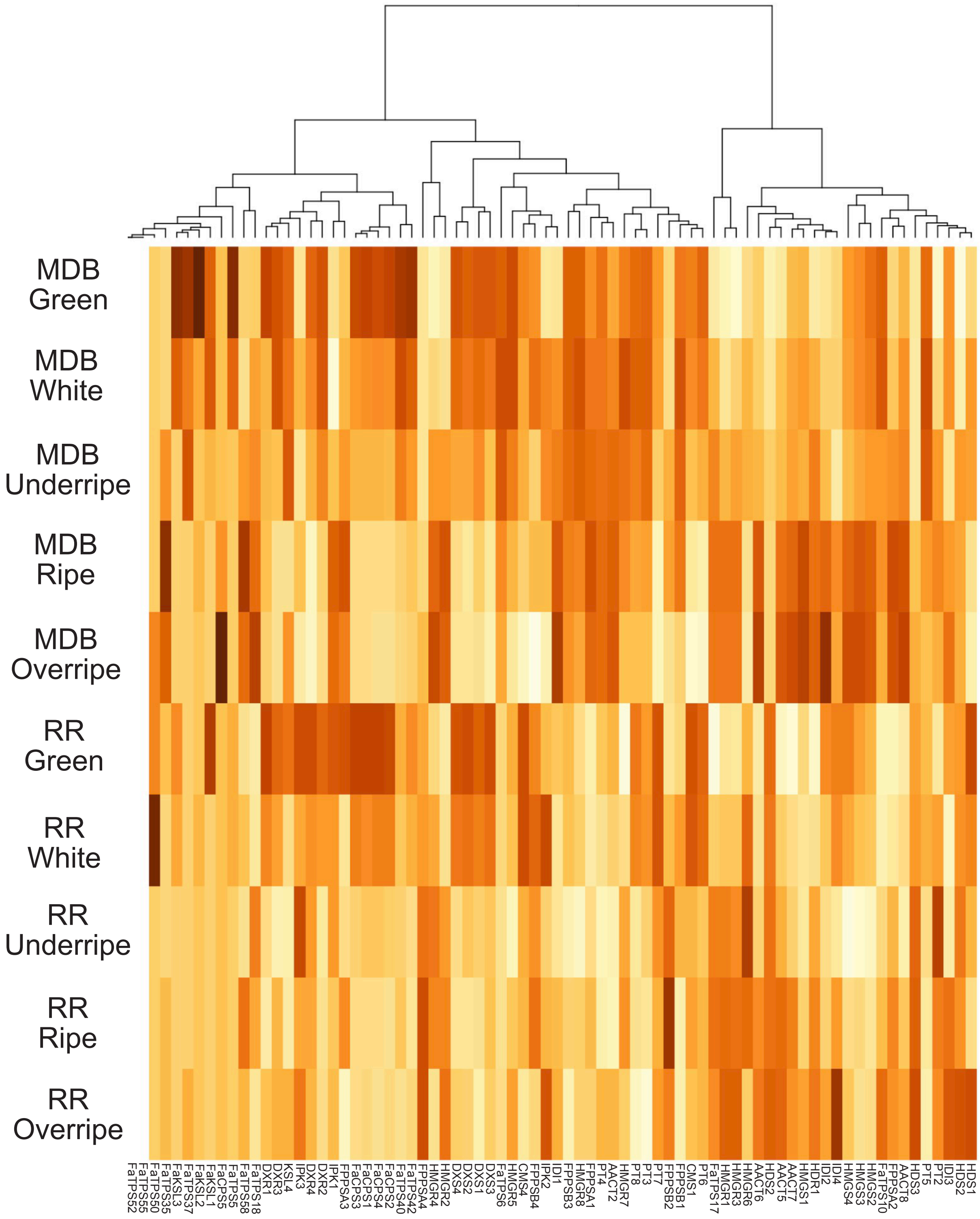
MDB Green
MDB White
MDB Underripe
MDB Ripe
MDB Overripe
RR Green
RR White
RR Underripe
RR Ripe
RR Overripe
FaTPS52
FaTPS55
FaTPS50
FaTPS35
FaKSL3
FaTPS37
FaKSL2
FaKSL1
FaCPS5
FaTPS5
FaTPS58
FaTPS18
DXR1
DXR3
KSL4
IPK3
DXR4
DXR2
IPK1
FPPSA3
FaCPS3
FaCPS1
FaCPS4
FaCPS2
FaTPS40
FaTPS42
FPPSA4
HMGR4
HMGR2
DXS4
DXS2
DXS1
DXS3
FaTPS6
HMGR5
CMS4
FPPSB4
IPK2
IDI1
FPPSB3
HMGR8
FPPSA1
PT4
AACT2
HMGR7
PT8
PT3
PT7
FPPSB2
FPPSB1
CMS1
PT6
FaTPS17
HMGR1
HMGR3
HMGR6
AACT6
HDS2
AACT5
AACT7
HMGS1
HDR1
IDI2
IDI4
HMGS4
HMGS3
HMGS2
FaTPS10
FPPSA2
AACT8
HDS3
PT5
PT2
IDI3
HDS2
HDS1

**Supplemental Fig. 16.** Hierarchical cluster analysis of select upstream terpene biosynthetic and terpene synthase (TPS) genes analyzed in this study from greenhouse-grown fruit from two cultivars, *Fa* Royal Royce and Mara des Bois, harvested from various developmental stages. TPSs were identified based on mapping functionally characterized TPSs in study against the FaRR1 genome (Hardigan et al. 2021). Gene expression data are based on three biological replicates.

**Supplemental Table 1.** Upstream Terpene biosynthetic genes of the mevalonate (MVA) and methylerythritol phosphate (MEP) pathways.

| FaRR1 Genome Hit | Name | Pathway | Length | Amino Acid Sequence | *F. vesca Hit* | Synteny Group |
|---|---|---|---|---|---|---|
| Fxa3Ag100033.2 | AACT1 | MVA | 314 | MIAAQTIQLGHNDIVVAGGMESMSNAPKYLPNARQGSRLGHDNIV<br>DGMLKDGLWDVYNDFGMGVCAELCAEKYTISREQQDNYAIQSFQ<br>RGISAQDAGLFSWEIVPVEVPGGRGKPSTIVDKDDGLQQFDAAKL<br>RKLRPSFKKSGGSVTAGNASIISDGAAALVLVSGEKALELGLQVIAKI<br>KGFADAAQAPELFTTAPALAIPRAISNAGLEASQIDYYEINEAFAVRI<br>QVVALANQKLLDLNPERVNAHGGAVSLGHPLGCSGARILVTLLGVL<br>RQKNGRYGVGGICNGGGGASAFVLELMPVARARPSKL* | FvH4_3g00290 | A |
| Fxa3Bg200036.1 | AACT2 | MVA | 411 | MDSCIKPRDVCVVGVARTPMGGFLGSLSSLSATRLGSIAIETALKR<br>AKVDPSLVQEVLFGNVLSANLGQAPARQAALGAGIPASVICTTINKV<br>CSSGMKAAMIAAQTIQLGHNDIVVAGGMESMSNAPKYLPNARQGS<br>RLGHDTIVDGMLKDGLWDVYNDFGMGVCAELCAEKYSISREQQD<br>NYAIQSFQRGISAQDAGLFSWEIVPVDVPGGRGKPSTIVDKDDGLQ<br>QFDAAKLRKLRPSFKKSGGSVTAGNASIISDGAAALVLVSGEKALEL<br>GLQVIAKIKGFADAAQAPELFTTAPALAIPRAISNAGLEASQIDYYEIN<br>EAFAVVALANQKLLGLNPERVNAHGGAVSLGHPLGCSGARILVTLL<br>GVLRQKNGRYGVGGICNGGGGASAFVLELMPAARARPSKL* | FvH4_3g00290 | A |
| Fxa3Cg100033.2 | AACT3 | MVA | 444 | MAASSDSCIKPRDVCIVGVARTPMRGFLGSLSSLSATRLGSIAIETA<br>LKRAKVGPSLVQEVLFGNVLSANLGQAPARQAALGAGIPASVICTTI<br>SKVCSSGMKGPSSFSLLLFSSSSSSSSSSFLACSAAMIAAQTIQLG<br>HNDIVVAGGMESMSNAPKYLPNARQGSRLGHDSIVDGMLKDGLW<br>DVYNDFGMGVCAELCAEKYSISREQQDNYAIQSFLRGISAQDAGLF<br>SWEIVPVEVPGGRGKPSTIVDKDDGLQQFDAAKLRKLRPSFKKSG<br>GSVTAGNSSIISDGAAALVLVSGEKALELGLQVIAKIKGFADAAQAP<br>ELFTTAPALAIPRAISNAGLEASQIDYYEINEAFAVRIQVVALANQKLL<br>GLNPERVNAHGGAVSLGHPLGCSGARILITLLGVLRQKNGRYGVG<br>GICNGGGGASAFVLELMPVARARPSKL* | FvH4_3g00290 | A |
| Fxa3Dg200030.2 | AACT4 | MVA | 442 | MAASSDSCIKPRDVCIVGVARTPMGGFLGSLSSLSATRLGSIAIETA<br>LKRAKVGPSLVQEVLFGNVLSANLGQAPARQAALGAGIPASVICTTI<br>NKVCSSGMKGPSSFSLLLFSSSSSSSSSFLACSAAMIAAQTIQLGHN<br>DIVVAGGMESMSNAPKYLPNARHGSRLGHDSIVDGMLKDGLWDV<br>YNDFGMGVCAELCAEKYSISREQQDNYAIQSFRRGISAQDAGLFS<br>WEIVPVEVPGGRGKPSTIVDKDDGLQQFDAAKLRKLRPSFKKSGG<br>SVTAGNASIISDGAAALVLVSGEKALELGLQVIAKIKGFADAAQAPEL<br>FTTAPALAIPRAISNAGLEASQIDYYEINEAFAVRIQVVALANQKLLG<br>LNPERVNAHGGAVSLGHPLGCSGARILVTLLGVLRQENGRYGVGG<br>ICNGGGGASAFVLELMPVARARPSKL* | FvH4_3g00290 | A |
| Fxa6Ag103901.1 | AACT5 | MVA | 409 | MAPVAAEACADSISPRGVYIVGVARTPMGAFLGALSSLPATKLGSIA<br>IEASLKRANVDPSLVEEVFFGNVLSANLGQAPARQAALGAGLSHSV<br>ICTTVNKVCASGLKATMLAAQSIQLGINDVVVAGGMESMSNVPKYL<br>AEARKGSRLGHDSLVDGMLKDGLWDVYNDYGMGVCAELCADQH<br>SVTREEQDNFACQSFERGIAAKDAGAFSWEIVPVEVSGGRGRPST | FvH4_6g40560 | B |

| | | | | | | |
|---|---|---|---|---|---|---|
| | | | | IVDKDEGLGKFDPAKLRKLRPSFKETGGSVTAGNASSISDGAAALV LVSGEKVLKLGLQVIAKISGFADAAQAPELFTTAPSLAIPKAISNAGL EASQIDYYEINEAFAVVALANQKLLGLDPAKVNVHGGAVALGHPLG CSGARILVTLLGVLKQKSAKYGVSGICNGGGGASALVLELL* | | |
| Fxa6Bg103557.1 | AACT6 | MVA | 409 | MAPVAAEACADSISPRGVYIVGVARTPMGAFLGALSSLPATKLGSIA IEAALKRANVDPSLVEEVFFGNVLSANLGQAPARQAALGAGLSHSV VCTTVNKVCASGLKATMLAAQSIQLGINDVVVAGGMESMSNVPKY LAEARKGSRLGHDSLVDGMLKDGLWDVYNDYGMGVCAELCADQ HSVTREEQDNFACQSFERGIAAKDAGAFSWEIVPVEVSGGRGRPS TIVDKDEGLGKFDAAKLRKLRPSFKETGGSVTAGNASSISDGAAAL VLVSGEKVLKLGLQVIAKISGFADAAQAPELFTTAPSLAIPKAISNAG LEASQIDYYEINEAFAVVALANQKLLGLDPAKVNVHGGAVALGHPL GCSGARILVTLLGVLKQKSAKYGVGGICNGGGGASALVLELL* | FvH4_6g40560 | B |
| Fxa6Cg103460.1 | AACT7 | MVA | 409 | MAPVAAEACADSISPRGVYIVGVARTPMGAFLGALSSLPATKLGSIA IEAALKRANVDPSLVEEVFFGNVLSANLGQAPARQAALGAGLSHSV ICTTVNKVCASGLKATMLAAQSIQLGINDVVVAGGMESMSNVPKYL AEARKGSRLGHDSLVDGMLKDGLWDVYNDYGMGVCAELCADQH SVTREEQDNFACQSFERGIAAKDAGAFSWEIVPVEVSGGRGRPST IVDKDEGLGKFDAAKLRKLRPSFKETGGSVTAGNASSISDGAAALV LVSGEKVLKLGLQVIAKISGFADAAQAPELFTTAPSLAIPKAISNAGL EASQIDYYEINEAFAVVALANQKLLGLDPAKVNVHGGAVALGHPLG CSGARILVTLLGVLKQKSAKYGVGGICNGGGGASALVLELL* | FvH4_6g40560 | B |
| Fxa6Dg103364.1 | AACT8 | MVA | 409 | MAPVAAEACADSISPRGVYIVGVARTPMGAFLGALSSLPATKLGSIA IEAALKRANVDPSLVEEVFFGNVLTANLGQAPARQAALGAGLSHSV ICTTVNKVCASGLKATMLAAQSIQLGINDVVVAGGMESMSNVPKYL AEARKGSRLGHDSLVDGMLKDGLWDVYNDYGMGVCAELCADQH SVTREEQDNFACQSFERGIAAKDAGAFSWEIVPVEVSGGRGRPST IVDKDEGLGKFDAAKLRKLRPSFKETGGSVTAGNASSISDGAAALV LVSGEKVLKLGLQVIAKISGFADAAQAPELFTTAPSLAIPKAISNAGL EASQIDYYEINEAFAVVALANQKLLGLDPAKVNVHGGAVALGHPLG CSGARILVTLLGVLKQKSAKYGVGGICNGGGGASALVLELL* | FvH4_6g40560 | B |
| Fxa5Ag202521.1 | CMS1 | MEP | 275 | MLSGHRFSFSGKPRRARIATSGFTCFTAKLAQEIKRETVVVKEKSV SVILLAGGKGTRMGASMPKQYLPLLSQPIALYSFYTFSRMPQVKEII VVCDPSYRDVFEDAKYKIQTALKFTLPGKERQDSVYSGLQAIDLTS ELVCIHDSARPLVSSENVEKVLKDGWLNGAAVLGVPVKATIKEANN ESFVVRTLDRRTLWEMQTPQVIKPELLKKGFELVNREGLEVTDDVS IVEHLPHPVYITEGSYTNIKVTTPDDLLLAERILSMDTKKSSE* | FvH4_5g27480 | C |
| Fxa5Bg102434.2 | CMS2 | MEP | 319 | MALLVHRLHFTLSPSSSSLLFSNDNNLRPISGNSLPPVPFLNSIHTL MFSGHRFTFSEKPRRARIATSGFTAKLAQEIAPETVVVKEKSVSVIL LAGGKGTRMGASMPKQYLPLLSQPIALYSFYTFSRMPQVKEIIVVC DPSYRDVFEDAKHKIQTELKFTLPGKERQDSVYSGLQAIDLTSELV CIHDSARPLVSSEDVEKVLEDGWLNGAAVLGVPVKATIKEANNESF VVRTLDRRTLWEMQTPQVIKPELLKKGFELVNREGLEVTDDVSIVE HLPHPVYITEGSYTNIKVTTPDDLLLAERILSMETKKSSE* | FvH4_5g27480 | C |
| Fxa5Cg202277.2 | CMS3 | MEP | 275 | MFSGHRFTFSEKHRRARIATSGFTCFTAKLAQEIKLETVVVKEKSVS VILLAGGKGTRMGASMPKQYLPLLSQPIALYSFYTFSCMPQVKEIIV VCDPSYRDVFEDAKNKIQTELKFTLPGKERQDSVYSGLQAIDLTSE LVCIHDSARPLVSSEDVEKVLKDGWLNGAAVLGVPVKATIKEANNE | FvH4_5g27480 | C |

| | | | | | | |
|---|---|---|---|---|---|---|
| | | | | SFVVRTLDRRTLWEMQTPQVIKPELLKKGFELVNREGLEVTDDVSIVEHLPHPVYITEGSYTNIKVTTPDDLLLAERILSMETKKSSE* | | |
| Fxa5Dg202343.1 | CMS4 | MEP | 217 | MGASMPKQYLPLLSQPIALYSFYTFSRMPQVKEIIVVCDPSHRDVFEDAKYKIQTALKFTLPGKERQDSVYSGLQAIDLTSELVCIHDSARPLVSSENVEKVLKDGWLNAAAVLGVPVKATIKEANNESFVVRTLDRRTLWEMQTPQVIKPELLKKGFELVNREGLEVTDDVSIVEHLPHPVYITEGSYTNINVTTPDDLLLAERILSMDTKKSSE* | FvH4_5g27480 | C |
| Fxa5Ag200475.1 | DXR1 | MEP | 472 | MALNLLSPAEIKAISFLDSTKSTHLPKLPGGFALRRKDCRTVIGRRIQCSAQAPPPAWPGSAFPEPGRRTWDGPKPISIVGSTGSIGTQTLDIVAENPDKFRVVALAAGSNVTLLVDQVKRFKPKLVAVRNESLVDELKEALSGLEDKPEIIPGEQGVIEVARHPDAVTVVTGIVGCAGLRPTVAAIEAGKDIALANKETLIAGGPFVLPLAHKHNVKILPADSEHSAIFQCIQGLPEGALRRIILTASGGAFRDWPVEKLKEVKVADALKHPNWTMGKKITVDSATLFNKGLEVIEAHYLYGADYDDIEIVIHPQSIIHSMIETQDSSVLAQLGWPDMRLPILYTMSWPERIYCSEVTWPRLDLCKLGSLTFKAPDNVKYPSMDLAYSAGRAGGTMTGVLSAANEKAVEMFIDEKISYLDIFKVVELTCAKHRAELVTSPSLEEIIHYDLWARDYAANLQNSTSSTPVFA* | FvH4_5g05300 | D |
| Fxa5Bg100449.1 | DXR2 | MEP | 472 | MALNLLSPAEIKAISFLDSTKSTHLPKLPGGFALRRKDCRTVVGRRIQCSAQAPPPAWPGSAFPEPGRRTWDGPKPISVVGSTGSIGTQTLDIVAENPDKFRVVALAAGSNVTLLVDQVKRFKPKLVAVRNESLVDELKEALSGLEDKPVIIPGEQGVIEVARHPDAVTVVTGIVGCAGLRPTVAAIEAGKDIALANKETLIAGGPFVLPLAHKHNVKILPADSEHSAIFQCIQGLPEGALRRIILTASGGAFRDWPVEKLKEVKVADALKHPNWSMGKKITVDSATLFNKGLEVIEAHYLYGADYDDIEIVIHPQSIIHSMIETQDSSVLAQLGWPDMRLPILYTMSWPERIYCSEVTWPRLDLCKLGSLTFKAPDNVKYPSMDLAYSAGRAGGTMTGVLSAANEKAVEMFIDEKISYLDIFKVVELTCAKHRAELVTSPSLEEIIHYDLWARDYAANLQNSTSSTPVFA* | FvH4_5g05300 | D |
| Fxa5Cg200440.1 | DXR3 | MEP | 472 | MALNLLSPAEIKAISFLDSTKSTHLPKFPGGFALRRKDCRTVIGRRIQCSAQAPPPAWPGSAFPEPGRRTWDGPKPISIVGSTGSIGTQTLDIVAENPDKFRVVALAAGSNVTLLEDQVKRFKPKLVAVRNESLVDELKEALSGLEDKPEIIPGEQGVIEVARHPDAVTVVTGIVGCAGLRPTVAAIEAGKDIALANKETLIAGGPFVLPLAHKHNVKILPADSEHSAIFQCIQGLPEGALRRIILTASGGAFRDWPVEKLKEVKVADALKHPNWSMGKKITVDSATLFNKGLEVIEAHYLYGADYDDIEIVIHPQSIIHSMIETQDSSVLAQLGWPDMRLPILYTMSWPERIYCSEVTWPRLDLCKLGSLTFKAPDNVKYPSMDLAYSAGRAGGTMTGVLSAANEKAVEMFIDEKISYLDIFKVVELTCAKHRAELVTSPSLEEIIHYDLWARDYAANLQNSTSSTPVFA* | FvH4_5g05300 | D |
| Fxa5Dg200439.1 | DXR4 | MEP | 472 | MAVNLLSPAEIKAISFLDSTKSTHLPKLPGGFALRRKDCRTVIGRRIQCSAQAPPPAWPGSAFPEPGRRTWDGPKPISIVGSTGSIGTQTLDIVAENPDKFRVVALAAGSNVTLLVDQVKRFKPKLVAVRNESLVDELKEALSGLEDKPEIIPGEQGVIEVSRHPDAVTVVTGIVGCAGLRPTVAAIEAGKDIALANKETLIAGGPFVLPLAHKHNVKILPADSEHSAIFQCIQGLPEGALRRIILTASGGAFRDWPVEKLKEVKVADALKHPNWSMGKKITVDSATLFNKGLEVIEAHYLYGADYDDIEIVIHPQSIIHSMIETQDSSVLAQLGWPDMRLPILYTMSWPERIYCSEVTWPRLDLCKLGSLTFKAP | FvH4_5g05300 | D |

| | | | | | | |
|---|---|---|---|---|---|---|
| | | | | DNVKYPSMDLAYSAGRAGGTMTGVLSAANEKAVEMFIDEKISYLDI<br>FKVVELTCAKHRAELVTSPSLEEIIHYDLWARDYAANLQNSTSSTPV<br>FA* | | |
| Fxa4Ag101691.1 | DXS1 | MEP | 717 | MALSTFSISTQKPTSKFSSHFSFGPALPCPQQQKLFNQVKKRTTGI<br>CASLSESGEYHSQRPPTPLLDTINYPIHMKNLSVKELKQLADELRS<br>DVIFNVSKTGGHLGSSLGVVELTVALHYVFNAPQDKILWDVGHQSY<br>PHKILTGRRDKMQTMRQTNGLAGFTKRSESEYDCFGTGHSSTTIS<br>AGLGMAVGRDLKGKNNHVVAVIGDGAMTAGQAYEAMNNAGYLDS<br>DMIIILNDNKQVSLPTANLDGPIPPVGALSSALSKLQSNRPLRELREV<br>AKVSILEAQGVTKQIGGSVHELAAKVDEYARGMISGSGSTLFEELG<br>LYYIGPVDGHNVDDLIAILQEVKTTQTTGPVLIHLITEKGRGYPYAEK<br>AADKYHGVAKFDPATGKQFKAAASTQSYTTYFAEALIAEAEADKDI<br>VAIHAAMGGGTGMNLFLRRFPTRCFDVGIAEQHAVTFAAGLACEG<br>LKPFCAIYSSFLQRAYDQVVHDVDLQKLPVRFAMDRAGLVGADGP<br>THCGAFDVTFMACLPNMVVMAPSDEAELFHMVATAAAIDDRPSCF<br>RYPRGNGIGVELPAGNKGTPLEIGKGRILIEGERVALLGYGSAVQT<br>CLAAATLVEPLGLSLTVADARFCKPLDHALIRKLAKSHEFLITVEEGS<br>IGGFGSHVVQFLALDGLLDGNLKWRPLVLPDRYIDHGAPADQLADA<br>GLTSSHIAATVLNMLGQTREALQVMS* | FvH4_4g16950 | E |
| Fxa4Bg101639.1 | DXS2 | MEP | 717 | MALSTFSISTQKPTSKFSSHFSFGPALSCPQQQKLFNQVKKRTTGI<br>CASLSESGEYHSQRPPTPLLDTINYPIHMKNLSVKELKQLADELRS<br>DVIFNVSKTGGHLGSSLGVVELTVALHYVFSAPQDKILWDVGHQSY<br>PHKILTGRREKMHTMRQTNGLAGFTKRSESEYDCFGTGHSSTTIS<br>AGLGMAVGRDLKGKNNHVVAVIGDGAMTAGQAYEAMNNAGYLDS<br>DMIIILNDNKQVSLPTANLDGPIPPVGALSSALSKLQSNRPLRELREV<br>AKVSILEAQGVTKQIGGSVHELAAKVDEYARGMISGSGSTLFEELG<br>LYYIGPVDAHNVDDLIAILQEVKTTQTTGPVLIHLITEKGRGYPYAEK<br>AADKYHGVAKFDPATGKQFKAAASTQSYTTYFAEALIAEAEADKDI<br>VAIHAAMGGGTGMNLFLRRFPTRCFDVGIAEQHAVTFAAGLACEG<br>LKPFCAIYSSFLQRAYDQVVHDVDLQKLPVRFAMDRAGLVGADGP<br>THCGAFDVTFMACLPNMVVMAPSDEAELFHMVATAAAIDDRPSCF<br>RYPRGNGIGVELPAGNKGTPLEIGKGRILIEGERVALLGYGSAVQT<br>CLAAATLVEPLGLSLTVADARFCKPLDHALIRKLAKSHEFLITVEEGS<br>IGGFGSHVVQFLALDGLLDGNLKWRPLVLPDRYIDHGAPADQLADA<br>GLTPSHIAATVLNMLGQTREALQVMS* | FvH4_4g16950 | E |
| Fxa4Cg201410.1 | DXS3 | MEP | 725 | MALSTFSISTQKSTSKFSSHFSFGPALSYPQQQKLFNQVKKRTTGI<br>CASLSESGEYHSQRPPTPLLDTINYPIHMKNLSVKELKQLADELRS<br>DVIFNVSKTGGHLGSSLGVVELTVALHYVFNAPQDKILWDVGHQSY<br>PHKILTGRREKMHTMRQTNGLAGFTKRSESEYDCFGTGHSSTTIS<br>AGLGMAVGRDLKGKNNHVVAVIGDGAMTAGQAYEAMNNAGYLDS<br>NMIIILNDNKQVSLPTANLDGPIPPVGALSSALSKLQSNRPLRELREV<br>AKVSILEAQGVTKQIGGSVHELAAKVDEYARGMISGSGSTLFEELG<br>LYYIGPVDGHNVDDLIAILQEVKTTQTTGPVLIHLITEKGRGYPYAEK<br>AADKYHGVAKFDPATGKQFKAAASTQSYTTYFAEALIAEAEADKDI<br>VAIHAAMGGGTGMNLFLRRFPTRCFDVGIAEQHAVTFAAGLACEG<br>LKPFCAIYSSFLQRAYDQVVHDVDLQKLPVRFAMDRAGLVGADGP<br>THCGAFDVTFMACLPNMVVMAPSDEAELFHMVATAAAIDDRPSCF<br>RYPRGNGIGVELPAGNKGTPLEIGKGRILIEGERVALLGYGSAVQT | FvH4_4g16950 | E |

| | | | | | | |
|---|---|---|---|---|---|---|
| | | | | CLAAATLVEPLGLSLTVADARFCKPLDHALIRKLAKSHEFLITVEEGS IGGFGSHVVQFLALDGLLDGNLKWRPLVLPDRYIDHGAPANQLADA GLTPSHIAATVLNMLGQTKRGTADNVMKGVLFSE* | | |
| Fxa4Dg101305.1 | DXS4 | MEP | 717 | MALSTFSISTQKPTSKFSSHFTFGPALSCPQQQKLFNQVKKRTTGI CASLSESGEYHSQRPPTPLLDTINYPIHMKNLSVKELKQLADELRS DVIFNVSKTGGHLGSSLGVVELTVALHYVFNAPQDKILWDVGHQSY PHKILTGRREKMHTMRQTNGLAGFTKRSESEYDCFGTGHSSTTIS AGLGMAVGRDLKGKNNYVVAVIGDGAMTAGQAYEAMNNAGYLDS DMIIILNDNKQVSLPTANLDGPIPPVGALSSALSKLQSNRPLRELREV AKVSILEAQGVTKQIGGSVHELAAKVDEYARGMISGSGSTLFEELG LYYIGPVDGHNVDDLIAILQEVKTTQTTGPVLIHLITEKGRGYPYAEK AADKYHGVAKFDPATGKQFKAAASTQSYTTYFAEALIAEAEADKDI VAIHAAMGGGTGMNLFLRRFPTRCFDVGIAEQHAVTFAAGLACEG LKPFCAIYSSFLQRAYDQVVHDVDLQKLPVRFAMDRAGLVGADGP THCGAFDVTFMACLPNMVVMAPSDEAELFHMVATAAAIDDRPSCF RYPRGNGIGVELPAGNKGTPLEIGKGRILIEGERVALLGYGSAVQT CLAAATLVEPLGLSLTVADTRFCKPLDHALIRKLAKSHEFLITVEEGS IGGFGSHVVQFLALDGLLDGNLKWRPLVLPDRYIDHGAPAHQLADA GLTYSHIAATVLNMLGQTREALQVMS* | FvH4_4g16950 | E |
| Fxa3Ag103964.1 | FPPS A1 | | 343 | MSNLKVKFLEVYSVLKSELINDPAFEFTDVSRQWIERMLDYNVPGG KLNRGLSVVDSLKLLKEGGELTDDEVFQSCALGWCIEWLQAYFLVL DDIMDGSHTRRGQPCWFRLPKIGMIAANDGIILRNHIPRILKKHFRV KPYYVDLLDLFNEVEFQTAHGQMIDLITTHEGEKDLSKYSLGIHRRI VQYKTAYYSFYLPVACALLMAGENLESHADVKNVLVEMGTYFQVQ DDYLDCFGDPEVIGKVGTDIQDFKCSWLVVKALELSNEEQKKLLHE NYGKDDQECIAKVKELYNALDLQGVFAEYESSSYDKITKSIEAHPSK AVQAVLKSFLAKIYKRLK* | FvH4_3g42290 | F |
| Fxa3Bg203613.1 | FPPS A2 | | 343 | MSNLKAKFLEVYSVLKSELINDPAFEFTDVSSQWIERMLDYNVPGG KLNRGLSVVDSLKLLKEGGELTDDEVFQSCALGWCIEWLQAYFLVL DDIMDGSHTRRGQPCWFRLPKIGMIAANDGIILRNHIPRILKKHFRV KPYYVDLLDLFNEVEFQTAHGQMIDLITTHEGEKDLSKYSLGIHRRI VQYKTAYYSFYLPVACALLMAGENLESHADVKNILIEMGTYFQVQD DYLDCFGDPEVIGKVGTDIQDFKCSWLVVKALELSNEEQKKLLHEN YGKDDRECIAKVKELYNALDLQGVFAEYESSSYNKITKSIEAHPSKA VQAVLKSFLAKIYKRLK* | FvH4_3g42290 | F |
| Fxa3Cg103545.1 | FPPS A3 | | 343 | MSNLKVKFLEVYSVLKSELINDPAFEFTDVSRQWIERMLDYNVPGG KLNRGLSVVDSLKLLKEGGELTDDEVFQSCALGWCIEWLQAYFLVL DDIMDGSHTRRGQPCWFRLPKIGMIAANDGIILRNHIPRILKKHFRV KPYYVDLLDLFNEVEFQTAHGQMIDLITTHEGEKDLSKYSLGIHRRI VQYKTAYYSFYLPVACALLMAGENLESHADVKNVLVEMGTYFQVQ DDYLDCFGDPEVIGKVGTDIQDFKCSWLVVKALELSNEEQKKLLHE NYGKDDQECIAKVKELYNALDLQGVFAEYESSSYDKITKSIEAHPSK AVQAVLKSFLAKIYKRLK* | FvH4_3g42290 | F |
| Fxa3Dg203403.1 | FPPS A4 | | 343 | MSNLKAKFLEVYSVLKSELINDPAFEFTDVSRQWIEQMLDYNVPGG KLNRGLSVVDSLKLLKEGGELTDDEVFQSCALGWCIEWLQAYFLVL DDIMDGSHTRRGQPCWFRLPKIGMIAANDGIILRNHIPRILKKHFRV KPYYVDLLDLFNEVEFQTAHGQMIDLITTHEGEKDLSKYSLGIHRRI VQYKTAYYSFYLPVACALLMAGENLESHADVKNVLIEMGTYFQVQ | FvH4_3g42290 | F |

| | | | | | | |
|---|---|---|---|---|---|---|
| | | | | DDYLDCFGDPEVIGKVGTDIQDFKCSWLVVKALELSNKEQKKLLHE NYGKDDQECIAKVKELYNALDLQGVFAEYESSSYDKITKSIEAHPSK AVQAVLKSFLAKIYKRLK* | | |
| Fxa3Ag100027.1 | FPPS B1 | | 343 | MADLRSKFMNVYSVLKSDLLEDPAFEFTDTSRQWVEQMLDYNVP GGKLNRGLSVIDSYQLLQHGRELTEDEIFLASALGWCIEWLQAFFL VLDDMMDGSHTRRGQPCWFRLPKVGLIAANDGVLLRNHIPRILKK HFRQKPYYVDLLDLFNEVEFQTASGQMIDLITTIDGEKDLSKYSLAIH RRIVQYKTAYYSFYLSVACALLMSGEELDKHIDVKNLLIDMGIYFQV QDDYLDCFGDPETIGKIGTDIEDFKCSWLVVKALELSNEEQKKTLH ENYGNPDPAKVAKVKALYKELDLQGVFAEYERQSYEKLTSSIEAHP SKAVQEVLKSFLGKIYKRLK* | FvH4_3g00240 | G |
| Fxa3Bg200029.1 | FPPS B2 | | 343 | MADLRSKFMNVYSVLKSELLEDPAFEFTDTSRQWVEQMLDYNVP GGKLNRGLSVIDSYQLLQHGRELTEDEIFLASALGWCIEWLQAFFL VLDDMMDGSHTRRGQPCWFRLPKVGLIAANDGVLLRNHIPRILKK HFRQKPYYVDLLDLFNEVEFQTASGQMIDLITTIDGEKDLSKYSLAIH RRIVQYKTAYYSFYLSVACALLMSGEELDKHIDVKNLLIDMGIYFQV QDDYLDCFGDPETIGKIGTDIEDFKCSWLVVKALELSNEEQKKTLH ENYGNPDPAKVAKVKALYKELDLQGVFAEYERLSYKKLTSSIEAHP SKAVQEVLKSFLGKIYKRQK | FvH4_3g00240 | G |
| Fxa3Cg100028.1 | FPPS B3 | | 343 | MADLRSKFMNVYSVLKSELLEDPAFEFTDTSRQWVEQMLDYNVP GGKLNRGLSVIDSYQLLQHGRELTEDEIFLASALGWCIEWLQAFFL VLDDMMDGSHTRRGQPCWFRLPKVGLIAANDGVLLRNHIPRILKK HFRQKPYYVDLLDLFNEVEFQTASGQMIDLITTIDGEKDLSKYSLAIH RRIVQYKTAYYSFYLSVACALLMSGEELDKHVDVKNLLIDMGIYFQV QDDYLDCFGDPETIGKIGTDIEDFKCSWLVVKALELSNEEQKKTLH ENYGNPDPAKVAKVKALYKELDLQGVFAEYERLSYEKLTSSIEAHP SKAVQEVLKSFLGKIYKRQK* | FvH4_3g00240 | G |
| Fxa3Dg200024.1 | FPPS B4 | | 343 | MADLRSKFMNVYSVLKSELLEDPAFEFTDTSRQWVEQMLDYNVP GGKLNRGLSVIDSYQLLQHGRKLTEDEIFLASALGWCIEWLQAFFL VLDDMMDGSHTRRGQPCWFRLPKVGLIAANDGVLLRNHIPRILKK HFRQKPYYVDLLDLFNEVEFQTASGQMIDLITTIDGEKDLSKYSLAIH RRIVQYKTAYYSFYLSVACALLMSGEELDKHVDVKNLLIDMGIYFQV QDDYLDCFGDPETIGKIGTDIEDFKCSWLVVKALELSNEEQKKTLH ENYGNPDPAKVAKVKALYKELDLQGVFAEYERLSYEKLMSSIEAHP SKAVQEVLKSFLGKIYKRLK* | FvH4_3g00240 | G |
| Fxa2Ag100784.1 | HDR1 | MEP | 462 | MAITLHLCRIPPRTDLFSSHSFAGSPCFRSRSSLSVKCSGDSSSSA AVESDFDAKVFRKNLTRSKNYNRRGFGHKEETLELMNREYTSDIIK KLKENGFEYTWGTVTVKLAEAYGFCWGVERAVQIAYEARKQFPDE RIWITNEIIHNPTVNKRLEEMEVKNIPIEEGRKQFEVVDKGDVVILPA FGAAVDEMLSLSERNVQIVDTTCPWVSKVWNTVEKHKKGDYTSIIH GKYAHEETVATASFAGKYVIVKNMDEARYVCDYILGGELDGSSSTR EAFLEKFKKAISPGFDPDRDLIKCGIANQTTMLKGETEDIGKLIEKTM MQKYGVENVNDHFVSFNTICDATQERQDAMYKLVEEKLDLLLVVG GWNSSNTSHLQEIAELRGIPSYWIDSEQRIGPGNRIAYKLNHGELV EKENWLPAGPVTIGVTSGASTPDKVVEDALIKVFDIKREEALQLA* | FvH4_2g07930 | H |
| Fxa2Bg200648.1 | HDR2 | MEP | 459 | MAITLHLCRIPPRTDLFPSHSLAGGPCFRSRSSLSVKCSGDAAAVE SDFDAKVFRKNLTRSKNYNRRGFGHKEETLELMNREYTSDIIKKLK ENGFEYTWGTVTVKLAEAYGFCWGVERAVQIAYEARKQFPDERIW | FvH4_2g07930 | H |

| | | | | | | |
|---|---|---|---|---|---|---|
| | | | | ITNEIIHNPTVNKRLEEMEVKNIPIEEGRKQFEVVDKGDVVILPAFGA AVDEMLSLSERNVQIVDTTCPWVSKVWNTVEKHKKGDYTSIIHGKY AHEETVATASFAGKYVIVKNMDEARYVCDYILGGELDGSSSTREAF LEKFKKAISPEFDPDRDLIKCGIANQTTMLKGETEDIGKLVEKTMMQ KYGVENVNDHFVSFNTICDATQERQDAMYKLVEEKLDLLLVVGGW NSSNTSHLQEIAELRGIPSYWIDSEQRIGPGNRIAYKLNHGELVEKE NWLPAGPVTIGVTSGASTPDKVVEDALTKVFDIKREEALQLT* | | |
| Fxa2Cg201772.2 | HDR3 | MEP | 459 | MAITLHLCHIPPRTDLFSSHSLAGGLCFRSRSSLSVKCSGDAAAVE SDFDAKVFRKNLTRSKNYNRRGFGHKEETLELMNREYTSDIIKKLK ENGFEYTWGTVTVKLAEAYGFCWGVERAVQIAYEARKQFPDERIW ITNEIIHNPTVNKRLEEMEVKNIPIEEGRKQFEVVDKGDVVILPAFGA AVDEMLSLSERNVQIVDTTCPWVSKVWNTVEKHKKGDYTSIIHGKY AHEETVATASFAGKYVIVKNMDEARYVCDYILGGELDGSSSTREAF LEKFKKAISPGFDPDRDLIKCGIANQTTMLKGETEDIGKLVEKTMMQ KYGVENVNDHFVSFNTICDATQERQDAMYKLVEEKLDLLLVVGGW NSSNTSHLQEIAELRGIPSYWIDSEQRIGPGNRIAYKLNHGELVEKE NWLPAGPVTIGVTSGASTPDKVVEDALIKVFDIKREEALQVA* | FvH4_2g07930 | H |
| Fxa2Dg200710.2 | HDR4 | MEP | 459 | MAITLHLCRIPPRTDLFSSHSLAGGPCFRSRSSLSVKCSGDAAAVE SDFDAKVFRKNLTRSKNYNRRGFGHKEETLELMNREYTSDIIKKLK ENGFEYTWGTVTVKLAEAYGFCWGVERAVQIAYEARKQFPDERIW ITNEIIHNPTVNKRLEEMEVKNIPIEEGRKQFEVVDKGDVVILPAFGA AVDEMLSLSERNVQIVDTTCPWVSKVWNTVEKHKKGDYTSIIHGKY AHEETVATASFAGKYVIVKNMDEARYVCDYILGGELDGSSSTREAF LEKFKKAISPGFDPDRDLIKCGIANQTTMLKGETEDIGKLVEKTMMQ KYGVENVNDHFVSFNTICDATQERQDAMYKLVEEKLDLLLVVGGW NSSNTSHLQEIAELRGIPSYWIDSEQRIGPGNRIAYKLNHGELVEKE NWLPAGPVTIGVTSGASTPDKVVEDALTKVFDIKREEALQLT* | FvH4_2g07930 | H |
| Fxa3Ag103081.1 | HDS1 | MEP | 741 | MATGAVPASFTGLTGRETTLGFTKSMDFVRVSDLKRFKSGRTRISV IRNSNPGSDIAELKPASEGSPLLVPRQKYCESIHKTVRRKTRTVMV GNVAIGSEHPIRIQTMTTSDTKEVSATVEEVMRIADKGADIVRITVQ GKKEADACFEIKNSLVQKNYNIPLVADIHFAPSVALRVAECFDKIRV NPGNFADRRAQFEKLEYTEEDYEKELEHIEQVFTPLVEKCKKYGRA MRIGTNHGSLSDRIMSYYGDSPRGMVESAFEFARICRKLDFHNFVF SMKASNPVIMVQAYRLLVAEMYIQGWDYPLHLGVTEAGEGEDGR MKSAIGIGTLLQDGLGDTIRVSLTEAPEEEIDPCKRLANLGMRAADL QQGVAPFEEKHRHYFDFQRRSGQLPVQKEGDEVDYRGVLHRDG SVLMSVSLDQLKTPELLYKSLAAKLVVGMPFKDLATVDSILLRQLPP VDDDDSRLALKRLIDVSMGVITPLSEQLTKPLPNAMVLVNTKELSSG AHKLLPEGTRLVVSLRGDEPYEVLDLIKRLDVTMLLHYLPYSEDKIS RVHAARRLFEYLGDNGLNYPVIHHIHFPSGIHRDDLVISAGTNVGAL LVDGLGDGLLLEAPDKDFNFIRNTSFNLLQGCRMRNTKTEYVSCPS CGRTLFDLQEISAQIREKTSHLPGVSIAIMGCIVNGPGEMADADFGY VGGAPGKIDLYVGKTVVKRAIEMEHATDALIQLIKDHGRWVDPPAE E* | FvH4_3g32460 | I |
| Fxa3Bg202785.1 | HDS2 | MEP | 741 | MATGAVPASFTGLTGRETTLGFTKSMGFVRVSDLKRFKSGRTRISV IRNSNPGSDIAELKPASEGSPLLVPRQKYCEYIHKTVRRKTRTVMV GNVAIGSEHPLRIQTMTTSDTKDVSATVEEVMRIADKGADIVRITVQ GKKEADACFEIKNSLVQKNYNIPLVADIHFAPSVALRVAECFDKIRV | FvH4_3g32460 | I |

| | | | | | | |
|---|---|---|---|---|---|---|
| | | | | NPGNFADRRAQFEKLEYTEEDYEKELEHIEQVFTPLVEKCKKYGRA MRIGTNHGSLSDRIMSYYGDSPRGMVESAFEFARICRKLDFHNFVF SMKASNPVIMVQAYRLLVAEMYVQGWDYPLHLGVTEAGEGEDGR MKSAIGIGALLQDGLGDTIRVSLTEAPEEEIDPCKRLANLGMRAADL QQGVAPFEEKHRHYFDFQRRSGQLPVQKEGDEVDYRGVLHRDG SVLMSVSLDQLKTPELLYKSLAAKLVVGMPFKDLATVDSILLRQLPP VDDDDSRLALKRLIDVSMGVITPLSEQLTKPLPNAIVLVNTKELSSG AHKLLPEGTRLVVSLRGDEPYEVLDLIKGLDVTMLLHDLPYSEDKIS RVHAARRLFEYLGDNGLNYPVIHHIQFPSGIHRDDLVISAGTNVGAL LVDGLGDGLLLEAPDKDFNFIRNTSFNLLQGCRMRNTKTEYVSCPS CGRTLFDLQEISAQIREKTSHLPGVSIAIMGCIVNGPGEMADADFGY VGGAPGKIDLYVGKTMVKRAIEMEHATDALIQLIKDHGRWVDPPAE E* | | |
| Fxa3Dg202713.1 | HDS3 | MEP | 741 | MATGAVPVSFTGLPGRETTLGLTKSMGFVRVSDLKRFKSGRTRISV IRNSNPGSDIAELKPASEGSPLLVPRQKYCEFIHKTVRRKTRTVMV GNVAIGSEHPIRIQTMTTSDTKDVSATVEEVMRIADKGADIVRITVQ GKKEADACFEIKNSLVQKNYNIPLVADIHFAPSVALRVAECFDKIRV NPGNFADRRAQFEKLEYTEEDYEKELEHIEQVFTPLVEKCKKHGRA MRIGTNHGSLSDRIMSYYGDSPRGMVESAFEFARICRKLDFHNFVF SMKASNPVIMVQAYRLLVAEMYVQGWDYPLHLGVTEAGEGEDGR MKSAIGIGTLLQDGLGDTIRVSLTEAPEEEIDPCKRLANLGMRAADL LQGVAPFEEKHRHYFDFQRRSGQLPVQKEGDEVDYRGVLHRDGS VLMSVSLDQLKTPELLYKSLAAKLVVGMPFKDLATVDSILLRQLPPV DDDDSRLALKRLIDVSMGVITPLSEQLTKPLPNAMVLVNMKELSSG THKLLPEGTRLVVSLRGDEPYEVLDLIKGLDVTMLLHDLPYREDNIS RVHAARRLFEYLGDNGLNYPVIHHIQFPSGIHRDDLVISAGTNVGAL LVDGLGDGLLLEAPDKDFDFIRNTSFNLLQGCRMRNTKTEYVSCPS CGRTLFDLQEISAQIREKTSHLPGVSIAIMGCIVNGPGEMADADFGY VGGAPGKIDLYVGKTMVKRAIEMEHATDALIQLIKDHGRWVDPPAE E* | FvH4_3g32460 | I |
| Fxa1Ag100908.1 | HMGR 1 | MVA | 563 | MDVRRRPPLPPRAGGVEPVKQHQRNHPKASDALPLPLYLTNGLFF TLFFTVAYFLLHRWREKIRSSTPLHVVTLSEIVALFSLFASVIYLLGFF GIDFVHPRAPHDTWNDVVRPTRRQNDVVKGSRPRGTTNINISHLS VPAMPEQEDEEIINSVVSGATPSYTLESNLGDCRRAAGIRREALQR TTRRSLEGLPLEGFDYDSILGQCCEMPIGYVQIPVGVAGPLLLNGE ELTVPMATTEGCLVASTNRGCKAIYASGGAHCVILKDGMTRAPVVR FNSAVRAAELKFFLEDPENFDSLAIVFSRSSRFAKLQGIKCAVAGKN LYMRFTCSNGDAMGMNMVSKGVQNVLDFLQSDFPDMDVIGISGN FCSDKKPAAVNWIEGRGKSVVCETVIKEEVVRKVLKTNVAALVELN MLKNLTGSAMAGALGGFNAHASNIVSAVYIATGQDPAQNVESSHCI TLMEAINNGKDLHVSVTMPSIEVGTVGGGTQLASQSACLNLLGVKG ASKESPGSNSRKLATIVAGSVLAGELSLMSALAAGQLVKSHMKYN RSSKDVSKLAC* | FvH4_1g09980 | J |
| Fxa1Bg200852.1 | HMGR 2 | MVA | 561 | MDVRRRPPLPPRAGGGEPVKQHQRNHPKASDALPLPLYLTNGLFF TLFFTVAYFLLHRWREKIRSSTPLHVVTLSEIVALFSLFASVIYLLGFF GIDFVQSFVHARAPHDPWNDVAKVMPRGHDDQLCGTTNIYHLSIP AMPEQEDEEIINSVVSGATPSYTLESNLGDCRRATAIRREALQRTT RRSLEGLPLEGFDYDSILGQCCEMPIGYVQIPVGVAGPLLLNGEEL | FvH4_1g09980 | J |

| | | | | | | |
|---|---|---|---|---|---|---|
| | | | | TVPMATTEGCLVASTNRGCKAIYASGGAHCVILKDGMTRAPVVRFNSAVRAAELKFFLEDPENFDSLAIVFNRSSRFAKLQGITCAVAGKNLYMRFTCSTGDAMGMNMVSKGVQNVLDFLQSDFPDMDVIGISGNFCSDKKPAAVNWIEGRGKSVVCEAVIKEEVVRKVLKTNVAALVELNMLKNLTGSAMAGALGGFNAHASNIVTAVYIATGQDPAQNVESSHCLTMMEAINDGKDLHVSVTMPSIEVGTVGGGTQLASQSACLNLLGAKGASKESPGSNSRKLATIVAGSVLAGELSLMSALAAGQLVKSHMKYNRSSKDVSKLAC* | | |
| Fxa1Cg100857.1 | HMGR 3 | MVA | 553 | MDVRRRPPLPPRAGGGEPVKQHQINHPKASDALPLPLPLYLTNGLFFTLFFTVAYFLLHRWREKIRSSTPLHVVTLSEIVALFSLFASVIYLLGFFGIDFVQSFVHPWNDVVKVPRGHDDQLCGTTHLSVPAMPEQEDEEIINSVVSGATPSYTLESNLGDCRRAAAIRREALQRTTRRSLEGLPLEGFDYDSILGQCCEMPIGYVQIPVGVAGPLLLNGEELTVPMATTEGCLVASTNRGCKAIYASGGAHCVILKDGMTRAPVVRFNSAVRAAELKFFLEDPENFDSLAIVFNRSSRFAKLQRITCAVAGKNLYMRFTCSTGDAMGMNMVSKGVQNVLDFLQSDFPDMDVIGISGNFCSDKKPAAVNWIEGRGKSVVCEAVIKEEVVRKVLKTNVAALVELNMLKNLTGSAMAGALGGFNAHASNIVTAVYIATGQDPAQNVESSHCLTMMEAINDGKDLHVSVTMPSIEVGTVGGGTQLASQSACLNLLGVKGASKVSPGSNSRKLATIVAGSVLAGELSLMSALAAGQLVKSHMKYNRSSKDVSKLAC* | FvH4_1g09980 | J |
| Fxa1Dg200784.1 | HMGR 4 | MVA | 551 | MDVRRRPPLPPRAGRVEPVKQHQINHPKASDALPLPLYLTNGLFFTLFFTVAYFLLHRWREKIRSSTPLHVVTLSEIVALFSLFASVIYLLGFFGIDFVQSFVHPWNDVVKVPRGHDDQLCGTTHLSVPAMPEQEDEEIINSVVSGATPSYTLESNLGDCRRAAAIRREALQRTTRRSLEGLPLEGFDYDSILGQCCEMPIGYVQIPVGVAGPLLLNGEELTVPMATTEGCLVASTNRGCKAIYVSGGAHCVILKDGMTRAPVVRFNSAVRAAELKFFLEDPENFDSLAIVFSRSSRFAKLQRITCAVAGKNLYMRFTCSTGDAMGMNMVSKGVQNVLDFLQSDFPDMDVIGISGNFCSDKKPAAVNWIEGRGKSVVCEAVIKEEVVRKVLKTNVAALVELNMLKNLTGSAMAGALGGFNAHASNIVTAVYIATGQDPAQNVESSHCLTMMEAINDGKDLHVSVTMPSIEVGTVGGGTQLASQSACLNLLGVKGASKESPGSNSRKLATIVAGSVLAGELSLMSALAAGQLVKSHMKYNRSSKDVSKLAC* | FvH4_1g09980 | J |
| Fxa2Ag101745.1 | HMGR 5 | MVA | 559 | MKVKFVDHEKDVVKASDALPLPLYFTNAVFFTLFFSVVYFLLTSWREKIRTNTPLHIIDLSEIVAIIAFVASFIYLLGFFGMDFVQSLILQPSNDIWAADDDEDIILKEDARKVPCAKAAPKVVPKVFEEKAMIETPPPTEEDEEIIKAVVAGTIPSYSLETKLGDCKRAASIRREALQRVTGKSLEGLPLEGFDYESILGQCCEMPVGYITIPVGIAGPLMLNGREFSVPMATTEGCLVASTNRGCKAINLSGGASSVLLKDGMTRAPVVRFNSAKRAAELKFFMEDPENYETISMVFNKSSRFGRLQTVKCAVAGKNLYMRFTCSTGDAMGMNMVSKGVQNVLDFLQNDFPDMDVIGISGNYCSDKKPAAVNWIEGRGKSVVCEAVIKGDIVTKVLKTNVAALVELNMLKNLTGSAIAGALGGFNAHASNIVSAVYLATGQDPAQNIESSHCITMMEAINDGKDLHVSVTMPSIEVGTVGGGTQLASQSACLNLLGVKGANREAPGTNARQLASVVAGSVLAGELSLMSAIAAGQLVKSHMKYNRSNKDVTTVTSA* | FvH4_2g17080 | K |

| | | | | | | |
|---|---|---|---|---|---|---|
| Fxa2Bg201569.1 | HMGR -Like | MVA | 558 | MKVKFVDDEKDVVKASDALPLPLYITNAVFFALFFSVVYFLLTSWRE KIRTNTPLHIINLSEIVAILAFVASFIYLLGFFGIDFVQSLILHPSNDIWA ADDALDFSITKMAPIASPEPAPKVAPKVFEEKTMIETPPPTAEDEGII KAVVAGTIPSYSLETKLGDCKRAASIRREALQRVTGKSLEGLPLEGF DYESILGQCCEMPVGYITIPVGIAGPLMLNGREFSVPMATTEGCLV ASTNRGCKAINLSGGASSVLLKDGMTRAPVVRFNSAKRAAELKFFL EDPENYETISMVFNKSSRFGRLQTVKCAVAGKNLYIRFTCSTGDAM GMNMVSKGVQNVLDFLQNDFPDMDVIGISGNYCSDKKPAAVNWIE GRGKSVVCEAVIKGDIVTKVLKTNVAALVELNMLKNLTGSAIAGALG GFNAHASNIVSAVYLATGQDPAQNIESSHCITMMEAINDGKDLHVS VTMPSIEVGTVGGGTQLASQSACLNLLGVKGANREAPGTNARQLA SVVAGSVLAGELSLMSAIAAGQLVKSHMKYNRSNKDVTAVASA* | FvH4_2g17080 | K |
| Fxa2Cg202539.1 | HMGR -like | MVA | 560 | MKVKFVDDEKDVVEASDALPLPLYITNAVFFALFFSVVYFLLTSWRE KIRTNTPLHIINLSEIVAILAFVASFIYLLGFFGIDFVQSLILQPSNDIWA ADDDEEEIIIKEDARKVPCAEPAPKVAPKVFEEKAMIETPPPTEEDE GIIKAVVAGTIPSYSLETKLGDCKRAASIRREALQRVTGKSLEGLPLE GFDYESILGQCCEMPVGYITIPVGIAGPLMLNGREFSVPMATTEGC LVASTNRGCKAINLSGGASSVLLKDGMTRAPVVRFNSAKRAAELKF FMEDPENYETISMVFNKSSRFGRLQTVKCAVAGKNLYMRFTCSTG DAMGMNMVSKGVQNVLDFLQNDFPDMDVIGISGNYCSDKKPAAV NWIEGRGKSVVCEAVIKGDIVSKVLKTNVAALVELNMLKNLTGSAIA GALGGFNAHASNIVSAVYLATGQDPAQNIESSHCITMMEAINDGKD LHVSVTMPSIEVGTVGGGTQLASQSACLNLLGVKGANREAPGTNA RQLASVVAGSVLAGELSLMSAIAAGQLVKSHMKYNRSNKDVTAVA SA* | FvH4_2g17080 | K |
| Fxa2Dg201512.1 | HMGR 8 | MVA | 557 | MKVKFVDDNEKDVVKASDALPLPLYITNAVFFALFFSVVYFLLTSWR EKIRTNTPLHIINLSEIVAILAFVASFIYLLGFFGIDFVQSLILQPSNDIW AAALDFSITKMAPIASPEPAPKVSPKVFEEKAMIETPPPTEEDEGIIK AVVAGTIPSYSLETKLGDCKRAASIRREALQRVTGKSLEGLPLEGF DYESILGQCCEMPVGYITIPVGIAGPLMLNGREFSVPMATTEGCLV ASTNRGCKAINLSGGASSVLLKDGMTRAPVVRFNSAKRAAELKFF MEDPENYETISMVFNKSSRFGRLQTVKCAVAGKNLYMRFTCSTGD AMGMNMVSKGVQNVLDFLQNDFPDMDVIGISGNYCSDKKPAAVN WIEGRGKSVVCEAVIKGDIVSKVLKTNVVALVELNMLKNLTGSAIAG ALGGFNAHASNIVSAVYLATGQDPAQNIESSHCITMMEAINDGKDL HVSVTMPSIEVGTVGGGTQLASQSACLNLLGVKGANREAPGTNAR QLASVVAGSVLAGELSLMSAIAAGQLVKSHMKYNRSNKDVTAVAS A* | FvH4_2g17080 | K |
| Fxa2Bg201570.1 | HMGR 6 | MVA | 558 | MVDHHHHHQNDAAKASDALPLPLYLTNAVFFTLFFSVVYFLLTRWR EKIRTSTPLHMVNLSEIVAILAFVASFIYLLGFFGIDFVQSLILRPSHD VWAVEDDEEALDCSVPKIAPLKPALKVAQKVFDKEVVTETAPPSAE DEEIIKAVVAGTIPSYSLETKLGDCKRAASIRREALQRVTGKSLEGLP LEGFDYESILGQCCEMPVGYVTIPVGVAGPIMLDGREFSVPMATTE GCLVASTNRGCKAINLSGGASSVLLKDGMTRAPVVRFNSAKRAAE LKFFMEDPENADTIATLFNKSSRFGRLQSVKCAVAGKNLYMRFCC STGDAMGMNMVSKGVQNVLDFLQDDFPDMDVIGISGNYCSDKKP AAVNWIEGRGKSVVCEALIKGDIVTKVLKTNVAALVELNMLKNLTGS AIAGALGGFNAHASNIVSAVYLATGQDPAQNIESSHCITMMEAINDG | | L |

| | | | | | | |
|---|---|---|---|---|---|---|
| | | | | KDLHVSVTMPSIEVGTVGGGTQLASQSACLNLLGVKGANREAPGA NARQLASVVAASVLAAELSLMSAIAAGQLVNSHMRYNRSSRDVSA MVSA* | | |
| Fxa2Cg202536.1 | HMGR 7 | MVA | 573 | MEQTRKPMKMKVKFAENEEAKVVKASDALPLPLYLTNAVFFTLFFS VAYFLLTSWREKIRSHTPLHVLNLSEIVSILVLVASFVYLLGFFGIDFV QTPSLKDEEEDIIVQEDARIVPCGQALECPRPQIAGIASPKPAPKAIV EKPAMETPPPTAEDEEIIKAVVAGTIPSYSLESKLGDCKRAASIRRE ALQRITGKSLEGLPLEGFDYQSILGQCCEMPVGYVTIPVGIAGPLML DGREFSIPMATTEGCLVASTNRGCKAINLSGGASSVLLKDGMTRAP VVRFNSARRAAELKFYMEDPENYETISMVFNKSSRFGRLQSVQCA VAGKNLYMRFCCSTGDAMGMNMVSKGVQNVLDFLQNDFPDMDVI GISGNYCSDKKPAAVNWIEGRGKSVVCEVVIKGDVVTKVLKTNVAA LVELNMLKNLTGSAIAGALGGFNAHASNIVSAVYLATGQDPAQNIES SHCLTMMEAINDGKDLHVSVTMPSIEVGTVGGGTQLASQSACLNL LGVKGANREAPGSNARQLATVVAGSVLAGELSLMSAIAAGQLVKS HMKYNRSSKDVTTVSSA* | | M |
| Fxa5Ag200128.1 | HMGS 1 | MVA | 463 | MAKNVGILAVDIYFPPSCIQQEALEAHDGASKGKYTIGLGQDCMAF CSEVEDVISMRYLTVVTNLLEKYGVDPKQIGRLEVGSETVIDKSKSI KTFLMQIFEKHGNTDIEGVDSTNACYGGTAALFNCVNWVESSSWD GRYGLVVCTDSAVYAEGPARPTGGAAAIAMLIGPDAPIAFESKIRGS HMSHVYDFYKPDLASEYPVVDGKLSQTCYLMALDSCYKQLCQKYE KIEGKQFSISDAEYFVFHSPYNKLVQKSFARLVFQDSLRNACSIDEA AKEKFAPFASLSSDESYASRDLEKVAQQVAKPLYDAKVQPTTLVPK QVGNMYTASLYAAFASLLHNKHDSLDGKRVILFSYGSGSTATMFSL QLRAGQHPFSLKNIAAVMNVGEKLKSRHEFAPEKFCEIMKIMEHRY GGKDFVTNKDISLLLPGTFYLTEVDCKYRRFYAQKDCAENGVVAN GH* | FvH4_5g01350 | N |
| Fxa5Bg100136.1 | HMGS 2 | MVA | 463 | MAKNVGILAVDIYFPPSCIQQEALEAHDGASKGKYTIGLGQDCMAF CSEVEDVISMRYLTVVTNLLEKYGVDPKQIGRLEVGSETVIDKSKSI KTFLMQIFEKHGNTDIEGVDSTNACYGGTAALFNCVNWVESSSWD GRYGLVVCTDSAVYAEGPARPTGGAAAIAMLIGPDAPIAFESKIRGS HMSHVYDFYKPDLASEYPVVDGKLSQTCYLMALDSCYKQLCKKYE KIEGKQFSISDAEYFVFHSPYNKLVQKSFARLVFQDSLRNACSIDEA AKEKFAPFASLSSDESYASRDLEKVAQQVAKPLHEAKVQPTTLVPK QVGNMYTASLYAAFASLLHNKHDSLDGKRVILFSYGSGSTATMFSL QLRAGQHPFSLKNIAAVMNIGEKLKSRHEFAPEKFCEIMKIMEHRY GGKDFVTNKDISLLLPGTFYLTEVDSKYRRFYAQKDGAENGVVAN GH* | FvH4_5g01350 | N |
| Fxa5Cg200112.1 | HMGS 3 | MVA | 464 | MAKNVGILAVDIYFPPSCIQQEVLEAHDGASKGKYTIGLGQDCMAF CSEVEDVISMRYLTVVTNLLEKYGVDPKQIGRLEVGSETVIDKSKSI KTFLMQIFEKHGNTDIEGVDSTNACYGGTAALFNCVNWVESSSWD GRYGLVVCTDSAVYAEGPARPTGGAAAIAMLIGPDAPIAFESKIRGS HMSHVYDFYKPDLASEYPVVDGKLSQTCYLMALDSCYKQLCKKYE KIEGKQFSISDAEYFVFHSPYNKLVQKSFARLVFQDSLRNACSIDEA AKEKFAPFASLSSDESYASRDLEKVAQQVAKPLYDAKVQPTTLVPK QVGNMYTASLYAAFASLLHNKHDSLDGKRVILFSYGSGSTATMFSL QLRAGQHPFSLKNIAAVMNVGEKLKSRHEFAPEKFCEIMKIMEHRY | FvH4_5g01350 | N |

| | | | | | | |
|---|---|---|---|---|---|---|
| | | | | GGKDFVTNKDISLLLPGTFYLTEVDSKYRRFYAQKEDGAENGVVAN<br>GH* | | |
| Fxa5Dg200121.1 | HMGS<br>4 | MVA | 463 | MAKNVGILAVDIYFPPSCIQQEALEAHDGASKGKYTIGLGQDCMAF<br>CSEVEDVISMRYLTVVTNLLEKYGVNPKQIGRLEVGSETVIDKSKSI<br>KTFLMQIFEKHGNTDIEGVDSTNACYGGTAALFNCVNWVESSSWD<br>GRYGLVVCTDSAVYAEGPARPTGGAAAIAMLIGPDAPIAFESKIRGS<br>HMSHVYDFYKPDLASEYPVVDGKLSQTCYLMALDSCYKQLCKKYE<br>KIEGKQFSISDAEYFVFHSPYNKLVQKSFARLVFQDSLRNACSIDEA<br>AKEKFAPFASLSSDESYANRDLEKVAQQVAKPLYDAKVQPTTLVPK<br>QVGNMYTASLYAAFASLLHNKHDSLDGKRVILFSYGSGSTATMFSL<br>QLRAGQHPFSLKNIAAVMNVGEKLKSRHEFAPEKFCEIMKIMEHRY<br>GGKDFVTNKDISLLLPGTFYLTEVDSKYRRFYAQKDGAENGVVAN<br>GH* | FvH4_5g01350 | N |
| Fxa6Ag100050.1 | IDI1 | | 302 | MSFTATRGLLNTATRATLSSLFFPKSHLLRLFSSSYSTFASRRPLSL<br>AKPFPSLLPARSSPSLRISTMADAPDAGMDAVQRRLMFEDECILVD<br>ENDRVVGHDTKYNCHLMEKIEAENLLHRAFSVFLFNSKNELLLQQR<br>SATKVTFPLVWTNTCCSHPLYRESELIDEECLGVRNAAQRKLFDEL<br>GIPAEDVPVDQFIPLGRILYKAPSDGKWGEHELDYLLFTVRDVSVH<br>PNPDEVADIKYVNQEELKELLRKVDAGEEGLKLSPWFRLVVDNFLF<br>KWWDHVEKCTIKEAADMKTIHRLT* | FvH4_6g00470 | O |
| Fxa6Bg100046.1 | IDI2 | | 235 | MADAPDAGMDAVQRRLMFEDECILVDENDRVVGHDTKYNCHLME<br>KIEAENLLHRAFSVFLFNSKNELLLQQRSATKVTFPLVWTNTCCSH<br>PLYRESELIDEECLGVRNAAQRKLLDELGIPAEDVPVDQFIPLGRML<br>YKAPSDGKWGEHELDYLLFTVRDVSVHPNPDEVADIKYVNQEELK<br>ELLRKADAGEEGLKLSPWFRLVVDNFLFKWWDHVEKRTIKEAADM<br>KTIHRLT* | FvH4_6g00470 | O |
| Fxa6Cg100047.1 | IDI3 | | 302 | MSFTATSGLLNTATRATLSSLFFPKSHLLRLFSSSYSTFASRRPLSL<br>AKPFPSLLPARSSPSLRISTMADAPDAGMDAVQRRLMFEDECILVD<br>ENDRVVGHDTKYNCHLMEKIEAENLLHRAFSVFLFNSKNELLLQQR<br>SATKVTFPLVWTNTCCSHPLYRESELIDEECLGVRNAAQRKLLDEL<br>GIPAEDVPVDQFIPLGRMLYKAPSDGKWGEHELDYLLFTVRDVSVH<br>PNPDEVADIKYVNQEELKELLRKADAGEEGLKLSPWFRLVVDNFLF<br>KWWDHVEKCTIKEAADMKTIHRLT* | FvH4_6g00470 | O |
| Fxa6Dg100047.1 | IDI4 | | 303 | MSFTSTTSGLLNTATRATLSSLFFPKSHLPRLFSSSYSTFASRRPLS<br>LSKPFPSLLPARPSPSLRISTMADAPDAGMDAVQRRLMFEDECILV<br>DENDRVVGHDTKYNCHLMEKIEAENLLHRAFSVFLFNSKNELLLQQ<br>RSATKVTFPLVWTNTCCSHPLYRESELIDEECLGVRNAAQRKLLDE<br>LGIPAEDVPVDQFIPLGRMLYKAPSDGKWGEHELDYLLFTVRDVSV<br>HPNPDEVADIKYVNQEELKELLRKADAGEEGLKLSPWFRLVVDNFL<br>FKWWDHVEKRTIKEAADMKTIHRLT* | FvH4_6g00470 | O |
| Fxa3Ag102669.1 | IPK1 | | 366 | MAFSQFVLGPSPCPPNRGRRLRSNLMVCGRRQVEIVYDPDERINK<br>LADKVEVSRLSLFSPCKINVFLRITNKRDDGFHDLASLFHVISLGDVI<br>KFSLSPSKSKDRLSTNVSGVPLDDRNLIIKALNLYRKKTGTHNFFWI<br>HLDKKVPTGAGLGGGSSNAATALWAANQFSGSLATEKELQEWSS<br>EIGSDVPFFFSQGAAYCTGRGEIVQNIPPPVPFDIPMVLIKPQQACS<br>TAEVYKRLQLDTTNNVDPLKLLERISVNGISQDVCINDLEPPAFQVL<br>PSLKRLKQRVLAASRGQYDAVFMSGSGSTIVGIGSPDPPQFVYDE<br>DEYKDVFLSEANFLTREENHWYREPSSRSASGQSSDLSESIP* | FvH4_3g27650 | P |

| | | | | | | |
|---|---|---|---|---|---|---|
| Fxa3Bg202352.1 | IPK2 | | 370 | MAFSQFVLGPSPCPPKLFENRRRRLRSNLMVCGRRQVEIVYDPDE RINKLADKVELSRLSLFSPCKINVFLRITNKRDDGFHDLASLFHVISL GDVIKFSLSPSKSKDRLSTNVSGVPLDDRNLIIKALNLYRKKTGTHN FFWIHLDKKVPTGAGLGGGSSNAATALWAANQFSGSLATEKELQE WSSEIGSDVPFFFSQGAAYCTGRGEIVQNIPPPVPFDIPMVLIKPQQ ACSTAEVYKRLQLDTTNNVDPLKLLERISVNGISQDVCINDLEPPAF QVLPSLKRLKQRVLAASRGQYDAVFMSGSGSTIVGIGSPDPPQFV YDEDEYKDVFLSEANFLTREENHWYREPSSRSASGQSSDLSESIP* | FvH4_3g27650 | P |
| Fxa3Dg202308.1 | IPK3 | | 370 | MAFSQFVLGPGPCPPKLFENRRRRLHSNLMVCGRRQVEIVYDPDE RINKLADKVELSRLSLFSPCKINVFLRITNKRDDGFHDLASLFHVISL GDVIKFSLSPSKSKDRLSTNVSGVPLDDRNLIIKALNLYRKKTGTLN FFWIHLDKKVPTGAGLGGGSSNAATALWAANQFSGSLATEKELQE WSSEIGSDVPFFFSQGAAYCTGRGEIVQNIPPPVPFDIPMVLIKPQQ ACSTAEVYKRLQLDTTNNVDPLKLLERISVNGISQDVCINDLEPPAF QVLPSLKRLKQRVLAASRGQYDAVFMSGSGSTIVGIGSPDPPQFV YDEDEYKDVFLSEANFLTREENHWYREPSSRSASGQSSDLSESIP* | FvH4_3g27650 | P |
| Fxa1Ag101298.1 | PT1 | | 361 | MSCVNLSSWGQACSMLNRSRSSPSLLLLHPTRPVSFSSHKRRRP SSVAAILTKKESAVNEESTKPSINFKTYMVEKAESVNRALDAAVSLK DPVTIHEAMRYSLLAGGKRVRPVLCLAACELVGGDESVAMPAACA VEMIHTMSLIHDDLPCMDNDDLRRGKPTNHKVFGEDVAVLAGDAL LSFAFEHLAVSTVGVEPSRIVRAVEELARSIGSEGLVAGQVVDIHSE GLSDVGLEQLEYIHIHKTAALLECAVVLGSILGGGSDTEIEKLRTFAR YIGLLFQVVDDILDVTKSSQELGKTAGKDLLADKVTYPKLMGIEKSR EFAEKLNRDAQEQLVGFNSKKAAPLIALANYIAYRQN* | FvH4_1g14070 | Q |
| Fxa1Bg201207.1 | PT2 | | 360 | MSCVNLSSWGQACSMLNRSRSSPSLLLLHPTRPFSFAPPIRRRSS VAAILTKQESAVNEESAKPSFNFKTYMVEKAESVNRALDAAVSLKD PVTIHEAMRYSLLAGGKRVRPVLCLAACELVGGDESVAMPAACAV EMIHTMSLIHDDLPCMDNDDLRRGKPTNHKVFGEDVAVLAGDALL SFAFEHLAVSTVGVEPSRLVRAVEELARSIGSEGLVAGQVVDIHSE GLSDVGLEHLEYIHLHKTAALLECAVVLGSILGGGSDSEIEKLRTFA RNIGLLFQVVDDILDVTKSSQELGKTAGKDLLADKVTYPKLMGIEKS REFAEKLNRNAQEQLVGFDSEKAAPLIALANYISYRQN* | FvH4_1g14070 | Q |
| Fxa1Cg101268.1 | PT3 | | 361 | MSCVNLSSWGHACSMLNRSRSSPSLLLLHPTRPVSFSSHKRSRPS SVAAILTKQESAVNEESVKPGFNFKIYMVEKAESVNRALDAAVSLK DPVTIHEAMRYSLLAGGKRVRPVLCLAACELVGGDESVAMPAACA VEMIHTMSLIHDDLPCMDNDDLRRGKPTNHKVFGEDVAVLAGDAL LSFAFEHLAVSTVGVVPSRIVRAVEELARSIGSEGLVAGQVVDIHSE GLSDVGLEHLEYIHLHKTAALLECAVVLGSILGGGSDIEIEKLRTFAR YIGLLFQVVDDILDVTKSSQELGKTAGKDLLADKVTYPKLMGIEKSR EFAEKLNRDAQEQLVGFDSEKAAPLIALANYIAYRQN* | FvH4_1g14070 | Q |
| Fxa1Dg201159.1 | PT4 | | 361 | MSCVNLSSWGQACSMLNRSRSSPSLLLLHPTRPVSFSSHKRRRP SSVAAILTKQESAVNEESVKPSFNFKTYMVEKAESVNRALDAAVSL KDPVTIHEAMRYSLLAGGKRVRPVLCLAACELVGGDESVAMPAAC AVEMIHTMSLIHDDLPCMDNDDLRRGKPTNHKVFGEDVAVLAGDA LLSFAFEHLAGSTVGVEPSRIVRAVEELARSIGSEGLVAGQVVDIHS EGLSDVGLEHLEYIHLHKTAALLECAVVLGSILGGGSDIEIEKLRTFA RYIGLLFQVVDDILDVTKSSQELGKTAGKDLLADKVTYPKLMGIEKS REFAEKLNRDAQEQLVGFDSEKAAPLIALANYIAYRQN* | FvH4_1g14070 | Q |

| | | | | | | |
|---|---|---|---|---|---|---|
| Fxa2Ag101616.1 | PT5 | | 330 | MAFSVVSTPAHFHLPKKLTFRIRCCAASSVSTRSESTRFDLKTYWT TLIADINSKLNEAVPVRYPELIYESMRYSVLADGAKRASPVMCVAAC ELFGGDRLAAFPTACALEMVHAASLIHDDLPCMDDDSSRRGQPSN HTVYGEDMAILAGDALFPLGFQHIVSNTPSNLVPEARLLRVITEIART VGSTGMAAGQFLDLEGGPNAVEFVQEKKFGEMGECSAVCGGLLA GAEDEEVDRLRRYGRAVGVLYQVVDDILEEKKNGKDENEKKEKKG KSYVKVYGVEKAMEVAERLRAQAKQELDGFEKYGDGVVPLHSFV DYAVDRSFTL* | FvH4_2g15810 | R |
| Fxa2Bg201429.1 | PT6 | | 331 | MAFSVVSTPAHFHLPKKLTFRIRCCAASSVSTRSKSTRFDLKTYWT TLIADINSKLNEAVPVRYPELIYESMRYSVLADGAKRASPVMCFAAC ELFGGDRLAAFPTACALEMVHAASLIHDDLPCMDDDPSRRGQPSN HTIYGEDMAILAGDALFPLGFQHIVSNTPSNLVPEARLLRVITEIART VGSTGMAAGQFLDLEGGPNAVEFVQEKKFGEMGKCSAVCGGLLA GAEDDEVDRLRRYRRAVGVLYQVVDDILEEKKKNGKDENEKKGKK GKSYVKVYGVEKALEVAERLRAQAKLELDGFEKYGDGVVPLHSFV DYAVDRSFTL* | FvH4_2g15810 | R |
| Fxa2Cg202421.1 | PT7 | | 331 | MAFSVVSTPAHFHLPKKLTFRIRCCAASSVSTRSKLTRFDLKTYWT TLIADINSKLNEAVPVRYPELIYESMRYSVLADGAKRASPVMCVAAC ELFGGDRLAAFPTACALEMVHAASLIHDDLPCMDDDPSRRGQPSN HTIYGEDMAILAGDALFPLGFQHIVSNTPSNLVPEARLLRVITEIART VGSTGMAAGQFLDLEGGPHAVEFVQEKKFGEMGECSAVCGGLLA GAEDEEVDRLRRYGRAVGVLYQVVDDILEETKKNGKDENEKKKKK GKSYVKVYGVEKALEVAERLRAQAKLELDGFEKYGDGVVPLHSFV DYAVDRSFTL* | FvH4_2g15810 | R |
| Fxa2Dg201392.1 | PT8 | | 331 | MAFSVVSTPAHFHLPKKITFRIRCCAASSVSSRSKSTRFDLKTYWT TLIADINSKLNEAVPVRYPELIYESMRYSVLADGAKRASPVMCVAAC ELFGGDRLAAFPTACALEMVHAASLIHDDLPCMDDDPSRRGQPSN HTIYGEDMAILAGDALFPLGFQHIVSNTPSNLVPEARLLRVITEIART VGSTGMAAGQFLDLEGGPNAVEFVQEKKFGVMGECSAVCGGLLA GAEDEEVDRLRRYGRAVGVLYQVVDDILEEKKKNGKDENEKKEKK GKSYVKVYGVEKALEVAERLRAQAKLELDGFEKYGDGVVPLHSFV DYAVDRSFTL* | FvH4_2g15810 | R |

**Supplemental Table 2:** Amino acid sequence similarity matrices of upstream terpene biosynthetic genes.

**AACT- acetoacetyl-CoA thiolase**

| AACT1 | Fxa3Ag100033.2 | Fxa3Bg200036.1 | Fxa3Cg100033.2 | Fxa3Dg200030.2 |
|---|---|---|---|---|
| Fxa3Ag100033.2 | | 73.3 | 69.1 | 69.5 |
| Fxa3Bg200036.1 | 73.3 | | 90.1 | 91 |
| Fxa3Cg100033.2 | 69.1 | 90.1 | | 98 |
| Fxa3Dg200030.2 | 69.5 | 91 | 98 | |

| AACT2 | Fxa6Ag103901.1 | Fxa6Bg103557.1 | Fxa6Cg103460.1 | Fxa6Dg103364.1 |
|---|---|---|---|---|
| Fxa6Ag103901.1 | | 99 | 99.3 | 99 |
| Fxa6Bg103557.1 | 99 | | 99.8 | 99.5 |
| Fxa6Cg103460.1 | 99.3 | 99.8 | | 99.8 |
| Fxa6Dg103364.1 | 99 | 99.5 | 99.8 | |

***CMS- 4-diphosphocytidyl-2-C-methyl-D-erythritol synthase***

| CMS | Fxa5Ag202521.1 | Fxa5Bg102434.2 | Fxa5Cg202277.2 | Fxa5Dg202343.1 |
|---|---|---|---|---|
| Fxa5Ag202521.1 | | 81.4 | 96 | 77.8 |
| Fxa5Bg102434.2 | 81.4 | | 82.3 | 65.5 |
| Fxa5Cg202277.2 | 96 | 82.3 | | 76 |
| Fxa5Dg202343.1 | 77.8 | 65.5 | 76 | |

**DXR- 1-deoxy-D-xylulose 5-phosphate reductoisomerase**

| DXR | Fxa5Ag200475.1 | Fxa5Bg100449.1 | Fxa5Cg200440.1 | Fxa5Dg200439.1 |
|---|---|---|---|---|
| Fxa5Ag200475.1 | | 99.2 | 99.4 | 99.4 |
| Fxa5Bg100449.1 | 99.2 | | 98.9 | 98.9 |
| Fxa5Cg200440.1 | 99.4 | 98.9 | | 99.2 |
| Fxa5Dg200439.1 | 99.4 | 98.9 | 99.2 | |

**DXS- 1-deoxy-D-xylulose 5-phosphate synthase**

| DXS | Fxa4Ag101691.1 | Fxa4Bg101639.1 | Fxa4Cg201410.1 | Fxa4Dg101305.1 |
|---|---|---|---|---|

| | | | | |
|---|---|---|---|---|
| Fxa4Ag101691.1 | | 99.2 | 97.1 | 98.9 |
| Fxa4Bg101639.1 | 99.2 | | 97.4 | 99 |
| Fxa4Cg201410.1 | 97.1 | 97.4 | | 97.1 |
| Fxa4Dg101305.1 | 98.9 | 99 | 97.1 | |

**FPPS- farnesyl diphosphate synthase**

| FPPSA | Fxa3Ag103964.1 | Fxa3Bg203613.1 | Fxa3Cg103545.1 | Fxa3Dg203403.1 |
|---|---|---|---|---|
| Fxa3Ag103964.1 | | 98.3 | 100 | 98.8 |
| Fxa3Bg203613.1 | 98.3 | | 98.3 | 98.3 |
| Fxa3Cg103545.1 | 100 | 98.3 | | 98.8 |
| Fxa3Dg203403.1 | 98.8 | 98.3 | 98.8 | |

| FPPSB | Fxa3Ag100027.1 | Fxa3Bg200029.1 | Fxa3Cg100028.1 | Fxa3Dg200024.1 |
|---|---|---|---|---|
| Fxa3Ag100027.1 | | 98.5 | 98.8 | 98.5 |
| Fxa3Bg200029.1 | 98.5 | | 99.1 | 98.3 |
| Fxa3Cg100028.1 | 98.8 | 99.1 | | 99.1 |
| Fxa3Dg200024.1 | 98.5 | 98.3 | 99.1 | |

**HDS- 4-hydroxy-3-methylbut-2-enyl diphosphate synthase**

| HDS | Fxa3Ag103081.1 | Fxa3Bg202785.1 | Fxa3Dg202713.1 |
|---|---|---|---|
| Fxa3Ag103081.1 | | 98.5 | 97.6 |
| Fxa3Bg202785.1 | 98.5 | | 98.1 |
| Fxa3Dg202713.1 | 97.6 | 98.1 | |

**HMGR- 3-hydroxy-3-methylglutaryl-CoA reductase**

| HMGR1 | Fxa1Ag100908.1 | Fxa1Bg200852.1 | Fxa1Cg100857.1 | Fxa1Dg200784.1 |
|---|---|---|---|---|
| Fxa1Ag100908.1 | | 93.5 | 92.3 | 92.8 |
| Fxa1Bg200852.1 | 93.5 | | 96.4 | 96.3 |
| Fxa1Cg100857.1 | 92.3 | 96.4 | | 98.7 |
| Fxa1Dg200784.1 | 92.8 | 96.3 | 98.7 | |

| HMGR2 | Fxa2Ag101745.1 | Fxa2Bg201569.1 | Fxa2Cg202539.1 | Fxa2Dg201512.1 |
|---|---|---|---|---|
| Fxa2Ag101745.1 | | 94.5 | 97 | 94.5 |
| Fxa2Bg201569.1 | 94.5 | | 96.1 | 98 |
| Fxa2Cg202539.1 | 97 | 96.1 | | 96.3 |
| Fxa2Dg201512.1 | 94.5 | 98 | 96.3 | |

**HMGS- 3-hydroxy-3-methylglutaryl-CoA synthase**

| HMGS | Fxa5Ag200128.1 | Fxa5Bg100136.1 | Fxa5Cg200112.1 | Fxa5Dg200121.1 |
|---|---|---|---|---|
| Fxa5Ag200128.1 | | 98.7 | 98.9 | 98.9 |
| Fxa5Bg100136.1 | 98.7 | | 98.9 | 98.9 |
| Fxa5Cg200112.1 | 98.9 | 98.9 | | 99.1 |
| Fxa5Dg200121.1 | 98.9 | 98.9 | 99.1 | |

**IDI- isopentenyl diphosphate isomerase**

| IDI | Fxa6Ag100050.1 | Fxa6Bg100046.1 | Fxa6Cg100047.1 | Fxa6Dg100047.1 |
|---|---|---|---|---|
| Fxa6Ag100050.1 | | 76.5 | 98.7 | 96.7 |
| Fxa6Bg100046.1 | 76.5 | | 77.5 | 77.6 |
| Fxa6Cg100047.1 | 98.7 | 77.5 | | 98 |
| Fxa6Dg100047.1 | 96.7 | 77.6 | 98 | |

**IDS- 4-hydroxy-3-methylbut-2-enyl diphosphate reductase**

| IDS | Fxa2Ag100784.1 | Fxa2Bg200648.1 | Fxa2Cg201772.2 | Fxa2Dg200710.2 |
|---|---|---|---|---|
| Fxa2Ag100784.1 | | 97.6 | 97.8 | 98.1 |
| Fxa2Bg200648.1 | 97.6 | | 98.5 | 99.6 |
| Fxa2Cg201772.2 | 97.8 | 98.5 | | 98.9 |
| Fxa2Dg200710.2 | 98.1 | 99.6 | 98.9 | |

**IPK- isopentenyl monophosphate kinase**

| IPK | Fxa3Ag102669.1 | Fxa3Bg202352.1 | Fxa3Dg202308.1 |
|---|---|---|---|
| Fxa3Ag102669.1 | | 98.4 | 97.6 |
| Fxa3Bg202352.1 | 98.4 | | 99.2 |

| | | |
|---|---|---|
| Fxa3Dg202308.1 | 97.6 | 99.2 |

**PT- Prenyltransferase (Geranyl Diphosphate Synthase/Geranylgeranyl diphosphate synthase)**

| PT1 | Fxa1Ag101298.1 | Fxa1Bg201207.1 | Fxa1Cg101268.1 | Fxa1Dg201159.1 |
|---|---|---|---|---|
| Fxa1Ag101298.1 | | 95 | 96.4 | 97.5 |
| Fxa1Bg201207.1 | 95 | | 95.3 | 96.4 |
| Fxa1Cg101268.1 | 96.4 | 95.3 | | 98.3 |
| Fxa1Dg201159.1 | 97.5 | 96.4 | 98.3 | |

| PT2 | Fxa2Ag101616.1 | Fxa2Bg201429.1 | Fxa2Cg202421.1 | Fxa2Dg201392.1 |
|---|---|---|---|---|
| Fxa2Ag101616.1 | | 96.7 | 97.3 | 97.3 |
| Fxa2Bg201429.1 | 96.7 | | 97.6 | 97.6 |
| Fxa2Cg202421.1 | 97.3 | 97.6 | | 97.9 |
| Fxa2Dg201392.1 | 97.3 | 97.6 | 97.9 | |

**Supplemental Table 3** Terpene synthase sequences identified in this study. 64 Genomic *F. × ananassa (Fa)* MSTP S; 12 Genomic *F. × ananassa* diTPS; 4 additional *F. × ananassa* transcripts; 5 additional *F. chiloensis (Fc)* transcripts; 5 additional *F. virginiana (Fvr)* transcripts; 4 additional *F. vesca (Fv)* transcripts. Additionally discarded pseudogenes are included from synteny analysis. Functionally tested candidates are shown in bold.

| **FaRR1 Genome Hit** | **Type** | **Name** | **Synteny Group** | **Species** | ***F. vesca Hit*** | **RRX8W** | **DDXXD** | **AA** | Tested? |
|---|---|---|---|---|---|---|---|---|---|
| Fxa1Ag100480 | M/S TPS | *FaTPS1* | a | *F. x ananassa* | FvH4_1g05400 | RRTANFKSPVW | DDMYD | 557 | No |
| Fxa1Ag100363 | Syntenic Psuedogene | (Fragment) | b | *F. x ananassa* | FvH4_1g04000 | NA | DDIYD | 425 | |
| Fxa1Bg200351 | M/S TPS | *FaTPS2* | b | *F. x ananassa* | FvH4_1g04000 | RRSANYHPSIW | DDIYD | 560 | No |
| Fxa1Dg200965 | M/S TPS | *FaTPS3* | c | *F. x ananassa* | - | NA | DDIFD | 557 | No |
| Fxa2Ag102276 | diTPS | *FaCPS6* | d | *F. x ananassa* | - | NA | DIDD | 563 | No |
| Fxa2Ag102357 | diTPS | *FaCPS7* | e | *F. x ananassa* | FvH4_2g23400 | NA | DLDD | 809 | No |
| Fxa2Cg203098 | Syntenic Psuedogene | (Fragment) | e | *F. x ananassa* | FvH4_2g23400 | NA | | 113 | |
| Fxa2Ag102359.1 | diTPS | *FaCPS1* | f | *F. x ananassa* | FvH4_2g23440 | NA | DIDD | 818 | No |
| Fxa2Bg202200.1 | diTPS | *FaCPS2* | f | *F. x ananassa* | FvH4_2g23440 | NA | DIDD | 818 | No |
| Fxa2Cg203102.1 | diTPS | *FaCPS3* | f | *F. x ananassa* | FvH4_2g23440 | NA | DIDD | 818 | No |
| Fxa2Dg202041.1 | diTPS | *FaCPS4* | f | *F. x ananassa* | FvH4_2g23440 | NA | DIDD | 818 | No |
| Fxa2Bg200794 | M/S TPS | *FaTPS4* | g | *F. x ananassa* | - | RPIANFQPSIW | DDIYD | 582 | No |
| Fxa2Ag100915 | Syntenic Psuedogene | (Fragment) | h | *F. x ananassa* | - | NA | DDIYD | 408 | |
| Fxa2Bg200800 | M/S TPS | *FaTPS5* | h | *F. x ananassa* | - | RPIANFQPSIW | DDIYD | 615 | No |
| **Fxa2Bg200800.1** | M/S TPS | ***FvrTPS5*** | h | *F. virginiana* | **NA** | **RPIANFQPSIW** | **DDIYD** | **498** | **Yes** |
| Fxa3Ag100266.1 | M/S TPS | *FaTPS6* | i | *F. x ananassa* | FvH4_3g03000 | RRSANYKPNIW | DDVYD | 584 | No |

| | | | | | | | | | |
|---|---|---|---|---|---|---|---|---|---|
| **Fxa3Ag100266.1** | M/S TPS | ***FvTPS6*** | i | *F. vesca* | **FvH4_3g03000** | **RRSANYKPNIW** | **DDVYD** | **583** | **Yes** |
| **Fxa3Ag100266.1** | M/S TPS | ***FaTPS7*** | i | *F. x ananassa* | **FvH4_3g03000** | **RRSANYKPNIW** | **DDVYD** | **584** | **Yes** |
| **Fxa3Bg200271.1** | M/S TPS | ***FaTPS10*** | i | *F. x ananassa* | **FvH4_3g03000** | **RRSANYKPNIW** | **DDVYD** | **568** | **Yes** |
| Fxa3Dg200246.1 | M/S TPS | *FaTPS21* | i | *F. x ananassa* | FvH4_3g03000 | RRSANYKPNIW | DDVYD | 580 | No |
| Fxa3Ag100275.1 | M/S TPS | *FaTPS8* | j | *F. x ananassa* | FvH4_3g03061 | NA | DDIFD | 522 | No |
| Fxa3Bg200281.1 | M/S TPS | *FaTPS12* | j | *F. x ananassa* | FvH4_3g03061 | RRGIAEDSLLP | DDIFD | 581 | No |
| Fxa3Dg200253.1 | M/S TPS | *FaTPS22* | j | *F. x ananassa* | FvH4_3g03061 | NA | DDIFD | 509 | No |
| **Fxa3Ag101306** | diTPS | ***FaKSL1*** | k | *F. x ananassa* | **FvH4_3g14116** | **NA** | **DDFFD** | **797*** | **Yes** |
| **Fxa3Bg201184** | diTPS | ***FaKSL2*** | k | *F. x ananassa* | **FvH4_3g14116** | **NA** | **DDFFD** | **797*** | **Yes** |
| **Fxa3Cg101173** | diTPS | ***FaKSL3*** | k | *F. x ananassa* | **FvH4_3g14116** | **NA** | **DDFFD** | **774*** | **Yes** |
| **Fxa3Dg2011124** | diTPS | ***FaKSL4*** | k | *F. x ananassa* | **FvH4_3g14116** | **NA** | **DDFFD** | **787*** | **Yes** |
| **Fxa3Dg201124** | diTPS | ***FcKSL5*** | k | *F. chiloensis* | **FvH4_3g14116** | **NA** | **DDLFD** | **571** | **Yes** |
| Fxa3Ag102016.1 | M/S TPS | *FaTPS9* | l | *F. x ananassa* | FvH4_3g21390 | NA | DDVYD/DDIFD | 1095 | No |
| Fxa3Bg201803 | Syntenic Psuedogene | (Fragment) | l | *F. x ananassa* | FvH4_3g21390 | NA | DDIFD | 329 | |
| Fxa3Cg101786.1 | M/S TPS | *FaTPS19* | l | *F. x ananassa* | FvH4_3g21390 | NA | DDVYD/DDIFD | 1019 | No |
| Fxa3Dg201726.1 | M/S TPS | *FaTPS23* | l | *F. x ananassa* | FvH4_3g21390 | NA | DDIFD | 557 | No |
| Fxa3Bg200279.1 | M/S TPS | *FaTPS11* | m | *F. x ananassa* | - | RRGIAEDSLLP | DDIFD | 585 | No |
| Fxa3Bg200284.1 | M/S TPS | *FaTPS13* | n | *F. x ananassa* | - | NA | DDIFD | 485 | No |
| Fxa3Bg200286.1 | M/S TPS | *FaTPS14* | o | *F. x ananassa* | - | NA | DDIFD | 522 | No |
| Fxa3Bg201829.1 | M/S TPS | *FaTPS15* | p | *F. x ananassa* | FvH4_3g21490 | NA | DDIYD | 484 | No |
| Fxa3Cg100132.1 | M/S TPS | *FaTPS16* | q | *F. x ananassa* | FvH4_3g01590 | RRSVNFKPSIW | DDMYD | 562 | No |

| | | | | | | | | | |
|---|---|---|---|---|---|---|---|---|---|
| **Fxa3Cg100266.2** | M/S TPS | ***FvrTPS17*** | r | *F. virginiana* | **(FvH4_3g03041)** | **RRGIAEDSLLP** | **DDIFD** | **495** | **Yes** |
| **Fxa3Cg100266.2** | M/S TPS | ***FaTPS17*** | r | *F. x ananassa* | **(FvH4_3g03041)** | **NA** | **DDIFD** | **519** | **Yes** |
| **Fxa3Cg100266.3** | M/S TPS | ***FcTPS18*** | r | *F. chiloensis* | **(FvH4_3g03041)** | **RRGIAEDSLLP** | **DDIFD** | **580** | **Yes** |
| **Fxa3Cg100266.3** | M/S TPS | ***FaTPS18*** | r | *F. x ananassa* | **(FvH4_3g03041)** | **RRGIAEDSLLP** | **DDIFD** | **580** | **Yes** |
| Fxa3Ag102106 | Syntenic Psuedogene | (Fragment) | s | *F. x ananassa* | FvH4_3g22162 | NA | DDIYD | 268 | |
| Fxa3Cg101875.1 | M/S TPS | *FaTPS20* | s | *F. x ananassa* | FvH4_3g22162 | RRSANFHPIW | DDIYD | 562 | No |
| **Fxa3Dg201123.1** | diTPS | ***FvrCPS5*** | t | *F. virginiana* | **Fv4_3g14111** | **NA** | **DLDD** | **800** | **Yes** |
| **Fxa3Dg201123.1** | diTPS | ***FaCPS5*** | t | *F. x ananassa* | **Fv4_3g14111** | **NA** | **DLDD** | **800** | **Yes** |
| Fxa4Ag102687 | Syntenic Psuedogene | (Fragment) | v | *F. x ananassa* | FvH4_4g27943 | RNKIVAEYFW | DDVYD | 311 | |
| Fxa4Bg102623 | Syntenic Psuedogene | (Too long) | v | *F. x ananassa* | FvH4_4g27943 | RPANYAPCMW | DDVYD | 879 | |
| Fxa4Cg202367.1 | M/S TPS | *FaTPS24* | v | *F. x ananassa* | FvH4_4g27943 | RPANYAPCMW | DDVYD/DDVYD | 1050 | No |
| Fxa4Dg102144.1 | M/S TPS | *FaTPS26* | v | *F. x ananassa* | FvH4_4g27943 | RPANYAPCIW | DDVYD | 564 | No |
| Fxa4Dg102142.1 | M/S TPS | *FaTPS25* | w | *F. x ananassa* | - | RCSPHYTPGIE | DDVYD | 560 | No |
| Fxa5Ag200582.1 | M/S TPS | *FaTPS27* | x | *F. x ananassa* | FvH4_5g06470 | RRSANFSPSVW | DDTFD | 582 | No |
| Fxa5Bg100567.1 | M/S TPS | *FaTPS29* | x | *F. x ananassa* | FvH4_5g06470 | RRSANFSPSVW | DDTFD | 572 | No |
| Fxa5Cg200524 | M/S TPS | *FaTPS31* | x | *F. x ananassa* | FvH4_5g06470 | RRSANFPPSVW | DDTFD | 570 | No |
| Fxa5Dg200537 | M/S TPS | *FaTPS34* | x | *F. x ananassa* | FvH4_5g06470 | RRSADFSPSVW | DDTFD | 573 | No |
| **Fxa5Dg200537.1** | M/S TPS | ***FaTPS35*** | x | *F. x ananassa* | **FvH4_5g06470** | **RRSADFSPSVW** | **DDTFD** | **570** | **Yes** |
| **Fxa5Dg200537.1** | M/S TPS | ***FvTPS35*** | x | *F. vesca* | **FvH4_5g06470** | **RRSADFSPSVW** | **DDTFD** | **570** | **Yes** |
| **Fxa5Dg200537.1** | M/S TPS | ***FaTPS36*** | x | *F. x ananassa* | **FvH4_5g06470** | **RRSADFSPSVW** | **DDTFD** | **570** | **Yes** |
| Fxa5Ag203430.1 | M/S TPS | *FaTPS28* | y | *F. x ananassa* | FvH4_5g35711 | LAYYEYAPCIW | DDIYD | 586 | No |

| | | | | | | | | | |
|---|---|---|---|---|---|---|---|---|---|
| Fxa5Bg103205 | M/S TPS | *FaTPS30* | y | *F. x ananassa* | FvH4_5g35711 | LAYYEHAPCIW | DDIYD | 579 | No |
| Fxa5Cg202929 | M/S TPS | *FaTPS32* | z | *F. x ananassa* | FvH4_5g35700 | RPSADYTPSIW | DDIYD | 555 | No |
| Fxa5Dg200534 | M/S TPS | *FaTPS33* | aa | *F. x ananassa* | - | RRSANFSPSIW | DDVYD | 553 | No |
| Fxa6Ag100563 | M/S TPS | *FaTPS37* | bb | *F. x ananassa* | FvH4_6g06540 | RPIANFQPSIW | DDIYD | 579 | No |
| **Fxa6Ag100563.1** | M/S TPS | ***FvTPS37*** | bb | *F. vesca* | **FvH4_6g06540** | **RPIANFQPSIW** | **DDIYD** | **569** | **Yes** |
| Fxa6Bg100527 | M/S TPS | *FaTPS46* | bb | *F. x ananassa* | FvH4_6g06540 | RPIANFQPSIW | DDIYD | 580 | No |
| Fxa6Ag101058 | M/S TPS | *FaTPS38* | cc | *F. x ananassa* | FvH4_6g11440 | PRPANYAPCIW | DDVYD | 673 | No |
| Fxa6Bg100956 | Syntenic Psuedogene | (Fragment) | cc | *F. x ananassa* | FvH4_6g11440 | NA | DDVYD | 392 | |
| Fxa6Cg100917 | M/S TPS | *FaTPS47* | cc | *F. x ananassa* | FvH4_6g11440 | PRPANYAPCIW | DDVYD | 564 | No |
| Fxa6Dg100848 | M/S TPS | *FaTPS49* | cc | *F. x ananassa* | FvH4_6g11440 | PRPANYAPCIW | DDVYD | 564 | No |
| Fxa6Ag101852 | M/S TPS | *FaTPS39* | dd | *F. x ananassa* | FvH4_6g19050 | RRTANYHPSIW | DDIYD | 564 | No |
| Fxa6Cg101643 | M/S TPS | *FaTPS48* | dd | *F. x ananassa* | FvH4_6g19050 | RRTAIYHPSIW | DDIYD | 750 | No |
| Fxa6Dg101605 | M/S TPS | *FaTPS50* | dd | *F. x ananassa* | FvH4_6g19050 | RRTANFKPSVW | DDMYD | 558 | No |
| **Fxa6D101605.1** | M/S TPS | ***FvTPS50*** | dd | *F. vesca* | **FvH4_6g19050** | **RRTANFKPSVW** | **DDMYD** | **556** | **Yes** |
| Fxa6Ag104161 | M/S TPS | *FaTPS40* | ee | *F. x ananassa* | FvH4_6g43720 | RRLAQYHPTIW | DDMYD | 614 | No |
| **Fxa6Ag104161.1** | M/S TPS | ***FcTPS40*** | ee | *F. chiloensis* | **FvH4_6g43720** | **RRLAQYHPTIW** | **DDMYD** | **602** | **Yes** |
| **Fxa6Ag104161.1** | M/S TPS | ***FvrTPS40*** | ee | *F. virginiana* | **FvH4_6g43720** | **RRLAQYHPTIW** | **DDMYD** | **604** | **Yes** |
| Fxa6Bg103821 | Syntenic Psuedogene | (Fragment) | ee | *F. x ananassa* | FvH4_6g43720 | NA | DDMYD | 395 | |
| Fxa6Cg103709 | Syntenic Psuedogene | (Fragment) | ee | *F. x ananassa* | FvH4_6g43720 | RRLAQYHPTIW | NA | 334 | |
| Fxa6Dg103609 | M/S TPS | *FaTPS51* | ee | *F. x ananassa* | FvH4_6g43720 | RRLAQYHPTIW | DDMYD | 630 | No |
| Fxa6Ag104162 | M/S TPS | *FaTPS41* | ff | *F. x ananassa* | - | RRLAQYHPTIW | DDMYD | 589 | No |
| **Fxa6Ag104162.1** | M/S TPS | ***FaTPS42*** | ff | *F. x ananassa* | **-** | **RRLAQYHPTIW** | **DDMYD** | **580** | **Yes** |

| | | | | | | | | | |
|---|---|---|---|---|---|---|---|---|---|
| Fxa6Ag104168 | M/S TPS | *FaTPS43* | gg | *F. x ananassa* | - | RRTADYKPSIW | DDVYD/DDVYD | 1067 | No |
| Fxa6Ag104224 | M/S TPS | *FaTPS44* | hh | *F. x ananassa* | - | TRTADYKPSIW | DDVYD | 572 | No |
| Fxa6Cg103766 | Syntenic Psuedogene | (Fragment*)* | hh | *F. x ananassa* | - | RTADYKPSIW | DDVYD | 363 | |
| Fxa6Ag104324 | M/S TPS | *FaTPS45* | ii | *F. x ananassa* | - | RVTPDYKPSIW | DDIYD | 563 | No |
| Fxa6Bg103826 | Syntenic Psuedogene | (Fragment) | jj | *F. x ananassa* | FvH4_6g43820 | NA | DDVYD | 259 | |
| Fxa6Dg103617 | M/S TPS | *FaTPS52* | jj | *F. x ananassa* | FvH4_6g43820 | RRTTDYKPSIW | DDVYD | 578 | No |
| **Fxa6Dg103617.2** | M/S TPS | ***FcTPS52*** | jj | *F. chiloensis* | **FvH4_6g43820** | **RRTTDYKPSIW** | **DDVYD** | **577** | **Yes** |
| Fxa6Dg103674 | M/S TPS | *FaTPS53* | kk | *F. x ananassa* | - | TRTAYYKPSIW | DDVYD | 566 | No |
| Fxa7Ag200397 | M/S TPS | *FaTPS54* | ll | *F. x ananassa* | FvH4_7g03620 | RRVANFSPSVW | DDIYD | 528 | No |
| Fxa7Bg200423 | M/S TPS | *FaTPS57* | ll | *F. x ananassa* | FvH4_7g03620 | RRVANFSPSVW | DDIYD | 564 | No |
| Fxa7Cg100355 | M/S TPS | *FaTPS59* | ll | *F. x ananassa* | FvH4_7g03620 | RRVANFSPSVW | DDIYD | 510 | No |
| Fxa7Dg100343 | M/S TPS | *FaTPS62* | ll | *F. x ananassa* | FvH4_7g03620 | RRVANFSPSVW | DDIYD | 565 | No |
| **Fxa7Ag203289.1** | M/S TPS | ***FvrTPS55*** | mm | *F. virginiana* | **FvH4_7g33640** | **PAYYEYAPCIW** | **DDIYD** | **561** | **Yes** |
| **Fxa7Ag203289.1** | M/S TPS | ***FaTPS55*** | mm | *F. x ananassa* | **FvH4_7g33640** | **PAYYEYAPCIW** | **DDIYD** | **561** | **Yes** |
| **Fxa7Bg203106** | M/S TPS | ***FcTPS58*** | mm | *F. chiloensis* | **FvH4_7g33640** | **PAYYEYAPCIW** | **DDIYD** | **561** | **Yes** |
| Fxa7Bg203106 | M/S TPS | *FaTPS58* | mm | *F. x ananassa* | FvH4_7g33640 | LAYYEYAPCIW | DDIYD | 568 | No |
| Fxa7Cg103018 | M/S TPS | *FaTPS61* | mm | *F. x ananassa* | FvH4_7g33640 | LAYYEYAPCIW | DDIYD | 584 | No |
| Fxa7Dg102851 | M/S TPS | *FaTPS64* | mm | *F. x ananassa* | FvH4_7g33640 | LAYYEYAPGIW | DDIYD | 568 | No |
| Fxa7Ag203294 | M/S TPS | *FaTPS56* | nn | *F. x ananassa* | FvH4_7g33760 | PAYYEYAPCIW | DDIYD | 1910 | No |
| Fxa7Bg203107 | Syntenic Psuedogene | (Too long, no DDxxD) | nn | *F. x ananassa* | FvH4_7g33760 | RRKFTGQTVA | NA | 1347 | |
| Fxa7Cg103019 | Syntenic Psuedogene | (Too long, no DDxxD) | nn | *F. x ananassa* | FvH4_7g33760 | NA | NA | 951 | |

| | | | | | | | | | |
|---|---|---|---|---|---|---|---|---|---|
| Fxa7Dg102852 | Syntenic Psuedogene | (Too long, no DDxxD) | nn | *F. x ananassa* | FvH4_7g33760 | RRKFTGQTVA | NA | 1355 | |
| Fxa7Cg101251 | M/S TPS | *FaTPS60* | oo | *F. x ananassa* | FvH4_7g13030 | RRSTNFKLSIW | DDIYD | 536 | No |
| Fxa7Dg102096 | M/S TPS | *FaTPS63* | pp | *F. x ananassa* | - | RRSTNFKPSIW | DDIYD | 472 | No |

**Supplemental Table 4:** Sequences of identified TPS genes. Tested candidates shown in bold.

| FaRR1 Genome Hit | Type | Name | Species | Clade | % similarity to genomic | Source | AA Sequence |
|---|---|---|---|---|---|---|---|
| Fxa2Ag102276 | diTPS | *FaCPS6* | *F. x ananassa* | c | - | FaRR1 Genomic | MMHDVVTTLLYSLEGMAGLDWEKLLKLQ<br>SRDGSFLCSPASTAYALQQTKDQNCMSY<br>LLSVVQYFNGGVPNVYLVDLFERMWAVD<br>RLQRLGLSRYFEPEIKECLEYVSRYWTEK<br>GISFVRNSEVPDIDDTSMGFRLLRLHGHK<br>VSAEVYEHFRKGSDFFCFPGQSNQAVTG<br>IYNLYRASQVALPGEKILGDAKEFAAKFLR<br>EKQASNELLDKWIITKDLPGEVDYALEIP<br>WYASLPRLETRFYIEQYGGENDVWIGKT<br>LYSMPHVNNNVYLELAILDYNNCQALHLK<br>EWDSIQKWYKEWKFEDYGVSKKSFLTAY<br>FVAAASIFEPERANERLAWAKMACLVDTI<br>VSYFKEETSDEDNKKAFVDEFRSFSNMQ<br>GYENARRSYTNKIAGQGIVRALLETISQLS<br>LVTMVLHGQDINESLYQAWEKWLLKWQ<br>QKEESYQDEAELLVETINRTAGLSLRKEL<br>SNYPEYEQLFGLTNKVCNQLRSYRNQKN<br>KVNDNGDCKTKINMTTPEIESYMQQLVE<br>MVLEKFLDDTKLLIKQSFFTVTRSFYYSTY<br>CDHEVISHHINKVLFQRAV* |
| Fxa2Ag102357 | diTPS | *FaCPS7* | *F. x ananassa* | c | - | FaRR1 Genomic | MSSHTTHHLVLSFQTSLPSSSSSSGSIPS<br>GFSGDEDRRVLPLRSSAVSKPHTRYDAP<br>VLHKIVRDDTQQNEETPLTKDDSAVSAST<br>QIIKKHVETVKSMLSSMEDGSISISAYDTA<br>WIALAEDVNGSASPQFPSSLEWIANNQL<br>EDGSWGYKDLFSAHDRLISTLACVVALKL<br>WNLHADKCHKGMKFFKENLHKLEDENS<br>EHIIVGFEVTFPSLLEIARSLNLDLPDDTPV<br>LHDIHARRDLKLSKIPRDILHNVATSLLHS<br>LEGMAGLDWEKLLKLQSEDGSILFSPAST<br>AYALQQTKDRNCMSYLSRLVHKFNGGAP<br>NVYPVDLFERLWAVDRLQRLGLSRYLEP<br>EIKECMNYVSRYWTEKGISWVRNSELPD<br>LDDTSMGFRLLRLHGHKVSADVFEHFKK<br>GSEFFCHPGELNNAVTVIHNLYRASQVAF<br>PGEKILGDAKEFATKFLRKKQASNEFTDK<br>WIIAKDLPGEVGYALDFPWYASLPRLETR<br>FYIEQYGGEDDVWIAKTLYNMPCISNNLY |

| | | | | | | | |
|---|---|---|---|---|---|---|---|
| | | | | | | | LELAKLDYNDCQTLHLKEWNSIQKWYKE<br>WKLGDYGLSEKSLLTAYFVAAASIFEPER<br>ANERLAWAKTTCLVDTIMSFKGETDDKY<br>KKAFVDEFKTFSTMRDHENARRLYKNKF<br>AGQGIVGALLATISQLSSATMVLHRQDITE<br>SLCQAWGKWLMKWQGTSKRYQDEAELL<br>VKIINQTAGLQNELLSNDPEYEQLFSLINK<br>VCNQLRFYRNQKIKVGDNGNCGTKINMT<br>TPEIESYMQQLVGLVLEKSSDDTVSLIKQ<br>TFFTVARSFYYATCCDPEVISHHINKVLFQ<br>RAI* |
| Fxa2Ag102359.1 | diTPS | *FaCPS1* | *F. x ananassa* | c | - | FaRR1 genomic | MASHSTHLLFLSLSSSLSPASSSFANHNQ<br>PKHVPGLWLFGAKNKRVLPSICNAVSKP<br>HTQDYPDVIKWRENPTIVGDYTQQDEEA<br>LPTKDDDSATTEIIREHVETVKSMLSSME<br>DGEISVSAYDTAWVSLVEDVNGTGSPQF<br>PSSLEWIANNQLEDGSWGYEDIYNAHDR<br>MISTLACVVALKSWNLHPDKCEKGIKFFK<br>ENLHKLEDENIEHMPIGFEVAFPSVLEIAR<br>SLKLDVPDDSPVLHKIYACRNLKLTKIPKD<br>ILHNVPTTLLHSMEGMAGLDWEKLLKLQ<br>CQDGSFLFSPASTAYALQQTKDQKCMSY<br>LSRAVRKFNGGVPNVYPVDLFEHMWAV<br>DRLQRLGLYRYFKPEIKECMNYVAKFWT<br>EKGICWARNSEVQDIDDTAMGFRLLRLH<br>GHKVSSDVFKHFKKGDKFFCFAGQSNQ<br>AVTGMYNLYRASQVALPGEKTLYEAKEF<br>STKFLREKQASNELLDKWIIMKDLPGEVE<br>YALDVPWYASLPRLETKVYIEQYGGGDD<br>VWIGKSLYSMPYVNNNVYLELAKLDFNN<br>CQALHLSEWDDLQKWYGEWKLGGYGLS<br>KKSLLMAYFVAAASIFEPERANERLAWAK<br>TTSLVDTIESYFKGETSHEDKKAFVDEFK<br>NLSNMREYEITRRSNKIAGQGILGALLATL<br>SQLSLGTMVLHGQDIIQCLRQSWVKWLL<br>KWLERGGMYQDEAELLVETINQTAGLSP<br>TQGLFSNNPEYEHIFSLTNKVCNQLRSC<br>QKQKHKVNDDGCYKSKMSMTTQAIESD<br>MQQLVELMLKSPDGIESLLKQSFFTVARS<br>FYYLNYCDPEIISHHIDKVLFERAI* |
| Fxa2Bg202200.1 | diTPS | *FaCPS2* | *F. x ananassa* | c | - | FaRR1 genomic | MASHSTHLLFLSLSSSLSPASSSFFNHNQ<br>PKHVPGLSLFGAKDKRVLPSICNAVSKPH<br>TQDYPDVIKWRENPTIVGDYTQQDEEAL<br>PTKDDDSATTEIIREHVETVKSMLSSMED |

| | | | | | | | |
|---|---|---|---|---|---|---|---|
| | | | | | | | GEISVSAYDTAWVALVEDVNGTGSPQFP<br>SSLEWIANNQLEDDSWGYEDIYNAHDRII<br>STLACVVALKTWNLHPDKCEKGIKFFKEN<br>LQKLEDENIEHMPIGFEVAFPSVLEIARSL<br>KLDVPDNSPVLHKIYACRNLKLTKIPKDIL<br>HNVPTTLLHSMEGMAGLDWEKLLKLQCQ<br>DGSFLFSPASTAYALQQTKDQKCMSYLS<br>RAVRKFNGGVPNVYPVDLFEHMWAVDR<br>LQRLGLYRYFEPEIKECMNYVAKFWTEK<br>GICWARNSEVQDIDDTAMGFRLLRLHGH<br>KVSSDVFKHFKKGNEFFCFAGQSNQAVT<br>GMYNLYRASQVALPGEKTLYEAKEFSTK<br>FLREKQASNELLDKWIIMKDLPGEVEYAL<br>DVPWYASLPRLETKVYIEQYGGGDDVWI<br>GKSLYSMPYVNNNVYLELAKLDFNNCQA<br>LHLSEWDNLQKWYGEWKLGGYGLSKKS<br>LLMAYFVAAASIFEPERANERLAWAKTAS<br>LVDTIESYFKGETSHEDKKTFVDEFKNLS<br>NMREYEIARRSNKIAGQGILGALLATLGQ<br>LSLGTMVLHGQDITQSLRQTWVKWLMK<br>WLERGGMYQDEAELLVETINQTAGLSPT<br>QGLFSNNPEYEHIFSLTNKVCNQLRSCQ<br>KQKHKVNDDGSYESKMNMTTQAIESDM<br>QQLVELVLKSPDGIESLLNQSFFTVARSF<br>YYFNYCDPEIISHHIDKVLFERAI* |
| Fxa2Cg203102.1 | diTPS | *FaCPS3* | *F. x ananassa* | c | - | FaRR1 genomic | MASHSTHLLFLSLSSSLSPASSSFFNHNQ<br>PKHVPGLSLFGAKDKRVLPSICNAVSKPH<br>TQDYPDVIKWRENPTIVGDYTQQVEEAL<br>PTKDDDSATTEIIREHVETVKSMLSSMED<br>GEISVSAYDTAWVALVEDVNGTGSPQFP<br>SSLEWIANNQLEDDSWGYEDIYNAHDRII<br>STLACVVALKTWNLHPDKCEKGIKFFKEN<br>LHKLEDENIEHMPIGFEVAFPSVLEIARSL<br>KLDVPDNSPVLHKIYACRNLKLTKIPKDIL<br>HNVPTTLLHSMEGMAGLDWEKLLKLQCQ<br>DGSFLFSPASTAYALQQTKDQKCMSYLS<br>RAVRKFNGGVPNVYPVDLFEHMWAVDR<br>LQRLGLYRYFEPEIKECMNYVAKFWTEK<br>GICWARNSEVQDIDDTAMGFRLLRLHGH<br>KVSSDVFKHFKKGDEFFCFAGQSNQAVT<br>GMYNLYRASQVALPGEKILYEAKEFSTKF<br>LREKQASNELLDKWIIMKDLPGEIGYALD<br>VPWYASLPRLETKVYIEQYGGGDDVWIG<br>KSLYSMPYVNNNVYLELAKLDFNNCQAL<br>HLSEWDNLQKWYGEWKLGGYGLSKKSL |

| | | | | | | | |
|---|---|---|---|---|---|---|---|
| | | | | | | | LMAYFVAAASIFEPERANERLAWAKTASL<br>VDTLESYFKGETSHKDKKTFVDEFKNLSN<br>MREYEIARRSNKIAGQGILGALLATLSQLS<br>LGTMVLHGQDITQSLRETWVKWLMKWL<br>ERGGMYQDEAELLVETINQTAGLSPTQG<br>LFSNNPEYEHIFSLTNKVCNQLRSCQKQK<br>HKVNDDGSYKSKMNMTTQAIESDMHQL<br>VELVLKSPDGIESLLKQSFFTVARSFYYF<br>NYCDPEIISHHIDKVLFQRAI* |
| Fxa2Dg202041.1 | diTPS | *FaCPS4* | *F. x ananassa* | c | - | FaRR1 genomic | MASHSTHLLFLSLSSSLSPASSSFFDHNQ<br>PKHVPGLSLFGAKDKRVLPSICNAVSKPH<br>TQDYPDVIKWPENPTIVGDYTQQDEEAL<br>PTKDDDSATTEIIREHVETVKSMLSSMEN<br>GEISVSAYDTAWVALVEDVNGTGSPQFP<br>SSLEWIANNQLEDDSWGYEDIYNAHDRII<br>STLACVVALKTWNLHPDKCEKGIKFFKEN<br>LHKLEDENIEHMPIGFEVAFPSVLEIARSL<br>KLDVPDNSPVLHKIYACRNLKLTKIPKDIL<br>HNVPTTLLHSMEGMADLDWEKLLKLQCQ<br>DGSFLFSPASTAYALQQTKDQKCMSYLS<br>RAVRKFNGGVPNVYPVDLFEHMWAVDR<br>LQRLGLYRYFEPEIKECMNYVAKFWTEK<br>GICWARNSEVQDIDDTAMGFRLLRLHGH<br>KVSSDVFKHFKKGDEFFCFAGQSNQAVT<br>GMYNLYRASQVALPGEKTLYEAKVFSTK<br>FLREKQASNELLDKWIIMKDLPGEVGYAL<br>DVPWYASLPRLETKVYIEQYGGGDDVWI<br>GKSLYSMPYVNNNVYLELAKLDFNNCQA<br>LHLSEWDNLQKWYGEWKLGGYGLSKKS<br>LLMAYFVAAASIFEPERANERLAWAKTAS<br>LVDTIESYFKGETSHQDKKTFVDEFKSLS<br>NMREYEIARRSNKIAGQGILGALLATLSQL<br>SLGTMVLHGQDITQSLRQTWVKWLMKW<br>LERDGMYQDEAELLVETINQTAGLSPTQ<br>GLFSNNPEYEHIFSLTNKVCNQLRSCQK<br>QKHKVNDDGSYKSKMNMTTQAFESDMQ<br>QLVELVLKSPDGIESLLKQSFFTVARSFY<br>YFNYCDPEIISHHIDKVLFERAI* |
| **Fxa3Ag101306** | diTPS | ***FaKSL1*** | *F. x ananassa* | **e/f** | **100** | **FaRR1 transcript** | MSFSQLNTFRCRTFSSSSGKDLLPQAAS<br>SIPRAIGTAEVNTAVLSLEGTKQRIKKLFN<br>KVDLSVSSYDTAWVAMVPSPNSGKEPFF<br>PECVNWLLDNQLHDGSWGPPDLHPLLT<br>KNALLSTLACILALKRWNIGEKQIDNGLHF<br>IALNLASATDGEQPSPVGFDIIFPSMIESA<br>FNLDVNLPVGGSTLDALLLRRDVELQRG |

| | | | | | | | |
|---|---|---|---|---|---|---|---|
| | | | | | | | NGSNSEGWRAYLAYISEGIGKSQDWEM<br>VMKYQKKNGSLFNSPSTTAAAFTHLNNA<br>GCLSYLRSLIEKFGSAVPTIYPLDKYARLS<br>MVSSLESMGVDRHFREEIRSVLDETYRC<br>WLQGDEDILSDAGTCAMAFRLLRLHGYD<br>VSADPLSQFSEDRFFNSLGGYLKDTGAA<br>LELFRASEVIIHPDESVLEKQHYWTTHFLK<br>QELSNNLTQAYRLNKHIDQVDDALRLPSY<br>ATLGRLSSRKAIKYYNTDSTKMLKSSYRC<br>TNVGNEDFLKLAVEDFNMCQSIHREELE<br>CLSRWVVESRLDKLNFARQKQAYCYFSA<br>AATLFPPELSDARISWAKNGVLTTVVDDF<br>FDVGGSEVELVNLVQLIEKWDVNERTDS<br>CSEQVEIIFSAIKSTVNEIGANAYPRQERS<br>VTNHFIEIWLDLVKSMLKEAQWLINNSAPI<br>LDEYMENAYVSFALGPIVLPALYFVGPKL<br>SEEAVRSSEFHQLYKLMSTCGRLLNDMQ<br>SFKRESAEGKLNSVTLAMLYGDGSVTEE<br>ETFTEMKSIITSKRRELQRLVLQKDSLVPR<br>ACKDLFWNMCKIVHLFYAKHDGFTGHGL<br>MKTVNGVTEEPIILSELQAESK* |
| **Fxa3Bg201184** | diTPS | ***FaKSL2*** | *F. x ananassa* | **e/f** | **100** | **FaRR1 transcript** | MSFSQLNSFRCRTFSSSSGKDLLPQAAL<br>SIPRAIGTAEVNTAVLSLEGTKQRIKKLFN<br>KVDLSVSSYDTAWVAMVPSPNSGKEPFF<br>PECVNWLLDNQLHDGSWGPPDLHPLLT<br>KDALLSTLACIIALKRWNIGEKQIDNGLHFI<br>ALNLASATDGEQPSPVGFDIIFSSMIESAF<br>NLDVNLPVGGSTLDALLLRRDVELQRGN<br>ESNSEGWRAYLAYISEGIGKSQDWEMV<br>MKYQKKNGSLFNSPSTTAAAFTHLNNAG<br>CLSYLRSLIEKFGSAVPTIYPLDKYARLSM<br>VSSLESMGVDRHFREEIKSVLDETYRCW<br>LQGDEDILSDAGTCAMAFRLLRLHGYDV<br>SADSLSQFSEDRFFNSLGGYLKDTGAAL<br>ELFRASEVIIHPDESVLEKQHYWTSHFLK<br>QELSNNLTQAHRLNKHIDQVDDALHLPSY<br>ATLGRISSRKAIKYYNTDSTKMLKSSYRC<br>SNVGNEDFLKLAVEDFNMCQSVHREELE<br>RLSRWVVESRLDKLNFARQKQAYCYFSA<br>AATLFPPELSDARISWAKNGVLTTVVDDF<br>FDVGGSEVELVNLVQLIEKWDVNARTDS<br>CSEQVEIIFSAIKSTVNETGANAYPRQER<br>SVTNHFIEIWLDLVKSMLKEAQWLINNSA<br>PTLDEYMENAYVSFALGPIVLPALYFVGP<br>KLSEEAVRSSEFHQLYKLMSTCGRLLND |

| | | | | | | | |
|---|---|---|---|---|---|---|---|
| | | | | | | | MQSFKRESAEGKLNSVTLAMLHGDGSVT EGETFTEMKSIITSKRRELQRLVLQKDSL VPRACKDLFWNMCKIVHLFYAKHDGFTG HDLMKTVNGVTEEPIILSELQAESK* |
| **Fxa3Cg101173** | diTPS | ***FaKSL3*** | *F. x ananassa* | **e/f** | **100** | **FaRR1 transcript** | MSFSQLNTFRCRTISSSSGKDLLPQAASS IPRAIGTAEINTAILSFEGTKQRIKKLFNKV DLSVSSYDTAWVAMVPSPNSGKEPFFPE CVNWLLDNQLHDGSWGPPDLHPLLTKD ALLSTLACILALKRWNIGEKQIDNGLHFIAL NLASATDGEQPSPVGFDIIFPSMIEYAFDL DVNLPVGGSTLDALLLRRDVELQRGNGS NSEGWRAYLAYISEGIGKSQDWEMVMK YQKKNGSLFNSPSTTAAAFTHLNNAGCL SYLRSLIEKFGSAVPTIYPLDKYARLSMVS SLESMGVDRHFREELRSVLDETYRCWLQ GDEDILSDAGTCAMAFRLLRLHGYDVSA DPLSQFSEDRFFNSLGGYLKDTGSALELF RASEVIVHPDESVLEKQHYWTSHFLKQE LSNNLTQAHRLNKHIDQIDDALRLPSYATL GRLSSRKAIKYYNTDSTKMLKSSYCCTNV GNEDFLKLAVEDFNMCQSIHREELECLS RWVVESRLDKLNFARQKQAYCYFSAAAT LFPPELSDARISWPKNGVLTTVVDDFFDV GGSEVELVNLVQLIEKWDVNERTDSCSE QVEIIFSAIKSTVNEIGANAYPRQERSVTN HFIEIWLDLVKSMLKEAQWLINNSAPILDE YMENAYVSFALGPIVLPALYFVGPKLSEE AVRSSEFHQLYKLMSTCGRLLNDMQSFK RESAEGKLNSVTLAMLYGDGSVTEEETF TEMKSIITSKRRELQRLVLQKDSLVPRAC KDLFWTCAKLCTYFMQSTMDLLGMT* |
| **Fxa3Dg2011124** | diTPS | ***FaKSL4*** | *F. x ananassa* | **e/f** | **100*** | **FaRR1 transcript** | MSSHLKIFRCCTSSSSGNYELPQVEVNT AGLNFEATKQRIKTMFNKVHLSVSSYDTA WVAMVPSPNSANGPFFPECVNWLLDNQ LHDGSWGWGPTDMHPLLTKDALSSTLA CILALKQWSVGEEQINKGLHFIESNIASAT DEEQPSPVGFDIIFPAMIESAMSLNMNFP QGGSTLKALFHRRDFELKRGHGSNSKG WKAYIAYISEGIGKSQDWKMVMKYQRKN GSLFNSPSATASAFTHSKNAGCIDYLRVLI KKFGNAVPTVYPLDHYARLSMVSTLDSL GIDRHFREEIRTVLDETYRCWLQGNEDV FSDAATCAMAFRLLRVNGYDVSADRLNQ FSDYKFFDSLGGYLRDTDAALELFRASEV IIHPDESVLENQNYWTRNFLKQELSNNLT |

| | | | | | | | |
|---|---|---|---|---|---|---|---|
| | | | | | | | QPHSINKYIGLEVDDALKFPSYATVGRLS<br>SRRAITSYNTNSTRILKSSYRCLNIGNEDF<br>LKLAVEDFNMCQSIHREELEYLSRWVTE<br>SRFDKLKFARQNLANCYFSAAGTLFPPEL<br>SDARISWTKNAVLVTVVDDFFDIEVSEEE<br>LINLIQLVEKWDVNPRTESCSENVEIIFSA<br>MKSLVSETGINAFARQERSVTNHVIENWL<br>DLLKAMLKEAKWLINKTAPTMEEYMENA<br>YVSFGMGPVVLSNLYLVGPKLSEDAVRS<br>SEYYHLNRLLSTVGRLLNDVQGFKKEAA<br>EGKLTAVTLAMIHGNGSVTEEEVINSMKR<br>IITSKRRELRRLVSQDKDSEVPRACKDVF<br>WNMSRFSHLFYERHDGFTRHDVMKTVN<br>ELIKEPIILSELQSRKWVVESRLDKLKFAR<br>QNLALCYFPAAGIIFTPELSDARRSCTKN<br>AVLATVVDDLFDVGGSEEWDVNPTANSC<br>SENVEIIFSAIKSIVGETGINAFARQDRSGT<br>NHVIEIWLDLLKAMLKEAEWVINKSVPTM<br>EDYMDNAYVSFGMGPIILSTLSGPKLSED<br>AVQSLEYYNLYRLMGTVGRLLNDLVTYK<br>KEAAVGKLNAATLAVIHGNGSVSEEEDK<br>DSAVPRACKELFWNMGKSLNLFYGKQD<br>GHTSHDMMKYANEANEIPIILNEFQSRKK<br>RLLPQAASSIPRAIGTAEINTANLSFDGTK<br>QRIKKLFNKVDLSVSSYDTAWVAMVPSP<br>NSGREPFFPECVNWLLDNQLHDGAWGT<br>PDLHPLLTKDALLSTLACILALKRWNIGEK<br>QIDNGLHFIALNLASATDGEQPSPVGFDII<br>FPSMIESAFDLDVNLPVGGSTLDALLLRR<br>DVELQRGNESNSEGWRAYLAYISEGIGK<br>SQDWEMVMKYQKKNGSLFNSPSTTAAA<br>FTHLNNAGCLSYLCSLIEKFGSAVPMIYHL<br>DKYARLSMVSSLESMGVDRHFREEIGSV<br>LDETYRCWLQGDEDILSDAGTCAMAFRL<br>LRLHGYDVSADPLSRFSEDRFFSSLGGY<br>LKDTGSALELFRASEVIIHPDESVLEKQHY<br>WTSHFLKQELSNNLTQAHRLNNHIDQVD<br>DALRLPSYATLGRLSSRKAIKYYNTDSTKI<br>CSNVGNEDFLKLAVEDFNMCQSIHREEL<br>KCLSRWVVESRLDKLNFARQKQAYCYFS<br>AAATLFPPELSDARISWAKNGVLTTVVDD<br>FFDVGGSEVELVNLVQLIEKWDVNARTD<br>SCSEQVEITFSAIKSTVNEIGANAYPRQE<br>RSVTNHFIEIWSDLVKSMLKEAQWLINNS<br>APTLDEYMENAYVSFALGPIVLPAPYFVG |

| | | | | | | | |
|---|---|---|---|---|---|---|---|
| | | | | | | | PKLSEEAVRSSEFHQLYKLMSTCGRLLN DMQSFKRESAEGKLNSVTLAMLHGDGS VTEEETFTEMKSILTSKRRELQRLVLQKD SLVPRACKDLFWNMCKIVHLFYAKHDGF TGHDLMKTVNGVTEEPIILSELQAESK* |
| **Fxa3Dg201123.1** | diTPS | ***FvrCPS5*** | *F. virginiana* | **c** | **98.5** | **NC transcript** | MHTSSHISHRLVLSFPTSLPSSSSLASFS THNQPKHLLAGFLGNEDKRVLPLICSAFS KPHPRDCAPLLHKIVGHDTRQNDASAST QNIKEHVEMVKFMLSSMEEGDISISAYDT AWVAVVEDVNGTGSPQFPSSLEWIANN QLEDGSWGYKDLFSAHDRLISTLACVVAL KSWNIHAEKCFKGMKFFKENLHKIEDENK EHILVGFDVAFPSLLKIARSLNLEVPDDNT LVLREIHARRNLKLSKIPWDILHNVATSLL YSLEGMAGLDWEKLLKLQSEDGSFLFSP ASTAYALKQTNDQDCMSYLSRIVQKFNG GVPHVYPVDLFERMWVVDRLQRLGLSR YLEPEIKECICYVSRYWTEKGISWERNSE VPDLDDTSMGFRLLRMHGHKVSAEVFEH FKKGSQFFCYPGQLNQSVTVMHNLYRAS QVAFPGEKILGDAKEFATKFLREKQVSNE FSDKWIITKDLPGEVVYALDFPWYASLPR LETRFYIEQYGGEDDVWIAKTLYSMPNIS NNVFLELARLDYNDCQALHLKEWDNIQK WYKEWKLGDYGLSEKSLLTTYFVATASIF EPETANVRLAWAKTACLVDTIVCSCISDG DKKAFVDEFKFFSNMQDHGTARRLYKNK IAGQGIVGALLATVNQLSSVTMVLHRQNII ESLCQAWTKWLLKWQETGETYKDEAELL VETINQTAGLSQELMSNNSEYEQLFGLTN KICNQLRSYRDQKNKVNHNGNCTPEIES YMQQLVELVLEKAPDDTGSLIKQSFFTVT RSFYYSTYCDSQVISHHINKVLFQQAI* |
| **Fxa3Dg201123.1** | diTPS | ***FaCPS5*** | *F. x ananassa* | **c** | **99.5** | **FaRR1 transcript** | MHTSSHISHRLVLSFPTSLPSSSSLASFS THNQPKHLLAGFLGNEDKRVLPLICSAFS KPHPRDCAPLLHKIVGHDTRQNDASAST QNIKEHVEKVKSMLSSMEDGDISISAYDT AWVAVVEDVNGTGSPQFPSSLEWIANN QLEDGSWGYKDLFSAHDRLISTLACVVAL KSWNIHAEKCFKGMKFFKKNLHKIEDENK EHILVGFDVAFPSLLKIARSLNLEVPDDNT LVLREIHARKNLKFSKIPWDILHNVATSLL YSLEGMAGLDWEKLLKLQSEDGSFLFSP ASTAYALKQTNDQDCMSYLSRIVQKFNG GVPHVYPVDLFERMWVVDRLQRLGLSR |

| | | | | | | | |
|---|---|---|---|---|---|---|---|
| | | | | | | | YLEPEIKECICYVSRYWTEKGISWERNSE VPDLDDTSMGFRLLRMHGHKVSAEVFEH FKKGSEFFCYPGQLNQSVTVMHNLYRAS QVALPGEKILGDAKEFATKFLREKQVSNE FSDKWIITKDLPGEVVYALDFPWYASLPR LETRFYIEQYGGEDDVWIAKTLYSMPNIS NNVFLELARLDYNDCQALHLKEWDNIQK WYKEWKLGDYGLSEKSLLTTYFVATASIF EPETANERLAWAKTACLVDTIVSSCISDG DKKDFVDEFKFFSNMQDHGTARRLYKNK IAGQGIVGALLATVNQLSSVTMVLHRQNII ESLCQAWTKWLLKWQEAGETYKDEAEL LVETINQTAGLSQELMSNNSEYEQLFGLT NKICNQLRSYRDQKNKVNHNGNCTPEIE SYMQQLVELVLEKAPDDTGSLIKQSFFTV TRSFYYSTYCDSQVISHHINKVLFQQAI* |
| **Fxa3Dg201124** | diTPS | ***FcKSL5*** | *F. chiloensis* | **e/f** | **81.6** | **Amb transcript** | MVMKYQRKNGSLFNSPSTTASAFTHTKN AGCLDYLHALIEKLGNAVPTVYPLDHYAR LSMVSTLDSLGIDRHFREEIRSVLDKTYR CWLQGDEDVFSDAATCAMAFRLLRVNG YDVSADWLNRFSEDNFFDSLGGYLRDTD AALELFRASQVIIHPDESVLEKQNCWTSN FLKQEFSHNKTQPHRVNKLIGLEVDDALK FPSYATFSRLSSRRVILSYNTNSTRILKSS YRCLNIGNEDFLKLAAEDYNMCQSVYQE ELEHLSRWVVESRLDKLKFARQNLAFCY FTAAGILFTPELSDARRSCTKNAVLVTVV DDLFDVGGSEEELINLIQLVEKWDVTPRA DSCSENVEIIFSAIKSTVGETGINAFARQA RGVTNHVIEMWLDLLKAMLKEAEWVINKL APTMEDYMDNAYVSFGMGPIILSTLYLVG PKLSEDAVQSLEYYNLYRLMSTIGRLLND LVTYKKEAAVGKLNAVTLAVIHGNGSVTK EEAINETKSIITSKRRELQRLVLQDKDSAV PRACKELFWNMGKALNLFYGKQDGHTS HDMMKYANEVNEIPIILNELQSRK* |
| Fxa1Ag100480 | M/S TPS | *FaTPS1* | *F. x ananassa* | a | - | FaRR1 Genomic | MPVQATPAAECQIISKPEVVRRTANFKPS VWGDRFTNYAEDIITQAQMQEQVEELKQ VVRKEVFTDAAADSSRQLKLIAAIQRLGV AYHFETEIEQALERIHATYQDIHDDDGDL YNVALRFRLLRRHGYNVSCDVFNKFIDTN GDFKKSLVTDVSGMLSFYEAAHLRVHGE TLLEEALVFTTTHLESASAISSLLKTPITEA LERPLLKTMERLGARRYMSIYQDEASHS ENLLKLAKLDFNVVQCLHKKELSDILRWY |

| | | | | | | | |
|---|---|---|---|---|---|---|---|
| | | | | | | | KELDFARRMPFARDRVVELFFWIAGIYFE<br>PQYVFGRHILTKLIEIVTVMDDMYDAFGTF<br>EELVILTEAIDRWDASCMDQLPDYMQPF<br>YITLLDVIDEVEEELTKQGRSYRIHYAKDIL<br>KNQARLYFAEARWFHEGCTPKMDEYMR<br>VAASSVGNTMLSVVSLVGMGDIITKEEFE<br>WLTNEPKILRASNTIFRLMDDIAGYKFEKE<br>RGHVASSIDCYMNEYGVSEQETIDIFNKR<br>IVDSWKDINEEFLRPTAAPVPVLNRVLNL<br>TRVVDLLYKRGDAFTHVGKLMKDCIAAM<br>FIDPVPL* |
| Fxa1Bg200351 | M/S TPS | *FaTPS2* | *F. x ananassa* | a | - | FaRR1 Genomic | MSLTSVILASQSPQTLNNADTRRSANYHP<br>SIWGDWFLSYNNTMETNMQGKQQVLQQ<br>LKEEVKRMLMAPPVETGLGKLELIDDIQR<br>LGVSYHFEYEIDQTMQQIHENLNGTYDD<br>DDLHTCALRFRLLRQHGYNVSCDMFNKF<br>KDCDGKFDESLHHDVAGLQSLYEATHLR<br>VRGEDFLEEALAFTTTHLQSAANRLSPPL<br>SKQVRHALNQPLRKGLQRLEARHYMSLH<br>QELHGSRNHVLLRFAKLDFNLLQQVHQK<br>ELSDIARWWKKLDFASKLPFARDRVIECY<br>FWILGVYFEPKYYFARRTLTKVIAMTSIID<br>DIYDVYGTLEELDLFTRAIERWDISAMDVL<br>PEYMKFCYEALLDVYAEAEQGLASEGKS<br>YRIDYAKEAMKRQVRAYHAEAKWFHNNY<br>TPTMDEYMEVALVTSAYSMLATTSFVGM<br>GDIVTKDSFEWIFSDPKMVKASAVVCRL<br>MDDIVSHKFEQKRGHVASAVECYMTQY<br>GATEEETIIEFRKQVSDAWKDINEECLHP<br>TSVDMPLLMRVLNLTRVIDVLYKSEDGYT<br>HAGTILKDFVASLLVDPVTV* |
| Fxa1Dg200965 | M/S TPS | *FaTPS3* | *F. x ananassa* | a | - | FaRR1 Genomic | MSSQSAAAPPSQTIMSEIAGLTAIYQPGIC<br>GDGFNNYESLDNITRAHIEKQINQLKVLV<br>RELFTSTASGLPHQLKLIDDIQRLGVSYHF<br>ETELTEALENIHVQLHDGDLDLYNDSLRF<br>RLLRQHGYNISSGDIFKKFKDANGNFKES<br>LITDIPGMLSLYEATHLRVHGEEILEEALV<br>FTTNHLELAKSQVSYPLKAQISEALERPM<br>WRSLERLSARNYIPIYQATTSHNEALLKL<br>AKLDFNLVQDLHKEELSEVSRWWKELDF<br>ERKMPFARHRIVEVFFWMVGLYYEPKYS<br>VGRKIGAKLCAMATMLDDIFDSYGTFEKL<br>KIFRQVIERWDVNCCTDDLPQYMKVYYH<br>SLCEIMNEIEEELEKLGMPDRVHYAKQVL<br>KDQARDYLVEAEWLHKGCTPSMEEYLPV |

| | | | | | | | |
|---|---|---|---|---|---|---|---|
| | | | | | | | RVQSVGVCMTVVFSLLGMNDNITKETFE WVLKYPKIVMASSFIFRFMDDIVGSKHEK QKGDVDSSIDCYMKQYGVSKEEAIEVLD KKIVDSWKEINEDLLRPTAVPMYVLLRVM NFSRGVDLVYKEEDGFTLVGKVVKHAVA ALFGDPLPLE* |
| Fxa2Bg200794 | M/S TPS | *FaTPS4* | *F. x ananassa* | a | - | FaRR1 Genomic | MSMIVEPSRSSPGPQIAMPEIVRPIANFQ PSIWGDQFLNYDSQDIKTEALWQQQVEE LKVKVKSEVFTNEGDFALRLKLIDAIQRLG VAYQFEEEIEDALRHIHATYRDQDQFNDD SDLPTVALCFRLLRQHGYEISSDIFNKFKD ENGSFKEGLVADGCAMLSLYEAAHLRVH GENILEEALVFTTTHLESAKSSACYSAQM TEQITQALARPLRTSLERICAKRHMSVYQ LVDDAALVKHHETLLKLAKMDFNLVQFLH KKELCEITRWWKELDFERKLPFARDRKV ELFFWIVGVYFEPQYSTGRIFMTKVAILLT VMDDIYDAYGTFEELVVFTEAIDRWDVKC IDELPDYLKIFYYELLNVFHEMDKVLAKEG RSYRVCYAIQAVPYELHVRLLDQMKDQA RSYFNEARWLHEGRTPSMEEYMSVATV SISYTFLTTISLLGMGDIVTKDSFEWLSNG PKIVRASNIIFRLMDDTVSTKFEKERGHAP SSIDCYTKQYGVSEQEAIDVFNKQIVESW KDINEEFLKPTAVPMPVLMRVLNLTRVAD LLYKGEDGFTRVGKMTKDSVAAVIIDSVP L* |
| Fxa2Bg200800 | M/S TPS | *FaTPS5* | *F. x ananassa* | a | - | FaRR1 Genomic | MSMIIEPSGSSPVPQIAMPEIVRPIANFQP SIWGDQFLNYDSQDIKTEALWQQQVEEL KVKVKSEMFTNEGDFALRLKLIDAIQRLG VAYQFEEEIEDALRHIHATYRDQDQFNDD SDLPTVALCFRLLRQHGYEISSDIFNKFKD ENGSFKEGLVADGCAMLSLYEAAHLRVH GENILEEALVFTTTHLESAKSSACYSAQM TEQITQALARPLRTSLERICAKRHMSVYQ LEDDAALVKHHETLLKLAKMDFNLVQFLH KKELCEITRWWKELDFERKLPFARDRKV ELFFWIVGVYFEPQYSTGRIFMTKVAILLT VMDDIYDAYGTFEELVVFTEAIDRWDVKC IDELPDYLKIFYYELLNVFHEMDKVLAKEG RSYRVCYAIQAMKDQARSYFNEARWLHE GRTPSMEEYMSVATVSISYTFLTTISLLG MGDIVTKDSFEWLSNGPKIVRASNIIFRLM DDIVSTKFEKERGHAPSSIDCYTKQYGVS EQEAIDVFNKQIVESWKDINEEFLKPTAV |

| | | | | | | | |
|---|---|---|---|---|---|---|---|
| | | | | | | | PMPVLMRVLNLTRVADLLYKGEDGFTRV GKMTKDSVAAVIIDSVPLCNPSERIRSNH FPCMLKLVLGSHMCNVAQVMRKDQTLLH NHVQSS* |
| **Fxa2Bg200800.1** | M/S TPS | ***FvrTPS5*** | *F. virginiana* | **a** | **98.9** | **HS transcript** | MSMIKEPSRSSPVPQIDMPEIVRPIANFQ PSIWGDQFLNYDSQDIKTEALWQQQVEE LKVKVKSEVFTNEGDFALRLKLIDAIQRLG VAYQFEEEIEDALRHIHATYRDQDQFNDD SDLPTVALCFRLLRQHGYEISSDIFNKFKD ENGSFIEGLVADGCAMLSLYEAAHLRVH GENILEEALVFTTTHLESAKSSACYSAQM TEQITQALARPLRTSLERICAKRHMSVYQ LEDDAALVKHHETLLKLAKMDFNLVQFLH KKELCEITRWWKELDFERKLPFARDRKV ELFFWIVGVYFEPQYSTGRIFMTKVAILLT VMDDIYDAYGTFEELVVFTEAIDRWDVKC IDELPDYLKIFYYELLNVFHEMDKVLAKEG RSYRVCYAIQAMKDQARSYFNEARWLHE GRTPSMEEYMSVATVSISYTFLTTISLLG MGDIVTKDSFEWLSNGPKIVRASNIIFRLM DDIVSTKFEKERGHAPSSIDCYTKQYGVS EQEA* |
| Fxa3Ag100266.1 | M/S TPS | *FaTPS6* | *F. x ananassa* | b | - | FaRR1 Genomic | MDCSKQLRNPQAEEQQIHQWQTKSELS ADSVHDQSRRSANYKPNIWKYDFLESLN SKFDGECYLIQMQKLIKDVKKLFVECKES DVIAKLELIDSIRKVGLNNHFEQEIKAALDA IASAELENNSNPCISSEGDLYAAALFFKIL RHQGYQVSQDIFGRFMDEMGALKKSTS GNVKGMIELLEASNLAFEGEDILEKAQAF LIATLRDTNTMWDEIDSSISKHVTCSLELS SQRRVQWFNVKWHIKAYEQNRTNQPYM TTLLELAKLNFNVVQATLQKDLREASKW WYNLGLTKNLDFARDRMVECFMCAVGL AFETDHKSFRKWLTKVINLILIIDDVYDVY GSLEELKHFTRAVERWDVMETEQLPEC MKICFQVLYNTTYEIAYEIEEENGWNQVL PHLCKVWADFCKALLVEAEWYNEAYTPS FEEYLSNGYISSSASLIFTHAFFATKHEEG VDDSLHMNEDLVYNISVILRLLNDLGTSA AEQERGDAASSILCYMREMNVSEEIARK NIKSMIDYAWKKINENCLTRNPKISSSYINI TTNIARVGHILYQDGDGFGDQEQGTRALI QSLLVQPLS* |

| | | | | | | | |
|---|---|---|---|---|---|---|---|
| **Fxa3Ag100266.1** | M/S TPS | ***FvTPS6*** | *F. vesca* | **b** | **97.9** | **UC06 transcript** | MDCSKQLRNPQAEEQQIHQWQTKSELS AYSVHDQSRRSANYKPNIWKYDFLESLN SKFDGECYLIQMQKLIKDVKKLFVECKES DVIAKLELIDSIRKVGLNNHFEQEIKAALDA IASAELENNSNPCISSEGDLYAAALFFKIL RQQGYQVSQDIFGRFMDEMGTLKKSTS GNVKGMIELLEASNLAFEGEDILEKAQAF LIATLRDTNTMWDEIDSSISKHVTCSLELS SQRRVQWFNVKWHIKAYEQNHTNQPYM TTLLELAKLNFNEVQATLQKDLREASKW WYNLGLTKNLDFARDRMVECFMCAVGL AFETDHKSFRKWLTKVINLILIIDDVYDVY GSLEELKHFTRAVERWDVMETEQLPEC MKICFQVLYNTTCEIANETEEENGWNQVL PQLCKVWADFCKALLVEAEWYNEAYTPS FEEYLSNGYISSSASLIFTHAFFATKHEEG VDDSLHMNEDLVYNISVILRLLNDLGTSA AEQERGDAASSILCYKREMNVSEEIARKN IKSMIDNAWKKINENCLTRNPKISSSYINIT TNIARVGHILYQDGDGFGDQEQGTRALIQ SLLVEPLS* |
| **Fxa3Ag100266.1** | M/S TPS | ***FaTPS7*** | *F. x ananassa* | **b** | **94.9** | **Prim transcript** | MDCSKQLRNPQAEEQQIHQWQTKSEFS AHSVHDQSRRSANYKPNIWKYDFLESLN SKFDGGGYLIQMQKLIKDVKKLFVECKES DVIAKLELIDSIRKLGLNNHFEQEIKAALDA IASAELENNRNPCISSEDDDLYAAALFFKI LRQQGYQVSQDIFGRFMDEMGTLKKSSF GNVKGMIELLETSNLAFEGENFLEKAQAF LIATLRDTNTMWDEIDSSISKHVTYALQIS SQRRAQWFNVKWHIKAYEQNHTNQPYM TTLLELAKLNFNVVQATLQKDLREASKW WYNLGLTKNLDFARDRMVECFMCAVGL AFETDHKSFRKWLTKVINLILIIDDVYDVY GSLEELKHFTRAVERWDVMETEQLPDC MKICFQVLYNTTCEIAYEIDEENGWNQVL PHLCKVWADFCKALLVEAEWYNEAYTPS FEEYLSNGYISSSASLIFTHAFFATKHEEG VDDFLHMNEDLLYNISVILRLLNDLGTSAA EQERGDAASSILCYMREMNVSEEIARKNI KSMIDNAWKKINENCLTRNPKISSSYINIT TNIARVGHSLYQDGDGFGDQEQGTRALI QSLLVQPLS* |
| Fxa3Ag100275.1 | M/S TPS | *FaTPS8* | *F. x ananassa* | g | - | FaRR1 Genomic | MNVETKHTRTMDDIFVQHSRKLELLRNVL RNAAEVDALEGLNMIDAVQRLGIDYHFQ REIDAILHKQMSIVSASDDLHEVALRFRLL |

| | | | | | | | |
|---|---|---|---|---|---|---|---|
| | | | | | | | RQHGYSVPEDVFNNFKESKGTFKQVLGE<br>DIKGLMSLYEASQLGTEGEDTLVEAEKFS<br>GHLLKTSLSHLDHHQARIVGNTLRNPHH<br>RSLASFMARNFFVTSQATRLNSWLTLLK<br>EVAKIDFNMVRSRHQNEIVQISKWWKDL<br>GLAKELKFARDQPLKWYIWSMAVLTDPK<br>LSEERVELTKPISFVYLIDDIFDVYGTLDDL<br>YLFTEAVNRWEITAIDHLPDYMKICFKALY<br>DMTNEFSCKVYQKHGWNPLQSLKISWA<br>SLCNAFLVEAKWFASGKLPKSEEYLKNGI<br>VSSGVNVVLVHMFFLLGQNITRKSVELLN<br>ETPAIISSAAAILRLWDDLGSAKDENQDG<br>NDGSYVRCYLEEHEGCSIEEAREKTINMI<br>SDEWKKLNRELLSPNPFPATFTLASLNVA<br>RMIPLMYSYDGNQCLPSLKEYVKLMLYE<br>TVSM* |
| Fxa3Ag102016.1 | M/S TPS | *FaTPS9* | *F. x ananassa* | a | - | FaRR1 Genomic | MPSQSAPAPPSQTTMSEIAGLTAFPPGIC<br>GDGFTTFETLDNVTRAHMEQQIAKLKVLV<br>RELLTSTAAGLSHQLKFIDDIQRLGVSYHF<br>ETELAEALENIHARLHDGDLDLYNESLRF<br>RLLRQHGYNISSGDIFKKFKDANGNFKES<br>LIADVPGMLSLYEATHLRVHGEEILEEALV<br>FTTNHLELAMSQVSYPLKAQISAALERPL<br>RRSLERLSVRNYITIYQATTSHNEALLKLV<br>KLDFNLLQSLHKEELSEVTRWWKEQDYE<br>SKIPFARHRIVECFFWMVGMYYEPQYSA<br>ARKIGSKLCTFATMLDDVYDSYGGTVEEL<br>KIFRAAMDRWDVNCCMDGLPPYMIGFYH<br>SLLDMMNAIEEELEKQGLSYRVQYAKQV<br>LKNQARDYLVEAKWLQEECTPSMEEYM<br>PVRTRSAGSCMTIVLSLLGMNDNIPKETF<br>EWILKYPKIIRAASLIFRLMDDIEGCKSEKE<br>QGDVASSIDCYMKQYRVSEEEAIDVFNK<br>QIVEAWKDINEDLLQPIAKPMYVLKRALN<br>FTRNVDVVYKGEDGFKYVGKVVKQGVA<br>ALFTMSEIAGLTAIYQPGICGDGFNNFESL<br>DNITRAHMEKQINQLKVLVRELLTSTAAG<br>LSHQLKFIDDIQRLGVSYHFETELAEALEN<br>IHSRLHDGDLDLYNDTLRFRLLRQHGYNI<br>SSGDIFKKFKDANGNFKESLIADVPGMLS<br>LYEATHLRVHGEEILEEALVFTTNHLELAI<br>SQVSCPLKAQISEALERPMWRSLERLSA<br>RNYIPIYQATTSHNEALLKHAKLDFNLVQA<br>LYKEELSEVSRWWKELDFERKMPFARH<br>RIVECFFWMVGMYYEPKYSVGRKIGTKL |

| | | | | | | | |
|---|---|---|---|---|---|---|---|
| | | | | | | | CAMATMLDDIFDSYGTFEELKIFRQVIERWDVNCCMDDLPRYMKVYYHSLCDVMNEIEEELEKLGMPDRIHYAKQVLKDQARDYLVEAEWLHEGCTLSMEEYLPVRVQSVGVCMTVVFSLLGMNDNITKETFEWVLKYPKIVMASSFIFRLMDDIGGSKHEKQKGDVDSSIDCYMKQYGVSKEEAIEVLDKQIVDSWKEINEDLLRPTAVPMYVLLRVMNFSRDVDLVYKGEDGFTHVGKVVKHAVAALFGDPLPLE* |
| **Fxa3Bg200271.1** | M/S TPS | ***FaTPS10*** | *F. x ananassa* | **b** | **99.5** | **EM transcript** | MDCSKQLRNPQAEEQQIHQSRRSANYKPNIWKYDFLESLNSKFDGECYLIQMQKLIKDVKELFVECKESDVIAKLELIDSIRKLGLNNHFEQEIKATLDTKASVGLENNSNPCISSEGDLYAAALLFKILRQHGYQVSQDIFGRFMDEMGTLKKSSFGNVKGMIELLEASNLAFEGEDILEKAQAFLIASLRDTNTMWDEIDSSISKHVTYALELSSQRRVQWFNVKWHIKACEQNRTNQRYMTTLLELAKLNFNVVQATLQKDLREASKWWYNLGLTKNLDFARDRMVECFMCAVGLAFETDHKSFRTWLTKVINLILIVDDVYDVYGSLEELKHFTRAVERWDLMETEQLPECMKICFQVLYNTTCEIAYEIDEENGWNQVLPHLCKVWADFCKALLVEAEWYNEAYTPSFEEYLSNGYISSSASLIFTHAFFATKHEEGVEDFLHMNEDLVYNISVILRLLNDLGTSAAEQERGDAASSILCYMREMNVSEEIARKNIKSMIDNAWKKINENCFTRNPKISSSYINITTNIARVGHILYQDGDGFGDQEQGTRALIQSLLVEPLS* |
| Fxa3Bg200279.1 | M/S TPS | *FaTPS11* | *F. x ananassa* | g | - | FaRR1 Genomic | MLIIDQMASSSRAFFKVFNPAPKSIPHIGQSNLMQLTHKKQLPTFQRRGIAEDSLLPSSTTPIKPMNVETKHTRTMGDIFVQHSQKLELFRNVLRNVAELDALEGLNMIDAVHRLGIDFHFQREIDEILHKQMSIVSASDDLHEVALRFRLLRQHGYFVPEDVFNNFKDSKGTFKQVLGEDIKGLMSLYEASQLGTEGEDTLVEAEKFSGHLLKTSLSHLDHHQARIVGNTLRNPHHKSLASFMARNFFVTSQATNSWLNLLKDVAKTDFNMVRSQHQNEVVQISKWWKELGLAKELKFARDQPQKWYIWSMACLTDPKLSEERVELTKPISFVYLIDDIFDVYGTLDDLILFTEAVNRWEITAIDHVPDYMKICFKALYDMTNEISCKVYQKHGWNPLQSLKNSWASLCNAFLVEAKWFASGQLPKSEEYLK |

| | | | | | | | |
|---|---|---|---|---|---|---|---|
| | | | | | | | NGIVSSGVNVVLVHMFFILGQNITRKSVELLNETPAMISSSAAILRLWDDLGSAKDENQDGNDGSYVRCYLEEHEGCSIEEAREKTINMISDEWKKLNRELLSPNPFPATFTLASLNLARMIPLMYSYDGNQCLPSLKEYMKLMLYETVSM* |
| Fxa3Bg200281.1 | M/S TPS | *FaTPS12* | *F. x ananassa* | g | - | FaRR1 Genomic | MASSSWAFFKVFNPQIAPKSISHIGQSDLMQLTHKKQLPTFQRRGIAEDSLLPSSTTPIKPMNVETKHTRTMGDIFVQHSQKLELFRNVLRNVAELDALEGLNMIDAVQRLGIDYHFQREIDEILHKQMGIVSACDDLYDVALRFRLLRQHGYFVPEDVFNNFKDSKGTFKQVLGEDIKGLMSLYEASQLGTEGEDTLVEAEKFSGHLLKTSLSHLDHHRARIVGNTLRNPHHKSLASFMARNFFVTSQATNSWLNLLKEVAKTDFNMVRSLHQKEIVQISKWWKELGLVKELKFARDQPLKWYTWSMAGLTDPKLSEERVELTKPISFVYLIDDIFDVYGTLDDLILFTEAVNRWEITAIDHLPDYMKICFKALYDMTNEFSCKVYQKHGWNPLRSLKISWASLCNAFLVEAKWFASGQLPKSEEYLKNGIVSSGVNVGLVHMFFLLGQNITRKSVELLNETPAMISSSAAILRLWDDLGSAKDENQDGNDGSYVRCYLEEHEGCSIEEAREKTINMISDEWKKLNRELLTPSPFPATFTLASLNLARMIPLMYSYDGNQSLPSLKEYMKLMLYETVSM* |
| Fxa3Bg200284.1 | M/S TPS | *FaTPS13* | *F. x ananassa* | g | - | FaRR1 Genomic | MHVETKHTRTMGDFFVQHSQKLELLRNVFRNVAELDALEGLNMIDAVQRLGIDYHFQQEIDEILHNQMSIVSACDDLHEVALRFRLLRQHGYFVPEDVFNNFKNSKGKFKQVLGEDIKGLMSLYEASQLGTVLAQRPQRGCDEGKKQGSGNEAPGSVGTEGEDTLVEAEKFSGHLLKTSLSHLDHHRARIVGNTLRNPHHKSLASFMARNFFVTSQATNSWLNLLKEVAKTDFNMVRSLHQNEIVQISKWWKELGLAKELKLARDQPLKWYIWSMACLTDPKLSEERVELTKPISFVYLIDDIFDVYGTLDDLILFTEAVNRWEITAIDHLPDNMKICFRALYDITNEISCKVYLKHGWNPLQSLKISWASLCNAFLVEAKWFASGQLPKSEEYSKNGIVSSGVNVGLVHLFFLLGQNITRKSVELLNETPAIISSSAAILRLWNDLGSAKDENQDGNDGSYVRCYLEEHESCSIEEHEKRRLI* |

| | | | | | | | |
|---|---|---|---|---|---|---|---|
| Fxa3Bg200286.1 | M/S TPS | *FaTPS14* | *F. x ananassa* | g | - | FaRR1 Genomic | MNVETKHTRTMDDIFVQHSRKLELLRNVLRNAAEVDALEGLNMIDAVQRLGIDYHFQREIDAILHKQMSIVSASDDLHEVALRFRLLRQHGYSVPEDVFNNFKESKGTFKQVLGEDIKGLMSLYEASQLGTEGEDTLVEAEKFSGHLLKTSLSHLDHHQARIVGNTLRNPHHRSLASFMARNFFVTSQATRPNSWLTLLKEVAKIDFNMVRSRHQNEIVQISKWWKDLGLAKELKFARDQPLKWYIWSMAVLTDPKLSEERVELTKPISFVYLIDDIFDVYGTLDDLCLFTEAVNRWEITAIDHLPDYMKICFKALYDMTNEFSCKVYQKHGWNPLQSLKISWASLCNAFLVEAKWFASGKLPKSEEYLKNGIVSSGVNVVLVHMFFLLGQNITRKSVELLNETPAIISSSAAILRLWDDLGSAKDENQDGNDGSYVRCYLEEHEGCSIEEAREKTINMISDEWKKLNRELLSPNPFPATFTLASLNLARMIPLMYSYDGNQCLPSLKEYVKLMLYETVSM* |
| Fxa3Bg201829.1 | M/S TPS | *FaTPS15* | *F. x ananassa* | a | - | FaRR1 Genomic | MSSQVSASEAQTPKDVTAEENDTKTDQQVQELKEEVRTMLMSPVKKISQKLELIDDITRLGVSRHFKEVIDEILRKVYDHANDLRVSSEIFSKFKNGEGLFEESLVNDVVGLLSLYEATHLMRHGEDILEEALTFATTHLESAVHSLSSLLLKQVKHALYQPYWKGLPRIEAVHYLSIYGEIDSHNKALLSLAKWWKDLDFVNKLPFARDRIVEAYFWALAVYFEPEYHLARMMCCKFVTIITIVDDIYDVYGTYEELKLFTEAVERWNISAADQLPGKLAMKGNLYRIRYGREAFKVQVRAYIQEAKWFKQKYTPTMDEYMEVELNTSFFAFSTTAFLGLGAIVTKDSMDRVFDDPKIVKVTSVVDRLLNDIVGHKFEQKREHIASSVECYMNQYGVTEEKAKTELMKQVNDAWKDINEEWLQANTSIPKTLRWFILNLARSGEVLYKNEDLFTHSETVLKSFLVSLFVEPVPVFVTHQCKT* |
| Fxa3Cg100132.1 | M/S TPS | *FaTPS16* | *F. x ananassa* | a | - | FaRR1 Genomic | MSFQLPSAASQAPTPNAAAHVMRRSVNFKPSIWGEHFLSYDSMEVDIILEQQVQQLKEEVKRMLMDHVKDDSQKLSLIDDIQRLGVSYHFENEICEILQQIFDKDHVSYDDVLHQNDSDLYAVALGFRLLRQQGHNTSSDKYFSNLKDNDGKFKESLVNDIKGLLSLYEATHLRIHGDDILDEALTFTTTHLESATRRGLSSPLSKQVTHALYQPLWKGMTRIEVSHYLAIYQEDKSHNATLLNLARLDFNLLQQLHKKE |

| | | | | | | | |
|---|---|---|---|---|---|---|---|
| | | | | | | | LCELTRWWKDVDFVKNLPFARDRVVECY<br>LSTLGVFFEPKYYIARRTLCKFIYMVAVID<br>DMYDVYGTLEELELFNEAIQRWDISAMD<br>PLPDYMKVCYKTLLDVYTELEENLASEGS<br>LYGIHHARESMKMHVGGYMKEARWFNS<br>NYTPTSDEYMPLAALTSLYSLTIVAAVGM<br>GVAASKDSLDWLLTDPGIVNAPSVIGRLL<br>NDIMSHQREHSATAVQLYMKEHGATEEA<br>AIVELINQVNDAWKIINKACLNPTRIPLPVL<br>MRPLNLARVMEMLYKNEDAFSNPGDEIK<br>GFVTSVLIEPVLIA* |
| **Fxa3Cg100266.2** | M/S TPS | ***FvrTPS17*** | *F. virginiana* | **g** | **95** | **NC transcript** | MASSSRAFFKVFNPAPKSIPHIGQSNLMQ<br>LTHKKRLPTFQRRGIAEDSLLPSSTTPIKP<br>MNVETKHTRTMGDIFVQHSQKLELLKTVL<br>RNVAELDALEGLNMIDAVQRLGIDFHFQR<br>EIDEILHKQMSIVSASDDLHEVALRFRLLR<br>QHGYFVPEDVFNNFKDSKGTFKQVLGED<br>IKGLMSLYEASQLGTEGEDILVEAEKFSG<br>HLLKTSLSHLDHHQARIVGNTLRNPHHKS<br>LASFMARNFFVTSQATNSWLNLLKDVAK<br>TDFNMVRSLHQNEIVQISKWWKELGLAK<br>ELKFARDQPLKWYIWSMACLTDPKLSEE<br>RVELTKPISFVYLIDDIFDVYGTLDDLILFT<br>EAVNRWEITAIDHLPDYMKICFKALYDMT<br>NEFSSKVYLKHGWNPLQSLKISWASLCN<br>AFLVEAKWFASGQLPKSEEYLKNGIVSS<br>GVNVVLVHMFFILGQNITRKSVELLNETP<br>AMISSSAAILRLWDDLGSAKDENQDGND<br>GSYVR* |
| **Fxa3Cg100266.2** | M/S TPS | ***FaTPS17*** | *F. x ananassa* | **g** | **96.5** | **FaRR1 transcript** | MNVETKHTRTMGDIFVQHSQKLELLKTVL<br>RNVAELDALEGLNMIDAVQRLGIDYNFQR<br>EIDEILHKQMSIVSARDDLHEVALRFRLLR<br>QHGYFVPEDVFNNFKDSKGTFKQVLGED<br>IKGLMSLYEASQLGTEGEDILVEAEKFSG<br>HLLKTSLSHLDHHRVRIVANTLRNPHHKS<br>LAPFMARNFFVTSQATNSWLNLLKEVAK<br>TDFNMVRSLHQNEIVQMSKWWKELGLA<br>KELKFARDQPLKWYIWSMACLTDPKLSE<br>ERVELTKPISFVYLIDDIFDVYGTLDDLILF<br>TEAVNRWEITAIDHLPDYMKICFKALYDM<br>TNEFSSKVYLKHGWNPLQSLKISWASLC<br>NAFLVEAKWFASGKLPKSEEYLKNGIVSS<br>GVNVVLVHMFFLLGQNITRKSVELLNETP<br>AIISSSAAILRLWDDLGSAKDENQDGNDG<br>SYVRCYLEEHEGCSIEEAREKTINMISDE |

| | | | | | | | |
|---|---|---|---|---|---|---|---|
| | | | | | | | WKKLNRELLSPNPFPASFTLASLNLARMI<br>PLMYSYDGNQCLPSLKEYMKLMLYETVS<br>M* |
| **Fxa3Cg100266.3** | M/S TPS | ***FcTPS18*** | *F. chiloensis* | **g** | **96.9** | **AMB transcript** | MALSTRAFFKVFNPQITPNSISHIGQSNL<br>MQLTQKKQLPTFQRRGIAEDSLLPSSTTP<br>IKPMNVETKHTRTMGDIFVQHSQKLELLK<br>TVLRNVAELDALEGLNMIDTVQRLGIDYN<br>FQREIDEILHKQMSIVSARDDLHEVALRF<br>RLLRQHGYFVPEDVFNNFKDSKGTFKQV<br>LGEDIKGLMSLYEASQLGTEGEDILVEAE<br>KFSGHLLKTSLSHLDHHRVRIVANTLRNP<br>HHKSLAPFMARNFFVTSQATNSWLNLLK<br>EVAKTDFNMVRSLHQNEIVQMSKWWKE<br>LGLAKELKFARDQPLKWYIWSMACLTDP<br>KLSEERVELTKPISFVYLIDDIFDVYGTLD<br>DLILFTEAVNRWEITAIDHLPDYMKICFKA<br>LYDMTNEFSSKVYLKHGWNPLQSLKISW<br>ASLCNAFLVEAKWFASGQLPKSEEYLKN<br>GIVSSGVNVVLVHMFFLLGQNITRKSVEL<br>LNETPAIISSSAAILRLWDDLGSAKDENQD<br>GNDGSYVRCYLEEHEGCSIEEAREKTIN<br>MISDEWKKLNRELLSPNPFPASFTLASLN<br>LARMIPLMYSYDGNQCLPSLKEYMKLML<br>YETVSM* |
| **Fxa3Cg100266.3** | M/S TPS | ***FaTPS18*** | *F. x ananassa* | **g** | **100** | **FaRR1 transcript** | MALSTRAFFKVFNPQITPNSISHIGQSNL<br>MQLTQKKQLPTFQRRGIAEDSLLPSSTTP<br>IKPMNVETKHTRTMGDIFVQHSQKLELLK<br>TVLRNVAELDALEGLNMIDAVQRLGIDYN<br>FQREIDEILHKQMSIVSACDDLHEVALRF<br>RLLRQHGYFVPEDVFNNFKDSKGMFKQV<br>LGEDIKGLMSLYEASQLGTEGEDTLVEAE<br>KFSGHLLKTSLSHLDHHRARIVANTLRNP<br>HHKSLAPFMARNFFVTSQATNSWLNLLK<br>EVAKTDFNMVRSLHQNEIVQISKWWKEL<br>GLAKELKFARDQPLKWYIWSMACLTDPK<br>LSEERVELTKPVSFVYLIDDIFDVYGTLDE<br>LILFTEAVNRWEITAIDHLPDYMKICFKALY<br>DMTNEFSSKVYLKHGWNPLQSLKISWAS<br>LCNAFLVEAKWFASGQLPKSEEYLKNGIV<br>SSGVHVGLVHMFFLLGQNITTKSVELLNE<br>TPAMISSSAAILRLWDDLGSAKDENQDG<br>NDGSYVRCYLEEHEGCSIEEAREKTINMI<br>SDEWKKLNRELLSPNPFSATFTLASLNLA<br>RMIPMMYSYDGNRCLPDLKEYMKLMLYE<br>TESM* |

| | | | | | | | |
|---|---|---|---|---|---|---|---|
| Fxa3Cg101786.1 | M/S TPS | *FaTPS19* | *F. x ananassa* | a | - | FaRR1 Genomic | MPSQSAPAPPSQTTMSEIAGLTPFPPGIC<br>GDGFTTFKTLDNVTRAHMELQIAQLKVLV<br>RELLTSTAAGLSHQLKLIDDIQRLGVSYHF<br>ETELAEVLENIHSRLHDGDLDLYNDSLRF<br>RLLRQHGYNISSGDIFKKFKDANGNFKES<br>LTVDVPGMLSLYEATHLRVHGEEILEEAL<br>VFTTNHLELAMSQVSYPLKAQISAALERP<br>LRRSLERLSARNYITIYQATTSHNEALLKL<br>AKLDFNLLQSLHKEELSEVTRWWKEQDY<br>ERKIPFARHRIVECFFWMVGMYYEPQYS<br>AARKIGSKLCTFATMLDDVYDSYGGTVE<br>ELEIFRAAMDRWDVNCCMDGLPPYMIGF<br>YHSLMDMMNAIEEELEKQGLSYRVQYAK<br>QVLKNQARDYLVEAKWLQEECTPSMEE<br>YMPVRSRSAGSCMTVVLSLLGMNDSIPK<br>ETFEWILKYPKIIRAASLIFRLMDDIEGCKS<br>EKEQGDVASSIDCYMKQYRVSEEEAIDV<br>FNKQIVDAWKDINEDLLQPIAKPMYVQKR<br>ALNFTRNVDVVYKGEDGFKYVGKVVKQA<br>HMEKQINQLKVLVRELLTSTAAGLSHQLK<br>LIDDIQRLGVSYHFETELAEALENIHARLH<br>DGDLDLYNDSLRFRLLRQHGYNISSGDIF<br>KKFKDANGNFKESLIVDVPGMLSLYEATH<br>LRVHGEEILEEALVFTTNHLELAISQKDLC<br>RSLERLSASNYIPIYQATTSQNEALLKLAK<br>LDFNLFQALYKEELSEVSRWWKELDFER<br>KMPFARHRIVECFFWMVGMYYYEPKYSV<br>GRMIGTKLCAMATMLDDIFDSYGTFEELK<br>IFRQVIERWDVNCCMDDLPRYMKVYYHS<br>LCDVMNEIEEELEKLGMPDRVHYAKRVL<br>KDQARDYLVEAEWLHEGCTPSMEEYLPV<br>RVQSVGVCMTVVFSLRGMNDNITKETFE<br>WVLKYPKIVMASSFIFRFMDDIGGSKHEK<br>QKGDVDSSIDCYMKQYGVSKEDAIEVLD<br>KQIVDSWKEINEDLLRPTVVPIFGTPTHID<br>FGLIKSIILEEI* |
| Fxa3Cg101875.1 | M/S TPS | *FaTPS20* | *F. x ananassa* | a | - | FaRR1 Genomic | MSVTAVILASQSPQSLNNADTRRSANFH<br>PSIWGDRFLSCNNTMETDMKGEQQVLQ<br>QLKEEVKRMLMAPPVETGLGKLELIDDIQ<br>RLGVSYHFEYEIDQTMQQIHENLNGTCD<br>DDDLHTCALRFRLLRQHGYNVSCADMFN<br>KFKDCDGNFDESLHHDIVGLQSLYEATHL<br>RVRGEEFLEEALAFTTTRLESAANLLSAP<br>LLKQVSHALNQPLWKGLPRLEARHYMSL<br>HQELHGSRNHVLLRFAKLDFNLLQQVHL |

| | | | | | | | |
|---|---|---|---|---|---|---|---|
| | | | | | | | KELSDIARWWKKLDFASKLPFARDRVIEC<br>YFWILGVYFEPKYYFARKTLTKVIAMTSIID<br>DIYDVYGTLEELDLFTRAIERWDISAMDEL<br>PEYMQVCYEALLDIYAEAEQGLASEGKS<br>YRIDYAKEAMKRLVRAYHAEAKWFHNNY<br>TPTMDEYMEVALVTSAYSMLATTSFVGM<br>GDIVTKDSFEWIFSDPKMVKASAVVCRL<br>MDDIVSHKFEQKRGHVASAVECYMTQY<br>GATEEETIIEFRKQVSDAWKDINEECLHP<br>TSLDMPLLMRVLNLTRVIDVVYKSEDGYT<br>HAGTVLKDFVASLLVDPVTV* |
| Fxa3Dg200246.1 | M/S TPS | *FaTPS21* | *F. x ananassa* | b | - | FaRR1 Genomic | MDCSKQLRNPQAEEQQIHQWQTKSEFS<br>AHSVHDQSRRSANYKPNIWKYDFLESLN<br>SKFDGGGYLIQMQKLIKEVKKLFVECKES<br>DVIAKLELIDSIRKLGLNNHFEQEIKAAVDA<br>IASAELENNRNPCISSEGDDLYAAALFFKI<br>LRQQGYQVSQDIFGRYMDEMGTLKNST<br>SGNVKGMIELLETSNLAFEGENFLEKAQA<br>FLIATLRDTDTMWDEIDSSNSKNVTYAFQI<br>SSQRRAQWFNVMKSERHTNQPYMTTLL<br>ELAKLNFNVVQATLQKDLREAYKWWYNL<br>GLTKNLDFARDRMVECFMCAVGLAFETD<br>HKSFRKWLTKVINLILIIDDVYDVYGSLEEL<br>KHFTRAVERWDVMETEQLPECMKICFQV<br>LYNTTCEIAYEILEENGRNQVLPHLCKVW<br>ADFCKALLVEAEWYNEAYTPSFEEYLSN<br>GYISSSASLIFTHAFFATKHEEGVDDFLH<br>MNEDLLYNISVILRLLNDLGTSAAEQERG<br>DAASSILCYMREMNVSEEIARKNIKSMID<br>NAWKKINENCFARNPKISSSYINITTNIAR<br>VGHSLYQDGDGFGDQEQGTRELIQSLLV<br>EPLS* |
| Fxa3Dg200253.1 | M/S TPS | *FaTPS22* | *F. x ananassa* | g | - | FaRR1 Genomic | MGDDIFVQHSQKLELLRNVLRNVAEVDA<br>LEGLNMIDAVQRLGIDYHFQREIDEILYKQ<br>MSIVSACDDLHEVALRFRLLRQHGYSVP<br>EDMFNNFKDSKGTFKQALGEDIKGLMSL<br>YEASQLGTEGEDTLVEAEKFSGHLLKTSL<br>SHIDHRRARTVGNTLRNPHHRSLASFMA<br>RNFFVTSQATNSWLNLLKEVAKTDFNMV<br>RSLHQNEIVQISKWWKELGLAKELKFARD<br>QPLKWYTWSMAGLTDPKLSEERVELTKP<br>ISFVYLIDDIFDVYGTLDDLILFTEAVNRWE<br>ITAIDHLPDYMKICFKALYDMTNEFSCKVY<br>QKHGWNPLQSLKISWASLCNAFLVEAKW<br>FAAGQLPKSEEYLKNGIVSSGVNVVLVH |

| | | | | | | | |
|---|---|---|---|---|---|---|---|
| | | | | | | | MFFLLGQNITRKSVELLNETPAIISSSAAIL<br>RLWDDLGSAKDENQDGNDGSYVRCYLE<br>EHEGCSIEEAREKTINMISDEWKKLNRELI<br>SPNPFSATFTLASLNLARMIPLMYSYDGN<br>QCLPSLKEYMKLMVYETVSM* |
| Fxa3Dg201726.1 | M/S TPS | *FaTPS23* | *F. x ananassa* | a | - | FaRR1 Genomic | MPSQSAAAPPSQTMSEIAGLTAIYQPGIC<br>GDGFNNFESLDNITRAHMEKQINQLKVLV<br>RELLTSTAAGLSHQLKLIDDIQRLGVSYHF<br>ETKLAEALENIHSRLHDGDLDLYNDSLRF<br>RLLRQHGYNISSGDIFKKFKDANGNFKES<br>LTADVQGMLSLYEATHLRVHGEEILEEAL<br>VFTTNHLELAISQVSCPLKAQISDALQRP<br>MWRSLERLSARNYIPIYQATTSHNEALLK<br>LAKLDFNLVQALYKEELSEVSRWWKELD<br>FEMKMPFARHRIVECFFWMVGMYYEPK<br>YSVGRKIGTKLCAMATMLDDIFDSYGTFE<br>ELKIFRQVIERWDVKCCMDDLPRYMKVY<br>YHSLCDVMNEIEEELEKLGMPDRVHYAK<br>QVLKDQARDYLVEAEWLHEGCTPSMEE<br>YLPVRVQSVGVCMTVVFSLLGMNDNITK<br>ETFEWVLKYPKIVMASSFIFRFMDDIGGS<br>KHEKQKGDVDSSIDCYMKQYGVSKEEAI<br>EVLDKQIVDSWKEINEDLLRPTAVPMYVL<br>LRVINFSRDVDLVYKGEDGFTHVGKVVK<br>HAVAALFGDPLPLE* |
| Fxa4Cg202367.1 | M/S TPS | *FaTPS24* | *F. x ananassa* | a | - | FaRR1 Genomic | MALHQVLASPAPTLNAAFDVKPRPANYA<br>PCMWGDHFLSNSSMEVDVKLEQRVQEL<br>KEEVKRMLMAAVNEPASHMLDMVDNIQR<br>LGLSYSFESEIDTILKHILERMVTVTTLYFR<br>LLRQQGYNVSCDVFNKFKESNGTFKEYL<br>TSDVVGLLSLYEATHLRVHGEDILEEALIF<br>TRTHLESAAPRLSSPLSKQVIHALYQPLW<br>KGFSRVEARHYMSVYEEYDSHNKNLLTL<br>AKLDFNLMQKVHQKELCDMTRWWKDLD<br>FANKLPFARDRIVETYFWALATIFEPEYQF<br>ARTMVCKYGALTTVQDDVYDVYGSYQEL<br>ELYTDAIERWDISAIDDQLPQCLKDCYKG<br>LLDSYSSYEEKLAKEGNLYRLDYAKEAIKI<br>QLRGYFQEAKWLKEKYTPTMDEYIPNGL<br>NSSFYPMAITSFVGMGGLVTKDTMDWVL<br>NDPKIVKATALTGRLLNDMAGHKFEQDIE<br>QGASSVECYMNQYGVTEEEAKIELTKQK<br>DDAWKDINQEWLDLHSSNSIPKPLLQIILN<br>LARSGEVLYKNEDLYTNAEAGLKGFIVSL<br>LIEPAAWHMLDMVDNIQRLGLLYCFKNEI |

| | | | | | | | |
|---|---|---|---|---|---|---|---|
| | | | | | | | DILLKHESNGTFKEYLTSDVVGLLSLYEAT<br>HLRVHGGDILEQALTFTTTHLESAAPRLS<br>SPLSKQVTHALYQPLWKGYSRLEARHYM<br>SVYEEYDSHNETLLILAKLDFNLMQKVYQ<br>KELCDMTRWWKDLDFANKLPFARDRIVS<br>YFWALATIFEPEYHFARTIACIYGALTTVR<br>DDVYDVYGSYQELEHFTEAIERWDISAID<br>DQLPQCLKDRYKGLLDSYSSYEEKLAKE<br>GKLYRLDYAKEAIKIQLRGYFQEAKWLKE<br>KYTPTMEEYIPNGLNSSFYPMAITSFVGM<br>GGLVTKDTMDWVLNHPNEPASHMLDMV<br>DNIQRLGLSYRFENEIDTILKHCERPFVLY<br>RPVPTTARLHPSGLQQRPRPLFLPGESA<br>KFQFPARSTLFEVLGDLAGQLEKNGLAD<br>VFNKFKESNGTFKEYLTSDVVGLLSLYEA<br>THLRFEQNNEQCASSVECYMNQYGVGE<br>EEAKIALTNQRMLHGKT* |
| Fxa4Dg102142.1 | M/S TPS | *FaTPS25* | *F. x ananassa* | a | - | FaRR1 Genomic | MASAQAQITNAASDVKRCSPHYTPGIEG<br>QHFLSNSAAADLEANVVLEQHFQELKEQ<br>VKRMLMAAVNQPLHILDMIDNIQRLGLSN<br>HFDKAIDAILEQVHNAFGTEDQENGGDLY<br>TTALRFRLLRQQGYNVSCDVFNKFKQSN<br>GTFKEHLMSDIVGLLSLYEATHLRVHGED<br>ILEEALTFTTSHLESAAPQLSSPLSKQVTH<br>ALYQPLWKGFSRLEARHYMSVYEEDDS<br>HNQTLLTLAKLDFNLAQKVDQKELSDMT<br>RWWKDLDFVNKLPFARDRLVEAYFWGL<br>ATCYEPEYYFARIMACKSVALSTVLDDVY<br>DVYGTYEELERFTEAIERWDISAVDQVQL<br>PQCMEVCYKGLLDSYSGYEEKLADEGNL<br>YRIEFAKEAIKIQLRGYFQEAKWLKEKYTP<br>TMDEYIPNALDTSFFPMAITTFVGMGGLV<br>TKHTMDWVLNHPKIVKVTALIGRLLNDMA<br>GHKFEQNNGQCASSVECYMNQYGVGEE<br>EAKIALTKQKDAAWKDLNQELFDLHSSNS<br>IPKPLLQIILNLARSSEVIYSNNEGYTNSSV<br>LKGFIDSLLIEPVPV* |
| Fxa4Dg102144.1 | M/S TPS | *FaTPS26* | *F. x ananassa* | a | - | FaRR1 Genomic | MALHQVLASPAQTLNATFDVKPRPANYA<br>PCIWGDHFLSYSSMEVDIKLEQRVQELKE<br>EVKRMLMAAVNESTSHMLDIVDNIQRLGL<br>SYRFENEIDTILKHVHDHFYGSKDLGKDG<br>DLYTTALYFRLLRQQGYNVSCDVFNKFK<br>ESNGTFKEYLTSDVVGLLSLYEATHLRVH<br>GEDILEEALIFTRTHLESAAPRLSSPLSKQ<br>VIHALYQPLWKGFSRVEARHYVSVYEEY |

| | | | | | | | |
|---|---|---|---|---|---|---|---|
| | | | | | | | DSHNKNLLTLAKLDFNLMQKVHQKELCD MTRWWKDLDFANKLPFARDRIVETYFWA LATIFEPEYQFARTMVCKYGAPTTVQDDV YDVYGSYQELELYTDAIQRWDISAIDDQL PQCLKDCYKGLLDSYSAYEEKLAKEGNL YRLDYAKEAIKIQLRGYFQEAKWLKEKYT PTMDEYIPNGLNSSFYPMAITSFVGMGGL VTKDTMDWVLNDPKIVKATALTGRLLND MAGHKFEQDIEQGASSVECYMNQYGVT EEEAKIELTKQKDDAWKDINQEWLDLHS SNSIPKPLLQIILNLARSGEVLYKNEDLYT NAEAGLKGFIVSLLIEPVPV* |
| Fxa5Ag200582.1 | M/S TPS | *FaTPS27* | *F. x ananassa* | a | - | FaRR1 Genomic | MDPQLVSASQAQTQNGSSTPDVERRSA NFSPSVWGDYFLKYASVETDDIEAEKSIK ELKEEVKGVLRLSSPQRLDLIDYIQRLGV AHHFEDEIHQLLQQIHNQYSSSYDDDDLH TVALRFRLLRQQGFKVSCDMFNKFIDVD GNLKESSVADVPGLLSLYEAAHLRTYGD DILDRAMSFTTTHLESAVAAQGRLIPPLS NQVAHALYQPLWMGNPRIEARHYLSSYQ ELNSAPHFSQSLLTFAKLDFNQLQRIHQK ELSEITRWWKELDFVNKLPFARNRIVEAY FWSLGVCFEPKYRLPRETICKALGLLTMI DDTFDTYGKFDELELFAEAVERWDLSAM DSLPDYMKIIYEAVLNTYTEIETELAKEGN SYRIEYLVEGMQCQVRAYLKEAKWYHLK CTPKTYDEYMRVGLITSGMFLVEAAVIVP LAGEISTRDSLESLFRDPNNKIVYASNILG RLLNDIRSHKHEQKRGNDFSAIQCYMKE HCVTEEEALIELNKQVNDAWKDVNEVLL QPPTTVPRPILLLCLNLFVLATILSVLSRGA TSDEEFTAQAIDTHGGVSRATIGKFPRIG ASSGEL* |
| Fxa5Ag203430.1 | M/S TPS | *FaTPS28* | *F. x ananassa* | a | - | FaRR1 Genomic | MSLQVLASQSQTPNARPSANYTPSIWGD HFLSFATAPDTGADVKLPLAYYEYAPCIW DDHFLSYSTTKVNIKLEQHVRELKEEVKR MLMDLANVSGSQQIDLVNDIQRLGVAYH FENEIDEILKQIHHNYSCGTDDDDLYTTAL RFRLLRQHGYNISCDTFNKFKEGNHQTF KASLHSDVSELLSLYEATHLRVHGEEILE EALAFTTTHLELIKHSLSPPLSNLVSHSLN QPLRTSVARVEARHYLSIYQECDSHNETL LTLAKLDFNLVQQLHQKELCEITRWWKNL DVKNKLPFARDRVTEVYFIWVLSVYFEPR FSFARRTSCKVTAILSIIDDIYDSQGTLEEL |

| | | | | | | | |
|---|---|---|---|---|---|---|---|
| | | | | | | | ELFTEAIQRWDICAIDPLPDYMKVFYKAM LEVYIEIEEELAKEGNLYRIHYAIEAMKKQ ATYYFHEAKLLQQKHIPTLDEYMDLALPS TAYHMLITTAFVGMGDIATQDSFDWLATY PRAVKGAEVVCRLMDDIADYKFELERGT DISSVECYMKEHGATEEEAMTELRRRLS DAWKDINESFFLPNALPRPLLTRVLNFAC AMDVAYKYEDSYTTHHASILKDFIVPTLVE SVPL* |
| Fxa5Bg100567.1 | M/S TPS | *FaTPS29* | *F. x ananassa* | a | - | FaRR1 Genomic | MDPQLVSASQAQTQNGSSTPDVERRSA NFSPSVWGDYFLKYASVETDDIEADKSIK ELKEDVKGVLLSSPQRLDLIDYIQRLGVA HHFVDEIHQLLQQIHNKYSSSYDDDDLHT VALRFRLLRQQGFKVSCDMFNKFIDVDG NLKESSVADVPGLLSLYEAAHLRTYGDDI LDRAMSFTTTHLESAVAAQGRLSPPLSN QVAHALYQPLWKGIEARHYLSCYQELNS APHFSQSLLTFAKLDFNQLQLIHQKELSEI TRWWKELDFVNKLPFARNRIVEAYFWSL GVCLEPKYRLPRETICKALGLLTMIDDTFD TYGKLDELELFAEAVERWDLSAMDSLPD SMKIIYEAVLNSYTEIEAELAKEGNSYRIA YLVEGMQCQVRAYMKEAKWYHLKCTPK TYDEYMRVALITSGMFLVEAAVIVPLAGEI STRDSLESLFRDPNNKIVYASNILGRLLND IRTQKQITNFHNKHEQKRGNDFSAIQCFM KEHCVTEEEALIELNKQVNDAWKDVNEV LLQPPTTVPRPILLLCLNLLRVTDVIYKND DGYTNGAVVLKDYITSLLVEPAPM* |
| Fxa5Bg103205 | M/S TPS | *FaTPS30* | *F. x ananassa* | a | - | FaRR1 Genomic | MSLPVLASQIHKPDTGADVKIPLAYYEHA PCIWDDHFLSYSSTKVDIKLEQHIQELKEE VKRMIMDPANVSGSQQIDLVNDIQRLGVA YHFENEIDEILKQIHHNYSCGIDDDDIYTT ALRFRLLRQQGYNISCGKLNANLCSSKEL PTDADIFTKFKDGNHQTFKASLQSDVSGL LSLYEATHLRVHGEEILEEALAFTTTHLES IKHSLSPPLSNLVSHSLNQPLRTGVARVK ARHYLSIYQDCDSHNETLLTLAKLDFNLV QQLHQKELCEITRWWKNLDVKKKLPFAR DRVTEVYFIWVLSVYFEPQFSFARRTSCK VTALLSIIDDIYDSQGTLEELELFTEAIQRW DICATDPLPDYMKVFYKAMLEVYIEIEEEL AKEGNLYRIHYAIEAMKKQATYYFHEAKL LQQKHIPTLDEYMDLALPSTAYHMLITTAF VGMGDITTQDSFDWLATYPRAVKGAEVV |

| | | | | | | | |
|---|---|---|---|---|---|---|---|
| | | | | | | | CRLMDDIADYKFEQERGTDISSVECYMK<br>EHGATEEEAMTELRRRLSDAWKDINESF<br>FLPNALPRPLLTRVLNFACAMDVAYKYED<br>SYTSHHASILKDFIFLTLVESVPL* |
| Fxa5Cg200524 | M/S TPS | *FaTPS31* | *F. x ananassa* | a | - | FaRR1 Genomic | MDPRLVSASQAQTQNGSSTPDVERRSA<br>NFPPSVWGDYFLKYASVETDDIKADKCIK<br>ELKEDVKGALLSSPQRLDLIDYIQRLGVA<br>HHFEDETHQLLQQIHNQYSSSSSDDDDD<br>LHTVALRFRLLRQEGFEVSSADMFNKFID<br>VDGNLKESCVADVPGLLSLYETAHLRKH<br>GEDFLDRAMSFTTTHLESAVAAQGRLSP<br>PLSNQVAHALYQPLWKGNPRIEARHYLS<br>SYQELNSAPHFSQSLLTFAKLDFNQLQRI<br>HQKELSEITRWWKELDFVNKLPFARNRIV<br>EAYFWSLGVCFEPKYRLPRETICKALGLL<br>TMIDDTFDTYGKLDELELFAEAVERWDLS<br>AMDSLPDYMKIIYEAVLNSYTEIEAELAKE<br>GNSYRIEYLVEGMQCLVRAYLKEAKWYH<br>LKCTPKTYDEYMRVGLITSGMFLVEAAVI<br>VPLAGEISTRDSLESLFRDPNNKIVYASNI<br>LGRLLNDIRSQKHEQKRGNDFSAIQCYM<br>KEHCVTEEEALIELNKQVNDAWKDVNEV<br>LLQPPTTVPRPILLLCLNFLRVTDVIYKND<br>DGYTNGGAVLRDYITSLLVEPAPM* |
| Fxa5Cg202929 | M/S TPS | *FaTPS32* | *F. x ananassa* | a | - | FaRR1 Genomic | MSLQVLASQSQTPNARPSANYTPSIWGD<br>HFLSYDTVEVDFSHKQQFQELKRQVKM<br>MLMVPVKEPSDKLNLVDDIQRLGVSYHF<br>ENEIDEVLEQIYQTTHGSDMANDDDLCTT<br>ALRFRLLRQQGYNVSCDIFNKFKEGNDQ<br>TFKQTLQSDVSGLLSLYEATHLRIHGEDIL<br>EEALVFTTTHLESIKHSLSHSLSKQITHSL<br>KQPLRKGIIRVEARYYLSIYHEFDSHNETL<br>LTFAKLDFNLLQQVHQIELCEVTRWWKD<br>LDVRNELPFTRDRVTEVYFCWALSICFEP<br>EYAFARRTLCKVTALTSILDDIYDLYGTLE<br>ELELFTEAIERWDIRVIDALPNYMKVCYKA<br>LLDVYTEVEEELAKERKLYLIHYAREAMK<br>LQARSYYLEAKWFHQKYMPTMDEYIALS<br>SLTSGYPLLITTTFVTMEEITTRDPFDWLA<br>TYPKSVKGSAVVARLTDDLVSHKFEQKR<br>GHVASAVECHMKEFGATEEETIIELRSKV<br>SDAWKDINESFLVPNDVSRPLLTRILNFA<br>CVMDVVYKYEDGYTTQHGMLKDLITSVLI<br>EQVPV* |

| | | | | | | | |
|---|---|---|---|---|---|---|---|
| Fxa5Dg200534 | M/S TPS | *FaTPS33* | *F. x ananassa* | a | - | FaRR1 Genomic | MDPQLNSASQPQTQNGSSGIPHVDRRS<br>ANFSPSIWGDYFLAYTSLETKDTEANQRI<br>QELKEEVKRVIISSLPSQRLDLIDYIQRLG<br>VSYRFEDEIHQLLQQIHNRYSSSSSSSSC<br>YDDDDDDLHTVALRFRLLRQQGFKVSCY<br>MFNSPTVELSKIRSVFSGAEKTGTLRFID<br>VDGNLKESCVADLPGLLSLYEAAHLRTY<br>GDDILYRAMSFTTIHLDGSTRPFKPPTLK<br>SSSSCLISITLEGKPRIEARHYLSSYQELN<br>STPHFSDFCKVGFQSIAAYPSERTMWWK<br>ELNSVNKLPFARDRIVEGYFWSLGVYFEL<br>KYHFGRTTLCKVIVLTTMLDDVYDVHGTP<br>EELEQFTEAVQRWDISATDSLREYMKVL<br>YHAMLNVYTDIEDELAKEGNSYRINYAIEA<br>IKIQVRFYMKEARWFHQSYTPKTLDEYM<br>SVARVTAAMFLLETTFTVATAGDIDSQTIG<br>RLLDDIRSHKFEQKRGHVASAVECYMKE<br>HCVTEEEAVIELTKQVSDGWKDVNEGLL<br>NPIATFPRPLVLHNFNFLRVTDVVYKYED<br>GSWSCA* |
| Fxa5Dg200537 | M/S TPS | *FaTPS34* | *F. x ananassa* | a | - | FaRR1 Genomic | MDPLLVSASQSQTQNGSSTADVERRSA<br>DFSPSVWGEYFLAYASVETADIEADKRIK<br>ELKEEVKRVLLSSPPSQRLDLIDYIQRLGV<br>SHHFEDEIHQLLQKIHNQYSSSSSSDDDD<br>DLHTVALRFRLLRQEGFEVSSADMFNKFI<br>GVDGNLKESCVADVPGLLSLYETAHLRK<br>HGEDFLDRAMSFTTTHLESAVAAQGRLS<br>PPLSNQVAHALYQPLWKGYPRIEARHYL<br>SNYQELNSAPHFSQSLLTFAKLDFKFFQR<br>VHQKELSEITRWWKELDFVNKLPFARNRI<br>VEAYFWSLGVCFEPKYRLPRETLCKALG<br>LLTMIDDTFDTYGKLDELELFAEAIQRWD<br>LSAMDSLPDYMKIIYEAVLNAYAEIEAELS<br>KEGNSYRIAYLVEGMQCLVRAYLKEAKW<br>YHLKCTPKTYDEYMRVGLITSGMFLVEAA<br>VIVPLAGEISTRDSLESLFRDPNNKFVYAS<br>NILGRLLNDIRSHKHEQKRGNDFSAIQCY<br>MKEHGATEEEALIEMNKQVNDAWKDVN<br>EVLLQPPTTVPRPILLLCLNFLRATDVIYK<br>NDDAYTNGGGVLKDYITSLLVEPAPM* |
| **Fxa5Dg200537.1** | M/S TPS | ***FaTPS35*** | *F. x ananassa* | **a** | **93.4** | **FaRR1 transcript** | MDPLLVSASQAQTQNGSSTADVERRSA<br>NFSPSVWGEYFLAYASVETADIQADKRV<br>KELKEEVKRVLLSSPPSQRLDLIDYIQRLG<br>LSHHFEDEIHQFLQQIHNQYSSSSSDDDD<br>LHTVALRFRLLRQEGFNKVSCDMFNKFID |

| | | | | | | | |
|---|---|---|---|---|---|---|---|
| | | | | | | | VDGNLKESCVADVPGLLSLYETAHLRKH<br>GEDFLDRAMSFTTTHLESAVAAPGRLSP<br>SLSNQVAHALYQPLWKGNPRIEARHYLS<br>SYQELNSAPHFSQSLLTFAKLDFNFFQRI<br>HQKELSEITRWWKELDFVNKLPFARDRIV<br>ECYFWSLGVCFEPKYRLPRETLCKILGLL<br>TMIDDTFDTYGKFDELELFTEAIQRWDLS<br>AMDSLPDYMKIIYEAVLNAYTEIEAELAKE<br>GNSYRIAYLVEGTQCLVRAYMKEAKWYH<br>LKCTPKTYDEYMRVGLITSGMFLVEAAVI<br>VPLAGEISTRDSLESLFRDPNNKIVYASNI<br>LGRLLNDIRSHKHEQKRGNDFSAIQCYM<br>KEHCVTEEEALIELNKQVNDAWKDVNEV<br>LLQPPTTVPRPILLLCLNFLRGTDVIYKND<br>DGYTNGGGALKDYITSLLVERAPM* |
| **Fxa5Dg200537.1** | M/S TPS | ***FvTPS35*** | *F. vesca* | **a** | **93** | **UC06 transcript** | MDPLLVSASQAQTQNGSSTADVERRSA<br>NFSPSVWGEYFLAYASVETADIQADKRV<br>KELKEEVKRVLLSSPPSQRLDLIDYIQRLG<br>LSHHFEDEIHQFLQQIHNQYSSASSDDDD<br>LHTVALRFRLLRQEGFNKVSCDMFNKFID<br>VDGNLKESCVADVPGLLSLYETAHLRKH<br>GEDFLDRAMSFTTTHLESAVAAPGRLSP<br>SLSNQVAHALYQPLWKGNPRIEARHYLS<br>SYQELNSAPHFSQSLLTFAKLDFNFFQRI<br>HQKELSEITRWWKELDFVNKLPFARDRIV<br>ECYFWSLGVCFEPKYRLPRETLCKILGLL<br>TMIDDTFDTYGKFDELELFTEAIQRWDLS<br>AMDSLPDYMKIIYEAVLNAYTEIEAELAKE<br>GNSYRIAYLVEGTQCLVRAYMKEAKWYH<br>LKCTPKTYDEYMRVGLITSGMFLVEAAVI<br>VPLAGEISTRDSLESLFRDPNNKIVYASNI<br>LGRLLNDIRSHKHAQKRGNDFSAIQCYM<br>KEHYVTEEEALIELNKQVNDAWKDVNEV<br>LLQPPTTVPRPILLLCLNLLRVTDVIYKND<br>DGYTNGGVVLKDYITSLLVEPAPM* |
| **Fxa5Dg200537.1** | M/S TPS | ***FaTPS36*** | *F. x ananassa* | **a** | **93** | **DPW transcript** | MDPLLVSASQAQTQNGSSTADVERRSA<br>NFSPSVWGEYFLAYASVETADIQADKRV<br>KELKEEVKRVLLSSPPSQRLDLIDYIQRLG<br>LSHHFEDEIHQFLQQIHNQYSSSSSSDDDD<br>LHTVALRFRLLRQEGFNKVSCDMFNKFID<br>VDGNLKESCVADVPGLLSLYETAHLRKH<br>GEDFLDRAMSFTTTHLESAVAAPGRLSP<br>SLSNQVAHALYQPLWKGNPRIEARHYLS<br>SYQELNSAPHFSQSLLTFAKLDFNFFQRI<br>HQKELSEITRWWKELDFVNKLPFARDRIV |

| | | | | | | | |
|---|---|---|---|---|---|---|---|
| | | | | | | | ECYFWSLGVCFEPKYRLPRETLCKILGLL<br>TMIDDTFDTYGKFDELELFTEAIQRWDLS<br>AMDSLPDYMKIIYEAVLNAYTEIEAELAKE<br>GNSYRIEYLVEGMQCQVRAYLKEAKWYH<br>LKCTPKTYDEYMRVGLITSGMFLVEAAVI<br>VPLAGEISTRDSLESLFRDPNNKIVYASNI<br>LGRLLNDIQSHKHEQKRGNDFSAIQCYM<br>KEHCVTEEEALIELNKQVNDAWKDVNEV<br>LLQPPTTVPRPILLLCLNLLRVTDVIYKND<br>DGYTNGGVVLKDYITSLLVERAPM* |
| Fxa6Ag100563 | M/S TPS | *FaTPS37* | *F. x ananassa* | a | - | FaRR1 Genomic | MSMIIEPSLSSPVPQIAMPEIVRPIANFQP<br>SIWGDQFLNYDSQDIKTGAFWQQQVEQL<br>KVNVKSKVFTNECDFALRLKLIDAIQRLGV<br>AYHFEEEIEDALRHIHATYRDQDHFNDDS<br>DLSTVALCFRLLRQHGYEISSDMFNKFKD<br>ENGSFKEGLVADGCAMLSLYEAAHLKVH<br>GENILEEALVFTTAHLESAKSSACYSTQM<br>TEQITQALVRPLRTSLERICAKRHMSVYQ<br>LEDDAALIKHNETLLKLAKMDFNLVQFLH<br>KKELCEITRWWKELDFERKLPFARDRKV<br>ELFFWIVGVYFEPQYSIGRIFMTKVAILLT<br>VMDDIYDAYGTFEELVVFTEAIDRWDVKC<br>IDELPDYLKIFYYELLNVFHEMDTVLAKEG<br>RSYRVCYAIQAVRNTQMIDMKDQARSYF<br>NEARWLHEGRTPSMEEYMSVATVSISYT<br>FLTTISLLGMGDIVTKDSFEWLSNGPKIVR<br>ASNIIFRLMDDIVSTKFEKERGGHAPSSID<br>CYTKQYGVSEQEAIDVFNKQIVESWKDIN<br>EEFLKPTAVPMPVLMRVLNLTRVADLLYK<br>GEDGFTRVGKMTKDSVAAVIIDSVPL* |
| **Fxa6Ag100563.1** | M/S TPS | ***FvTPS37*** | *F. vesca* | **a** | **96.2** | **UC06 transcript** | MSMIIEPSLSSPVPQIAMPEIVRPIANFQP<br>SIWGDQFLNYDSQDIKTGAFWQQQVEEL<br>KVNVKSKVFTNECDFALRLKLIDAIQRFG<br>VAYHFEEEIEDALRHIHATYRDQDHINDD<br>SDLSTVALCFRLLRQHGYAISSDIFNKFKD<br>ENGSFKEGLVADGCAMLSLYEAAHLKVH<br>GENILEEALVFTTAHLESAKSSACYSTQM<br>TEQITQALVRPLRTSLERICAKRHMSVYQ<br>LEDDAALIKHNETLLKLAKMDFNLVQFLH<br>KKELCEITRWWKELDFERKLPFARDRKV<br>ELFFWIVGVYFEPQYSTGRIFMTKVAILLT<br>VMDDIYDAYGTFEELVVFTEAIDRWDVKC<br>IDELPDYLKIFYYGLLNVFNEMDKVLAKEG<br>RSYRVCYAIQAMKDQARSYFNEARWLHE<br>GRTPSMEEYMSVATASISYTFLTTISLLG |

| | | | | | | | |
|---|---|---|---|---|---|---|---|
| | | | | | | | MGDIVTKDSFEWLSNGPKIVRASNIIFRLM DDIVSTKFEKERGHAPSSIDCYTKQYGVS EQEAIDVFNEQIVESWKDINEEFLKPTVV PMPVLMRVLNLTRVVDLLYKGEDGFTRV GKMTKDSVAAVIIDSVPL* |
| Fxa6Ag101058 | M/S TPS | *FaTPS38* | *F. x ananassa* | a | - | FaRR1 Genomic | MALHQVLASPAQTLNAAFDVKPRPANYA PCIWGDHFLSYSSMELDVKVEQHVQELK EEVKRMLMVVVNEPASHMLDMVDNIQRL GLSYHFKNEIDTILKHAHDHFYGSKDFGK DGDLYTTALHFRLLRQQGYNVSCDVFNK FKESNGTFKEYLTSDVVGLLSLYEATHLR VHGEDILEEALIFTTTHLESAPPRLSSPLS KQVTHALYQPLWKGFSRVEARHYMSVY EDYDSHNKTLLTLAKLDFNLMQKVHQKQ LSDMTRWWKDLDFANKLPFARDRIVETY FWALATIFEPEYHFARTMVCKYGALTTVQ DDVYDVYGSYQELELYTDAIERWDISAID DQLPQCLKDCYKGLLDSYSAYEEKLAKE GNLYRLDYAKEAIKIQLRGYFQEAKWLNK KYTPTMDEYIPNGLNSSFYPMAITSFVGM GGLVTKDTMDWVLNDPKIVKATALTGRLL NDMAGHKFEQDIEQGASSVECYMNQYG VTEEEAKIELTKQKDAAWKDINQEWLDLH SSNSIPKPLLQIILNLARSGEVLYKNEDLY TNAEAGLKGFIVSLLIEPEKGWGVDFTSH PIPSDLQLLEPGKVYLISKERNGFGDPGH SQLDFEDRKVTAINQNLCNTAGKSLDVK RTSALGLK* |
| Fxa6Ag101852 | M/S TPS | *FaTPS39* | *F. x ananassa* | a | - | FaRR1 Genomic | MSTQLLASLSIQTPKIVNPEILRRTANYHP SIWGDQFSNYEESQDPMTRSLLQKQVN QLKVVVKREVFRNASSDFSTQMKSIDAM QRLGIAYHFERDIENALEQAHAACVHDG NLYNVALRFRLLREHGYNVSSDIFKRFKD ANDNFKEGLTSDLPGMLSFYEATHLRIHG EDILEDGLHFTTAHLESSANDVNHPLAAQ ITQALERPLRKFPERLYARHYISIFPDDPK SLHHEAVALLLKLAKLDFNLVQSLHKKEL SEIIRWWDELDFERELPFARNRIVELYFW TLGVYAEPQYSKARHFLTKAIAFGSVLDDI YDAYGSFEELKIFTEAIQRWDVNCIDELP DYMKIFYRRLFILCNEIEVEMVKQGKSYR VHFAKEIIKAQARLYFAEAQWLHENYTPS MDEYMEVSVRSVGNSFLSTMSLVGMGDI VTKDAFEWLANHPKILRASNIIFRLMDDLV SGKFEKEREHIASSIDIYMKQYGVSEQET |

| | | | | | | | |
|---|---|---|---|---|---|---|---|
| | | | | | | | VDIFNKKVADSWKDINEEFLRPTAVPVPV LTRVLNLTRVVDLLYKKDDEYTRVGDVM KDGVASLLIHPVPL* |
| Fxa6Ag104161 | M/S TPS | *FaTPS40* | *F. x ananassa* | b | - | FaRR1 Genomic | MSLLFCSSSPSILFTKLTSATRYLRMLMP APVLVHVHHTTATKQSSFTLSNSSAKVID PPVAHQRRLAQYHPTIWGPKLIDSFSTHY THESYASQLEGLKQNVSRAWVVSSTKG DACSILKLIDSMQRLGVAYHFEKEIQAAIG ALVSSWNASTTTDLQTVALQFRILRQHGI SISTDLFNKFRNGEGSFKDSLSNDVEGLL SLYESSHLGIPGEDVMEEAKSFSSKNLKQ SMSTILNDDNLLKRVEKSLETPVHWRMP RIEAIDFISMYQRDDSSNLALLELAKLDYN LVQSVHQKEIKELSRWWRELDFKSKASF SRDRLMENYLWALGITYEPQFSECRIGLT KFVCILSAIDDMYDVYGFLDELELFTDAVT QWNLKANEDLPEYMKPIYSAMFTFGNEL ADQNYGLNTLPLIKKEWENLCKSYMVEA RWFYGGYTPTLEEYLKNAWTSVGGPGA MLHAYLLSQGSQLTKASLESYNHGSQLIY WASIITRLTDDLGTSKAESERGDVAKSVH CYMEEKGISEEEAQEYIKGFICYSWKKIS EESAKATIPKSIVNMSLNMARTAHCIFQH GDGIGTSVGVTKDRLVSLITNPIPIDDDHH HHVIHVTQ* |
| **Fxa6Ag104161.1** | M/S TPS | ***FcTPS40*** | *F. chiloensis* | b | **95.6** | **ILE transcript** | MAMPAPVLEHEHHTTATKQSSFILSNSSS KVIDPPVAHQRRLAQYHPTIWGPKLIDSF STRYSHESYASKLEGLKQNVSEYLHVLST KAKGDACLTLKLIESMQRLGVAYHFEKEI KAAIVTLVSSGTASTTTDLQTVALQFRIQR QYGISISSDLFNQFRNSEGSFKDSLSNDV EGLLSLYEASHLGMPGEDVLEEAKSFSS KNLKQSLTTILNDDNLRKRVDQSLETPVH WKMPRIEALDFINMYQRDDSRNLVLLELA KLDYNLVQSVHQNEIKELSRWWRELDFK SKASFSRDRLMENYLWALGITYEPQFSE CRIGLTKFVCILSAIDDMYDVYGFLDELEL FTDAVTQWNLKANEDLPEYMKPIYSAMF TFGNELADQNCGLNTLTLIKKEWENLCKS YMVEARWFYGGYTPTLEEYLKNAWTSV GGPGAMLHAYLLTQGSQLTEASLESFNH GSQLIYWASIITRLTDDLGTSKAESERGD VAKSVHCYMEEKGISEEEAQEYIKGLICY SWKKINEESAKATIPKSIVNMSLNMARTA |

| | | | | | | | |
|---|---|---|---|---|---|---|---|
| | | | | | | | HCIFQHGDGIGTSVGVTKDRLVSLITNPIPI DDDDHHVIHVNNSYTIANYDRPIMK* |
| **Fxa6Ag104161.1** | M/S TPS | ***FvrTPS40*** | *F. virginiana* | b | **95.6** | **HS transcript** | MAMPAPVLEHEHHTTATKQSSFILSNSSS KDVVIDPPVAHQRRLAQYHPTIWGPKLID SFSTRYSHESYASKLEGLKQNVSEYLHVL STKAKGDACFTLKLIDSMQRLGVAYHFG KEFKAAIVTLVSSGTASTTTDLQTVALQF RILRQYGISISSDLFNKFRNDEGSFKDSFN NDVDGLLSLYEASHLGMPGEDVLEEAKS FSSKNLKQSLTTILNDDNLRKRVDQSLET PVHWKMPRIEALDFINMYQRDDSRNLVL LELAKLDYNLVQSVHQNEIKELSRWWRE LDFKSKASFSRDRLMENYLWALGITYEP QFSECRIGLTKFVCILSAIDDMYDVYGFLD ELELFTEAVTQWNLKANEDLPEYMKPIYS AMFTFGNELADQNCGLNTLPLIKKEWEN LCKSYMVEARWFYGGYTPTLEEYLKNA WTSVGGPGAMLHAYLLTQGSQLTEASLE SFNHGSQLIYWASIITRLTDDLGTSKAESE RGDVAKSVHCYMEEKGISEEEAQEYIKG LIFYSWKKINEESAKATIPKSIVNMSLNMA RTAHCIFQHGDGIGTSVGVTKDRLVSLIT NPIPIDDDDHHVIHVNNSYTIANYDRPIMK * |
| Fxa6Ag104162 | M/S TPS | *FaTPS41* | *F. x ananassa* | b | - | FaRR1 Genomic | MALPALVLEHAHHTTATKQSSFILSNSSS KVIDPPVAHQRRLAQYHPTIWGPKLIDSF STRYTHESYASKLEDLKQNVSEDLHVLST KAKGDAWLTLKLIDSMQRLGVAYHFDKEI KAAIVTLVSSSTASTTTDLQTVALQFRILR QYGISISSDLFNQFRNCEGSFKDSLSNDV EGLLSLYEASHLGMPGEDVLEEAKRFCS KNLKQSMAKLKDDNLLKRVAQSLQTPLH WRMPRIEALNFITMYQREDSKNLQALLEL AKLDYNLVQSVHQMEIKELSRQVYMETW WRELDLKSKASFSRDRLMENYMWAMGI CYDPQFLQCRIGLTKFVCILTVIDDMYDVY GFLDELEHFTYAVTQWNMEAKEELPEYM KPIYIAMLEFGNDLADNIFKNTGLNTLPYIK NEWINLCKAYIVEARWFYGGYTPTLEEYL KNAWTSVGGPGAMLHAYLLTQGSQLTE ASLESFNHGSQMIYWASIITRLTDDLGTS KAERERGDVAKSIQCYMDEKGASEEEAC DYIKGLICHSWKKINQESAKTTIPRSIVNLS QNMARAAHCIFEHGDGIGNSIGVTKDRLV SLIANPIPLDD* |

| | | | | | | | |
|---|---|---|---|---|---|---|---|
| **Fxa6Ag104162.1** | M/S TPS | ***FaTPS42*** | *F. x ananassa* | b | **92.7** | **FaRR1 transcript** | MPGPVLEQAHHTTATKQSSFILSNSSSKV<br>IDPPVAHQRRLAQYHPTIWGPKLIDSFST<br>RYNHESYASKLEGLKQNVSEYLHVLSTK<br>AKGDACLTLKLIESMQRLGVAYHFEKEIK<br>AAIVTLVSSGTASTTTDLQTVALQFRILRQ<br>YGISISSDLFNQFRNSEGSFKDSLSNDVE<br>GLLSLYEASHLGMPGEDVLEEAKSFSSK<br>NLKQSMATLKDDNLLKRVAQSLQTPLHW<br>RMPRIEALNFITMYQKEDSKNLQAPLELA<br>KLDYNLVQSIHQMEIKELSRWWRELDLK<br>SKASFSRDRLMENYMWAMGIIYDPQFQQ<br>GRIGLTKFVCILTVIDDMYDVYGYLDELER<br>FTDAVTQWNMEAKEELPEYMKPIYIAMLE<br>FGNDLADNIFKNTGLNTLPYIKNEWINLCK<br>AYMVEARWFYGGYTPTLEVYLKNAWTS<br>VGGPGAVLHAYLLIQGSQLTEASLESFIN<br>GSQLIYWASIITRLTDDLGTSKAERERGD<br>VAKSIQCYMEEKGASEEEACDYIKGLICH<br>SWKKINQESAKTTIPRSIVNLSLNMARAA<br>HCIFEHGDGIGNSIGVTKDRLVSLIANPIPL<br>DD* |
| Fxa6Ag104168 | M/S TPS | *FaTPS43* | *F. x ananassa* | b | - | FaRR1 Genomic | MSTPIMASNGSTATETSYLDVQRRTADY<br>KPSIWSYEFLQSLQDELYEDDRWKSMEE<br>YVKHMIDYGNDAQSVETTLELIDDIQRLG<br>LGHRFEASIVGVLHRIMKNLDQITDEDAP<br>SLHLTALSFRLLRQHGFQVSQDVFKNFT<br>DCDGSFKDCLCKDVKGMLSLYEASFLGF<br>EGETLLDEALAFTSMHLKNLSCRLLDTLG<br>RNVLEQVSHALEMPLHHRMQRLEARWYI<br>EVYNKRQDANQMLLEFAKLDYNVVQQTY<br>QSDLKDLSRWWKDMGLAKKLSFSRDRL<br>MECFFWSVGIVSEPQFSNLRKGITKVSVL<br>ICTIDDVYDVYGTLDELKLFTAVVERWDV<br>NAVEILQDYQLKLSFLALYNTVNEMAYET<br>LKDQGVDVLPYLTKAWTDMCKAFLTEAE<br>WRYNNYTPTFEEYIANAWISVSGVLLLVH<br>TYFLLNQNISDEALECLVNHHDLLRWPSLI<br>FRLSNDLATSERGETATSISCIKRGSSDV<br>CDQASARQFIRNLIEKSWKKMNKDGLLN<br>GASGASSPFTKEFVAAATNLARIAQCIYQ<br>YGDGISAPDKIAKNRILAVLLQPITSQAFG<br>VTSFFVLFTLMTKEQYEDNKWKSMEEYV<br>KHMLGYDNDAQSVETTLEVIDDIQRLGLG<br>HRFEASIVGVLHRIMKNLDQITDEDAPSL<br>HLTALSFRLLRQHGFQVSQDVFKIFTDCD |

| | | | | | | | |
|---|---|---|---|---|---|---|---|
| | | | | | | | GSFKDCLCKDVKGMLSLYEASFLGFEGETLLDEALAFTSMHLKNLSCRLLDTQGIDVLEQVSHALEMPLHHRLQRLEARWYIEAYNKRADANQLLLESAKLDYNAVQQTYQRELSDLSRWWKDMGLAKKLPFAKDRLMECFFWSVGMVYEPQLSNLRKGLTKVGALVSTIDDVYDVYGTFDELQLFTSLVERWDVNAVEILQDYQLKLSFLALYNTVNEMAYETLKDQGVDVLPYLTKAWADLCKAFLREAEWSNNKYTPTFEEYLANAWIMDISAELERGETANSISCLIRGSSSDVCDEESARKYISNLIEKSWKKMNKDGQLFRVSSASSPFTKEFVAAATNLSRIAQCMYQYGDGIGAPDKIVKNRILAVILQPV* |
| Fxa6Ag104224 | M/S TPS | *FaTPS44* | *F. x ananassa* | b | - | FaRR1 Genomic | MSTLNLASSGSIATQTSDLDVKTRTADYKPSIWSYEFLQSLQEEQYEDDRWTLLEEDVKHIIDYDNDAKSVKTTLELIDDIQRLGLGHRFEASITGAIDRIMKNLDQITDEDTPSLHLTALSFRLLRQRGFQVSQDIFKIFTDCDRDGSFTDCLRKDVKGMLSLYEASFLGFEGETLLDEALAFTSTHLKNLSSRLLNTHGRNVLEQVSHALEMPLHHRMQRLEARWYIEVYNKRQDANQMLLEFAKLDYNVVQQTYQSELKDLSRWWKDMGLAKKLSFSRDRLMECFFWSVGILSEPQFSNLRKGITKVSVLICTIDDVYDVYGTLDKLELFTSVVERWDVNAMEILQDYQLELSFLALYNTVNEIAYETLKEQGVDVLPYLTKAWADMCKAFLREAEWRHNEYTPTFEEYIANAWISVSGVVILVHTYFLLNQNISGGALECLENHHDLLRWPSLIFRLSNDLATSEAELERGETATSISCLKRGSSVCDVESARKYISNLIENSWKKMNKDGQLFGANSASSPFTKEFEAAATNLARIAQCIYQYGDGISAPDKIVKNRMLAVILQSV* |
| Fxa6Ag104324 | M/S TPS | *FaTPS45* | *F. x ananassa* | b | - | FaRR1 Genomic | MSTQTSGVKRVTPDYKPSIWNDEFLHSLQIHDCKQDQPYEDNKWKSMEESVTHMIDDAQSVETTLEFIDDIQRLGLGHRLETNITRVLERIMKKLEQKTDEDTPSLHLTALRFRLLRQHGFQLSQDVFKIFTDCDGSFNDRLCKDVKGMLSLYEASFLGFEGETLLDEALAFTSMHLKNLSCRLLDTHGRNVLEQVSHALEMPLHHRMQRLEARWYIEAYNKRADANQVLLEFAKLDYNAVQQTYQRDLKDLSRWWKDMGLAKKLPFAKDRLMECFFWSVGIVFEPQLSNLRKGITKVSALISTVDDIYDVYAT |

| | | | | | | | |
|---|---|---|---|---|---|---|---|
| | | | | | | | YDELEQFTSLVERWDVNAAVETLPDYKL KLCFLALYNTVNEMAYETLKEQGVDVLPY LRKAWADLCKAFLREAEWSNNKYTPTFE EYLANAWISASAVVILVHAYFLLNQNISAE ALGCLENPHDLLRWPSLIFRLTNDLATSE AELERGETANSISCLIRGSSNEVCDEESA RKYISNLIENSWKKMNKDGQLFGASSAS SPFTKEFVAAGTNLSRIAQCIYQYGDGIG APDKIVKNRILAVIMQPV* |
| Fxa6Bg100527 | M/S TPS | *FaTPS46* | *F. x ananassa* | a | - | FaRR1 Genomic | MIIDPSLSSPVPQIAMPEIVRPIANFQPSIW GDQFLNYDSQDIKTEALWQQQVEELKVN VQSEVFTNEGDFALRLKLIDAIQRLGVAY HFEEEIEDALRHIHATYHDKDHFNDDSDL STAALCFRLLRQHGYEISSDIFNKFKDEN GSFKEGLVADGCAMLSLYEAAHLRVHGE NILEEALVFTTTHLESAKSSACYSAQMTE QITQALARPLRTSLERICAKRHMSVYQLE DDAALIKHNETLLKLAKMDFNLVQFLHKK ELCEITRWWKELDFERKLPFARDRKVELF FWIVGVYFEPQYSISRIFMTKVAILLTVMD DIYDAYGTFEELVVFTEAIDRWDVKRIDEL PDYLKIFYYELLNVFREMDTVLAKEGRSY RVCYAIQAVPHELHVRLLDQMKDQARSY FNEARWLHEGRTPSMEEYMSVATVSISY TFLTTISLLGMGDIVTKDSFEWLSNGPKIV RASNIIFRLMDDIVSTKFEKERGHAPSSID CYTKQYGVSEQEAIDVFNKQIVESWKDIN EEFLKPTAVPMPVLMRVLNLTRVADLLYK GEDGFTRVGKMTKDSVAAVIIDSVPL* |
| Fxa6Cg100917 | M/S TPS | *FaTPS47* | *F. x ananassa* | a | - | FaRR1 Genomic | MALHQVLASPAQTLNAAFDVKPRPANYA PCIWGDHFLSYSSMEVDVKFEQRVQELK EEVKKMLMAAVNEPALHMLDIVDNIQRLG LSYRFENEIDTILKHVRDHFYGSKDLGNY GDLYSIALHFRLLRQQGYNVSCDVFNKFK ESNGTFKEYLTSDVVGLLSLYEATHLRVH GEDILEEALIFTRTHLESAAPRLSSPLSKQ VTHALYQPLWKGFSRVEARHYMSVYED YDSHNKTLLTLAKLDFNLMQKVHQKELS DMTRWWKDLDFANKLPFARDRIVETYFW ALATIFEPEYHFARMMVCKYGALTTVQD DVYDVYGSYQELELYTDAIERWDISAIDD QLPQCLKDCYKGLLDSYSAYEEKLAKEG NLYRLEYAKEAIKIQLRGYFQEAKWLKEK YTPTMDEYIPNGLNSSFYPMAITSFVGMG GLVTKDTMDWVLNDPKIVKATALTGRLLN |

| | | | | | | | |
|---|---|---|---|---|---|---|---|
| | | | | | | | DMAGHKFEQDIEQGASSVECYMNQYGV<br>TEEEAKIELTKQKDAAWKDINQEWLDLHS<br>SNSIPKPLLQIILNLARSGEVLYKNEDLYT<br>NAEAGLKGFIVSLLIEPVPV* |
| Fxa6Cg101643 | M/S TPS | *FaTPS48* | *F. x ananassa* | a | - | FaRR1 Genomic | MSTQLASRFVIKRISASLSIQTLKIVNPEIL<br>RRTAIYHPSIWGDRFSNYESQDPMTRSL<br>QQKQVDQLKVVVKREVFRNASSDFSTQ<br>MKSIDAMQRLGIAYHFERDIENALEHAHA<br>ACVHAGDLYNVALRFRLLREHGYNVSSDI<br>FKRFKDANGNFKEGLTSDLHGMLSFYEA<br>THLRIHGEDILEDGLHFTTAHLESSANDV<br>NHPLAAQITQALERPLRKFPERLYARRYI<br>SIFPDDPKSLHPEAVALLLKLAKLDFNLVQ<br>SLHKKELSEIIRWWDELDFERELPFARNR<br>IVELYFWILGVYFEPQYSKARHFLTKVIAFI<br>SVLDDIYDAYGTFEELKMFTEAIQRWPDV<br>NCIDELPDYMKIYYRRLFNLCNEIEIKAQA<br>RLYFAEAQWLHENYTPSMDEYMEVSVR<br>SVGNSLLSTMSLVEMGDIVTKDAFEWLA<br>NNPKILRASNIIFRLMDDLVSGKFEKEREH<br>IASSIDCYMKQYGVSEQQTVDIFNKKVVD<br>SWKDINEEFLRPTAVPVAVLTRVLNLTRV<br>VDLLYKKDDEYTRVGDVMKDGVASLLIHP<br>TLYSVSSSSCDLNDLAGISNLRKLRIILSS<br>PFNYMENLLETAGSILNCIQSLIVYNEVES<br>NGCPEQVTQIVAMCRQLYKLTLIGPTLEL<br>PKELQYTSLTKVLLSQCCLKDDQMTILEK<br>LQNLTTLDLRGEVFGENTKILVFSKGGFP<br>QLKFLYLFSMLQISECMVEKGSMPGLCG<br>LSISNCSSSRWVEICHCPQGNNY* |
| **Fxa6D101605.1** | M/S TPS | ***FvTPS50*** | *F. vesca* | **a** | **96.4** | **UC06 transcript** | MPVQATPAAESQIISKPEVVRRTANFKPS<br>VWGDRFANYAEDIITQTQMQEQVEELKQ<br>VVRKEVFTNAADDSSHQLKLIDEIQRLGV<br>AYHFESEIDQALERIHETYQDIHDGGDLY<br>NVALRFRLLRRHGYNVSCDVFNKFKDTN<br>GDYKKSLVTDLSGMLSFYEAAHLRVHGE<br>KLLEEALVFTTTHLQSASAKSSLLKTQITE<br>AVERPLLKTMERLGARRYMSIYQDEASY<br>SENLLKLAKLDFNVVQCLHKKELSDILRW<br>YKELDFARRMPFARDRIVELFFWIAGIYFE<br>PEYVFGRHILTKLIEITTVMDDMYDAFGTF<br>EELVILTEAIDRWDASCMDQLPDYMQPF<br>YITLLDVIDEVEEELTKQGRSYRIHYAKEI<br>MKNQARLYFAEARWFHEGCTPKMDGYM<br>RVAASSVGNTMLSVVSLVGMGDIITKFEF |

| | | | | | | | |
|---|---|---|---|---|---|---|---|
| | | | | | | | EWLTNEPKILRASNTIFRLMDDIAGYKFEK ERGHVASSIDCYMNEYGVSEQETIDIFNK RIVDSWKDINEEFLRPTAAPVPVLNRVLN LTRVVDLLYKRGDAFTHVGKLMKDCIAA MFIDPVPL* |
| Fxa6Dg100848 | M/S TPS | *FaTPS49* | *F. x ananassa* | a | - | FaRR1 Genomic | MALHQVLASPAQTLNAAFDVKPRPANYA PCIWGDHFLSYSSMEVDVKFEQRVQELK EEVKRMLMAVVNEPASHMLDMVDNIQRL GLSYRFENEIDTILKHVHDHFCGSKYLGK NCDLYTTALHFRLLRQQGYNVSCDVFNK FKESNGTFKEYLTSDVVGLLSLYEATHLR VHGEDILEEALIFTRTHLESAAPRLSSPLS KQVTHALYQPLWKGFSRVEARHYMSVY EDYDSHNKTLLTLAKLDFNLMQKVHQKE LSDMTRWWKDLDFANKLPFARDRIVETY FWALATIFEPEYHFARTMVCKYGALTTVQ DDVYDVYGSYQELELYTDAIERWDISAID DQLPQCLKDCYIGLLDSYSAYEEKLAKEG NLYRLEYAKEAIKIQLRGYFQEAKWLKEK YTPTMDEYIPNGLNSSFYPMAITSFVGMG GLVTKDTMDWVLNDPKIVKATALTGRLLN DMAGHKFEQDIEQGASSVECYMNQYGV TEEEAKIELTKQKDAAWKDINQEWLDLHS SNSIPKPLLQIILNLARSGEVLYKNEDFYT NAEAGLKGFIVSLLIEPVPV* |
| Fxa6Dg101605 | M/S TPS | *FaTPS50* | *F. x ananassa* | a | - | FaRR1 Genomic | MPVQATPAAESQIISKPEVVRRTANFKPS VWGDRFANYAEDIITQAQMQEQVEELKQ VARKEVFTDAADDSSRQLKLIDAIQRLGV AYHFESEIDNALERIHETYQDIHDDDGDL YNVALRFRLLRRHGYNVSCDVFNKFKDT NGDYKKSLVTDVSGMLSFYEAAHLRVHG ETLLEETLVFTTTHLESASAISSLLKTQITE ALERPLLKTMERLGARRYMSIYQDEASH SENLLKLAKLDFNVVQCLHKKELSDILRW YKELDFARRMPFARDRVVELFFWIAGIYF EPEYVFGRHILTKLIEIVTVMDDMYDAFGT FEELVILTEAIDRWDASCMDQLPDYMQPF YITLLDVIDEVEEELTKQGRSYRIHYAKDI MKNQARLYFAEARWFHEGCTPKMDEYM RVAASSVGNTMLSVVSLVGMGDIITKEEF EWLTNEPKILRASNTIFRLMDDIAGYKFEK ERGHVASSIDCYMNEYGVSEQETIDIFNK RIVDSWKDINEEFLRPTAAPVPVLNRVLN LTRVVDLLYKRGDAFTHVGKLMKDCIAA MFIDPVPL* |

| | | | | | | | |
|---|---|---|---|---|---|---|---|
| Fxa6Dg103609 | M/S TPS | *FaTPS51* | *F. x ananassa* | | - | FaRR1 Genomic | MSRLLFFSSSSPSFLFTKLTNTSYLRLVM PGPVLEQAHHTTATKQSSFILSNSSSKVI DPPVAHQRRLAQYHPTIWGPKLIDSFSTR YSHESYASKLEGLKQNVSEYLHVLSTKAK GDACLTLKLIDSMQRLGVAYHFEKEFKAA IVTLVSSGTASTTTDLQTVALQFRILRQYG ISISSDLFNQFRNSEGSFKDSLSNDVEGL LSLYEASHLGMPEEEVLEEAKSFSSKNLK QSMATLKDDNLLKRVAQSLQTPLHWRM PRIEALNFITMYQREDSKNLQALLELAKLD YNLVQSVHQMEIKELSRQVWKRFRFPNP YDQAVDILLQWWRELDLKSKASFSRDRL MENYMWAMGIIYDPQFQQGRIGLTKFVCI LTVIDDMYDVYGYLDELERFTDAVTQWN MEAKEELPEYMKPIYIAMLEFGNDLADNIF KNTGLNTLPYIKNEWINLCKAYMVEARWF YGGYTPTLEEYLKNAWTSVGGPGAVLHA YLLIQGSQLTEASLESFNNGSQLIYWASII TRLTDDLGTSKAERERGDVAKSIQCYME EKGASEEEACDYIKGLICHSWKKINQESA KTTIPRSIVNLSLNMARAAHCIFEHGDGIG NSIGVTKDRLVSLIANPIPLDD* |
| Fxa6Dg103617 | M/S TPS | *FaTPS52* | *F. x ananassa* | b | - | FaRR1 Genomic | MSTPILASNGSTATQASYPDVQRRTTDY KPSIWSYEFLRSLHIDDRGYKEQYEDNK WKSMEEYVKHMLGYDNDAQSVETTLEVI DDIQRLGLGHRFEASIIGTLHRIIKNLDQVT DEDTPSLHLTALSFRLLRQHGFQVSQDV FKIFTDCYGSFKDCLCIDVKGMLSLYEAS FLGFEGETLLNEALAFTSMHLKNLSCRLL DTQGIGVLEHVSHALEMPLHHRMQRLES RWYIEAYNKRADANQVLLESAKLDYNAV QQTYQRDLSDLSRWWKDMGLANKLPFA KDRLMECFFWSVGMVFEPQLSNLRKGLT KVGALVSTIDDVYDVYGTFDELELFTSLV ERWDVNAVEILQDYRLKLSFLALYNTVNE MAYETLKDQGVDVLPYLTKAWADLCKAF LREAEWSNNKYTPTFEEYLANAWISASA VVILVHAYFQLNQNISDEALECLENHHDLL RWPSLIFRLTNDLATSEAELERGETANSI SCLIRGSSSDVCDEESARKYISNLIEKSW KKMNKDGQLFGVSSASSPFTKEFVAAAT NLCRIAQCMYQYGDGIGAPDKIVKNRILA VILQPV* |

| | | | | | | | |
|---|---|---|---|---|---|---|---|
| **Fxa6Dg103617.2** | M/S TPS | ***FcTPS52*** | *F. chiloensis* | **b** | **99.8** | **ILE transcript** | MSTPILASNGSTATQASYPDVQRRTTDY KPSIWSYEFLRSLHIDDRGYKEQYEDNK WKSMEEYVKHMLGYDNDAQSVETTLEVI DDIQRLGLGHRFEASIIGTLHRIIKNLDQVT DEDTPSLHLTALSFRLLRQHGFQVSQDV FKIFTDCYGSFKDCLCIDVKGMLSLYEAS FLGFEGETLLNEALAFTSMHLKNLSCRLL DTQGIGVLEQVSHALEMPLHHRMQRLES RWYIEAYNKRADANQVLLESAKLDYNAV QQTYQRDLSDLSRWWKDMGLANKLPFA KDRLMECFFWSVGMVFEPQLSNLRKGLT KVGALVSTIDDVYDVYGTFDELELFTSLV ERWDVNAVEILQDYRLKLSFLALYNTVNE MAYETLKDQGVDVLPYLTKAWADLCKAF LREAEWSNNKYTPTFEEYLANAWISASA VVILVHAYFQLNQNISDEALECLENHHDLL RWPSLIFRLTNDLATSEAELERGETANSI SCLIRGSSSDVCDEESARKYISNLIEKSW KKMNKDGQLFGVSSASSPFTKEFVAAAT NLCRIAQCMYQYGDGIGAPDKIVKNRILA VILQPV* |
| Fxa6Dg103674 | M/S TPS | *FaTPS53* | *F. x ananassa* | b | - | FaRR1 Genomic | MSTLSLASSGSIATQTSDMDVKTRTAYYK PSIWSYEFLQSLQEELYEDDRWTSLEED VKHMIDYDDVKSVKTTLELIDDIQRLGLGH RFEASITGALDRIMKNLDDQITDEGTPSLH LTALSFRLLRQHGFQVSQDVFKIFTDCDG SFKDCLCKDVKGMLSLYEASFLGFEGET RLDETLAFTSMHLKNLSCRLLDTHGRNVL EQVSHALEMPLHHRMQRLEARWYIEVYN KRQDANQMLLEFAKLDYNVVQQTYQSDL KDLSRWWKDMGLAKKLSFSRDRVMECF FWSVGIVFEPQFSNLRKGITKVSVLICTID DVYDVYGTLDKLELFTSVVERWDVNAME ILQDYQLELSFLALYNTVNEIAYETLKEQG VDVLPYLTKAWADMCKAFLREAEWRHN KYTPTFEEYIANAWISVSGVVILVHIYFLLN QNISGGALECLENHHDLLRWLSLIFRLSN DLATSEAELERGETATSISCLKRGSSVCD VESARKYISNLIDNSWKKMNKDGQLFGA SSASSPFTKEFEAAATNLARIAQCIYQYG DGISAPDKIVIAVILQSV* |
| Fxa7Ag200397 | M/S TPS | *FaTPS54* | *F. x ananassa* | a | - | FaRR1 Genomic | MSTVPAALAPELAHSSTPNITATLDVITRR VANFSPSVWGDHFLSYSSMEAADGKSK KHVHDLKEELKRMILAPAERPSQKLHFIN NIQRLGVSYHFQNEIDKILEQVHREEDDD |

| | | | | | | | |
|---|---|---|---|---|---|---|---|
| | | | | | | | DADLCTTALKFRLLRQQVSRQSISKDDD GKFKESDINDVVGLLSLYEASHLLVPGED ALMEALTFCTTHLESAAHRLIPSSQLSKQ VTHALYQPLWKGMPRIEARHYLSVYQED ESNNETLLNFAKLDFNFVQKVHQKELSEI TRWWKDLDFATKLPFARDRVVEAYFWAL GVYFEPEYYFARMISAKATSIITVIDDIYDV GLWDISAVDQLPDYMKVCYTALLNFFTEI EESLANKGISYRLHYAREGFKVQVRAYF QEAKWFKQKYTPTLEEYMSVERYTSFFM VAIVSFVGLGAIVTEDSMDWVFSEPKILKA TSIIGRLMNDLVGHKQYGATEEEAEVELT KQVNEAWKDINEEWLEATPIPRLLLSLILN FARTSELLYKGEDVYTHSGNVLKGYVAS LFIESVPM* |
| **Fxa7Ag203289.1** | M/S TPS | ***FvrTPS55*** | *F. virginiana* | **a** | **98.5** | **NC transcript** | MSPLVLASQVHEPETGADVKLPPAYYEY APCIWNDHFLSYSTTEVDIKLEQHVRELK EEVKRILMDLANVSGSQQIDLVNDLQRLG VAYHFENEIDEILKQIHHNYSCGTDDDDL YTTALRFRLLRQHGYNISCDIFNKFKDGN HQTFKKSLQSDVSGLLSLYEATHLRVHG EEILEEALAFTTTHLESIKHNLSPPLSNLVS HSLNQPLRTGVARVEARHYLSIYPDCDS HNETLLTLAKLDFNLVQQLHQKELCEITR WWKNLDVKNKLPFARDRVTEIYFIWVLSV YFEPQFSFARRTSCKVTAILSIIDDIYDTQ GTLEELELFTEAIQRWDICAIDPLPDYMKV FYKAMLEVYIEIEEELAKEGNSYRIHYAIE AMKKQATYYFHEAKLLQQKHIPTLDEYM DLALPSTAYHMLITTAFVGMGDIATQDSF DWLATYPRAVKGAEVVCRLMDDIADYKF ELERGTDISSVECYLKEHGATEEEAMTEL RRRLSDAWKDINESFFLPNALPRPLLTRV LNFACAMDVAYKYEDTYTTHHASILKDFI VPTLVESVPL* |
| **Fxa7Ag203289.1** | M/S TPS | ***FaTPS55*** | *F. x ananassa* | **a** | **99.4** | **MM transcript** | MSPLVLASQVHEPETGADVKLPPAYYEY APCIWNDHFLSYSTTEVDIKLEQHVRELK EEVKRILMDLANVSGSQQIDLVNDLQRLG VAYHFENEIDEILKQIHHNYSCGTDDDDL YTTALRFRLLRQHGYNISCDIFNKFKDGN HQTFKASLQSDVSGLLSLYEATHLRVHG EEILEEALAFTTTHLESIKHSLSPPLSNLVS HSLNQPLRTGVARVEARHYLSIYPDCDS HNETLLTLAKLDFNLVQQLHQKELCEITR WWKNLDVKNKLPFVRDRITEVYFIWVLSV |

| | | | | | | | |
|---|---|---|---|---|---|---|---|
| | | | | | | | YFEPQFSFARRTSCKVTAILSIIDDIYDTQ GTLEELELFTEAIQRWDICAIDPLPDYMKV FYKAMLEVYIEIEEELAKEGNSYRIHYAIE AMKKQATYYFHEAKLLQQKHIPTLDEYM DLALPSTAYHMLITTAFVGMGDIATQDSF DWLATYPRAVKGAEVVCRLMDDIADYKF ELERGTDISSVECYMKEHGATEEEAMTE LRRRLSDAWKDINESFFLPNALPRPLLTR VLNFACAMDVAYKYEDSYTTHHSSILKDF IIPTLVESVPL* |
| Fxa7Ag203294 | M/S TPS | *FaTPS56* | *F. x ananassa* | a | - | FaRR1 Genomic | MLNIDRCCPASPSWLGSGPSVRMPGPD PMWASQVHEPETGADVKLPPAYYEYAP CIWNDHFLSYSTTEVDIKLEQHVRELKEE VKRILMDLANVSGSQQIDLVNNLQRLGVA YHFENEIDEILKQIHHNYSCGTDDDDLYTT ALRFRLLRQHGYNISCDIFNKFKDGNHQT FKASLQSDVSGLLSLYEATHLRVHGEEIL EEALAFTTTHLESIKHSLSPPLSNLVSHSL NQPLRTGVARVEARHYLSIYQDCDSHNE TLLTLAKLDFNLVQQLHQKELCEITRWWK NLDVKNKLPFVRDRITEVYFIWVLSVYFE PQFSFARRTSCKVTAILSIIDDIYDTQGTLE ELELFTEAIQRWDICAIDPLPDYMKVFYKA MLEVYIEIEEELAKEGNSYRIHYAIEAMKK QATYYFHEAKLLQQKHIPTLDEYMDLALP STAYHMLITTAFVGMGDIATQDSFDWLAT YPRAVKGAEVVCRLMDDIADYKFELERG TDISSVECYLKEHGATEEEAMTELRRRLS DAWKDINESFFLPNALPRPLLTRVLNFAC AMDVAYKYEDSYTTHHSSEGSFGKVYKG RRKFTGQTVAMKFIMKHGKSDKDIHNLR QEIEILRKLKHENIIEMLDSFESPQEFCVV TEFAQVLISSLSKGELFEILEDDKHLPEEQ VQAIAKQLVRALHYLHSNRIIHRDMKPQNI LIGAGSIVKLCDFGFARAMSTNTVVLRSIK GTPLYMAPELVREQPYNHTADLWSLGVI LYELFVGQPPFFTNSVYALVRHIVKDPVK YPDNMSSSFKNFLKGLLNKVPQNRLTWP ALLEHPFVKETAEELEAREMRSATAAER GCVAAWRGEGNKIQTPGGLAVSSPGIVT STASSEDNSGISFQDDAQSNIPDSKAVNS SPNEEFPGFANPDEVKQSGCQILDRLEN NSRTVKGALIIGQDNEALAHVLLPIKRCSQ GSQNSSRDQGILTSNQSLRILSNLVAVGA ITSSGLLDEIIHEILVYTTFIVSIKSSELNELR |

| | | | | | | | |
|---|---|---|---|---|---|---|---|
| | | | | | | | AKSFSIIKVLVDNVGHDIGSSYFRHWVAL<br>AEIFSQVVGCSEDASGRVMLESIACITAM<br>LTRVNEGLKVLFSTPARQEVCGPNEAVK<br>QILDHAKTSGLVDQLCLCLATAGASLISG<br>SSNMLRSACEACRAIWLLVDASEFFSTK<br>GNVVSFPLNTMASPARDQDRGGSVILIES<br>AKLVAVVTRALVRSKPVQVAIHYCLHQRL<br>EASLYAGIQLLLRCCLQSGIVPGILCGLPS<br>SLPVTTVVSGGGDRSIISEIFSLLSLCISSQ<br>NKDPQAIETTNLKSKISDPNTLVMHSCLIL<br>ASVAQCLKATGRNSALFMLTTSSKNQLS<br>RLSVLAHYFSSGESTNTSSRAHSASAML<br>ALASILSLESGASVGSSVFEVAVPLIPQTT<br>TLCEYLKLPSNSEIVSGPNGTNGVLSNW<br>HGLRDGCVGLLEARLRWGGPSAVQQMC<br>ASNIPLLLINLLAKNQQYSSPEEESSASDQ<br>VGLSPIGVVWTVSSICQCLSGGALTFRQI<br>LLRSDHIKLFSDLISDTHLKLVKSWAGPG<br>GGMDGVRDITNAIIDLLAFPFVAVQNAPG<br>LPAATASVNSGILLNMGSPGVKVGMEDR<br>DMVKVIEEDLGKYIKILLEVGVPGIILWCLE<br>HLELKDLGRPVAFLAKMIAQRPLAVQLVG<br>KGLLDPTKMRRLLDRSSPKEVILDGLMIV<br>SDLARMDKGFYEYINRASLLEFFKGFLVH<br>EDPSVRSKACSALGNMCRHSSYFYSSLA<br>RNQIIGLLIDRCSDPDKRTRKFACFAIGNA<br>AYHNDMLYEELRRSIPKLANLLLSSEEDK<br>TKANAAGALSNLIRNSNKLCEDIVSKGAM<br>QSLLKLVAECSELALNPSRRDSAHESPLK<br>IALFSLAKMCSHPPCRDFLRSSDLFPVIG<br>RLRQSPESTIANYASAIINKVA* |
| Fxa7Bg200423 | M/S TPS | *FaTPS57* | *F. x ananassa* | a | - | FaRR1 Genomic | MSTVPAALASELAYSSTTNITATLDVITRR<br>VANFSPSVWGDHFLSYSSMEAADGKSK<br>KHVHDLKEELKRMILAPAERPSQKLHFIN<br>NIQRLGVSYHFQNEIDKILEQVHREEDDD<br>DADLCTTALKFRLLRQQGYNASCNMFNK<br>FKDDDGKFKESHINDVVGLLSLYEASHLL<br>VPGEDALMEALTFCTTHLESAAHRLSSSS<br>QLSKQVTHALYQPLWKGMPRIEARHYLS<br>VYQEDDSNNETLLNFAKLDFNFVQKVHQ<br>KELSEITRWWKNLDFATKLPFARDRVVE<br>AYFWALGVYFEPEYYFARMISAKATSIITV<br>IDDIYDVHGTYEELESFTEAIERWDISAVD<br>QLPDYMKVCYTVLLNFFTKLEESLANKGI<br>SYRLHYAREGFKVQVRAYFQEAKWFKQ |

| | | | | | | | |
|---|---|---|---|---|---|---|---|
| | | | | | | | KYTPTLEEYMSVERDTSFFMVAVVSFVG<br>LGAIVTEDSMDWVFSEPKILKATSIIGRLM<br>NDLVGHKFEQKREHNASAVECYMKQYG<br>ATEEEAEVELTKQVNEAWKDINKEWLEA<br>TPIPRLLLSLILNFARTSELLYKGEDVYTH<br>SGNVLKGYVASLFIESVPI* |
| **Fxa7Bg203106** | M/S TPS | ***FcTPS58*** | *F. chiloensis* | **a** | **96.1** | **Amb transcript** | MSPLVLASQVHEPETGADVKLPLAYYEY<br>APCIWDDHFLSYSTTEVDIKLEQHVRELK<br>EEVKRILKDPANVSGSQQIHLVNDLQRLG<br>VAYHFENEIDEILKQIHHNYSCGTADDDL<br>YTTALRFRLLRQHGYNISCDIFNKFKDEN<br>HQTFKASLQSDVSGLLSLYEATHLRVHG<br>EEILEEALAFTTTHLESIKHNLSPPLSNLVS<br>HSLNQPLRTGVARVEARHYLSIYQECDS<br>HNETLLTLAKLDFNLVQQLHQKELCEITR<br>WWKNLDLKNNIPFARDRITEVYFIWVLSV<br>YFEPQFSFARRTSCKVTAILSIIDDIYDSQ<br>GTLEELELLTEAIQRWDICAIDPLPDYMKV<br>FYKAMLEVYIEIEEELAKEGNLYRIHYAIEA<br>MKKQATYYFHEAKLLQQKHIPTLDEYMDL<br>ALPSTAYHMLITTAFVGMGDIATQDSFDW<br>LATYPRAVKGAEVVCRLMDDIADYKFELE<br>RGTDISSVECYMKEHGATEEEAMTELRR<br>RLSDAWKDINESFFLPNALPRPLLTRVLN<br>FACAMDVAYKYEDSYTTHHSSILKDFIIPT<br>LVESVPL* |
| Fxa7Bg203106 | M/S TPS | *FaTPS58* | *F. x ananassa* | a | - | FaRR1 Genomic | MSLLVLASQVHEPDTGADVKLPLAYYEY<br>APCIWDDHFLSYSTTEVDIKLKQHVRELK<br>EEVKRILMDHANVSGSQQIDLVNDLQRL<br>GVAYHFENEIDEILKQIHHNYSCGTADDD<br>LYTTALRFRLLRQHGYNISCDIFNKFKDG<br>NHQTFKKSLQSDVSGLLSLYEATHLRVH<br>GEEILEEALAFTTTHLESIKHNLSPPLSNL<br>VSHSLNQPLRTGVARVEARHYLSIYQEC<br>HSHNETLLTLAKLDFNLVQQLHQKELCEI<br>TRWWKNLDVKNKLPFARDRVTEVYFIWV<br>LSVYFEPQFSFARRTSCKVTAILSIIDDIYD<br>SQGTLEELELLTEAIQRWDICAIDPLPDYM<br>KVFYKAMLEVYIEIEEELAKEGNLYRIHYAI<br>EAVSPIYLMKKQATYYFHEAKLLQQKHIP<br>TLDEYMDLALPSTAYHMLITTAFVGMGDI<br>ATQDSFDWLATYPRAVKGAEVVCRLMD<br>DIADYKFELERGTDISSVECYMKEHGATE<br>EEAMTELRRRLSDAWKDINESFFLPNALP |

| | | | | | | | |
|---|---|---|---|---|---|---|---|
| | | | | | | | RPLLTRVLNFACAMDVAYKYEDTYTTHH<br>ASILKDFIVPTLVESVPL* |
| Fxa7Cg100355 | M/S TPS | *FaTPS59* | *F. x ananassa* | a | - | FaRR1 Genomic | MSTVPATVDVITRRVANFSPSVWGDHFL<br>SYSSMEAADGKSKKHVHNLKEELKRMIL<br>APAERPSQKLHFINNIQRLGVSYHFQNEI<br>DKILEQVHREEDDDDADLCTTALKFRFLR<br>QEGYNASCNMFNKFKDDDGKFKESHIND<br>IVGLLSLYEASHLLVPGEDALMEALTFCTT<br>HLESATHRLSSSSQLSKQVTHALYQPLW<br>KGMPRIEARHYLSVYQEDDSNNETLLNF<br>AKLDFNFVQKVHQKELSEITRMISAKATSI<br>ITVIDDIYDVHGTYEELESFTEAIERWDISA<br>VDQLPDYMKVCYTALLNFFTELEESLANK<br>GISYRLHYAREGFKVQVRAYFQEAKWFK<br>QKYTPTLEEYMSVERDTSFFIVAVVSFVG<br>LGSIVTEDSMDWVFSEPKISKATSIIGRLM<br>NDLVGHKVERKREHNASAVECYMKEYG<br>ATEEEAEVELTKQVNEAWKDINEEWLEA<br>TPIPRLLLSLILNFARTSELLYKGEDVYTH<br>SGNVLKGYVASLFIESVPI* |
| Fxa7Cg101251 | M/S TPS | *FaTPS60* | *F. x ananassa* | a | - | FaRR1 Genomic | MAASVALARVEFGRISTTSPHLDLKLGIR<br>RMPSQLSSESEALSPKATADVKRRSTNF<br>KLSIWGDYFLSYASMEVDSTLEQKVQQL<br>KEEVKRMLVDNVKDGSQKLSLIDDIQRLG<br>VSYHFENEICEILQQIYDNDHVSYDDLHQ<br>NNSDLHAVSLRFRLLRQQGYNASCVKYF<br>SNLKDNDGKFKESLVDDTKGLLGLYEAT<br>HLRIHGDDILDEALTFTTTHLESATRRGLS<br>SPLSKQVTHALNQPFWKGMPRIEARHYI<br>STYQEDKYSHNETLLNFARLDFNLLQQLH<br>QKKLCELTRWWKDLDFVKKLPFTRNRLV<br>ECYFIALGIFFEPEYYFGRRTFCKVIYMLS<br>AIDDIYDVHGTLEELDSSMKLFRGLNQLIP<br>YHVGHLFPGSVAGLYESLCFQAQLDVFT<br>EIEEKLASGGNVYAIHHARESLKMTVAGY<br>MKEARWFYREHAASAVELYMKEHGATE<br>EEAIVELTNQVTDAWKVIIKACLHPTPFPL<br>PDLMRPLRVASTWEVLYKSADSYTSPGA<br>EFKGIVTSLLIEPVPNALA* |
| Fxa7Cg103018 | M/S TPS | *FaTPS61* | *F. x ananassa* | a | - | FaRR1 Genomic | MSPQVLASQVQEPDTGADVKLPLAYYEY<br>APCIWDDHFLSYSTTEVDIKLEQHVRELK<br>EEVKRILMDPANVSGSQQIDLVNDLQRLG<br>VAYHFENEIDEILKQIHHNYSCGTDDDDL<br>YTTALRFRLLRQHGYNISCDARSSKELPT |

| | | | | | | | |
|---|---|---|---|---|---|---|---|
| | | | | | | | DADIFNKFKDGNHQTFKKSLQSDVSGLLS<br>LYEATHLRVHGEEILEEALAFTTTHLESIK<br>HNLSPPLSNLVSHSLNQPLRTGVARVEA<br>RHYLSIYQECDSHNETLLTLAKLDFNLVQ<br>QLHQKELCEITRWWKNLDLKNNIPFARD<br>RITEVYFIWVLSVYFEPQFSFARRTSCKF<br>SFARRTSCKVTAILSIIDDIYDSKGTLEELE<br>LLTEAIQRWDICAIDPLPDYMKVFYKAML<br>EVYIEIEEELAKEGNLYRIHYAIEAMKKQA<br>TYYFDEAKLLQQKHIPTLDEYMDLALPST<br>AYHMLITTAFVGMGDIATQDSFDWLATYP<br>RAVKGAEVVGRLMDDIADYKFELERGTNI<br>SSVECYMKEHGATEEEAMTELRRRLSDA<br>WKDINESFFLPNALPWPLLTRVLNFACAM<br>DVAYKYEDGYTTHHASILEDFIVPTLVESV<br>PL* |
| Fxa7Dg100343 | M/S TPS | *FaTPS62* | *F. x ananassa* | a | - | FaRR1 Genomic | MSTVPAALASELAHSSSPNITATLDVITRR<br>VANFSPSVWGDHFLSYSSMEAADGKSK<br>KHVHDLKEELKRMILAPAERPSQKLHFIN<br>NIQRLGVSYHFQNEIDKILEQVHREEDDD<br>DDADLCTTALKFRVLRQEGYNASCNMFN<br>KFKDDDGKFKESHINDVVVLLSLYEASHL<br>LVPGEDALMEALTFCTTHLESATHRLSSS<br>SQLSKQVTHALYQPLWKGMPRIEARHYL<br>SVYQEDDSNNETLLNFAKLDFNFVQKVH<br>QKEQSEITRWWKDLDFATKLPFARDRVV<br>EAYFWALGVYFEPEYYFARMISAKPTSIIT<br>VIDDIYDVHGTYEELESFTEAIERWDISAV<br>DQLPDYMKVCYTALLNFFTELEESLANKG<br>ISYRLHYAREGFKVQVRAYFQEAKWFKQ<br>KYTPTLEEYMSVERDTSFFMVAVVSFVG<br>FGAIVTEDSMDWVFSEPKILKATSIIGRLM<br>NDLVGYKFEQKREHNASAVECYMKQYG<br>ATEEEAEVELTKQVNEAWKDINEEWLEA<br>TPIPRLLLSLILNFARTSELLYKGEDVYTH<br>SGNVLKGYVASLFIESVPI* |
| Fxa7Dg102096 | M/S TPS | *FaTPS63* | *F. x ananassa* | a | - | FaRR1 Genomic | MPSQLSSESEALSPKATAVVKRRSTNFK<br>PSIWGDYFLSYASMEVDITLEQQVQQLKE<br>EVKRMLVDNVKDGSQKLSLIDDIQRLGVS<br>YHFENEICEILQQIYDNDHVSYNDLYQNN<br>SDLHAVSLRFRLLRQQGYNASCDKYFSN<br>LKDNVGKFKESLVDDTKGLLGLYEATHLR<br>IHGDDILDEALPFTTTHLQSATRRDLSSPL<br>SKQVTHALNQPFWKGMPRIEARHYISTY<br>QEDKYSHNETLLNFARLDFNLLQQLHQK |

| | | | | | | | |
|---|---|---|---|---|---|---|---|
| | | | | | | | ELCELTSWCKDLDFVKKLPFTRNRLVEC YFIALAIYFEPAYYFGRTFCKVIYMLSAIDD IYDVHGTLEELELFNEAFLRWDISALDQS SDYMKVCFQAQLDVFTEIEEKLASEGNLY AIHHPRESLKMTVAGYMKEARWFYSKYT PTFDEYMPLGTLTSSYYLSITAVVGMGLA ATKDSLDWLFTDPGILNATSVIGRLLNDIR THQVFLARC* |
| Fxa7Dg102851 | M/S TPS | *FxaTPS64* | *F. x ananassa* | a | - | FaRR1 Genomic | MSPQILASQVQEPDTGADVKLPLAYYEY APGIWDDHFLSYSTTEVDIKLEQHVRELK EEVKRILMDPANVSGSQQIDLVNDIQRLG VAYHFENEIDEILKQIHHNYSCGTDDDDL YTTALRFRLLRQHGYNISCDIFNKFKDGN HQTFKKSLQSDVSGLLSLYEATHLRVHG EEILEEALAFTTTHLESIKHNLSPPLSNLVS HSLNQPLRTGVSRVEARHYLSIYQECDS HNETLLTLAKLDFNLVQQLHQKELCEITR WWKNLDLKNKIPFARDRITEVYFIWVLSV YFEPQFSFARRTSCKVTAILSIIDDIYDTQ GTLEELELFTEAIQRWDICAIDPLPDYMKF FYKAMLEVYIEIEEELAKEGNSYRIHYAIE AVSPIYLMKKQATYYFHEAKLLQQKHIPTL DEYMDLALPSTAYHMLITTAFVGMGDIAT QDSFDWLATYPRAVKGAEVVCRLMDDIA DYKFELERGTDISSVECYLKEHGATEEEA MTELRRRLSDAWKDINESFFLPNALPRPL LTRVLNFACAMDVAYKYEDSYTTHHSSIL KDFIIPTLVESVPL* |

**Supplemental Table 5:** Strawberry genotypes used in this study. Modern cultivars 'Mara Des Bois' (MDB) and 'Royal Royce' were also grown under greenhouse conditions for a developmental time course study. No MDB fruit could be harvested in the field for RNA sequencing.

| **Species** | **Type** | **Code** | **Accessions** (Identifier) Release Date & Location |
|---|---|---|---|
| *F. vesca (Fv)* | Diploid | UC04<br>UC06 | **UC04** (PI 551598) California, USA<br>**UC06** (PI 551514) California, USA |
| *F. virginiana (Fvr)* | North American Octoploid | NC<br>HS | **NC_96-35-2** (PI 612323) Alabama, USA<br>**Harris Springs** (17X004P001) California, USA |
| *F. chiloensis (Fc)* | South American Octoploid | ILE<br>Amb | **Isle De Lemuy (02A White)** (PI 552038) Chile<br>**Ambato** (PI 551736) Ecuador |
| *F.* × *ananassa (Fa)* | Breeding Cultivar Octoploid | MM<br>DPW<br>EM<br>Head<br>Linn<br>Prim<br>Tan<br>MDUS<br>BB<br>MDB<br>17<br>RR | **Madame Moutot** (PI 551632) 1910 France<br>**Direktor Paul Wallbaum** (PI 551436) 1953 Germany<br>**EarliMiss** (PI 551862) 1955 Mississippi, USA<br>**Headliner** (PI 551652) 1957 Louisiana, USA<br>**Linn** (PI 551500) 1967 Oregon, USA<br>**Primella** (PI 551422) 1969 Netherlands<br>**Tangi** (PI 551481) 1975 Louisiana, USA<br>**MDUS 5130** (PI 551946) 1981 Maryland, USA<br>**Beaver Belle** (PI 551839) 1989 Canada<br>***Mara Des Bois** (17Z001P001) 1991 France<br>**17C224P011** [2017] California, USA<br>***Royal Royce** (08C123P001) 2019 California, USA |

**Supplemental Table 6:** Amino acid sequence similarity matrices of identified/tested TPS and syntenic pseudogenes from synteny analysis.

| TPS Clade C | FvrCPS5 | Fxa2Cg203098.1 | FaCPS1 | FaCPS2 | FaCPS3 | FaCPS4 | FaCPS5 | FaCPS6 | FaCPS7 |
|---|---|---|---|---|---|---|---|---|---|
| FvrCPS5 | | 10.9 | 69.2 | 69.2 | 69.2 | 69 | 98.5 | 54.7 | 78.5 |
| Fxa2Cg203098.1 | 10.9 | | 9.6 | 9.8 | 9.9 | 9.8 | 10.9 | 16.3 | 11.6 |
| FaCPS1 | 69.2 | 9.6 | | 96.7 | 96.2 | 96.1 | 69.4 | 51.6 | 71 |
| FaCPS2 | 69.2 | 9.8 | 96.7 | | 98.3 | 98.2 | 69.6 | 51.7 | 71.3 |
| FaCPS3 | 69.2 | 9.9 | 96.2 | 98.3 | | 98 | 69.6 | 51.7 | 71.7 |
| FaCPS4 | 69 | 9.8 | 96.1 | 98.2 | 98 | | 69.2 | 51.7 | 71.3 |
| FaCPS5 | 98.5 | 10.9 | 69.4 | 69.6 | 69.6 | 69.2 | | 55 | 78.4 |
| FaCPS6 | 54.7 | 16.3 | 51.6 | 51.7 | 51.7 | 51.7 | 55 | | 54.8 |
| FaCPS7 | 78.5 | 11.6 | 71 | 71.3 | 71.7 | 71.3 | 78.4 | 54.8 | |

| TPS Clade e/f | FcKSL5 | Fxa3Dg201124.1 | FaKSL1 | FaKSL2 | FaKSL3 | FaKSL4 |
|---|---|---|---|---|---|---|
| FcKSL5 | | 21.9 | 51.6 | 52.1 | 48.8 | 59 |
| Fxa3Dg201124.1 | 21.9 | | 40.7 | 40.6 | 38.8 | 31.8 |
| FaKSL1 | 51.6 | 40.7 | | 97.4 | 93 | 73.9 |
| FaKSL2 | 52.1 | 40.6 | 97.4 | | 91.3 | 73.8 |
| FaKSL3 | 48.8 | 38.8 | 93 | 91.3 | | 70.6 |
| FaKSL4 | 59 | 31.8 | 73.9 | 73.8 | 70.6 | |

| TPS Clade g | FcTPS18 | FvNES1 | FvrTPS17 | Fxa3Cg100266.1 | FaNES1 | FaNES2 | FaTPS8 | FaTPS11 | FaTPS12 | FaTPS13 | FaTPS14 | FaTPS17 | FaTPS18 | FaTPS 22 |
|---|---|---|---|---|---|---|---|---|---|---|---|---|---|---|
| FcTPS18 | | 93.6 | 82.1 | 23.4 | 88.8 | 93.6 | 83.4 | 92.7 | 94 | 69.2 | 83.4 | 88.8 | 96.2 | 81.6 |
| FvNES1 | 93.6 | | 80.5 | 22.6 | 84.2 | 93.8 | 83.4 | 92.8 | 98.1 | 68.9 | 83.4 | 84.2 | 92.8 | 81.8 |
| FvrTPS17 | 82.1 | 80.5 | | 19.7 | 73.1 | 82.6 | 70.6 | 81.5 | 80.9 | 75 | 70.4 | 73.1 | 81.1 | 68.4 |
| Fxa3Cg10026 6.1 | 23.4 | 22.6 | 19.7 | | 21 | 22.6 | 20.1 | 22.6 | 22.7 | 17.6 | 20.1 | 21 | 24.1 | 19.7 |
| FaNES1 | 88.8 | 84.2 | 73.1 | 21 | | 84.9 | 93.9 | 84.2 | 84.7 | 76.5 | 93.9 | 100 | 85.9 | 91.2 |
| FaNES2 | 93.6 | 93.8 | 82.6 | 22.6 | 84.9 | | 83.8 | 96.1 | 94.7 | 69.1 | 83.8 | 84.9 | 92.9 | 81.7 |
| FaTPS8 | 83.4 | 83.4 | 70.6 | 20.1 | 93.9 | 83.8 | | 83.4 | 83.3 | 75 | 98.9 | 93.9 | 82 | 91 |
| FaTPS11 | 92.7 | 92.8 | 81.5 | 22.6 | 84.2 | 96.1 | 83.4 | | 93.3 | 68.4 | 83.5 | 84.2 | 91.7 | 81.2 |
| FaTPS12 | 94 | 98.1 | 80.9 | 22.7 | 84.7 | 94.7 | 83.3 | 93.3 | | 69.3 | 83.4 | 84.7 | 93.5 | 82.3 |
| FaTPS13 | 69.2 | 68.9 | 75 | 17.6 | 76.5 | 69.1 | 75 | 68.4 | 69.3 | | 75 | 76.5 | 69.2 | 74 |
| FaTPS14 | 83.4 | 83.4 | 70.4 | 20.1 | 93.9 | 83.8 | 98.9 | 83.5 | 83.4 | 75 | | 93.9 | 82.6 | 91.8 |
| FaTPS17 | 88.8 | 84.2 | 73.1 | 21 | 100 | 84.9 | 93.9 | 84.2 | 84.7 | 76.5 | 93.9 | | 85.9 | 91.2 |
| FaTPS18 | 96.2 | 92.8 | 81.1 | 24.1 | 85.9 | 92.9 | 82 | 91.7 | 93.5 | 69.2 | 82.6 | 85.9 | | 81.4 |
| FaTPS22 | 81.6 | 81.8 | 68.4 | 19.7 | 91.2 | 81.7 | 91 | 81.2 | 82.3 | 74 | 91.8 | 91.2 | 81.4 | |

| TPS Clade b, 1 | FaTPS51 | FaTPS43 | FaTPS42 | FaTPS41 | FaTPS40 | FaTPS18 | FaTPS17 | Fxa6Cg103709.1 | Fxa6Bg103821.1 | FvrTPS40 | FcTPS40 |
|---|---|---|---|---|---|---|---|---|---|---|---|
| **FaTPS51** | | 20.1 | 89.9 | 87.5 | 75.7 | 31.4 | 30.1 | 41.1 | 60 | 75.7 | 77 |
| **FaTPS43** | 20.1 | | 20.5 | 20.2 | 20 | 16.5 | 16.3 | 9.5 | 15.6 | 20.5 | 20.6 |
| **FaTPS42** | 89.9 | 20.5 | | 93 | 76.6 | 33.7 | 32.6 | 40.9 | 64.8 | 80.8 | 82.9 |
| **FaTPS41** | 87.5 | 20.2 | 93 | | 76.2 | 34 | 32.2 | 40.8 | 63.4 | 80.5 | 81.7 |
| **FaTPS40** | 75.7 | 20 | 76.6 | 76.2 | | 31.9 | 30.2 | 42.3 | 51.7 | 83.2 | 84.3 |
| **FaTPS18** | 31.4 | 16.5 | 33.7 | 34 | 31.9 | | 86.4 | 16.8 | 23.7 | 33.3 | 33.3 |
| **FaTPS17** | 30.1 | 16.3 | 32.6 | 32.2 | 30.2 | 86.4 | | 14.9 | 26.5 | 31.8 | 31.9 |
| **Fxa6Cg103709.1** | 41.1 | 9.5 | 40.9 | 40.8 | 42.3 | 16.8 | 14.9 | | 13.1 | 40.8 | 40.9 |
| **Fxa6Bg103821.1** | 60 | 15.6 | 64.8 | 63.4 | 51.7 | 23.7 | 26.5 | 13.1 | | 52.6 | 53 |
| **FvrTPS40** | 75.7 | 20.5 | 80.8 | 80.5 | 83.2 | 33.3 | 31.8 | 40.8 | 52.6 | | 97.2 |
| **FcTPS40** | 77 | 20.6 | 82.9 | 81.7 | 84.3 | 33.3 | 31.9 | 40.9 | 53 | 97.2 | |

| TPS Clade B, 2 | FaTPS21 | FaTPS10 | FaTPS7 | FaTPS6 | FvTPS6 |
|---|---|---|---|---|---|
| **FaTPS21** | | 88.7 | 95.2 | 92 | 92.1 |
| **FaTPS10** | 88.7 | | 92 | 92.6 | 92.5 |
| **FaTPS7** | 95.2 | 92 | | 95.2 | 95 |
| **FaTPS6** | 92 | 92.6 | 95.2 | | 97.9 |
| **FvTPS6** | 92.1 | 92.5 | 95 | 97.9 | |

| TPS Clade b, 3 | FaTPS53 | FaTPS52 | FaTPS45 | FaTPS44 | FaTPS43 | FaTPS10 | Fxa6Cg103766.1 | Fxa6Bg103826.1 | FcTPS52 |
|---|---|---|---|---|---|---|---|---|---|
| **FaTPS53** | | 81.5 | 79.8 | 92.5 | 46.9 | 37.4 | 62.8 | 38.9 | 80.7 |
| **FaTPS52** | 81.5 | | 84.3 | 80.2 | 46.5 | 37.5 | 50.7 | 42.3 | 98.8 |
| **FaTPS45** | 79.8 | 84.3 | | 78.2 | 43.8 | 37 | 49 | 40.2 | 83.5 |
| **FaTPS44** | 92.5 | 80.2 | 78.2 | | 46.8 | 36.3 | 59.2 | 37.4 | 80 |
| **FaTPS43** | 46.9 | 46.5 | 43.8 | 46.8 | | 20.2 | 30.6 | 20.8 | 46.4 |
| **FaTPS10** | 37.4 | 37.5 | 37 | 36.3 | 20.2 | | 24.6 | 18.2 | 37.4 |
| **Fxa6Cg103766.1** | 62.8 | 50.7 | 49 | 59.2 | 30.6 | 24.6 | | 60.9 | 50.9 |
| **Fxa6Bg103826.1** | 38.9 | 42.3 | 40.2 | 37.4 | 20.8 | 18.2 | 60.9 | | 42.5 |
| **FcTPS52** | 80.7 | 98.8 | 83.5 | 80 | 46.4 | 37.4 | 50.9 | 42.5 | |

| TPS Clade a1, Pinene | FaTPS50 | FaTPS48 | FaTPS46 | FaTPS39 | FaTPS37 | FaTPS23 | FaTPS19 | FaTPS9 | FaTPS5 | FaTPS4 | FaTPS3 | FaTPS1 | Fxa3Bg201803.1 | Fxa2Ag100915.1 | FvrTPS5 | FvTPS50 | FvTPS37 | FvPINS |
|---|---|---|---|---|---|---|---|---|---|---|---|---|---|---|---|---|---|---|
| **FaTPS50** | | 42.7 | 60.4 | 60.5 | 60.4 | 55.1 | 29.5 | 27.5 | 57 | 60.1 | 54.5 | 95.4 | 32.5 | 46.6 | 53.1 | 94 | 61 | 93.4 |
| **FaTPS48** | 42.7 | | 42.4 | 65.5 | 42.8 | 37.8 | 28.9 | 27.1 | 43.5 | 42.4 | 38.7 | 42.7 | 25.3 | 32.1 | 37.6 | 42.5 | 43.1 | 42.5 |
| **FaTPS46** | 60.4 | 42.4 | | 59.1 | 94 | 53.9 | 29.3 | 27.8 | 86.5 | 94.9 | 53.8 | 61 | 32.8 | 67.4 | 81.4 | 59.8 | 91.6 | 59.6 |
| **FaTPS39** | 60.5 | 65.5 | 59.1 | | 59.8 | 53.7 | 28.8 | 26.9 | 55.7 | 59.1 | 54.4 | 60.7 | 33.6 | 45.4 | 52.1 | 60.2 | 59.9 | 60.1 |
| **FaTPS37** | 60.4 | 42.8 | 94 | 59.8 | | 53.8 | 29.1 | 27.6 | 87.4 | 93.3 | 53.9 | 60.9 | 32.9 | 67.1 | 82.2 | 59.7 | 95.2 | 59.6 |
| **FaTPS23** | 55.1 | 37.8 | 53.9 | 53.7 | 53.8 | | 42.9 | 39.5 | 50.6 | 53.2 | 91.2 | 55.7 | 55.8 | 42.5 | 46.6 | 54.6 | 54.1 | 54.4 |
| **FaTPS19** | 29.5 | 28.9 | 29.3 | 28.8 | 29.1 | 42.9 | | 87.1 | 29.6 | 28.9 | 42.1 | 30 | 27.3 | 22.7 | 25.5 | 29.1 | 29.3 | 29.2 |
| **FaTPS9** | 27.5 | 27.1 | 27.8 | 26.9 | 27.6 | 39.5 | 87.1 | | 28.1 | 27.4 | 39 | 27.8 | 25.5 | 21.2 | 24 | 27.1 | 27.7 | 27.1 |
| **FaTPS5** | 57 | 43.5 | 86.5 | 55.7 | 87.4 | 50.6 | 29.6 | 28.1 | | 89.8 | 50.9 | 58 | 31 | 65 | 80.3 | 57.1 | 87.8 | 56.6 |
| **FaTPS4** | 60.1 | 42.4 | 94.9 | 59.1 | 93.3 | 53.2 | 28.9 | 27.4 | 89.8 | | 53.5 | 61.3 | 32.7 | 68 | 84.5 | 60.1 | 92.1 | 59.6 |
| **FaTPS3** | 54.5 | 38.7 | 53.8 | 54.4 | 53.9 | 91.2 | 42.1 | 39 | 50.9 | 53.5 | | 55.2 | 54.4 | 41.9 | 47.5 | 54.4 | 55 | 54.3 |

| | | | | | | | | | | | | | | | | | | |
|---|---|---|---|---|---|---|---|---|---|---|---|---|---|---|---|---|---|---|
| **FaTPS1** | 95.4 | 42.7 | 61 | 60.7 | 60.9 | 55.7 | 30 | 27.8 | 58 | 61.3 | 55.2 | | 32.9 | 47.2 | 53.5 | 94.3 | 61.7 | 93 |
| **Fxa3Bg201803.1** | 32.5 | 25.3 | 32.8 | 33.6 | 32.9 | 55.8 | 27.3 | 25.5 | 31 | 32.7 | 54.4 | 32.9 | | 21.2 | 37.9 | 32.2 | 33.2 | 32.2 |
| **Fxa2Ag100915.1** | 46.6 | 32.1 | 67.4 | 45.4 | 67.1 | 42.5 | 22.7 | 21.2 | 65 | 68 | 41.9 | 47.2 | 21.2 | | 58.1 | 46.7 | 69.6 | 46.2 |
| **FvrTPS5** | 53.1 | 37.6 | 81.4 | 52.1 | 82.2 | 46.6 | 25.5 | 24 | 80.3 | 84.5 | 47.5 | 53.5 | 37.9 | 58.1 | | 52.5 | 83.3 | 52.3 |
| **FvTPS50** | 94 | 42.5 | 59.8 | 60.2 | 59.7 | 54.6 | 29.1 | 27.1 | 57.1 | 60.1 | 54.4 | 94.3 | 32.2 | 46.7 | 52.5 | | 60.8 | 98.7 |
| **FvTPS37** | 61 | 43.1 | 91.6 | 59.9 | 95.2 | 54.1 | 29.3 | 27.7 | 87.8 | 92.1 | 55 | 61.7 | 33.2 | 69.6 | 83.3 | 60.8 | | 60.8 |
| **FvPINS** | 93.4 | 42.5 | 59.6 | 60.1 | 59.6 | 54.4 | 29.2 | 27.1 | 56.6 | 59.6 | 54.3 | 93 | 32.2 | 46.2 | 52.3 | 98.7 | 60.8 | |

| **TPS Clade a1, 2** | **FaTPS63** | **FaTPS60** | **FaTPS36** | **FaTPS35** | **FaTPS34** | **FxTPS33** | **FaTPS31** | **FaTPS29** | **FaTPS27** | **FaTPS16** | **FvTPS35** |
|---|---|---|---|---|---|---|---|---|---|---|---|
| **FaTPS63** | | 59.4 | 42.7 | 43 | 42.1 | 38.1 | 42.3 | 41.7 | 42 | 60.6 | 42.7 |
| **FaTPS60** | 59.4 | | 39.5 | 39.7 | 39.4 | 34.9 | 39.2 | 38.2 | 37.7 | 57.8 | 39.6 |
| **FaTPS36** | 42.7 | 39.5 | | 98.4 | 93 | 56.5 | 91.8 | 87.4 | 83.6 | 51.5 | 97.7 |
| **FaTPS35** | 43 | 39.7 | 98.4 | | 93.4 | 56.6 | 91.4 | 86.9 | 82.9 | 51.1 | 97.9 |
| **FaTPS34** | 42.1 | 39.4 | 93 | 93.4 | | 55.8 | 91.8 | 86.6 | 82.1 | 50.3 | 92.3 |
| **FaTPS33** | 38.1 | 34.9 | 56.5 | 56.6 | 55.8 | | 55.7 | 56.5 | 55 | 41.7 | 56.1 |
| **FaTPS31** | 42.3 | 39.2 | 91.8 | 91.4 | 91.8 | 55.7 | | 90.9 | 86 | 49.7 | 91.6 |
| **FaTPS29** | 41.7 | 38.2 | 87.4 | 86.9 | 86.6 | 56.5 | 90.9 | | 86.4 | 48.6 | 87.3 |
| **FaTPS27** | 42 | 37.7 | 83.6 | 82.9 | 82.1 | 55 | 86 | 86.4 | | 47.4 | 82.6 |
| **FaTPS16** | 60.6 | 57.8 | 51.5 | 51.1 | 50.3 | 41.7 | 49.7 | 48.6 | 47.4 | | 51.6 |
| **FvTPS35** | 42.7 | 39.6 | 97.7 | 97.9 | 92.3 | 56.1 | 91.6 | 87.3 | 82.6 | 51.6 | |

| **TPS Clade a1, 3** | **FaTPS 62** | **FaTPS 59** | **FaTPS 57** | **FaTPS 54** | **FaTPS 49** | **FaTPS 47** | **FaTPS 38** | **FaTPS 26** | **FaTPS 25** | **FaTPS 24** | **FaTPS 15** | **Fxa6Bg10095 6.1** | **Fxa4Bg10262 3.1** | **Fxa4Ag1026 87.1** |
|---|---|---|---|---|---|---|---|---|---|---|---|---|---|---|
| **FaTPS62** | | 87.2 | 97.2 | 88.8 | 57.8 | 57.1 | 51.1 | 57.2 | 55.5 | 29.9 | 51.4 | 42.4 | 36.7 | 31.8 |
| **FaTPS59** | 87.2 | | 87.4 | 79 | 52.7 | 51.9 | 46.5 | 52.1 | 50.9 | 27 | 46.6 | 37.9 | 33.4 | 27.3 |
| **FaTPS57** | 97.2 | 87.4 | | 89.5 | 58.6 | 57.9 | 51.7 | 58.1 | 56.3 | 30.4 | 51.7 | 42.8 | 37.3 | 32 |
| **FaTPS54** | 88.8 | 79 | 89.5 | | 53.8 | 53.1 | 47.4 | 53.4 | 50.8 | 27.8 | 46.9 | 39.6 | 34.2 | 27.5 |
| **FaTPS49** | 57.8 | 52.7 | 58.6 | 53.8 | | 97.3 | 85.1 | 95.7 | 76.4 | 49.1 | 52.9 | 68.3 | 61.1 | 43.9 |
| **FaTPS47** | 57.1 | 51.9 | 57.9 | 53.1 | 97.3 | | 84.4 | 95.7 | 76.4 | 48.9 | 53.1 | 68.4 | 61 | 44.1 |
| **FaTPS38** | 51.1 | 46.5 | 51.7 | 47.4 | 85.1 | 84.4 | | 83.6 | 67.2 | 49.5 | 47.5 | 59.9 | 61.5 | 38.3 |
| **FaTPS26** | 57.2 | 52.1 | 58.1 | 53.4 | 95.7 | 95.7 | 83.6 | | 75.4 | 49.5 | 53.1 | 67.2 | 62.2 | 43.2 |
| **FaTPS25** | 55.5 | 50.9 | 56.3 | 50.8 | 76.4 | 76.4 | 67.2 | 75.4 | | 39.5 | 49.1 | 56.9 | 48.5 | 45.9 |
| **FaTPS24** | 29.9 | 27 | 30.4 | 27.8 | 49.1 | 48.9 | 49.5 | 49.5 | 39.5 | | 28.3 | 36 | 63.2 | 23.1 |
| **FaTPS15** | 51.4 | 46.6 | 51.7 | 46.9 | 52.9 | 53.1 | 47.5 | 53.1 | 49.1 | 28.3 | | 45.1 | 34.1 | 36.6 |
| **Fxa6Bg10095 6.1** | 42.4 | 37.9 | 42.8 | 39.6 | 68.3 | 68.4 | 59.9 | 67.2 | 56.9 | 36 | 45.1 | | 42.9 | 60.6 |
| **Fxa4Bg10262 3.1** | 36.7 | 33.4 | 37.3 | 34.2 | 61.1 | 61 | 61.5 | 62.2 | 48.5 | 63.2 | 34.1 | 42.9 | | 27.5 |
| **Fxa4Ag10268 7.1** | 31.8 | 27.3 | 32 | 27.5 | 43.9 | 44.1 | 38.3 | 43.2 | 45.9 | 23.1 | 36.6 | 60.6 | 27.5 | |

| TPS Clade a1, 4 | FaTPS 64 | FaTPS 61 | FaTPS 58 | FaTPS 56 | FaTPS 55 | FaTPS 32 | FaTPS 30 | FaTPS 28 | FaTPS 20 | FaTPS 2 | Fxa7Dg1 02852.1 | Fxa7Cg1 03019.1 | Fxa7Bg2 03107.2 | Fxa3Ag102 106.1 | Fxa1Ag1 00363.1 | FvrTPS 55 | FcTPS 58 |
|---|---|---|---|---|---|---|---|---|---|---|---|---|---|---|---|---|---|
| **FaTPS64** | | 89.4 | 96.1 | 6.2 | 93.9 | 62.2 | 85.1 | 88.8 | 55.7 | 56.6 | 8.6 | 12.5 | 8.8 | 30.3 | 43.8 | 93.7 | 94.1 |
| **FaTPS61** | 89.4 | | 89.4 | 6.3 | 88.1 | 59.2 | 85.4 | 83.9 | 52.4 | 52.7 | 8.7 | 12.5 | 8.9 | 28.6 | 41.2 | 88.3 | 89.3 |
| **FaTPS58** | 96.1 | 89.4 | | 6.3 | 93.5 | 62.5 | 85.8 | 89.4 | 55.5 | 56.5 | 8.7 | 12.7 | 8.9 | 30.6 | 44 | 94.4 | 94.2 |
| **FaTPS56** | 6.2 | 6.3 | 6.3 | | 6.1 | 5.2 | 5.9 | 6.2 | 5.7 | 5.6 | 69.4 | 46.7 | 68.2 | 2.9 | 4.5 | 6.3 | 6.3 |
| **FaTPS55** | 93.9 | 88.1 | 93.5 | 6.1 | | 64.3 | 86.4 | 89.8 | 56.2 | 57 | 8.6 | 12.5 | 8.8 | 30.2 | 43.6 | 97.7 | 96.3 |
| **FaTPS32** | 62.2 | 59.2 | 62.5 | 5.2 | 64.3 | | 60 | 60.9 | 57.8 | 59.8 | 7.3 | 10.7 | 7.5 | 30.5 | 45.6 | 63.7 | 63.6 |
| **FaTPS30** | 85.1 | 85.4 | 85.8 | 5.9 | 86.4 | 60 | | 83.6 | 53 | 53.5 | 8.2 | 12.1 | 8.4 | 28.6 | 41.9 | 86.3 | 86 |
| **FaTPS28** | 88.8 | 83.9 | 89.4 | 6.2 | 89.8 | 60.9 | 83.6 | | 53.9 | 55.2 | 8.7 | 12.8 | 9 | 29.3 | 42.2 | 89.6 | 89.3 |
| **FaTPS20** | 55.7 | 52.4 | 55.5 | 5.7 | 56.2 | 57.8 | 53 | 53.9 | | 93.8 | 8 | 11.6 | 8.5 | 43.8 | 69 | 56.9 | 56.4 |
| **FaTPS2** | 56.6 | 52.7 | 56.5 | 5.6 | 57 | 59.8 | 53.5 | 55.2 | 93.8 | | 7.9 | 11.3 | 8.2 | 45.2 | 72.8 | 57.5 | 57 |
| **Fxa7Dg102 852.1** | 8.6 | 8.7 | 8.7 | 69.4 | 8.6 | 7.3 | 8.2 | 8.7 | 8 | 7.9 | | 66.1 | 97.3 | 4.1 | 6.3 | 8.8 | 8.8 |
| **Fxa7Cg103 019.1** | 12.5 | 12.5 | 12.7 | 46.7 | 12.5 | 10.7 | 12.1 | 12.8 | 11.6 | 11.3 | 66.1 | | 66.2 | 5.9 | 9 | 12.8 | 12.8 |
| **Fxa7Bg203 107.2** | 8.8 | 8.9 | 8.9 | 68.2 | 8.8 | 7.5 | 8.4 | 9 | 8.5 | 8.2 | 97.3 | 66.2 | | 4.3 | 6.6 | 9 | 9 |
| **Fxa3Ag102 106.1** | 30.3 | 28.6 | 30.6 | 2.9 | 30.2 | 30.5 | 28.6 | 29.3 | 43.8 | 45.2 | 4.1 | 5.9 | 4.3 | | 42.1 | 30.8 | 30.3 |
| **Fxa1Ag100 363.1** | 43.8 | 41.2 | 44 | 4.5 | 43.6 | 45.6 | 41.9 | 42.2 | 69 | 72.8 | 6.3 | 9 | 6.6 | 42.1 | | 44 | 43.7 |
| **FvrTPS55** | 93.7 | 88.3 | 94.4 | 6.3 | 97.7 | 63.7 | 86.3 | 89.6 | 56.9 | 57.5 | 8.8 | 12.8 | 9 | 30.8 | 44 | | 96.1 |
| **FcTPS58** | 94.1 | 89.3 | 94.2 | 6.3 | 96.3 | 63.6 | 86 | 89.3 | 56.4 | 57 | 8.8 | 12.8 | 9 | 30.3 | 43.7 | 96.1 | |

| Zhang et al., FaTPS1 (FaTPS26) | FaTPS49 | FaTPS47 | FaTPS38 | FaTPS26 | FaTPS24 | Fxa6Bg100956.1 | Fxa4Bg102623.1 | Fxa4Ag102687.1 | *FaTPS1 |
|---|---|---|---|---|---|---|---|---|---|
| **FaTPS49** | | 97.3 | 85.2 | 95.7 | 49.3 | 68.1 | 61.4 | 43.9 | 96.3 |
| **FaTPS47** | 97.3 | | 84.6 | 95.7 | 49.2 | 68.3 | 61.3 | 44.1 | 96.1 |
| **FaTPS38** | 85.2 | 84.6 | | 83.8 | 49.7 | 59.9 | 62 | 38.5 | 84.4 |
| **FaTPS26** | 95.7 | 95.7 | 83.8 | | 49.7 | 67 | 62.5 | 43.2 | 97.9 |
| **FaTPS24** | 49.3 | 49.2 | 49.7 | 49.7 | | 36 | 64.6 | 23.2 | 50.2 |
| **Fxa6Bg100956.1** | 68.1 | 68.3 | 59.9 | 67 | 36 | | 42.9 | 61.5 | 67 |
| **Fxa4Bg102623.1** | 61.4 | 61.3 | 62 | 62.5 | 64.6 | 42.9 | | 27.6 | 63.5 |
| **Fxa4Ag102687.1** | 43.9 | 44.1 | 38.5 | 43.2 | 23.2 | 61.5 | 27.6 | | 43.2 |
| ***FaTPS1** | 96.3 | 96.1 | 84.4 | 97.9 | 50.2 | 67 | 63.5 | 43.2 | |

| Mehmood et al. | FaTPS43 | FaTPS42 | FaTPS40 | FaTPS18 | FaTPS17 | FaTPS14 | FaCPS1 | FnTPS6 | FnTPS4 | FnTPS3 | FnTPS2 | FnTPS1 |
|---|---|---|---|---|---|---|---|---|---|---|---|---|
| **FaTPS43** | | 18.9 | 18.7 | 16.8 | 16.7 | 16.8 | 8.9 | 5.1 | 16.4 | 47.8 | 14.1 | 12.2 |
| **FaTPS42** | 18.9 | | 75.5 | 32.8 | 33 | 32.4 | 13.5 | 12 | 31.5 | 34.8 | 20.9 | 49.6 |
| **FaTPS40** | 18.7 | 75.5 | | 31.6 | 31.6 | 31.4 | 13.6 | 11.1 | 30.8 | 32.4 | 19.9 | 43.1 |
| **FaTPS18** | 16.8 | 32.8 | 31.6 | | 83.4 | 79.3 | 15 | 10.5 | 76.7 | 29.7 | 18.4 | 20.9 |
| **FaTPS17** | 16.7 | 33 | 31.6 | 83.4 | | 93.1 | 14.5 | 10.1 | 88.7 | 30.2 | 18.8 | 20.5 |
| **FaTPS14** | 16.8 | 32.4 | 31.4 | 79.3 | 93.1 | | 13.9 | 10.5 | 91.9 | 30.1 | 19.5 | 20.7 |
| **FaCPS1** | 8.9 | 13.5 | 13.6 | 15 | 14.5 | 13.9 | | 4.2 | 13.6 | 13.7 | 10.9 | 11.5 |
| **FnTPS6** | 5.1 | 12 | 11.1 | 10.5 | 10.1 | 10.5 | 4.2 | | 10.2 | 10 | 8 | 7.4 |
| **FnTPS4** | 16.4 | 31.5 | 30.8 | 76.7 | 88.7 | 91.9 | 13.6 | 10.2 | | 30.3 | 19.8 | 20.7 |
| **FnTPS3** | 47.8 | 34.8 | 32.4 | 29.7 | 30.2 | 30.1 | 13.7 | 10 | 30.3 | | 26.3 | 23 |
| **FnTPS2** | 14.1 | 20.9 | 19.9 | 18.4 | 18.8 | 19.5 | 10.9 | 8 | 19.8 | 26.3 | | 26.9 |
| **FnTPS1** | 12.2 | 49.6 | 43.1 | 20.9 | 20.5 | 20.7 | 11.5 | 7.4 | 20.7 | 23 | 26.9 | |

**Supplemental Table 7**. Sample collection and strawberry accession information for this study.

| | Alias | RNA | UC_ID | 2021 Reps | 2021 Harvests | 2022 Reps | 2022 Harvests | Species | Subspecies | Breeding Date | Origin |
|---|---|---|---|---|---|---|---|---|---|---|---|
| **1** | WLSP-08 | | PI551453 | 5 | 2 | 6 | 2 | *F. chiloensis* | lucinda | NA | WA |
| **2** | LCM-10 | | PI551468 | 4 | 2 | 0 | 0 | *F. chiloensis* | lucinda | NA | OR, Pacific coast |
| **3** | Ambato | ** | PI551736 | 4 | 2 | 7 | 3 | *F. chiloensis* | chiloensis | NA | Ecuador |
| **4** | Isle of Lemuy (02A White) | ** | PI552038 | 7 | 3 | 0 | 0 | *F. chiloensis* | NA | NA | Chile |
| **5** | CFRA 688 | | PI612487 | 8 | 3 | 0 | 0 | *F. chiloensis* | pacifica | NA | BC, Pacific coast + Alaska |
| **6** | CFRA 1267 | | PI612488 | 6 | 2 | 4 | 2 | *F. chiloensis* | pacifica | NA | BC, Pacific coast + Alaska |
| **7** | UC04 | ** | PI551498 | 7 | 4 | 0 | 0 | *F. vesca* | NA | NA | CA bred; Asia-Temperate, Europe |
| **8** | UC06 | ** | PI551514 | 9 | 5 | 7 | 3 | *F. vesca* | NA | NA | CA West coast, south/central states/Mexico |
| **9** | Harris Springs | ** | 17X004P001 | 4 | 2 | 0 | 0 | *F. virginiana* | platypetala | NA | CA |
| **10** | LH_18-2 | | PI551876 | 4 | 2 | 0 | 0 | *F. virginiana* | glauca | NA | Wyoming. Pacific Coast and Midwest |
| **11** | Hinesburg | | PI552277 | 6 | 2 | 3 | 3 | *F. virginiana* | virginiana | NA | Vermont |
| **12** | JP_95-9-6 | | PI612320 | 6 | 3 | 6 | 2 | *F. virginiana* | grayana | NA | Georgia, East and central States |
| **13** | NC_96-35-2 | ** | PI612323 | 7 | 3 | 3 | 1 | *F. virginiana* | virginiana | NA | Alabama North and East North America |
| **14** | Fredrk._9 | | PI612493 | 6 | 4 | 4 | 2 | *F. virginiana* | NA | NA | Ontario, Canada |
| **15** | RH_30 | | PI612499 | 3 | 2 | 3 | 1 | *F. virginiana* | virginiana | NA | Minnesota, North and East North America |

| | | | | | | | | | | | |
|---|---|---|---|---|---|---|---|---|---|---|---|
| **16** | KY-17 | | PI616574 | 3 | 2 | 3 | 1 | *F. virginiana* | grayana | NA | Kentucky, East and central States |
| **17** | NC_95-11-1 | | PI616691 | 8 | 4 | 5 | 2 | *F. virginiana* | virginiana | NA | South Carolina North and East North America |
| **18** | NC_95-21-5 | | PI616720 | 5 | 3 | 3 | 1 | *F. virginiana* | grayana | NA | Mississippi, East and central States |
| **19** | NC_96-20-3 | | PI616789 | 8 | 3 | 0 | 0 | *F. virginiana* | virginiana | NA | Alabama North and East North America |
| **20** | NC_96-33-1 | | PI616815 | 3 | 3 | 0 | 0 | *F. virginiana* | virginiana | NA | Alabama North and East North America |
| **21** | NC_96-14-1 | | PI616902 | 9 | 3 | 11 | 4 | *F. virginiana* | virginiana | NA | North Carolina North and East North America |
| **22** | UC11 | | PI551495 | 8 | 3 | 0 | 0 | *F. virginiana* | NA | NA | CA North and East North America |
| **23** | UC12 | | PI551497 | 3 | 2 | 0 | 0 | *F. virginiana* | NA | NA | CA North and East North America |
| **24** | Jucunda | | PI551623 | 5 | 3 | 6 | 4 | *F. x ananassa* | NA | 1854 | England |
| **25** | Weisse Anasa | | PI270464 | 5 | 2 | 4 | 4 | *F. x ananassa* | NA | 1867 | Germany |
| **26** | Sitka | | PI616777 | 14 | 6 | 4 | 2 | *F. x ananassa* | NA | 1905 | Alaska |
| **27** | Howard 17 | | PI551593 | 5 | 2 | 5 | 2 | *F. x ananassa* | NA | 1907 | Massachusetts |
| **28** | Ettersburg 121 | | PI551904 | 8 | 3 | 4 | 2 | *F. x ananassa* | NA | 1907 | CA |
| **29** | Madame Moutot | ** | PI551632 | 7 | 3 | 5 | 2 | *F. x ananassa* | NA | 1910 | France |
| **30** | Kaiser Samling | | PI270471 | 7 | 4 | 4 | 2 | *F. x ananassa* | NA | 1912 | Germany |
| **31** | Aberdeen | | PI551630 | 6 | 3 | 5 | 3 | *F. x ananassa* | NA | 1917 | New Jersey |
| **32** | Blakemore | | PI551421 | 13 | 5 | 9 | 3 | *F. x ananassa* | NA | 1929 | Maryland |

| | | | | | | | | | | | |
|---|---|---|---|---|---|---|---|---|---|---|---|
| **33** | Sparkle | | PI551559 | 8 | 3 | 3 | 1 | *F. x ananassa* | NA | 1942 | New Jersey |
| **34** | Shasta | | 35C035P008 | 8 | 4 | 6 | 2 | *F. x ananassa* | NA | 1945 | UC Berkeley (Davis) |
| **35** | Morioka 17 | | PI551428 | 9 | 4 | 9 | 3 | *F. x ananassa* | NA | 1945 | Japan |
| **36** | Freja | | PI551628 | 5 | 2 | 3 | 1 | *F. x ananassa* | NA | 1948 | Denmark |
| **37** | Albritton | | PI551435 | 9 | 3 | 10 | 4 | *F. x ananassa* | NA | 1951 | North Carolina |
| **38** | Empire | | PI551569 | 15 | 6 | 9 | 3 | *F. x ananassa* | NA | 1951 | New York |
| **39** | Wiltguard | | 52C016P007 | 8 | 4 | 6 | 2 | *F. x ananassa* | NA | 1952 | UC Davis |
| **40** | Direktor Paul Wallbaum | ** | PI551436 | 12 | 4 | 4 | 2 | *F. x ananassa* | NA | 1953 | Germany |
| **41** | Senga Sengana | | PI264680 | 6 | 2 | 6 | 2 | *F. x ananassa* | NA | 1954 | Germany |
| **42** | EarliMiss | ** | PI551862 | 12 | 5 | 15 | 5 | *F. x ananassa* | NA | 1955 | Mississippi |
| **43** | K1_1953 | | PI616778 | 8 | 3 | 12 | 4 | *F. x ananassa* | NA | 1955 | Alaska |
| **44** | Red Gauntlet | | PI551530 | 6 | 2 | 9 | 3 | *F. x ananassa* | NA | 1957 | Scotland |
| **45** | Headliner | ** | PI551652 | 9 | 3 | 9 | 3 | *F. x ananassa* | NA | 1957 | Louisiana |
| **46** | Tioga | | 53C009P002 | 8 | 4 | 10 | 4 | *F. x ananassa* | NA | 1964 | UC Davis |
| **47** | Hood | | PI551502 | 4 | 2 | 7 | 3 | *F. x ananassa* | NA | 1965 | Oregon |
| **48** | Linn | ** | PI551500 | 9 | 3 | 9 | 4 | *F. x ananassa* | NA | 1967 | Oregon |
| **49** | Guardian | | PI551407 | 6 | 2 | 6 | 3 | *F. x ananassa* | NA | 1969 | Maryland |
| **50** | Primella | ** | PI551422 | 9 | 4 | 6 | 2 | *F. x ananassa* | NA | 1969 | Netherlands |
| **51** | Douglas | | 70C003P108 | 6 | 2 | 11 | 5 | *F. x ananassa* | NA | 1970 | UC Davis |
| **52** | Titan | | PI551398 | 5 | 2 | 5 | 2 | *F. x ananassa* | NA | 1971 | North Carolina |

| | | | | | | | | | | | |
|---|---|---|---|---|---|---|---|---|---|---|---|
| **53** | Totem | | PI551501 | 8 | 3 | 6 | 3 | *F. x ananassa* | NA | 1971 | British Colombia, Canada |
| **54** | Tufts | | 63C120P011 | 5 | 2 | 11 | 4 | *F. x ananassa* | NA | 1972 | UC Davis |
| **55** | EarliGlow | | PI551394 | 8 | 3 | 6 | 2 | *F. x ananassa* | NA | 1975 | Maryland |
| **56** | Florida Belle | | PI551396 | 9 | 4 | 6 | 2 | *F. x ananassa* | NA | 1975 | Florida |
| **57** | Tangi | ** | PI551481 | 6 | 3 | 9 | 3 | *F. x ananassa* | NA | 1975 | Louisiana |
| **58** | Kaoling | | PI551537 | 7 | 3 | 8 | 4 | *F. x ananassa* | NA | 1975 | Taiwan |
| **59** | ORUS_4816_ORUSM_173 | | PI551858 | 7 | 3 | 6 | 2 | *F. x ananassa* | NA | 1975 | Oregon |
| **60** | Chandler | | 77C032P103 | 9 | 3 | 8 | 4 | *F. x ananassa* | NA | 1977 | UC Davis |
| **61** | Hecker | | 69C141P101 | 6 | 3 | 12 | 4 | *F. x ananassa* | NA | 1979 | UC Davis |
| **62** | Brighton | | PI551494 | 6 | 3 | 5 | 2 | *F. x ananassa* | NA | 1979 | UC Davis |
| **63** | Elsanta | | PI551579 | 9 | 3 | 6 | 3 | *F. x ananassa* | NA | 1981 | Netherlands |
| **64** | MDUS 5130 | ** | PI551946 | 6 | 2 | 6 | 2 | *F. x ananassa* | NA | 1981 | Maryland |
| **65** | Selva | | 75C071P107 | 5 | 2 | 11 | 4 | *F. x ananassa* | NA | 1982 | UC Davis |
| **66** | Glooscap | | PI551580 | 10 | 4 | 8 | 3 | *F. x ananassa* | NA | 1983 | Nova Scotia |
| **67** | Tillikum | | PI551832 | 7 | 3 | 10 | 4 | *F. x ananassa* | NA | 1983 | Washington |
| **68** | Seascape | | 83C049P001 | 6 | 3 | 6 | 3 | *F. x ananassa* | NA | 1984 | UC Davis |
| **69** | Camarosa | | 88C029P603 | 7 | 3 | 7 | 3 | *F. x ananassa* | NA | 1988 | UC Davis |
| **70** | Beaver Belle | ** | PI551839 | 6 | 2 | 8 | 4 | *F. x ananassa* | NA | 1989 | Canada |
| **71** | Pelican | | PI637960 | 8 | 4 | 8 | 4 | *F. x ananassa* | NA | 1989 | Maryland |
| **72** | Mara des Bois | ** | 17Z001P001 | 6 | 3 | 9 | 3 | *F. x ananassa* | NA | 1991 | France |

| | | | | | | | | | | | |
|---|---|---|---|---|---|---|---|---|---|---|---|
| **73** | Diamante | | 91C248P006 | 7 | 3 | 9 | 4 | *F. x ananassa* | NA | 1991 | UC Davis |
| **74** | Camino Real | | 94C003P011 | 7 | 3 | 5 | 2 | *F. x ananassa* | NA | 1994 | UC Davis |
| **75** | Puget Reliance | | PI664321 | 6 | 2 | 3 | 1 | *F. x ananassa* | NA | 1995 | Washington |
| **76** | Ventana | | 96C042P601 | 9 | 3 | 9 | 5 | *F. x ananassa* | NA | 1996 | UC Davis |
| **77** | Albion | | 97C117P003 | 8 | 4 | 5 | 3 | *F. x ananassa* | NA | 1997 | UC Davis |
| **78** | Monterey | | 01C132P003 | 7 | 3 | 3 | 1 | *F. x ananassa* | NA | 2001 | UC Davis |
| **79** | San Andreas | | 01C139P002 | 9 | 3 | 0 | 0 | *F. x ananassa* | NA | 2001 | UC Davis |
| **80** | Portola | | 01C206P005 | 7 | 3 | 9 | 4 | *F. x ananassa* | NA | 2001 | UC Davis |
| **81** | Petaluma | | 08C020P602 | 8 | 4 | 6 | 3 | *F. x ananassa* | NA | 2008 | UC Davis |
| **82** | Grenada | | 08C055P002 | 6 | 3 | 4 | 2 | *F. x ananassa* | NA | 2008 | UC Davis |
| **83** | Fronteras | | 08C132P608 | 7 | 3 | 4 | 2 | *F. x ananassa* | NA | 2008 | UC Davis |
| **84** | UCD_Warrior | | 08C138P003 | 7 | 3 | 6 | 3 | *F. x ananassa* | NA | 2008 | UC Davis |
| **85** | Cabrillo | | 08C181P001 | 5 | 2 | 8 | 4 | *F. x ananassa* | NA | 2008 | UC Davis |
| **86** | 11-58_FVC | | FVC_11-58 | 9 | 6 | 5 | 4 | *F. x ananassa* | NA | 2010 | Michigan |
| **87** | UCD_Victor | | 11C057P001 | 5 | 2 | 3 | 1 | *F. x ananassa* | NA | 2011 | UC Davis |
| **88** | UCD_Valiant | | 11C103P001 | 7 | 3 | 5 | 2 | *F. x ananassa* | NA | 2011 | UC Davis |
| **89** | UCD_Moxie | | 11C141P001 | 6 | 2 | 8 | 3 | *F. x ananassa* | NA | 2011 | UC Davis |
| **90** | UCD_Royal_Royce (Royal Royce) | ** | 08C123P001 | 6 | 2 | 3 | 1 | *F. x ananassa* | NA | 2008 | UC Davis |
| **91** | UCD_Finn | | 12C112P004 | 7 | 3 | 3 | 1 | *F. x ananassa* | NA | 2012 | UC Davis |
| **92** | UCD_Mojo | | 12C166P002 | 5 | 2 | 9 | 3 | *F. x ananassa* | NA | 2012 | UC Davis |
| **93** | 16C111P068 | | 16C111P068 | 11 | 5 | 13 | 5 | *F. x ananassa* | NA | 2016 | UC Davis |

| | | | | | | | | | | | |
|---|---|---|---|---|---|---|---|---|---|---|---|
| **94** | 17EDN012 | | 12C104P004 | 6 | 3 | 8 | 4 | *F. x ananassa* | NA | 2017 | UC Davis |
| **95** | 17EDN013 | | 12C104P005 | 8 | 3 | 3 | 2 | *F. x ananassa* | NA | 2017 | UC Davis |
| **96** | 17C224P011 | ** | 17C224P011 | 14 | 6 | 10 | 4 | *F. x ananassa* | NA | 2017 | UC Davis |

**Supplemental Table 8.** Compound names, CAS numbers, GC-MS retention times, and compound classifiers for metabolites analyzed in this study.

| Compound | Retention Time | CAS | Compound Type |
|---|---|---|---|
| (-)-Myrtenol | 21.7496 | 515-00-4 | Terpene |
| 1R-(-)-Myrtenal | 18.6072 | 18486-69-6 | Terpene |
| 4,11-Selinadiene | 19.5932 | 28290-20-2 | Terpene |
| $\alpha$-Farnesene | 20.8517 | 502-61-4 | Terpene |
| $\alpha$-Guaiene | 17.9 | 3691-12-1 | Terpene |
| Allo-Ocimene | 13.1479 | 7216-56-0 | Terpene |
| $\alpha$-Muurolene | 20.4108 | 10208-80-7 | Terpene |
| $\alpha$-Pinene | 4.442 | 80-56-8 | Terpene |
| $\alpha$-Terpineol | 19.9054 | 10482-56-1 | Terpene |
| $\beta$-Myrcene | 7.5282 | 123-35-3 | Terpene |
| $\beta$-Phellandrene | 8.6986 | 555-10-2 | Terpene |
| $\beta$-Pinene | 6.1892 | 18172-67-3 | Terpene |
| $\beta$-Selinene | 20.3466 | 17066-67-0 | Terpene |
| Caryophyllene | 18.038 | 87-44-5 | Terpene |
| *Cis*-1-p-Menthanol | 14.1981 | 3901-95-9 | Terpene |
| *Cis*-Linalool Oxide | 14.751 | 5989-33-3 | Terpene |
| 7-epi-α-Selinene | 21.2205 | 106-22-9 | Terpene |
| $\delta$-guaiene | 20.3466 | 3691-11-0 | Terpene |
| D-Limonene | 8.3857 | 5989-27-5 | Terpene |
| E- $\beta$ -Farnesene | 19.3607 | 18794-84-8 | Terpene |
| Eremophilene | 20.1864 | 10219-75-7 | Terpene |
| Geraniol | 22.6234 | 106-24-1 | Terpene |
| Humulene | 19.4489 | 6753-98-6 | Terpene |
| Isopulegol | 17.6773 | 7786-67-6 | Terpene |

| | | | |
|---|---|---|---|
| Linalool | 17.028 | 78-70-6 | Terpene |
| Myrtenyl Acetate | 19.6894 | 1079-01-2 | Terpene |
| Nerolidol | 25.9102 | 7212-44-4 | Terpene |
| Selinane | 20.5391 | 30824-81-8 | Terpene |
| Terpinen-4-ol | 18.1422 | 562-74-3 | Terpene |
| *Trans*-Linalool Oxide | 15.3843 | 34995-77-2 | Terpene |
| *Trans*-Myrtanyl Acatate | 20.9315 | 90934-53-5 | Terpene |
| *Trans*-Pinocarvyl Acetate | 19.04 | 1686-15-3 | Terpene |
| 1-Hexanol | 12.7151 | 111-27-3 | Alcohol |
| 1-Octanol | 17.2844 | 111-87-5 | Alcohol |
| 1-Octen-3-ol | 14.9677 | 3391-86-4 | Alcohol |
| 1-Octen-3-yl Acetate | 16.7392 | 2442-10-6 | MCAAE |
| 1-Pentanol | 10.0695 | 71-41-0 | Alcohol |
| 2,4-Heptadienal E,E | 15.8094 | 4313-03-5 | Aldehyde |
| 2,4-Hexadienal E,E | 13.7011 | 142-83-6 | Aldehyde |
| 2,6-Nonadienal E,Z | 17.8056 | 557-48-2 | Aldehyde |
| 2(3H)-Furanone, dihydro-5-methyl-5-(2-methylpropyl)- | 28.4514 | 10200-21-2 | Phenolic |
| 2-Adamantanol-2-Bromomethyl | 21.6294 | 1000142-34-3 | Misc |
| 2-Butenoic Acid Methyl Ester, E | 6.1496 | 4358-59-2 | SCBAE |
| 2-Butyl Furan | 6.7267 | 4466-24-4 | Furan |
| 2-Decenal, E | 18.9038 | 3913-81-3 | Aldehyde |
| 2-Ethyl Furan | 3.464 | 3208-16-0 | Furan |
| 2-Heptanol | 11.8893 | 543-49-7 | Alcohol |
| 2-Heptanone | 8.1056 | 110-43-0 | Ketone |
| 2-Heptenal,E | 11.7771 | 18829-55-5 | Aldehyde |
| 2-Heptyl Butryrate | 13.8133 | 39026-94-3 | MCBAE |
| 2-Hexen-1-ol,E | 13.9977 | 928-95-0 | Alcohol |

| | | | |
|---|---|---|---|
| 2-Hexenal | 9.0114 | 505-57-7 | Aldehyde |
| 2-Hexenal,E | 8.5625 | 6728-26-3 | Aldehyde |
| 2-Hexenyl Butyrate,E | 15.4326 | 53398-83-7 | MCBAE |
| 2-Methyl-but-2-En-1-yl Acetate | 9.9413 | 33425-30-8 | MCAAE |
| 2-Methylbutyric Acid | 19.4489 | 116-53-0 | Acid |
| 2-Nonanol | 16.4588 | 628-99-9 | Alcohol |
| 2-Nonanol Acetate | 15.144 | 14936-66-4 | MCAAE |
| 2-Octenal, E | 14.3584 | 2548-87-0 | Aldehyde |
| 2-Penten-1-ol Acetate, Z | 6.2137 | 42125-10-0 | SCAAE |
| 2-Pentenal, E | 6.6626 | 1576-87-0 | Aldehyde |
| 2-Pentyl Furan | 9.3802 | 3777-69-3 | Furan |
| 3-Ethyl-4-Methyl pentan-1-ol | 8.851 | 38514-13-5 | Alcohol |
| 3-Formylamino-Succinimide | 24.1865 | 99417-77-3 | Misc |
| 3-Hexen-1-ol | 12.9555 | 928-96-1 | Alcohol |
| 3-Hexen-1-ol Acetate, E | 11.5687 | 3681-82-1 | MCAAE |
| 3-Hexen-1-ol Acetate, Z | 11.697 | 3681-71-8 | MCAAE |
| 3-Hexen-1-ol, E | 13.44 | 928-97-2 | Alcohol |
| 3-Hexenal | 6.8791 | 4440-65-7 | Aldehyde |
| 3-Hexenyl Butyrate | 15.144 | 16491-36-4 | MCBAE |
| 3-Pentanol | 6.3417 | 584-02-1 | Alcohol |
| 4,4-Dimethylpent-2-enal | 8.0334 | 22597-46-2 | Aldehyde |
| 4-Hexenoic Acid | 24.2824 | 1577-20-4 | Acid |
| 4-Methyl-5-nonanone | 21.461 | 35900-26-6 | Ketone |
| 4-Methyl-Pentanoic Acid | 21.926 | 646-07-1 | Acid |
| 5-Methyl-Hexanoic Acid | 23.6494 | 628-46-6 | Acid |
| Benzaldehyde | 16.2984 | 100-52-7 | Phenolic |
| Benzyl Acetate | 20.5391 | 140-11-4 | Phenolic |
| Benzyl Butyrate | 23.0163 | 103-37-7 | Phenolic |
| Butanethioic Acid-S-Methyl Ester | 8.53 | 2432-51-1 | Sulfur |

| | | | |
|---|---|---|---|
| Butanoic Acid | 18.6072 | 107-92-6 | Acid |
| Butyl Acetate | 5.6203 | 123-86-4 | SCAAE |
| Butyl Butyrate | 9.0676 | 109-21-7 | SCBAE |
| Butyl Hexanoate | 13.9977 | 626-82-4 | SCHAE |
| Cinnamyl Acetate | 27.5451 | 103-54-8 | Phenolic |
| *Cis*-3-Hexenyl Hexanoate | 19.2405 | 31501-11-8 | MCHAE |
| Decanal | 15.9938 | 112-31-2 | Aldehyde |
| Decyl Acetate | 19.6894 | 112-17-4 | MCAAE |
| Decyl Butyrate | 22.2866 | 5454-09-1 | MCBAE |
| Decyl Isobutyrate | 22.1184 | 5454-22-8 | MCAAE |
| Ethyl-(Z)-Cinnamate | 27.2565 | 4610-69-9 | Phenolic |
| Ethyl-2-Methyl Butyrate | 5.0673 | 7452-79-1 | SCBAE |
| Ethyl Benzoate | 19.3607 | 93-89-0 | Phenolic |
| Ethyl Butyrate | 4.74 | 105-54-4 | SCBAE |
| Ethyl Caprate | 18.8156 | 110-38-3 | SCDAE |
| Ethylene Glycol Dibutyrate | 20.9315 | 105-72-6 | Misc |
| Ethyl Hexanoate | 9.5085 | 123-66-0 | SCHAE |
| Ethyl Isobutyrate | 20.1864 | 97-62-1 | SCBAE |
| Ethyl Isocaproate | 8.4258 | 25415-67-2 | SCVAE |
| Ethyl Isovalerate | 5.4 | 108-64-5 | SCBAE |
| Ethyl Octanoate | 14.5508 | 106-32-1 | SCOAE |
| Ethyl Pentanoate | 6.8791 | 539-82-2 | SCVAE |
| Ethyl Propinoate | 3.5122 | 105-37-3 | SCPAE |
| Eugenol | 27.7539 | 97-53-0 | Phenolic |
| Furaneol | 25.7574 | 3658-77-3 | Furan |
| Heptanal | 8.1377 | 111-71-7 | Aldehyde |
| Heptanoic Acid | 24.4511 | 111-14-8 | Acid |
| Heptyl Acetate | 13.1479 | 112-06-1 | MCAAE |
| Hexanal | 5.5964 | 66-25-1 | Aldehyde |

| | | | |
|---|---|---|---|
| Hexane-3-Ethyl-4-Methyl | 10.1097 | 3074-77-9 | Misc |
| Hexanoic Acid | 22.7597 | 142-62-1 | Acid |
| Hexyl-2-Methylbutyrate | 14.4624 | 10032-15-2 | MCBAE |
| Hexyl Acetate | 10.6628 | 142-92-7 | MCAAE |
| Hexyl Butyrate | 14.1981 | 626-82-4 | MCBAE |
| Hexyl Hexanoate | 18.3506 | 6378-65-0 | MCHAE |
| Hexyl Isovalerate | 14.8635 | 10032-13-0 | MCBAE |
| Hexyl Propanoate | 12.2742 | 2445-76-3 | MCPAE |
| Hydrocinnamyl Isobutyrate | 24.2824 | 103-58-2 | Phenolic |
| Isoamyl Acetate | 6.5018 | 123-92-2 | SCAAE |
| Isoamyl Alcohol | 8.8991 | 123-51-3 | Alcohol |
| Isoamyl Butyrate | 10.3662 | 106-27-4 | SCBAE |
| Isobutyl Butyrate | 7.4643 | 539-90-2 | SCBAE |
| Isobutyl Hexanoate | 12.6429 | 105-79-3 | SCHAE |
| Isobutyric Acid | 17.4127 | 79-31-2 | Acid |
| Isopentyl Hexanoate | 15.088 | 2198-61-0 | SCHAE |
| Isopentyl Isovalerate | 11.1679 | 659-70-1 | SCBAE |
| Isopropyl Butyrate | 4.8429 | 638-11-9 | SCBAE |
| Isovaleric Acid | 19.5932 | 503-74-2 | Acid |
| Mesifurane | 17.9418 | 4077-47-8 | Furan |
| Methyl-2-Hydroxybutyrate | 13.2681 | 29674-47-3 | SCBAE |
| Methyl-2-Methyl Butyrate | 4.2895 | 868-57-5 | SCBAE |
| Methyl-3-Methylthio-Propionate | 16.4588 | 13532-18-8 | Sulfur |
| Methyl Anthranilate | 28.5476 | 134-20-3 | Phenolic |
| Methyl Butyrate | 3.9 | 623-42-7 | SCBAE |
| Methyl Cinnamate | 26.3912 | 103-26-4 | Phenolic |
| Methyl Decanoate | 17.9418 | 110-42-9 | SCDAE |
| Methyl Hexanoate | 8.1773 | 106-70-7 | SCHAE |
| Methyl Isobutyl Ketone | 4.2095 | 108-10-1 | Ketone |

| | | | |
|---|---|---|---|
| Methyl Isohexanoate | 7.0074 | 2177-83-5 | SCVAE |
| Methyl Isovalerate | 4.426 | 556-24-1 | SCBAE |
| Methyl Nonanoate | 15.8896 | 1731-84-6 | SCNAE |
| Methyl Octanoate | 13.4204 | 111-11-5 | SCOAE |
| Methyl Salicylate | 21.3568 | 119-36-8 | Phenolic |
| Methyl Thiolacetate | 4.9631 | 1534-08-3 | Sulfur |
| Methyl Valerate | 5.6846 | 624-24-8 | SCVAE |
| Nonanal | 13.4766 | 124-19-6 | Aldehyde |
| Nonanoic Acid | 27.7539 | 112-05-0 | Acid |
| Octan-8-o-l2,5-Diaza-2,5-Dimethyl | 10.8312 | 26439-05-4 | Misc |
| Octanal | 10.9754 | 124-13-0 | Aldehyde |
| Octanoic Acid | 26.1827 | 124-07-2 | Acid |
| Octyl-2-Methyl Butyrate | 18.7034 | 29811-50-5 | MCBAE |
| Octyl Acetate | 15.4326 | 112-14-1 | MCAAE |
| Octyl Butyrate | 18.4228 | 110-39-4 | MCBAE |
| Octyl Hexanoate | 22.0383 | 4887-30-3 | MCHAE |
| Octyl Isovalerate | 19.04 | 7786-58-5 | MCBAE |
| Octyl Propionate | 16.8676 | 142-60-9 | MCPAE |
| Pent-1-en-3-ol | 7.6005 | 616-25-1 | Alcohol |
| Pentanoic acid, 2,2,4-trimethyl-3-carboxyisopropyl, isobutyl ester | 23.2086 | 1000140-77-5 | Misc |
| Pentanoic acid, 2-Methyl Anhydride | 11.9534 | 63169-61-9 | MCHAE |
| Pentanoic Acid | 20.7152 | 109-52-4 | Acid |
| Pentyl Acetate | 7.849 | 628-63-7 | SCAAE |
| Phenethyl Acetate | 22.1184 | 103-45-7 | Phenolic |
| S-Methyl Isovalerate | 9.2038 | 23747-45-7 | Sulfur |
| Styrene | 10.0376 | 100-42-5 | Misc |
| *Trans*-2-Hexenyl Acetate,E | 12.2261 | 2497-18-9 | MCAAE |
| *Trans*-2-Hexenyl Isovalerate | 15.9938 | 68698-59-9 | MCBAE |

| | | | |
|---|---|---|---|
| $\gamma$-Decalactone | 27.4413 | 706-14-9 | Lactone |
| $\gamma$-Dodecalactone | 29.8943 | 2305-05-7 | Lactone |
| $\gamma$-Octalactone | 23.8099 | 104-50-7 | Lactone |
| $\gamma$-Pentalactone | 21.926 | 108-29-2 | Lactone |
| $\gamma$-Undecalactone Standard | 28.8201 | 104-67-6 | Standard |
| Z-Pent-2-enyl Butyrate | 13.0117 | 42125-13-3 | SCVAE |

**Supplemental Table 9:** Reagents and authentic standards used in this study.

| Standard | CAS # | Curve R2 | Range | Equation |
|---|---|---|---|---|
| Linalool | 78-70-6 | .991 | 2 ng/mL - 1110 ng/mL | 4345.18x |
| (1R)-(-)-Myrtenal | 18486-69-6 | .988 | 0.1 ng/mL - 5 ng/mL | 59902.65x |
| (-)-Myrtenol | 19894-97-4 | .957 | 1 ng/mL - 400 ng/mL | 2532.21x |
| (-)-Myrtenyl Acetate | 36203-31-3 | .98 | 0.2 ng/mL - 150 ng/mL | 7014.49x |
| Nerolidol | 7212-44-4 | .971 | 0.2 ng/mL - 120 ng/mL | 1423.46x |
| α-Pinene | 80-56-8 | .943 | 0.04 ng/mL - 2 ng/mL | 1622687x |
| (-)-β-Pinene | 18172-67-3 | .967 | 0.04 ng/mL - 2 ng/mL | 1438854x |
| Terpineol | 8000-41-7 | .99 | 1 ng/mL - 200 ng/mL | 397.25x |
| $\gamma$-Undecalactone Internal Standard | 104-67-6 | .987 | 0.2 ng/mL - 120 ng/mL | 753.85x |
| Citronellol | 106-22-9 | NA | NA | NA |
| Farnesene | 502-61-4 | NA | NA | NA |
| Farnesyl Pyrophosphate | 13059-04-3 | NA | NA | NA |
| Geranyl Pyrophosphate | 763-10-0 | NA | NA | NA |
| Borneol | 464-45-9 | NA | NA | NA |
| Caryophyllene | 87-44-5 | NA | NA | NA |
| Germacrene D | 23986-74-5 | NA | NA | NA |
| (R)-(+)- Limonene | 5989-27-5 | NA | NA | NA |
| (S)-(-)-Limonene | 5989-54-8 | NA | NA | NA |
| Isobutyl Benzene Internal Standard | 538-93-2 | NA | NA | NA |
| Linalool oxide | 60047-17-8 | NA | NA | NA |
| β-Elemene | 515-13-9 | NA | NA | NA |
| Humulene | 6753-98-6 | NA | NA | NA |
